## Supplementary material for "Age-associated changes in lineage composition of the enteric nervous system regulate gut health and disease": All supplemental data

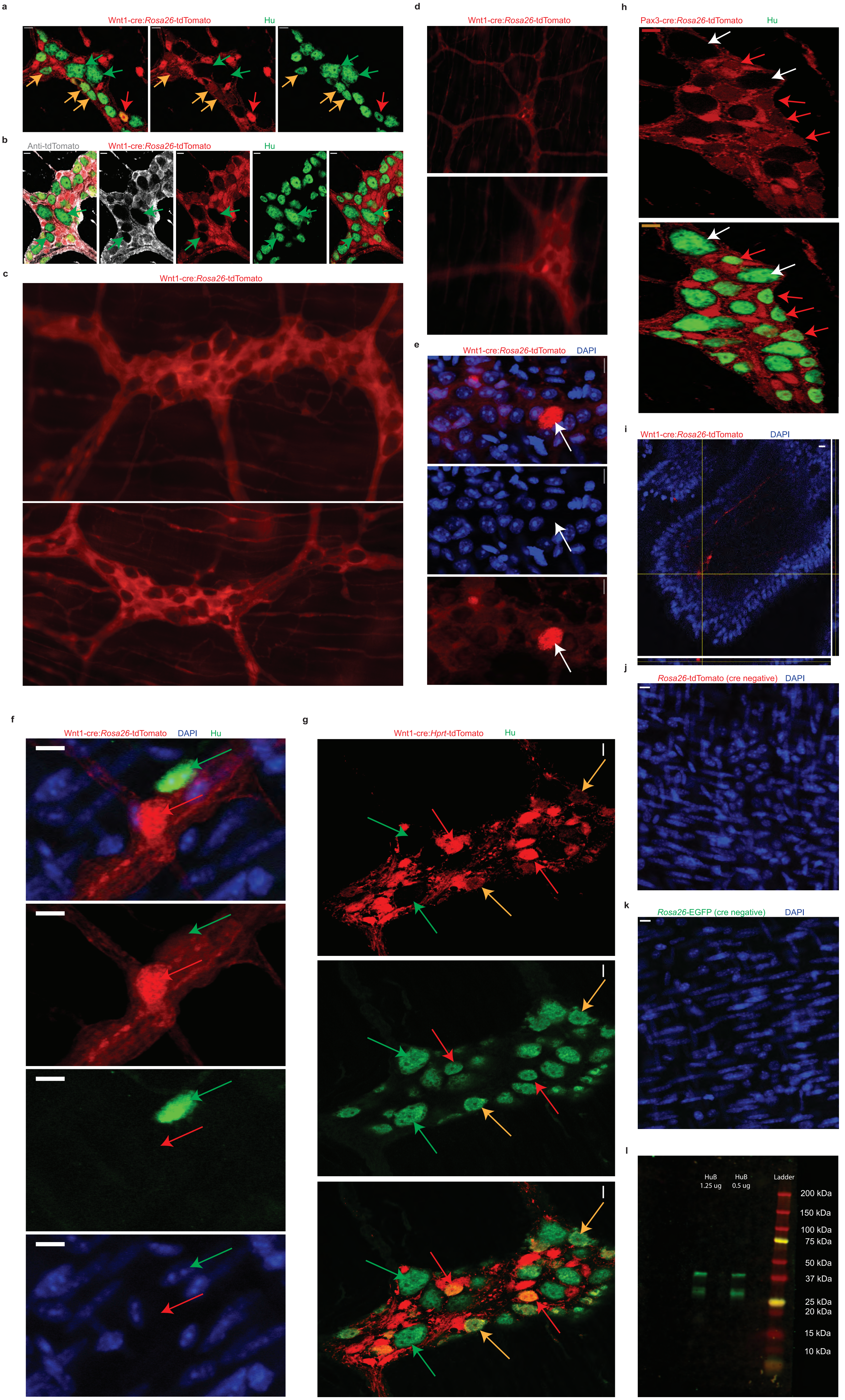


Supplementary Figure 1

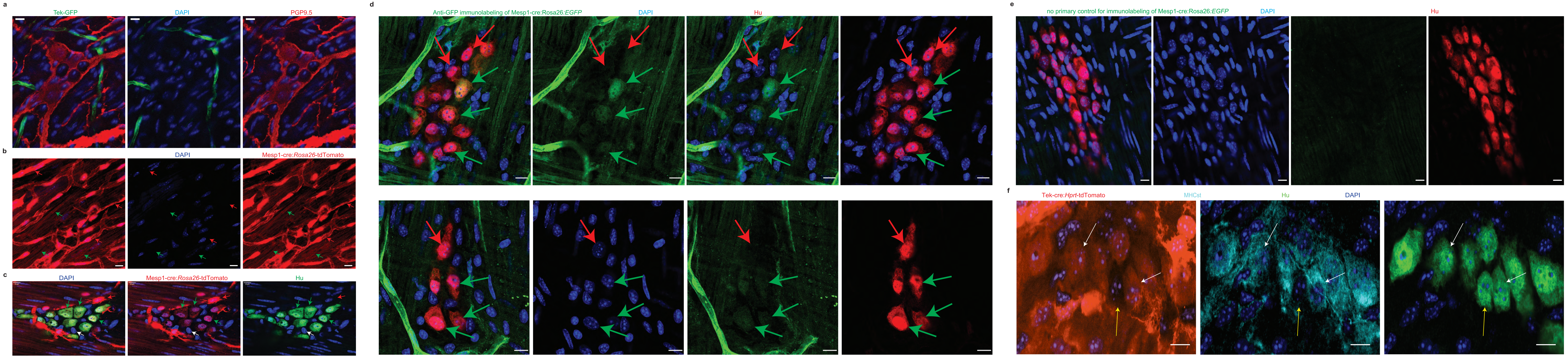


Supplementary Figure 2

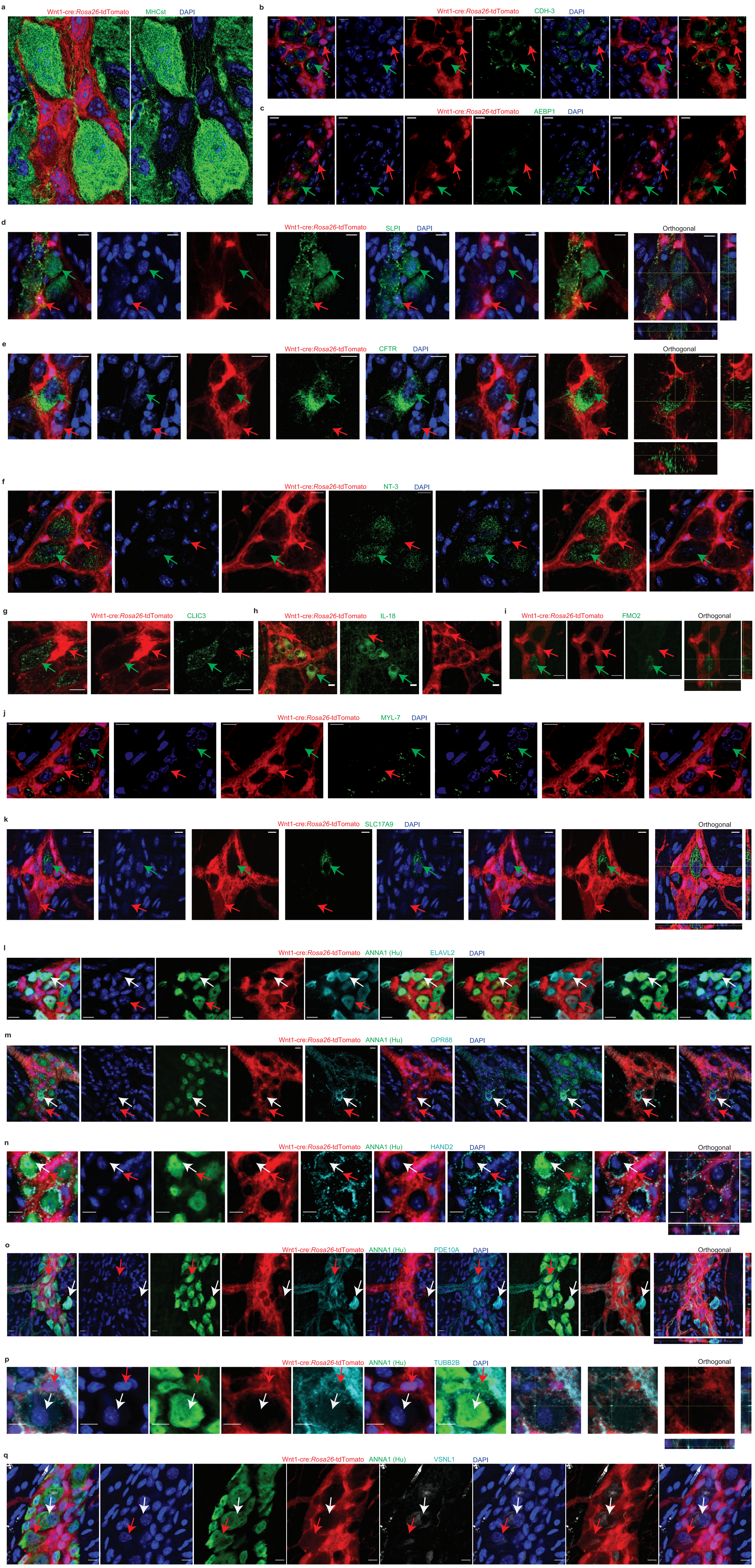


Supplementary Figure 3

a

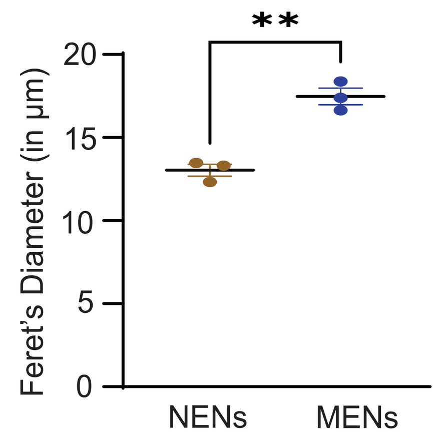


b

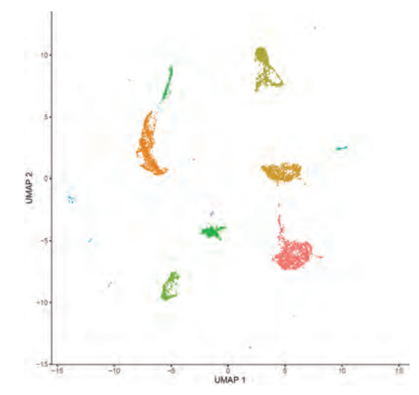

c

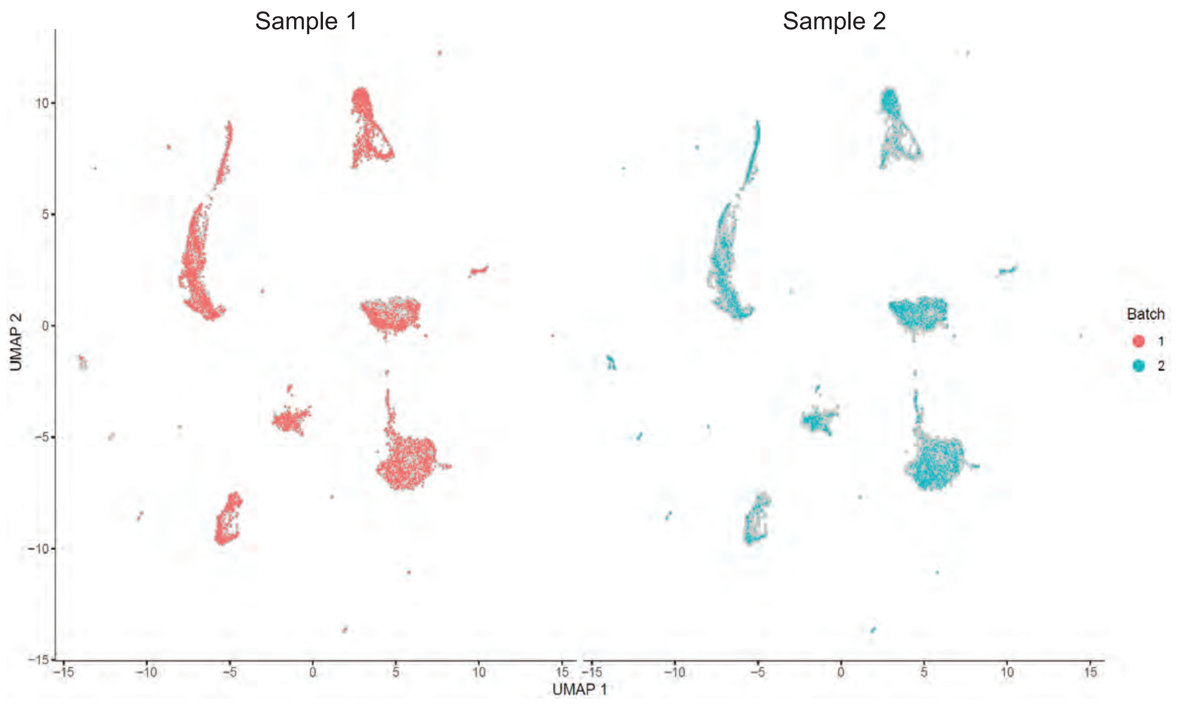


e

| Pattern 16 | | Pattern 27 | | Pattern 32 | | Pattern 41 | |
|---|---|---|---|---|---|---|---|
| | Weight | | Weight | | Weight | | Weight |
| Fth1 | 31.65 | C3 | 34.22 | S100a6 | 33.71 | Slpi | 33.71 |
| Crip1 | 28.76 | Aebp1 | 16.00 | Tmsb10 | 17.71 | Igfbp6 | 17.71 |
| Dcn | 25.00 | Crip1 | 15.56 | Rarres2 | 11.60 | Dcn | 11.60 |
| Krt19 | 18.27 | Rarres2 | 15.31 | Igfbp6 | 9.41 | Rarres2 | 9.41 |
| Fmo2 | 17.60 | Igfbp6 | 14.51 | S100a11 | 5.69 | Gpx3 | 5.69 |
| Vim | 17.11 | Msln | 13.76 | S100a10 | 4.46 | mt.Cytb | 4.46 |
| Tmsb4x | 16.67 | Dcn | 12.58 | Anxa2 | 3.93 | Timp2 | 3.93 |
| Gas1 | 15.81 | Clu | 7.84 | Gpx3 | 3.45 | mt.Nd1 | 3.45 |
| Timp2 | 15.54 | Pcolce | 7.47 | Cox6c | 2.35 | Tmsb4x | 2.35 |
| Upk3b | 15.36 | Efemp1 | 6.84 | Vim | 2.22 | Rpl39 | 2.22 |
| ... | ... | ... | ... | ... | ... | ... | ... |

f

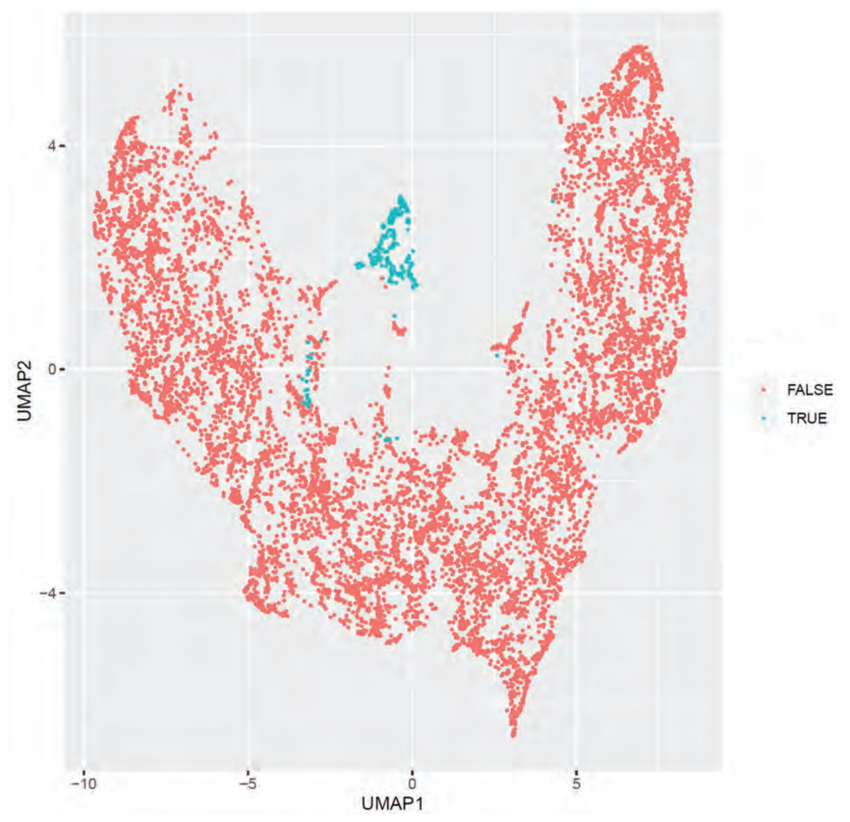


g

| Gene name | FDR | summary.logFC |
|---|---|---|
| Slpi | 1.39E-101 | 3.236117851 |
| Dcn | 4.36E-58 | 2.638143129 |
| C3 | 5.84E-51 | 2.422087511 |
| Igfbp6 | 4.36E-39 | 1.805873578 |
| Rarres2 | 1.26E-34 | 1.633843396 |
| Sparc | 1.20E-28 | 1.403776417 |
| Upk3b | 1.31E-27 | 1.340156034 |
| Fmo2 | 7.84E-27 | 1.434445935 |
| Ahnak | 2.50E-26 | 1.23139421 |
| Aebp1 | 8.95E-25 | 1.113210119 |
| Abi1 | 5.04E-24 | 1.243538323 |
| Lrrn4 | 5.50E-24 | 1.037438294 |
| Serping1 | 1.03E-23 | 1.022387499 |
| Ifitm3 | 1.17E-23 | 1.042651541 |
| Crip1 | 1.06E-22 | 1.258975664 |
| Gas1 | 8.49E-21 | 1.019994737 |
| Col3a1 | 6.90E-20 | 1.137131018 |
| Msln | 5.29E-19 | 0.753319531 |
| Mylk | 7.32E-19 | 1.113912447 |
| Igfbp7 | 1.16E-18 | 0.895910629 |
| Col1a1 | 1.87E-18 | 0.815599605 |
| Serpinh1 | 2.27E-18 | 0.821736594 |
| Col1a2 | 5.35E-18 | 0.852757795 |
| Krt19 | 3.07E-17 | 0.849525515 |
| Acta2 | 4.86E-17 | 0.996287023 |
| Gas6 | 4.90E-17 | 0.926085712 |
| Ezr | 1.08E-16 | 0.79191645 |
| Vim | 1.16E-16 | 0.990418893 |
| Sdc4 | 1.94E-16 | 0.698274093 |
| Rspo1 | 7.27E-16 | 0.719466458 |
| Actg2 | 4.86E-15 | 0.966294003 |
| Aldh1a2 | 5.62E-15 | 0.722238389 |
| C1ra | 2.16E-13 | 0.577407504 |
| Ptgis | 4.81E-13 | 0.585542774 |
| Pcolce | 5.69E-13 | 0.554316803 |
| Rnase4 | 6.98E-13 | 0.602302869 |
| Il6st | 7.29E-13 | 0.735840079 |
| Gpc3 | 7.29E-13 | 0.663125765 |
| Sema3c | 7.46E-13 | 0.813464997 |
| Cav1 | 1.00E-12 | 0.782988385 |
| Fgl2 | 1.01E-08 | 0.382322613 |
| Ahnak2 | 1.37E-08 | 0.384653085 |
| Mgp | 1.37E-08 | 0.451261281 |
| Dab2 | 1.78E-08 | 0.543382519 |
| Cald1 | 1.78E-08 | 0.629648566 |
| Ifi27l2a | 1.78E-08 | 0.443266751 |
| Actb | 2.35E-08 | 0.494312297 |
| Pbxip1 | 3.06E-08 | 0.451131524 |
| Creb3l1 | 4.11E-08 | 0.34684942 |
| Sulf1 | 4.11E-08 | 0.773533306 |
| Wwc1 | 4.64E-08 | 0.446842216 |
| Cfh | 4.85E-08 | 0.638220537 |
| Clic4 | 6.00E-08 | 0.42964409 |
| Tgm2 | 7.13E-08 | 0.398995609 |
| Ccnd2 | 7.29E-08 | 0.402770991 |
| Arhgap18 | 1.31E-07 | 0.600447138 |
| Cobll1 | 1.41E-07 | 0.635331276 |
| Cybrd1 | 1.68E-07 | 0.323950559 |
| Flna | 1.82E-07 | 0.668207013 |

d

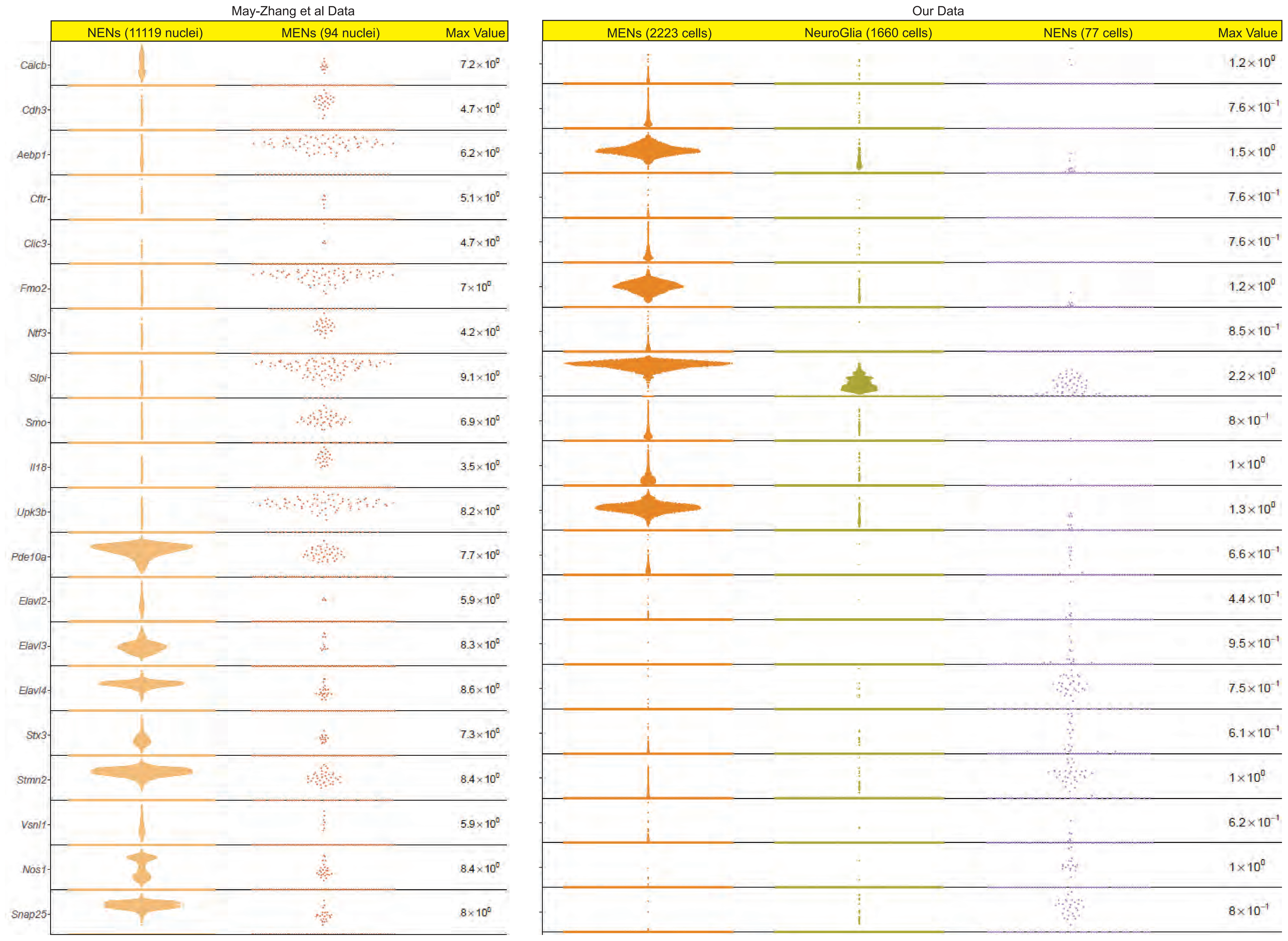


h

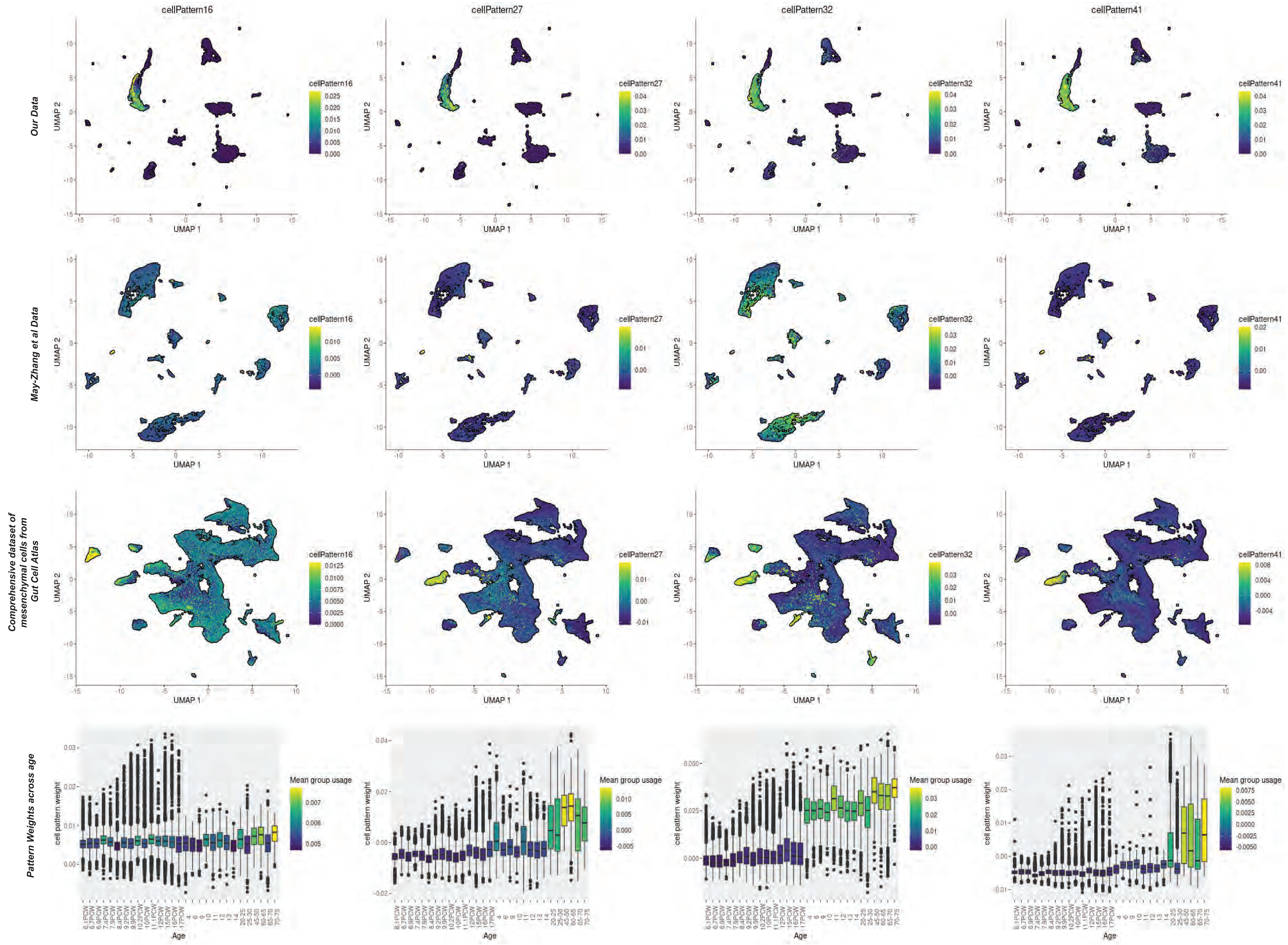


Supplementary Figure 4

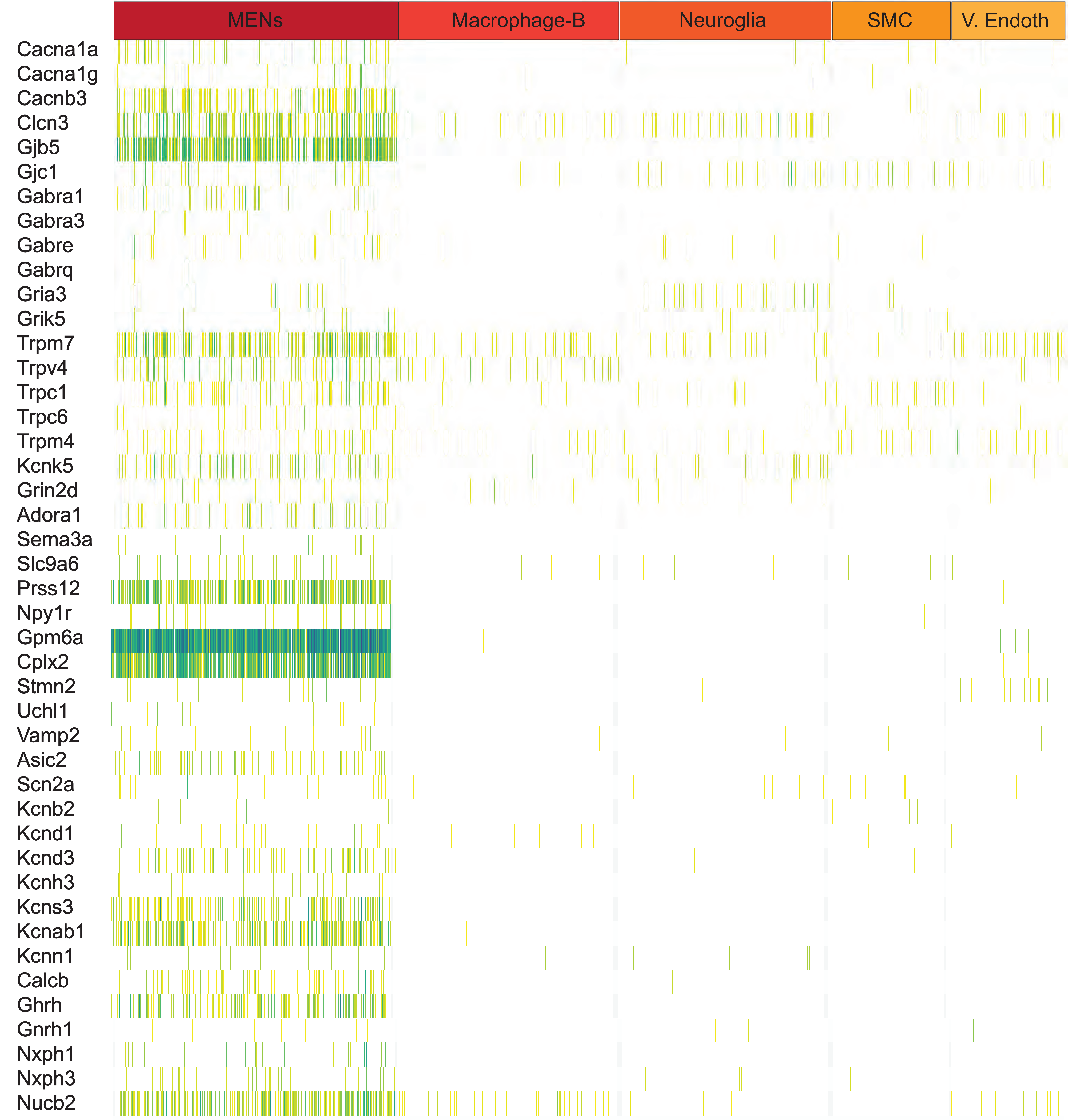
MENs
Macrophage-B
Neuroglia
SMC
V. Endoth
Cacna1a
Cacna1g
Cacnb3
Clcn3
Gjb5
Gjc1
Gabra1
Gabra3
Gabre
Gabrq
Gria3
Grik5
Trpm7
Trpv4
Trpc1
Trpc6
Trpm4
Kcnk5
Grin2d
Adora1
Sema3a
Slc9a6
Prss12
Npy1r
Gpm6a
Cplx2
Stmn2
Uchl1
Vamp2
Asic2
Scn2a
Kcnb2
Kcnd1
Kcnd3
Kcnh3
Kcns3
Kcnab1
Kcnn1
Calcb
Ghrh
Gnrh1
Nxph1
Nxph3
Nucb2

| | Gene | Protein name | Neuronal association |
|---|---|---|---|
| 1 | Cacna1a | CaV2.1 alpha 1 Encodes neuronal P/Q-type calcium channel | Exocytotic Calcium channel in CNS neurons[1], mutations associated with trigeminal neuralgia[2], congenital ataxia[3], and hemiplegic migrane[4] |
| 2 | Cacna1g | low-voltage-activated Ca(v)3.1 T-type calcium channel | Expressed highly in Purkinje Cells and by deep cerebellar nuclei neurons of the cerebellum[5]. Mutations associated with Spinocerebellar ataxia 42[5,6]. Involved in generation of spike-and-wave discharges in thalamocortical relay neurons[7] |
| 3 | Cacnb3 | Voltage-dependent L-type calcium channel subunit beta-3 | Accessory beta 3 subunits are required for normal functions of N-type calcium channels in neurons[8] |
| 4 | Clcn3 | Chloride Voltage-Gated Channel 3 CLC3 | Expressed in synaptic vesicles, loss of this protein causes selective degeneration of hippocampus. Contributes to vesicular acidification by providing an electrical shunt for efficient proton pumping by the $H^+$-ATPase[9] |
| 5 | Gjb5 | Connexin 31.1 | Expressed by electrically coupled striatal output neurons[10] |
| 6 | Gjc1 | Connexin 45 | Expressed by neurons of the adult spinal dorsal horn[11] |
| 7 | Gabra1 | GABA A receptor subunit 1 | Expressed by myenteric neurons[12] |
| 8 | Gabra3 | GABA A receptor subunit 3 | Expressed by myenteric neurons[12] |
| 9 | Gabre | GABA A receptor subunit epsilon | Expressed by locus coeruleus neurons[13] |
| 10 | Gabrq | GABA A receptor subunit theta | Expressed by monoaminergic neurons of the brain[14] |
| 11 | Gria3 | Glutamate Ionotropic receptor AMPA type subunit 3 (GluA3) | Involved in excitatory synaptic transmission in Glutamate responsive neurons[15] |
| 12 | Grik5 | Glutamate Ionotropic Receptor Kainate Type Subunit 5 (GluK5) | Contribute to excitatory post-synaptic transmission in Glutamate responsive neurons[16] |
| 13 | Trpm7 | Transient receptor potential melastatin 7 | Regulates plasticity of synaptic strength[17] |
| 14 | Trpv4 | Transient Receptor Potential Cation Channel Subfamily V Member 4 | Expressed by myenteric neurons[18] |
| 15 | Trpc1 | Transient Receptor Potential Cation Channel Subfamily C Member 1 | Expressed by pyramidal neurons and by a subset of somatostatin-expressing cortical interneurons[19], and mutations are associated with pyloric stenosis[20] |
| 16 | Trpc6 | Transient Receptor Potential Cation Channel Subfamily C Member 6 | Expressed by dentate granule cells[21] |
| 17 | Trpm4 | Transient Receptor Potential Cation Channel Subfamily M Member 4 | Regulates large subthreshold membrane potential oscillations that underlie tonic action |

| | | | |
|---|---|---|---|
| | | | potential discharge in locus coeruleus and suprachiasmatic nucleus[22] |
| 18 | Kcnk5 | TASK2 | Regulates pH sensitivity in chemosensory neurons of the retrotrapezoid nucleus[23] |
| 19 | Grin2d | Glutamate Ionotropic Receptor NMDA Type Subunit 2D (GluN2D) | NMDA receptor that contributes to Glutamatergic signaling and mutations are associated with developmental and epileptic encephalopathy[24] |
| 20 | Adora1 | Adenosine receptor 1 | Neuronal nucleoside signaling[25], expressed in myenteric ganglia[26] |
| 21 | Sema3a | Semaphorin 3A | Important for plexin-mediated control of neurite outgrowth, and is expressed by adult enteric neurons[27] |
| 22 | Slc9a6 | Sodium Hydrogen exchanger 6 (NHE6) | Mutations associated with Christianson syndrome that show predominantly neuronal and behavioral symptoms including postnatal microcephaly, intellectual disability, epilepsy, non-verbal status, autistic features[28] |
| 23 | Prss12 | Neurotrypsin | Synaptic serine protease that regulates maintenance of excitatory synapses in the CNS[29] |
| 24 | Npy1r | Neuropeptide Y receptor type 1 | Neuronally expressed gene that responds to NPY, PYY and other hormones[30] |
| 25 | Gpm6a | Neuronal membrane glycoprotein M6-A | Involved in neuron development and synapse formation and plasticity, and was also recently proposed as a gene-target in various neuropsychiatric disorders[31] |
| 26 | Cplx2 | Complexin 2 | Regulates neurotransmitter release from presynaptic terminals of mature neurons[32] |
| 27 | Stmn2 | Stathmin 2 | Neuronal growth associated gene and is expressed in motor neurons[33] |
| 28 | Uchl1 | Ubiquitin C-Terminal Hydrolase L1 (PGP9.5) | Pan-neuronal marker in the myenteric plexus[34] |
| 29 | Vamp2 | Synaptobrevin 2 | Mediates Tetanus and Botulinum toxin-sensitive vesicular release of neurotransmitters[35] |
| 30 | Asic2 | Amiloride-sensitive cation channel 1, Neuronal (ACCN1), Brain sodium channel 1 (BNaC1) | This acid sensing ion channel functions as a neuronal proton receptor and is expressed by mechanosensitive neurons[36] |
| 31 | Scn2a | NaV1.2 | Plays an important role in the initiation and conductance of an action potential[37] |
| 32 | Kcnb2 | Potassium Voltage-Gated Channel Subfamily B Member 2 (Kv2.2) | Controls delayed rectifier currents and is expressed by pyramidal cortical neurons[38] |
| 33 | Kcnd1 | Potassium Voltage-Gated Channel Subfamily D Member 1 (Kv4.1) | Regulates low frequency firing of Dentate Granule Neurons[39] |

| | | | |
|---|---|---|---|
| 34 | Kcnd3 | Potassium Voltage-Gated Channel Subfamily D Member 3 (Kv4.3) | Encodes for an A-type K+ channel, which is expressed in cerebellar neurons[40] |
| 35 | Kcnh3 | Potassium Voltage-Gated Channel Subfamily H Member 3 (Kv12.2) | Regulates excitability in hippocampal neurons[41] |
| 36 | Kcns3 | Potassium Voltage-Gated Channel Modifier Subfamily S Member 3 (Kv9.3) | Kv9.3 subunits are present in voltage-gated potassium channels that contribute to the precise detection of coincident excitatory synaptic inputs to parvalbumin neurons of the prefrontal cortex[42] |
| 37 | Kcnab1 | Potassium Voltage-Gated Channel Subfamily A Regulatory Beta Subunit 1 (Kvb1.3) | Expressed by axons of certain hippocampal neurons[43] |
| 38 | Kcnn1 | Potassium intermediate/small conductance calcium-activated channel, subfamily N, member 1 (SK1) | Regulates neuronal excitability by contributing to the slow component of synaptic after-hyperpolarization current[44] |
| 39 | Calcb | CGRPb | Encodes ENS-expressed isoform of CGRP[45] |
| 40 | Ghrh | Growth hormone releasing hormone | Expressed by hypothalamic neurons[46] |
| 41 | Gnrh1 | Gonadotropin releasing hormone | Expressed by hypothalamic and myenteric neurons[47] |
| 42 | Nxph1 | Neurexophilin 1 | Alpha Neurexin ligand, which is expressed by subpopulation of inhibitory neurons in the brain, and supports GABA-B receptor based neurotransmission[48] |
| 43 | Nxph3 | Neurexophilin 3 | Highly localized in cortical and cerebellar regions and is functionally important for sensorimotor gating and motor coordination[49] |
| 44 | Nucb2 | Nucleobindin 2/Nesfatin-1 | Expressed by NPY and CART-expressing neurons of the brainstem[50]; expressed by myenteric neurons[51] |

**References:**


1 Dunlap, K., Luebke, J. I. & Turner, T. J. Exocytotic Ca2+ channels in mammalian central neurons. *Trends in Neurosciences* **18**, 89-98, doi:https://doi.org/10.1016/0166-2236(95)80030-6 (1995).

2 Gambeta, E., Gandini, M. A., Souza, I. A., Ferron, L. & Zamponi, G. W. A CACNA1A variant associated with trigeminal neuralgia alters the gating of Cav2.1 channels. *Molecular Brain* **14**, 4, doi:10.1186/s13041-020-00725-y (2021).

3 Izquierdo-Serra, M., Fernández-Fernández, J. M. & Serrano, M. Rare CACNA1A mutations leading to congenital ataxia. *Pflügers Archiv-European Journal of Physiology* **472**, 791-809 (2020).

4 Hullugundi, S., Ansuini, A., Ferrari, M., van den Maagdenberg, A. & Nistri, A. A hyperexcitability phenotype in mouse trigeminal sensory neurons expressing the R192Q Cacna1a missense mutation of familial hemiplegic migraine type-1. *Neuroscience* **266**, 244-254 (2014).

5 Coutelier, M. *et al.* A Recurrent Mutation in CACNA1G Alters Cav3.1 T-Type Calcium-Channel Conduction and Causes Autosomal-Dominant Cerebellar Ataxia. *Am J Hum Genet* **97**, 726-737, doi:10.1016/j.ajhg.2015.09.007 (2015).

6 Chemin, J. *et al.* De novo mutation screening in childhood-onset cerebellar atrophy identifies gain-of-function mutations in the CACNA1G calcium channel gene. *Brain* **141**, 1998-2013, doi:10.1093/brain/awy145 (2018).

7 Kim, D. *et al.* Lack of the burst firing of thalamocortical relay neurons and resistance to absence seizures in mice lacking alpha(1G) T-type Ca(2+) channels. *Neuron* **31**, 35-45, doi:10.1016/s0896-6273(01)00343-9 (2001).

8 Murakami, M. *et al.* Pain perception in mice lacking the beta3 subunit of voltage-activated calcium channels. *J Biol Chem* **277**, 40342-40351, doi:10.1074/jbc.M203425200 (2002).

9 Stobrawa, S. M. *et al.* Disruption of ClC-3, a Chloride Channel Expressed on Synaptic Vesicles, Leads to a Loss of the Hippocampus. *Neuron* **29**, 185-196, doi:https://doi.org/10.1016/S0896-6273(01)00189-1 (2001).

10 Venance, L., Glowinski, J. & Giaume, C. Electrical and chemical transmission between striatal GABAergic output neurones in rat brain slices. *J Physiol* **559**, 215-230, doi:10.1113/jphysiol.2004.065672 (2004).

11 Chapman, R. J. *et al.* Localization of neurones expressing the gap junction protein Connexin45 within the adult spinal dorsal horn: a study using Cx45-eGFP reporter mice. *Brain Struct Funct* **218**, 751-765, doi:10.1007/s00429-012-0426-1 (2013).

12 Seifi, M. *et al.* Molecular and functional diversity of GABA-A receptors in the enteric nervous system of the mouse colon. *J Neurosci* **34**, 10361-10378, doi:10.1523/JNEUROSCI.0441-14.2014 (2014).

13 Belujon, P. *et al.* Inhibitory Transmission in Locus Coeruleus Neurons Expressing GABAA Receptor Epsilon Subunit Has a Number of Unique Properties. *Journal of Neurophysiology* **102**, 2312-2325, doi:10.1152/jn.00227.2009 (2009).

14 Bonnert, T. P. *et al.* theta, a novel gamma-aminobutyric acid type A receptor subunit. *Proc Natl Acad Sci U S A* **96**, 9891-9896, doi:10.1073/pnas.96.17.9891 (1999).

15 Italia, M., Ferrari, E., Di Luca, M. & Gardoni, F. GluA3-containing AMPA receptors: From physiology to synaptic dysfunction in brain disorders. *Neurobiology of Disease* **161**, 105539, doi:https://doi.org/10.1016/j.nbd.2021.105539 (2021).

16 Breustedt, J. & Schmitz, D. Assessing the Role of GLU<sub>K5</sub> and GLU<sub>K6</sub> at Hippocampal Mossy Fiber Synapses. *The Journal of Neuroscience* **24**, 10093-10098, doi:10.1523/jneurosci.3078-04.2004 (2004).

17 Jiang, Z.-J. *et al.* TRPM7 is critical for short-term synaptic depression by regulating synaptic vesicle endocytosis. *eLife* **10**, e66709, doi:10.7554/eLife.66709 (2021).

18 Fichna, J. *et al.* Transient receptor potential vanilloid 4 inhibits mouse colonic motility by activating NO-dependent enteric neurotransmission. *Journal of Molecular Medicine* **93**, 1297-1309, doi:10.1007/s00109-015-1336-5 (2015).

19 Martinez-Galan, J. R., Verdejo, A. & Caminos, E. TRPC1 Channels Are Expressed in Pyramidal Neurons and in a Subset of Somatostatin Interneurons in the Rat Neocortex. *Front Neuroanat* **12**, 15, doi:10.3389/fnana.2018.00015 (2018).

20 Everett, K. V. *et al.* Infantile hypertrophic pyloric stenosis: evaluation of three positional candidate genes, TRPC1, TRPC5 and TRPC6, by association analysis and re-sequencing. *Hum Genet* **126**, 819-831, doi:10.1007/s00439-009-0735-5 (2009).

21 Kim, J. E., Park, H., Choi, S. H., Kong, M. J. & Kang, T. C. TRPC6-Mediated ERK1/2 Activation Increases Dentate Granule Cell Resistance to Status Epilepticus Via Regulating Lon Protease-1 Expression and Mitochondrial Dynamics. *Cells* **8**, doi:10.3390/cells8111376 (2019).

22 Li, K., Shi, Y., Gonye, E. C. & Bayliss, D. A. TRPM4 Contributes to Subthreshold Membrane Potential Oscillations in Multiple Mouse Pacemaker Neurons. *eneuro* **8**, ENEURO.0212-0221.2021, doi:10.1523/eneuro.0212-21.2021 (2021).

23 Wang, S. *et al.* TASK-2 Channels Contribute to pH Sensitivity of Retrotrapezoid Nucleus Chemoreceptor Neurons. *The Journal of Neuroscience* **33**, 16033-16044, doi:10.1523/jneurosci.2451-13.2013 (2013).

24 Camp, C. R. & Yuan, H. GRIN2D/GluN2D NMDA receptor: Unique features and its contribution to pediatric developmental and epileptic encephalopathy. *Eur J Paediatr Neurol* **24**, 89-99, doi:10.1016/j.ejpn.2019.12.007 (2020).

25 Province, H. S. *et al.* Activation of neuronal adenosine A1 receptors causes hypothermia through central and peripheral mechanisms. *PLOS ONE* **15**, e0243986, doi:10.1371/journal.pone.0243986 (2020).

26 Li, T., Yuan, P.-Q., Million, M., Larauche, M. & Taché, Y. Transcriptomic profiling of the enteric nervous system (ENS) in the pig colon: regional heterogeneity and implication in physiological functions. *The FASEB Journal* **34**, 1-1, doi:https://doi.org/10.1096/fasebj.2020.34.s1.09672 (2020).

27 Gonzales, J. *et al.* Semaphorin 3A controls enteric neuron connectivity and is inversely associated with synapsin 1 expression in Hirschsprung disease. *Scientific Reports* **10**, 15119, doi:10.1038/s41598-020-71865-3 (2020).

28 Ouyang, Q. *et al.* Christianson syndrome protein NHE6 modulates TrkB endosomal signaling required for neuronal circuit development. *Neuron* **80**, 97-112, doi:10.1016/j.neuron.2013.07.043 (2013).

29 Stephan, A. *et al.* Neurotrypsin cleaves agrin locally at the synapse. *The FASEB Journal* **22**, 1861-1873, doi:https://doi.org/10.1096/fj.07-100008 (2008).

30 Bertocchi, I., Oberto, A., Longo, A., Palanza, P. & Eva, C. Conditional inactivation of Npy1r gene in mice induces sex-related differences of metabolic and behavioral functions. *Hormones and Behavior* **125**, 104824, doi:https://doi.org/10.1016/j.yhbeh.2020.104824 (2020).

31 Leon, A., Aparicio, G. I. & Scorticati, C. Neuronal Glycoprotein M6a: An Emerging Molecule in Chemical Synapse Formation and Dysfunction. *Front Synaptic Neurosci* **13**, 661681, doi:10.3389/fnsyn.2021.661681 (2021).

32 Ono, S. *et al.* Regulatory roles of complexins in neurotransmitter release from mature presynaptic nerve terminals. *Eur J Neurosci* **10**, 2143-2152, doi:10.1046/j.1460-9568.1998.00225.x (1998).

33 Gagliardi, D. *et al.* Stathmins and Motor Neuron Diseases: Pathophysiology and Therapeutic Targets. *Biomedicines* **10**, 711 (2022).

34 Krammer, H. J., Karahan, S. T., Rumpel, E., Klinger, M. & Kuhnel, W. Immunohistochemical visualization of the enteric nervous system using antibodies against

protein gene product (PGP) 9.5. *Ann Anat* **175**, 321-325, doi:10.1016/s0940-9602(11)80029-4 (1993).
35 Schiavo, G. *et al.* Tetanus and botulinum-B neurotoxins block neurotransmitter release by proteolytic cleavage of synaptobrevin. *Nature* **359**, 832-835, doi:10.1038/359832a0 (1992).
36 Cabo, R. *et al.* ASIC2 is present in human mechanosensory neurons of the dorsal root ganglia and in mechanoreceptors of the glabrous skin. *Histochem Cell Biol* **143**, 267-276, doi:10.1007/s00418-014-1278-y (2015).
37 Ogiwara, I. *et al.* Nav1.2 haplodeficiency in excitatory neurons causes absence-like seizures in mice. *Communications Biology* **1**, 96, doi:10.1038/s42003-018-0099-2 (2018).
38 Kihira, Y., Hermanstyne, T. O. & Misonou, H. Formation of heteromeric Kv2 channels in mammalian brain neurons. *J Biol Chem* **285**, 15048-15055, doi:10.1074/jbc.M109.074260 (2010).
39 Kim, K.-R. *et al.* Kv4.1, a Key Ion Channel For Low Frequency Firing of Dentate Granule Cells, Is Crucial for Pattern Separation. *The Journal of Neuroscience* **40**, 2200-2214, doi:10.1523/jneurosci.1541-19.2020 (2020).
40 Hsu, Y. H., Huang, H. Y. & Tsaur, M. L. Contrasting expression of Kv4.3, an A-type K+ channel, in migrating Purkinje cells and other post-migratory cerebellar neurons. *Eur J Neurosci* **18**, 601-612, doi:10.1046/j.1460-9568.2003.02786.x (2003).
41 Zhang, X. *et al.* Deletion of the potassium channel Kv12.2 causes hippocampal hyperexcitability and epilepsy. *Nat Neurosci* **13**, 1056-1058, doi:10.1038/nn.2610 (2010).
42 Georgiev, D. *et al.* Lower gene expression for KCNS3 potassium channel subunit in parvalbumin-containing neurons in the prefrontal cortex in schizophrenia. *Am J Psychiatry* **171**, 62-71, doi:10.1176/appi.ajp.2013.13040468 (2014).
43 Schulte, U. *et al.* The epilepsy-linked Lgi1 protein assembles into presynaptic Kv1 channels and inhibits inactivation by Kvbeta1. *Neuron* **49**, 697-706, doi:10.1016/j.neuron.2006.01.033 (2006).
44 Adelman, J. P., Maylie, J. & Sah, P. Small-Conductance Ca2+-Activated K+ Channels: Form and Function. *Annual Review of Physiology* **74**, 245-269, doi:10.1146/annurev-physiol-020911-153336 (2012).
45 STERNINI, C. Enteric and Visceral Afferent CGRP Neurons. *Annals of the New York Academy of Sciences* **657**, 170-186, doi:https://doi.org/10.1111/j.1749-6632.1992.tb22766.x (1992).
46 Balthasar, N. *et al.* Growth hormone-releasing hormone (GHRH) neurons in GHRH-enhanced green fluorescent protein transgenic mice: a ventral hypothalamic network. *Endocrinology* **144**, 2728-2740, doi:10.1210/en.2003-0006 (2003).
47 Ohlsson, B. Gonadotropin-Releasing Hormone and Its Role in the Enteric Nervous System. *Front Endocrinol (Lausanne)* **8**, 110, doi:10.3389/fendo.2017.00110 (2017).
48 Born, G. *et al.* Modulation of synaptic function through the α-neurexin–specific ligand neurexophilin-1. *Proceedings of the National Academy of Sciences* **111**, E1274-E1283, doi:doi:10.1073/pnas.1312112111 (2014).
49 Beglopoulos, V. *et al.* Neurexophilin 3 Is Highly Localized in Cortical and Cerebellar Regions and Is Functionally Important for Sensorimotor Gating and Motor Coordination. *Molecular and Cellular Biology* **25**, 7278-7288, doi:doi:10.1128/MCB.25.16.7278-7288.2005 (2005).
50 Psilopanagioti, A., Makrygianni, M., Nikou, S., Logotheti, S. & Papadaki, H. Nucleobindin 2/nesfatin-1 expression and colocalisation with neuropeptide Y and cocaine- and amphetamine-regulated transcript in the human brainstem. *J Neuroendocrinol* **32**, e12899, doi:10.1111/jne.12899 (2020).

51 Varricchio, E. *et al.* Expression and immunohistochemical detection of nesfatin-1 in the gastrointestinal tract of Casertana pig. *Acta Histochem* **116**, 583-587, doi:10.1016/j.acthis.2013.11.006 (2014).

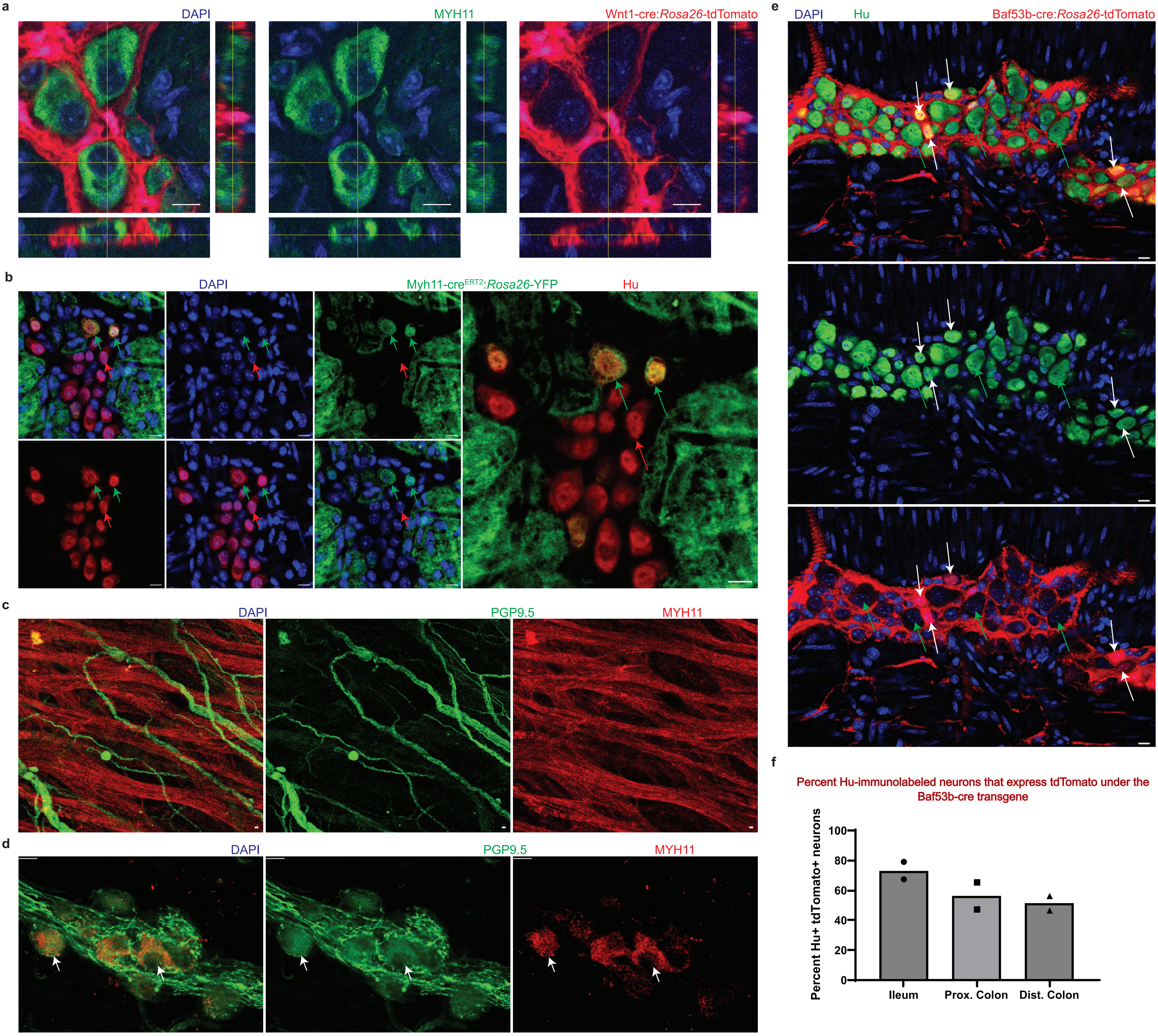


Supplementary Figure 6

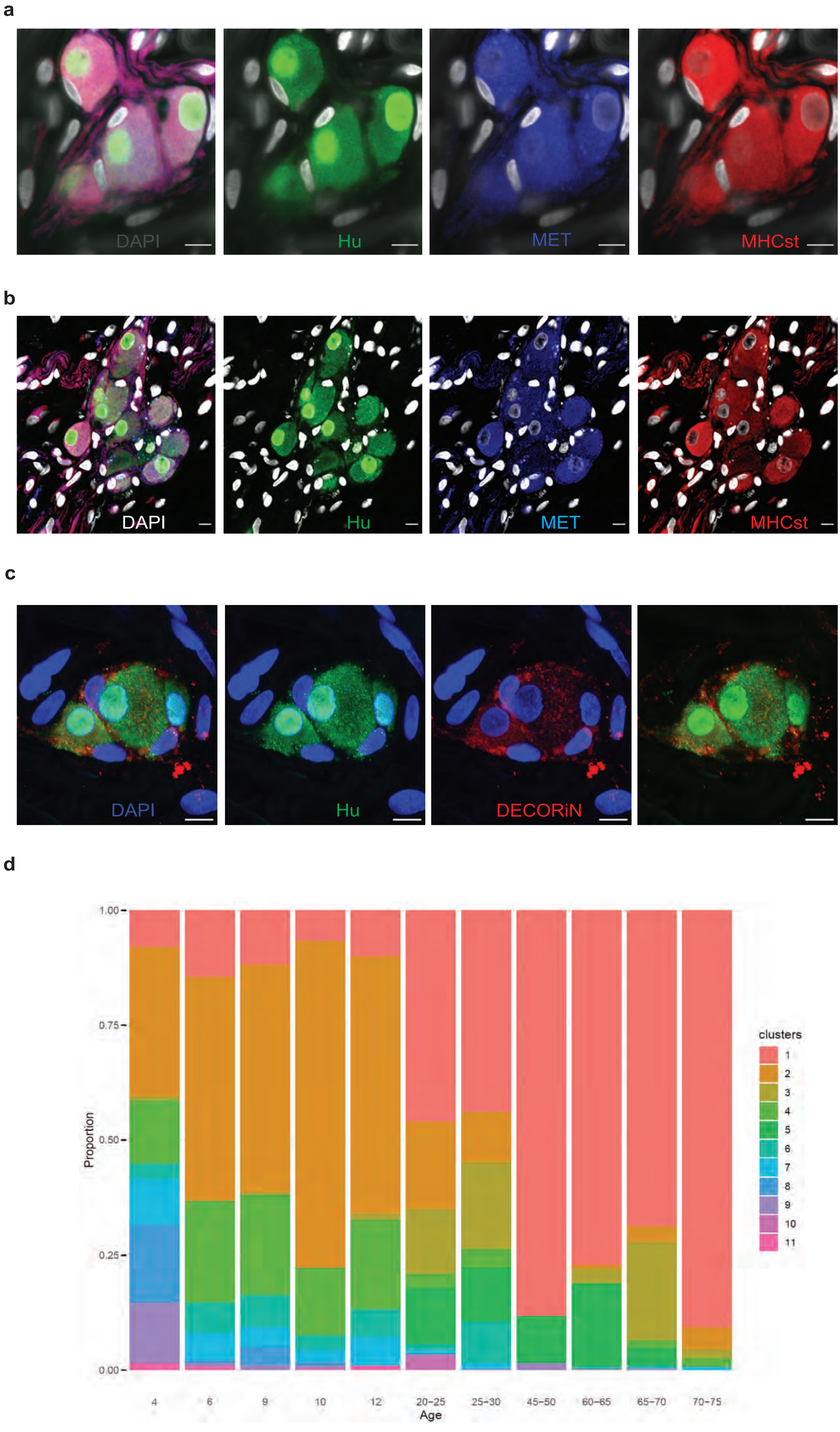


Supplementary Figure 7

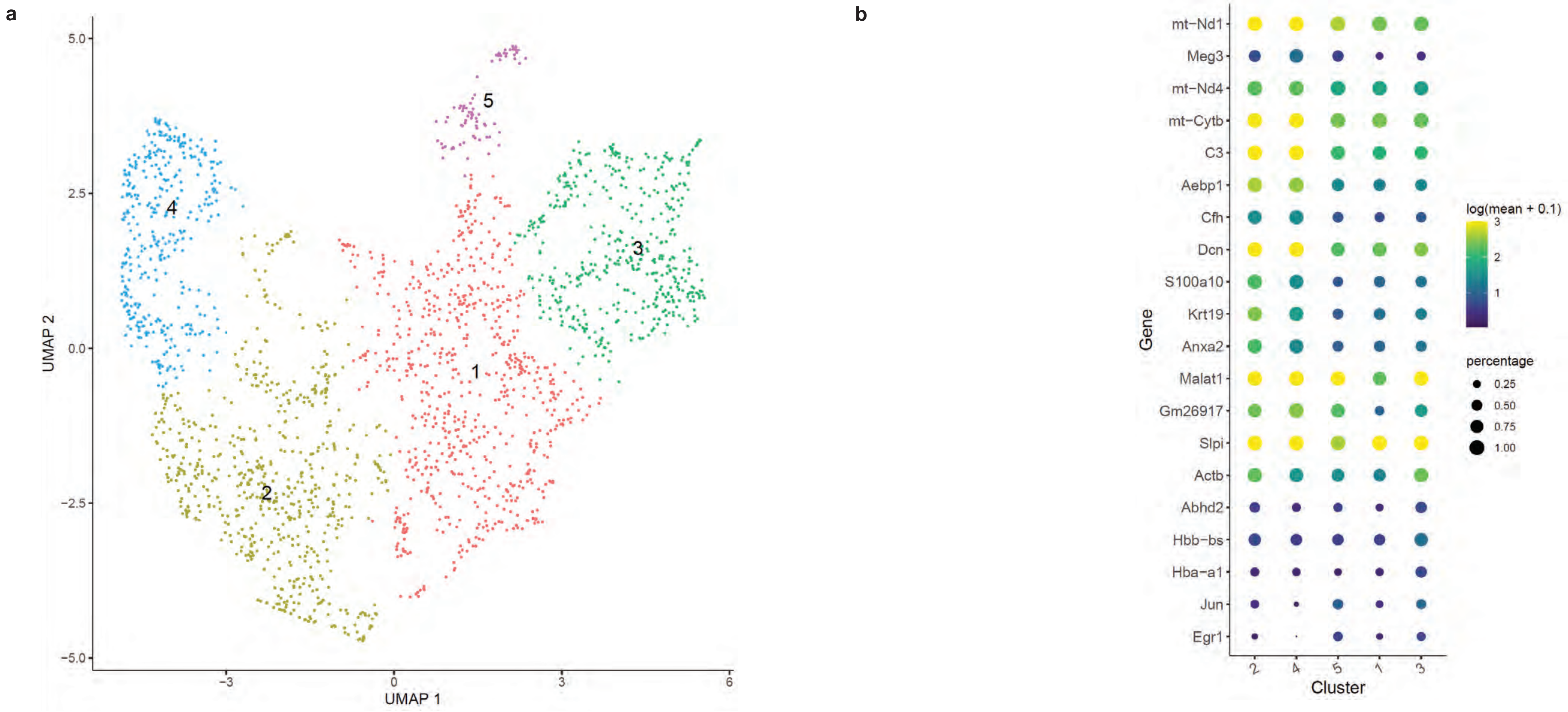


Supplementary Figure 8

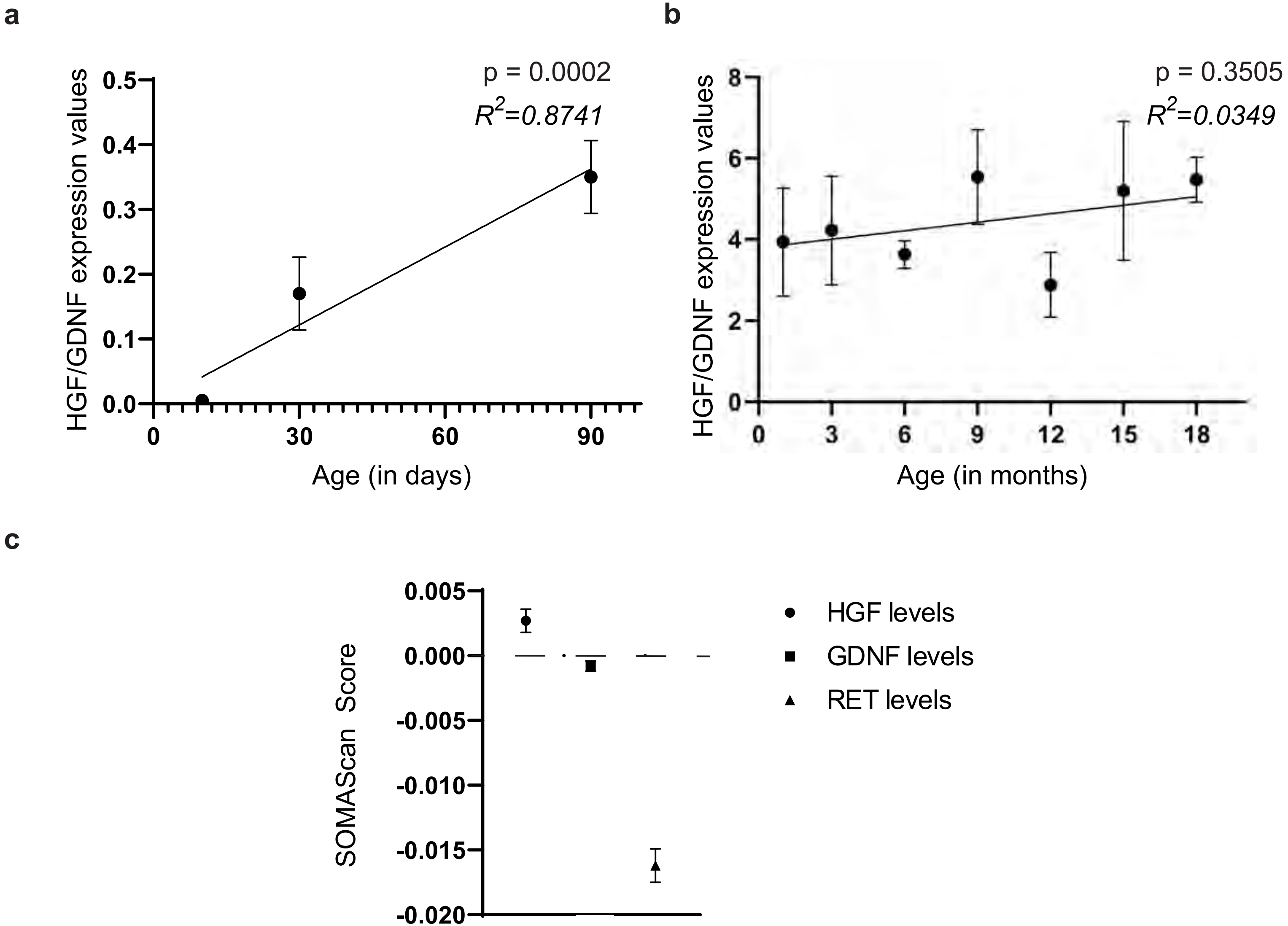


Supplementary Figure 9

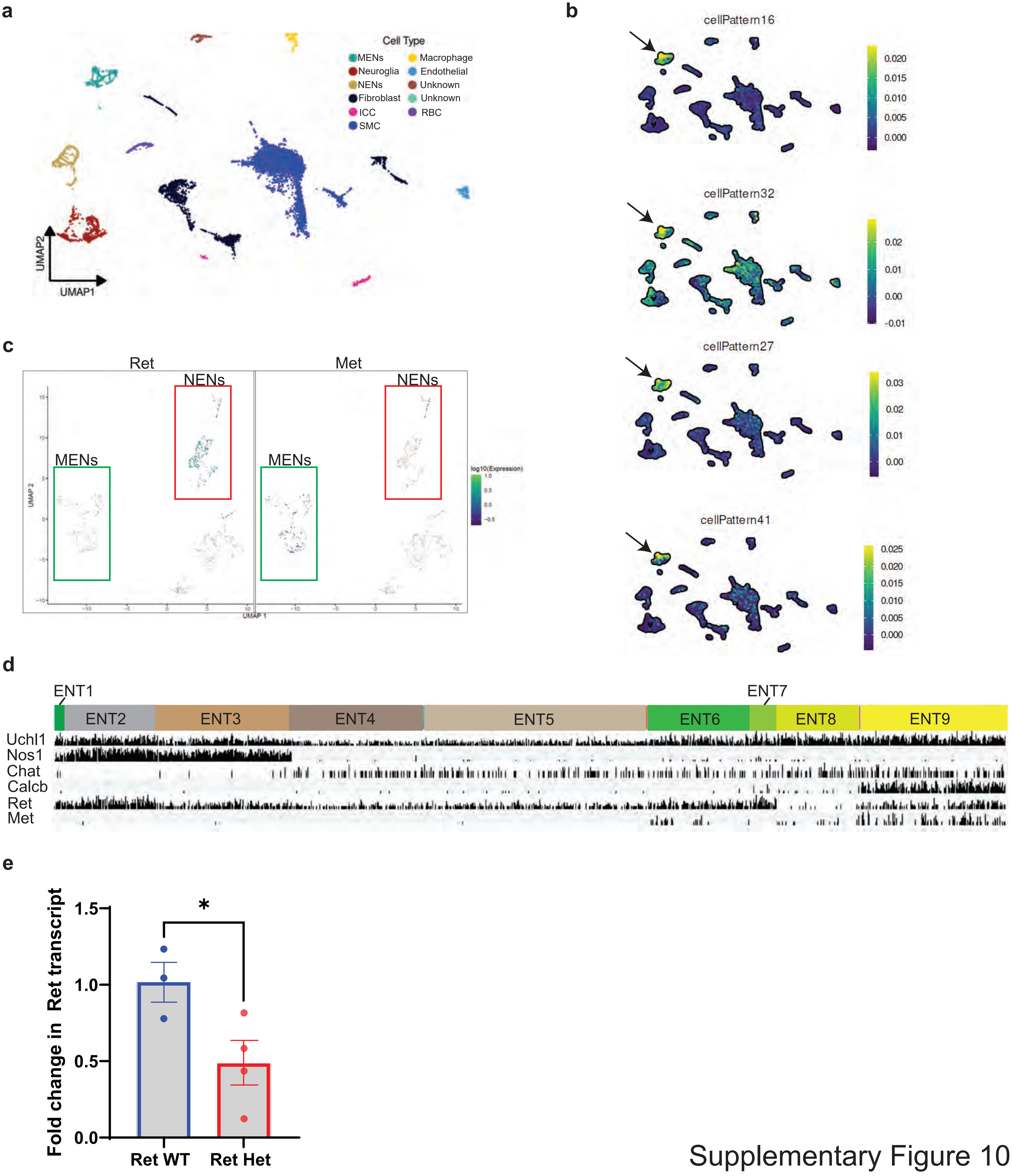


Supplementary Figure 10

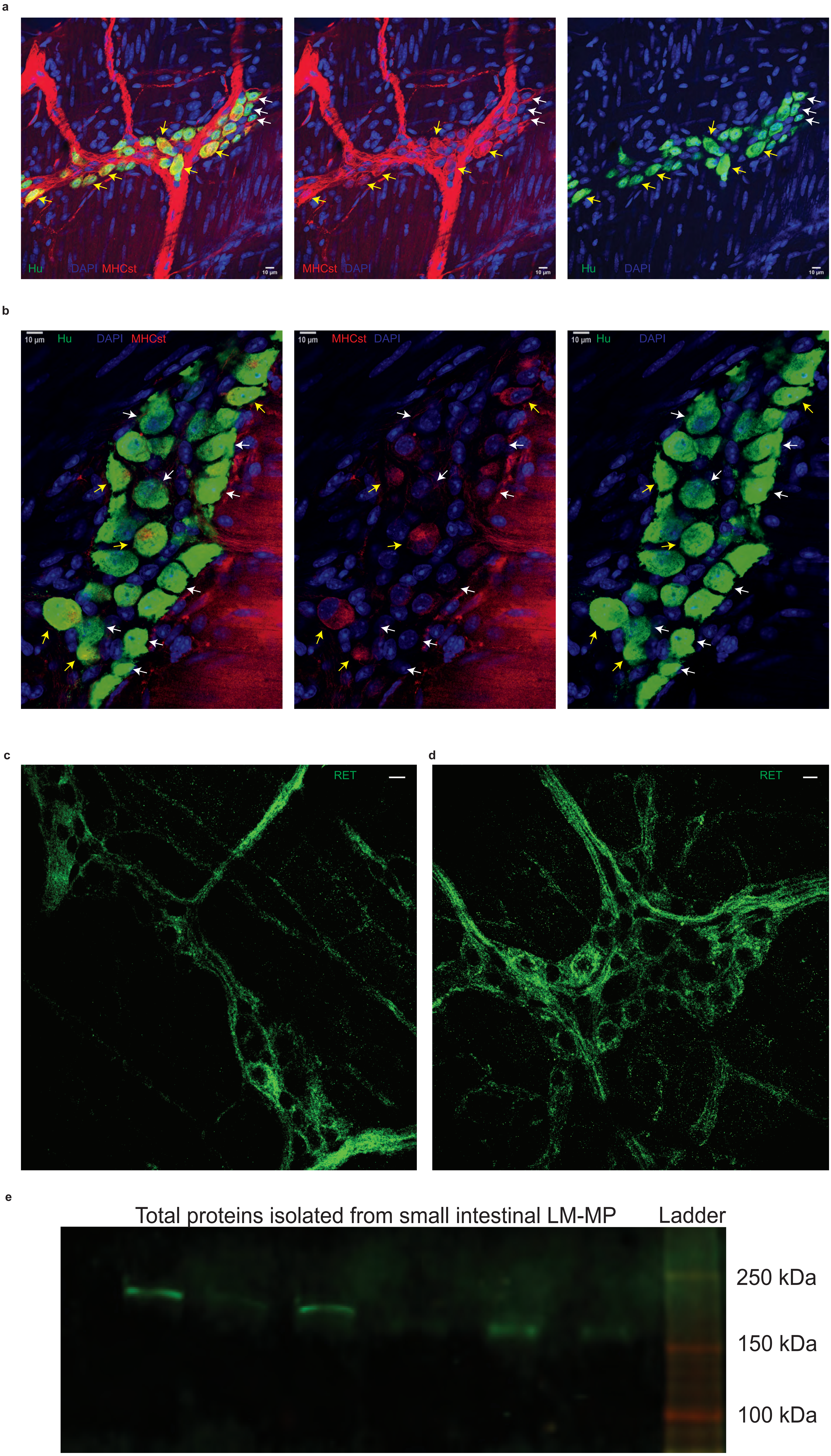


Supplementary Figure 11

Bulk RNA sequencing data from Control and Patients with Obstructed Defecation (OD) projected into murine NMF patterns

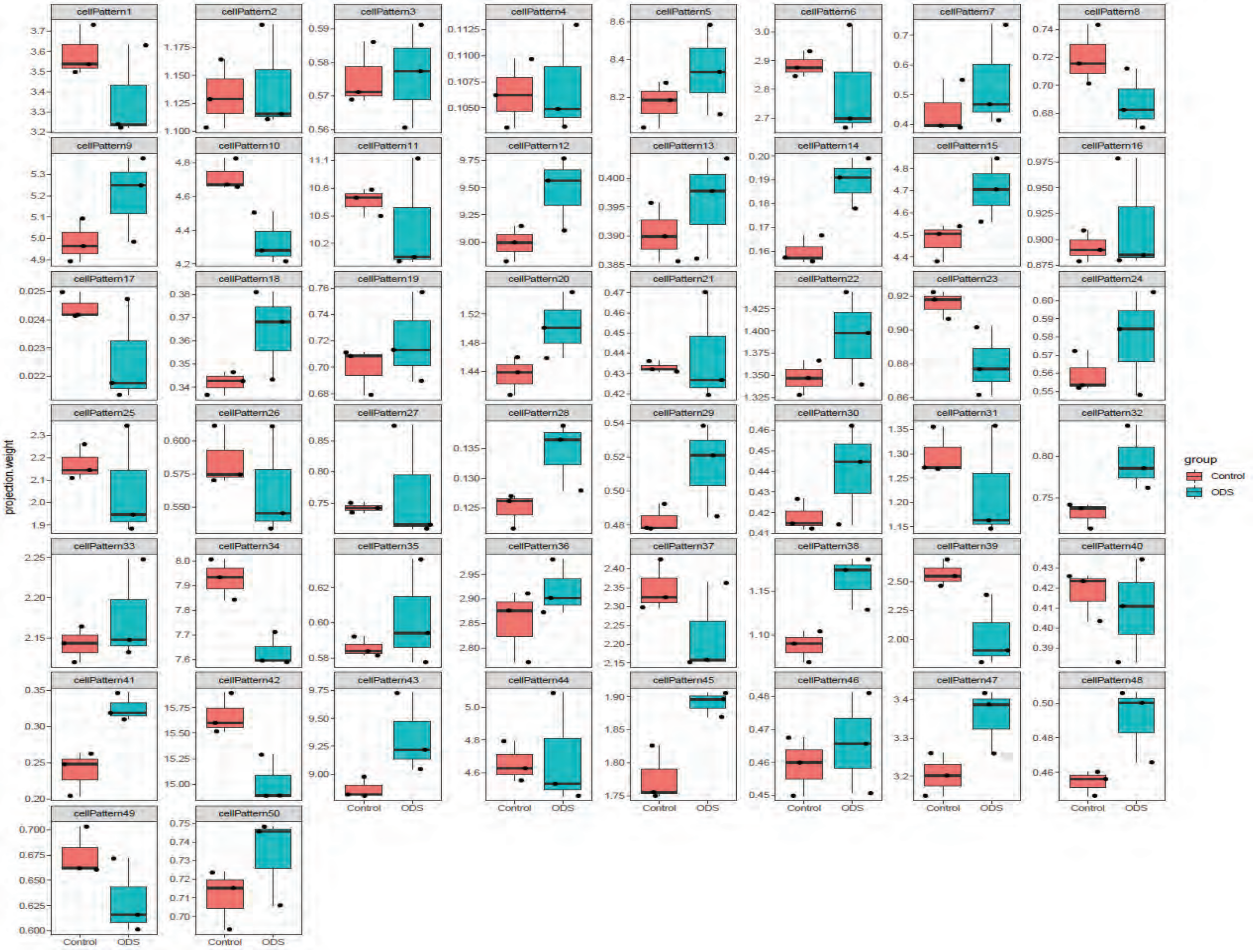


Supplementary Figure 12

**Supplementary Table 1**

**Primary Antibodies and Antisera**

| Antigenic target | Species | Company | Catalogue number | Dilution |
|---|---|---|---|---|
| HuC/D | Human | Human antisera ANNA1 | ANNA1 | 1:750 - 1:1000 |
| HuC/D | Rabbit | Abcam | ab184267 | 1: 500 |
| NOS1 | Rabbit | Invitrogen | 61-7000 | 1: 200 |
| CGRP | Rabbit | Immunostar | 24112 | 1: 500 |
| MET | Goat | R&D systems | AF276 | 1: 250 |
| MHCst (S46) | Mouse | DHSB Iowa | S46 | 1: 250 |
| RET | Rabbit | Abcam | ab134100 | 1: 500 |
| CDH-3 | Rabbit | Abcam | ab137729 | 1: 500 |
| RFP | Rabbit | Rockland | 600-401-379 | 1: 500 |
| GFAP | Rabbit | DAKO | GA-524 | 1: 1000 |
| β ACTIN | Mouse | MP Biomedicals | 0869100-CF | 1: 10000 |
| HGF | Rabbit | Abcam | ab178395 | 1: 500 |
| GDNF | Rabbit | Abcam | ab18956 | 1: 500 |
| GFP | Chicken | Aves | GFP-1020 | 1: 1000 |
| SLPI | Rabbit | Abbexa | abx104088 | 1: 200 |
| AEBP1 | Rabbit | Biorbyt | orb537407 | 1: 250 |
| CLIC3 | Rabbit | Proteintech | 15971-1-AP | 1: 250 |
| CFTR | Rabbit | Abcam | ab181782 | 1: 500 |
| SMO | Rabbit | Proteintech | 20787-1-AP | 1: 250 |
| FMO2 | Rabbit | Biorbyt | orb539506 | 1: 250 |
| MYH11 | Mouse | Thermo-Fisher | MA5-11971 | 1:250 |
| SLC17A9 | Rabbit | Proteintech | 26731-1-AP | 1:200 |
| IL-18 | Rabbit | Abcam | ab207323 | 1:400 |
| NT-3 | Rabbit | Chemicon | ab1532SP | 1:200 |
| MYL-7 | Rabbit | Abcam | ab127001 | 1:100 |
| SNAP-25 | Rabbit | ProteinTech | 14903-1-AP | 1:200 |
| PDE10A | Rabbit | Invitrogen | PA531293 | 1:250 |
| TUBB2B | Rabbit | Invitrogen | PA5-48314 | 1:250 |
| ELAVL2 | Rabbit | Invitrogen | 14008-1-AP | 1:250 |
| HAND2 | Rabbit | Invitrogen | PA5-115339 | 1:250 |
| VSNL1 | Mouse | Invitrogen | 67134-1-Ig | 1:200 |
| GPR88 | Rabbit | Novus Biologicals | NBP1-02330 | 1:200 |
| STMN2 | Rabbit | Invitrogen | 10586-1-AP | 1:250 |
| STX3 | Rabbit | Invitrogen | 15556-1-AP | 1:250 |
| ChAT | Rabbit | Abnova | PAB14536 | 1:100 |
| DECORIN | Mouse | DSHB | 6D6 | 1:250 |
| ECT2 | Rabbit | Bioss | BS-4102R | 1:250 |
| PHOX2B | Rabbit | Proteintech | 25276-1-AP | 1:250 |

**Secondary Antibodies**

| Antigenic target | Species | Company | Catalogue number | Dilution |
|---|---|---|---|---|
| Anti-Rabbit 488 | Goat | Invitrogen | A-11008 | 1:500 |
| Anti-Rabbit 647 | Goat | Invitrogen | A-21245 | 1:500 |
| Anti-Rabbit 647 | Donkey | Invitrogen | A-31573 | 1:500 |
| Anti-Chicken 647 | Donkey | Invitrogen | A-78952 | 1:500 |
| Anti-Chicken 488 | Donkey | Invitrogen | A-78948 | 1:500 |
| Ant-Mouse 647 | Donkey | Invitrogen | A-32787 | 1:500 |
| Anti-Mouse IRdye 680RD | Donkey | Li-Cor | 925-68072 | 1:10000 |
| Anti-Rabbit IRdye 800CW | Donkey | Li-Cor | 925-32213 | 1:10000 |

**TaqMan Probes**

| Probe ID | Target | Company |
|---|---|---|
| Mm01135184_m1 | Hgf | Invitrogen |
| Mm00599849_m1 | Gdnf | Invitrogen |
| Mm00446968_m1 | Hprt | Invitrogen |
| Mm00436304_m1 | Ret | Invitrogen |

| | cell_type | cell_group | gene_id | gene_short_name | marker_score |
|---|---|---|---|---|---|
| 1 | Macrophage-A | 1 | ENSMUSG00000003429 | Rps11 | 0.061354976 |
| 2 | Macrophage-A | 1 | ENSMUSG00000021939 | Ctsb | 0.061056079 |
| 3 | Macrophage-A | 1 | ENSMUSG00000006333 | Rps9 | 0.065563158 |
| 4 | Macrophage-A | 1 | ENSMUSG00000007892 | Rplp1 | 0.08632253 |
| 5 | Macrophage-A | 1 | ENSMUSG00000021242 | Npc2 | 0.057865841 |
| 6 | Macrophage-A | 1 | ENSMUSG00000022108 | Itm2b | 0.083967704 |
| 7 | Macrophage-A | 1 | ENSMUSG00000024397 | Aif1 | 0.148549596 |
| 8 | Macrophage-A | 1 | ENSMUSG00000024608 | Rps14 | 0.064703869 |
| 9 | Macrophage-A | 1 | ENSMUSG00000024661 | Fth1 | 0.143079328 |
| 10 | Macrophage-A | 1 | ENSMUSG00000027447 | Cst3 | 0.121007699 |
| 11 | Macrophage-A | 1 | ENSMUSG00000029373 | Pf4 | 0.1171735 |
| 12 | Macrophage-A | 1 | ENSMUSG00000030579 | Tyrobp | 0.124438212 |
| 13 | Macrophage-A | 1 | ENSMUSG00000031320 | Rps4x | 0.061680629 |
| 14 | Macrophage-A | 1 | ENSMUSG00000032554 | Trf | 0.059881634 |
| 15 | Macrophage-A | 1 | ENSMUSG00000034892 | Rps29 | 0.108320203 |
| 16 | Macrophage-A | 1 | ENSMUSG00000036594 | H2-Aa | 0.150412226 |
| 17 | Macrophage-A | 1 | ENSMUSG00000073421 | H2-Ab1 | 0.135365962 |
| 18 | Macrophage-A | 1 | ENSMUSG00000038274 | Fau | 0.075862596 |
| 19 | Macrophage-A | 1 | ENSMUSG00000036887 | C1qa | 0.156058866 |
| 20 | Macrophage-A | 1 | ENSMUSG00000036896 | C1qc | 0.119350241 |
| 21 | Macrophage-A | 1 | ENSMUSG00000036905 | C1qb | 0.15394097 |
| 22 | Macrophage-A | 1 | ENSMUSG00000024610 | Cd74 | 0.18406852 |
| 23 | Macrophage-A | 1 | ENSMUSG00000045128 | Rpl18a | 0.063317371 |
| 24 | Macrophage-A | 1 | ENSMUSG00000079017 | Ifi27l2a | 0.076158299 |
| 25 | Macrophage-A | 1 | ENSMUSG00000046330 | Rpl37a | 0.078167962 |
| 26 | Macrophage-A | 1 | ENSMUSG00000062006 | Rpl34 | 0.058457762 |
| 27 | Macrophage-A | 1 | ENSMUSG00000060938 | Rpl26 | 0.061960827 |
| 28 | Macrophage-A | 1 | ENSMUSG00000060586 | H2-Eb1 | 0.17119384 |
| 29 | Macrophage-A | 1 | ENSMUSG00000057322 | Rpl38 | 0.058504818 |
| 30 | Macrophage-A | 1 | ENSMUSG00000060636 | Rpl35a | 0.077783791 |
| 31 | Macrophage-A | 1 | ENSMUSG00000058715 | Fcer1g | 0.151309839 |
| 32 | Macrophage-A | 1 | ENSMUSG00000009927 | Rps25 | 0.05938735 |
| 33 | Macrophage-A | 1 | ENSMUSG00000062997 | Rpl35 | 0.059566007 |
| 34 | Macrophage-A | 1 | ENSMUSG00000057841 | Rpl32 | 0.064390862 |
| 35 | Macrophage-A | 1 | ENSMUSG00000064339 | mt-Rnr2 | 0.098665241 |
| 36 | Macrophage-A | 1 | ENSMUSG00000064341 | mt-Nd1 | 0.108844678 |
| 37 | Macrophage-A | 1 | ENSMUSG00000064345 | mt-Nd2 | 0.092335043 |
| 38 | Macrophage-A | 1 | ENSMUSG00000064357 | mt-Atp6 | 0.069185731 |
| 39 | Macrophage-A | 1 | ENSMUSG00000064358 | mt-Co3 | 0.088653046 |
| 40 | Macrophage-A | 1 | ENSMUSG00000064363 | mt-Nd4 | 0.111394777 |
| 41 | Macrophage-A | 1 | ENSMUSG00000064370 | mt-Cytb | 0.109911834 |
| 42 | Macrophage-A | 1 | ENSMUSG00000025508 | Rplp2 | 0.057979361 |
| 43 | Macrophage-A | 1 | ENSMUSG00000069516 | Lyz2 | 0.094195639 |
| 44 | Macrophage-A | 1 | ENSMUSG00000050708 | Ftl1 | 0.170054521 |

| | | | | | |
|---|---|---|---|---|---|
| 45 | Macrophage-A | 1 | ENSMUSG00000073412 | Lst1 | 0.101378574 |
| 46 | Macrophage-A | 1 | ENSMUSG00000029580 | Actb | 0.081340731 |
| 47 | Macrophage-A | 1 | ENSMUSG00000049775 | Tmsb4x | 0.160430527 |
| 48 | Macrophage-A | 1 | ENSMUSG00000025290 | Rps24 | 0.058450828 |
| 49 | Macrophage-A | 1 | ENSMUSG00000079641 | Rpl39 | 0.063696167 |
| 50 | Macrophage-A | 1 | ENSMUSG00000093674 | Rpl41 | 0.06685247 |
| 51 | Penk+ Fibroblasts | 10 | ENSMUSG00000000753 | Serpinf1 | 0.281868791 |
| 52 | Penk+ Fibroblasts | 10 | ENSMUSG00000001119 | Col6a1 | 0.248476123 |
| 53 | Penk+ Fibroblasts | 10 | ENSMUSG00000020241 | Col6a2 | 0.216900959 |
| 54 | Penk+ Fibroblasts | 10 | ENSMUSG00000001506 | Col1a1 | 0.308236247 |
| 55 | Penk+ Fibroblasts | 10 | ENSMUSG00000003477 | Inmt | 0.393969285 |
| 56 | Penk+ Fibroblasts | 10 | ENSMUSG00000058135 | Gstm1 | 0.206195673 |
| 57 | Penk+ Fibroblasts | 10 | ENSMUSG00000005397 | Nid1 | 0.349665379 |
| 58 | Penk+ Fibroblasts | 10 | ENSMUSG00000015354 | Pcolce2 | 0.540784376 |
| 59 | Penk+ Fibroblasts | 10 | ENSMUSG00000017493 | Igfbp4 | 0.215026487 |
| 60 | Penk+ Fibroblasts | 10 | ENSMUSG00000020363 | Gfpt2 | 0.301843593 |
| 61 | Penk+ Fibroblasts | 10 | ENSMUSG00000020810 | Cygb | 0.304453972 |
| 62 | Penk+ Fibroblasts | 10 | ENSMUSG00000021047 | Nova1 | 0.236825438 |
| 63 | Penk+ Fibroblasts | 10 | ENSMUSG00000021186 | Fbln5 | 0.205177361 |
| 64 | Penk+ Fibroblasts | 10 | ENSMUSG00000021876 | Rnase4 | 0.223700407 |
| 65 | Penk+ Fibroblasts | 10 | ENSMUSG00000022032 | Scara5 | 0.567938182 |
| 66 | Penk+ Fibroblasts | 10 | ENSMUSG00000022371 | Col14a1 | 0.318301321 |
| 67 | Penk+ Fibroblasts | 10 | ENSMUSG00000075602 | Ly6a | 0.225719455 |
| 68 | Penk+ Fibroblasts | 10 | ENSMUSG00000022894 | Adamts5 | 0.23078803 |
| 69 | Penk+ Fibroblasts | 10 | ENSMUSG00000023224 | Serping1 | 0.201258951 |
| 70 | Penk+ Fibroblasts | 10 | ENSMUSG00000024076 | Vit | 0.307463757 |
| 71 | Penk+ Fibroblasts | 10 | ENSMUSG00000025784 | Clec3b | 0.511313122 |
| 72 | Penk+ Fibroblasts | 10 | ENSMUSG00000026478 | Lamc1 | 0.266532987 |
| 73 | Penk+ Fibroblasts | 10 | ENSMUSG00000026674 | Ddr2 | 0.243373949 |
| 74 | Penk+ Fibroblasts | 10 | ENSMUSG00000026879 | Gsn | 0.662758607 |
| 75 | Penk+ Fibroblasts | 10 | ENSMUSG00000027204 | Fbn1 | 0.419276537 |
| 76 | Penk+ Fibroblasts | 10 | ENSMUSG00000029061 | Mmp23 | 0.201315212 |
| 77 | Penk+ Fibroblasts | 10 | ENSMUSG00000029661 | Col1a2 | 0.352093693 |
| 78 | Penk+ Fibroblasts | 10 | ENSMUSG00000030116 | Mfap5 | 0.54569428 |
| 79 | Penk+ Fibroblasts | 10 | ENSMUSG00000064080 | Fbln2 | 0.293453979 |
| 80 | Penk+ Fibroblasts | 10 | ENSMUSG00000039405 | Prss23 | 0.222049065 |
| 81 | Penk+ Fibroblasts | 10 | ENSMUSG00000034463 | Scara3 | 0.24476053 |
| 82 | Penk+ Fibroblasts | 10 | ENSMUSG00000028369 | Svep1 | 0.199300255 |
| 83 | Penk+ Fibroblasts | 10 | ENSMUSG00000054203 | Ifi205 | 0.230358477 |
| 84 | Penk+ Fibroblasts | 10 | ENSMUSG00000022665 | Ccdc80 | 0.24105144 |
| 85 | Penk+ Fibroblasts | 10 | ENSMUSG00000032334 | Loxl1 | 0.290317309 |
| 86 | Penk+ Fibroblasts | 10 | ENSMUSG00000028517 | Plpp3 | 0.232461178 |
| 87 | Penk+ Fibroblasts | 10 | ENSMUSG00000045573 | Penk | 0.379125541 |
| 88 | Penk+ Fibroblasts | 10 | ENSMUSG00000056481 | Cd248 | 0.403669781 |
| 89 | Penk+ Fibroblasts | 10 | ENSMUSG00000057098 | Ebf1 | 0.200985086 |

| | | | | | |
|---|---|---|---|---|---|
| 90 | Penk+ Fibroblasts | 10 | ENSMUSG00000033327 | Tnxb | 0.345536074 |
| 91 | Penk+ Fibroblasts | 10 | ENSMUSG00000029096 | Htra3 | 0.440656542 |
| 92 | Penk+ Fibroblasts | 10 | ENSMUSG00000026043 | Col3a1 | 0.378685992 |
| 93 | Penk+ Fibroblasts | 10 | ENSMUSG00000027074 | Slc43a3 | 0.239706114 |
| 94 | Penk+ Fibroblasts | 10 | ENSMUSG00000029659 | Medag | 0.205830577 |
| 95 | Penk+ Fibroblasts | 10 | ENSMUSG00000070436 | Serpinh1 | 0.193920545 |
| 96 | Penk+ Fibroblasts | 10 | ENSMUSG00000071984 | Fndc1 | 0.274119495 |
| 97 | Penk+ Fibroblasts | 10 | ENSMUSG00000019929 | Dcn | 0.31992032 |
| 98 | Penk+ Fibroblasts | 10 | ENSMUSG00000024011 | Pi16 | 0.298930222 |
| 99 | Penk+ Fibroblasts | 10 | ENSMUSG00000022816 | Fstl1 | 0.301602581 |
| 100 | Penk+ Fibroblasts | 10 | ENSMUSG00000038521 | C1s1 | 0.251531426 |
| 101 | T cells | 11 | ENSMUSG00000002033 | Cd3g | 0.593485338 |
| 102 | T cells | 11 | ENSMUSG00000005947 | Itgae | 0.508136894 |
| 103 | T cells | 11 | ENSMUSG00000015437 | Gzmb | 0.582090145 |
| 104 | T cells | 11 | ENSMUSG00000021108 | Prkch | 0.20503606 |
| 105 | T cells | 11 | ENSMUSG00000023132 | Gzma | 0.671809652 |
| 106 | T cells | 11 | ENSMUSG00000024910 | Ctsw | 0.374402304 |
| 107 | T cells | 11 | ENSMUSG00000025163 | Cd7 | 0.284760988 |
| 108 | T cells | 11 | ENSMUSG00000025647 | Shisa5 | 0.148245644 |
| 109 | T cells | 11 | ENSMUSG00000026117 | Zap70 | 0.160315007 |
| 110 | T cells | 11 | ENSMUSG00000026826 | Nr4a2 | 0.3870486 |
| 111 | T cells | 11 | ENSMUSG00000079110 | Capn3 | 0.161355733 |
| 112 | T cells | 11 | ENSMUSG00000027843 | Ptpn22 | 0.199290417 |
| 113 | T cells | 11 | ENSMUSG00000028071 | Sh2d2a | 0.143231368 |
| 114 | T cells | 11 | ENSMUSG00000029603 | Dtx1 | 0.144809734 |
| 115 | T cells | 11 | ENSMUSG00000030124 | Lag3 | 0.230031024 |
| 116 | T cells | 11 | ENSMUSG00000030165 | Klrd1 | 0.321639897 |
| 117 | T cells | 11 | ENSMUSG00000032094 | Cd3d | 0.423193245 |
| 118 | T cells | 11 | ENSMUSG00000032380 | Dapk2 | 0.188187408 |
| 119 | T cells | 11 | ENSMUSG00000035042 | Ccl5 | 0.657312485 |
| 120 | T cells | 11 | ENSMUSG00000036478 | Btg1 | 0.18303405 |
| 121 | T cells | 11 | ENSMUSG00000037754 | Ppp1r16b | 0.247327356 |
| 122 | T cells | 11 | ENSMUSG00000040957 | Cables1 | 0.151907801 |
| 123 | T cells | 11 | ENSMUSG00000038304 | Cd160 | 0.241552054 |
| 124 | T cells | 11 | ENSMUSG00000070691 | Runx3 | 0.308620597 |
| 125 | T cells | 11 | ENSMUSG00000052374 | Actn2 | 0.190069567 |
| 126 | T cells | 11 | ENSMUSG00000052837 | Junb | 0.153737577 |
| 127 | T cells | 11 | ENSMUSG00000053977 | Cd8a | 0.691523506 |
| 128 | T cells | 11 | ENSMUSG00000000409 | Lck | 0.334227347 |
| 129 | T cells | 11 | ENSMUSG00000054435 | Gimap4 | 0.160339721 |
| 130 | T cells | 11 | ENSMUSG00000004612 | Nkg7 | 0.408414339 |
| 131 | T cells | 11 | ENSMUSG00000064109 | Hcst | 0.152992717 |
| 132 | T cells | 11 | ENSMUSG00000068129 | Cst7 | 0.182240395 |
| 133 | T cells | 11 | ENSMUSG00000068227 | Il2rb | 0.205621928 |
| 134 | T cells | 11 | ENSMUSG00000070803 | Cited4 | 0.268734299 |

| | | | | | |
|---|---|---|---|---|---|
| 135 | T cells | 11 | ENSMUSG00000075010 | AW112010 | 0.258465808 |
| 136 | T cells | 11 | ENSMUSG00000090176 | Cd200r2 | 0.161809761 |
| 137 | T cells | 11 | ENSMUSG00000032093 | Cd3e | 0.481197893 |
| 138 | T cells | 11 | ENSMUSG00000076490 | Trbc1 | 0.15974128 |
| 139 | T cells | 11 | ENSMUSG00000076498 | Trbc2 | 0.302630756 |
| 140 | T cells | 11 | ENSMUSG00000076749 | Tcrg-C1 | 0.301007895 |
| 141 | T cells | 11 | ENSMUSG00000076752 | Tcrg-C2 | 0.309285426 |
| 142 | T cells | 11 | ENSMUSG00000076928 | Trac | 0.306932905 |
| 143 | T cells | 11 | ENSMUSG00000026395 | Ptprc | 0.144339901 |
| 144 | T cells | 11 | ENSMUSG00000062082 | Cd200r4 | 0.188410814 |
| 145 | T cells | 11 | ENSMUSG00000026358 | Rgs1 | 0.369041325 |
| 146 | T cells | 11 | ENSMUSG00000030775 | Trat1 | 0.146925843 |
| 147 | T cells | 11 | ENSMUSG00000076757 | Tcrg-C4 | 0.231642042 |
| 148 | T cells | 11 | ENSMUSG00000102112 | 1810041H14Rik | 0.170229197 |
| 149 | T cells | 11 | ENSMUSG00000100150 | Gm19585 | 0.287087328 |
| 150 | T cells | 11 | ENSMUSG00000114333 | 1700084D21Rik | 0.304306721 |
| 151 | NENs | 12 | ENSMUSG00000006575 | Rundc3a | 0.321553502 |
| 152 | NENs | 12 | ENSMUSG00000021087 | Rtn1 | 0.323537805 |
| 153 | NENs | 12 | ENSMUSG00000021647 | Cartpt | 0.199037817 |
| 154 | NENs | 12 | ENSMUSG00000023484 | Prph | 0.493389128 |
| 155 | NENs | 12 | ENSMUSG00000024268 | Celf4 | 0.277638792 |
| 156 | NENs | 12 | ENSMUSG00000025777 | Gdap1 | 0.202042766 |
| 157 | NENs | 12 | ENSMUSG00000026204 | Ptprn | 0.326552748 |
| 158 | NENs | 12 | ENSMUSG00000027273 | Snap25 | 0.347553426 |
| 159 | NENs | 12 | ENSMUSG00000027500 | Stmn2 | 0.306553158 |
| 160 | NENs | 12 | ENSMUSG00000027801 | Tm4sf4 | 0.254405112 |
| 161 | NENs | 12 | ENSMUSG00000028137 | Celf3 | 0.318222696 |
| 162 | NENs | 12 | ENSMUSG00000061601 | Pclo | 0.216977049 |
| 163 | NENs | 12 | ENSMUSG00000029223 | Uchl1 | 0.259136384 |
| 164 | NENs | 12 | ENSMUSG00000030000 | Add2 | 0.199082787 |
| 165 | NENs | 12 | ENSMUSG00000031137 | Fgf13 | 0.202911151 |
| 166 | NENs | 12 | ENSMUSG00000032303 | Chrna3 | 0.242814369 |
| 167 | NENs | 12 | ENSMUSG00000032549 | Rab6b | 0.211236233 |
| 168 | NENs | 12 | ENSMUSG00000034891 | Sncb | 0.203382079 |
| 169 | NENs | 12 | ENSMUSG00000033061 | Resp18 | 0.2282748 |
| 170 | NENs | 12 | ENSMUSG00000039278 | Pcsk1n | 0.441654137 |
| 171 | NENs | 12 | ENSMUSG00000004113 | Cacna1b | 0.229604789 |
| 172 | NENs | 12 | ENSMUSG00000070695 | Cntnap5a | 0.29451253 |
| 173 | NENs | 12 | ENSMUSG00000032826 | Ank2 | 0.274719411 |
| 174 | NENs | 12 | ENSMUSG00000042750 | Bex2 | 0.211689045 |
| 175 | NENs | 12 | ENSMUSG00000002265 | Peg3 | 0.249571405 |
| 176 | NENs | 12 | ENSMUSG00000029245 | Epha5 | 0.312925541 |
| 177 | NENs | 12 | ENSMUSG00000044288 | Cnr1 | 0.228104128 |
| 178 | NENs | 12 | ENSMUSG00000048978 | Nrsn1 | 0.202933713 |
| 179 | NENs | 12 | ENSMUSG00000055567 | Unc80 | 0.22880753 |

| | | | | | |
|---|---|---|---|---|---|
| 180 | NENs | 12 | ENSMUSG00000035864 | Syt1 | 0.204257423 |
| 181 | NENs | 12 | ENSMUSG00000052727 | Map1b | 0.239069513 |
| 182 | NENs | 12 | ENSMUSG00000061576 | Dpp6 | 0.219034797 |
| 183 | NENs | 12 | ENSMUSG00000074483 | Bglap | 0.222031338 |
| 184 | NENs | 12 | ENSMUSG00000037217 | Syn1 | 0.288534064 |
| 185 | NENs | 12 | ENSMUSG00000018865 | Sult4a1 | 0.197197165 |
| 186 | NENs | 12 | ENSMUSG00000029361 | Nos1 | 0.24302464 |
| 187 | NENs | 12 | ENSMUSG00000014602 | Kif1a | 0.266600226 |
| 188 | NENs | 12 | ENSMUSG00000022658 | Tagln3 | 0.206379642 |
| 189 | NENs | 12 | ENSMUSG00000031837 | Necab2 | 0.228305948 |
| 190 | NENs | 12 | ENSMUSG00000018411 | Mapt | 0.209892124 |
| 191 | NENs | 12 | ENSMUSG00000028546 | Elavl4 | 0.431189449 |
| 192 | NENs | 12 | ENSMUSG00000076441 | Ass1 | 0.272813998 |
| 193 | NENs | 12 | ENSMUSG00000027581 | Stmn3 | 0.269574121 |
| 194 | NENs | 12 | ENSMUSG00000006930 | Hap1 | 0.227019763 |
| 195 | NENs | 12 | ENSMUSG00000019986 | Ahi1 | 0.303765359 |
| 196 | NENs | 12 | ENSMUSG00000038738 | Shank1 | 0.219337403 |
| 197 | NENs | 12 | ENSMUSG00000044349 | Snhg11 | 0.648027475 |
| 198 | NENs | 12 | ENSMUSG00000087620 | 5330434G04Rik | 0.249457376 |
| 199 | NENs | 12 | ENSMUSG00000021700 | Rab3c | 0.220071922 |
| 200 | NENs | 12 | ENSMUSG00000097451 | Rian | 0.378275766 |
| 201 | NK cells | 13 | ENSMUSG00000001281 | Itgb7 | 0.142470509 |
| 202 | NK cells | 13 | ENSMUSG00000003882 | Il7r | 0.382680281 |
| 203 | NK cells | 13 | ENSMUSG00000007892 | Rplp1 | 0.121948633 |
| 204 | NK cells | 13 | ENSMUSG00000008668 | Rps18 | 0.130535555 |
| 205 | NK cells | 13 | ENSMUSG00000015312 | Gadd45b | 0.145639565 |
| 206 | NK cells | 13 | ENSMUSG00000016559 | H3f3b | 0.141769338 |
| 207 | NK cells | 13 | ENSMUSG00000017404 | Rpl19 | 0.13420677 |
| 208 | NK cells | 13 | ENSMUSG00000018239 | Zcchc10 | 0.212013263 |
| 209 | NK cells | 13 | ENSMUSG00000019987 | Arg1 | 0.271168023 |
| 210 | NK cells | 13 | ENSMUSG00000020395 | Itk | 0.161714817 |
| 211 | NK cells | 13 | ENSMUSG00000020423 | Btg2 | 0.138428109 |
| 212 | NK cells | 13 | ENSMUSG00000020577 | Tspan13 | 0.121243679 |
| 213 | NK cells | 13 | ENSMUSG00000020644 | Id2 | 0.267325862 |
| 214 | NK cells | 13 | ENSMUSG00000021025 | Nfkbia | 0.138086745 |
| 215 | NK cells | 13 | ENSMUSG00000021298 | Gpr132 | 0.1849573 |
| 216 | NK cells | 13 | ENSMUSG00000021728 | Emb | 0.184521752 |
| 217 | NK cells | 13 | ENSMUSG00000022015 | Tnfsf11 | 0.156866447 |
| 218 | NK cells | 13 | ENSMUSG00000024399 | Ltb | 0.317065109 |
| 219 | NK cells | 13 | ENSMUSG00000025647 | Shisa5 | 0.131705638 |
| 220 | NK cells | 13 | ENSMUSG00000026360 | Rgs2 | 0.218149554 |
| 221 | NK cells | 13 | ENSMUSG00000026770 | Il2ra | 0.182087007 |
| 222 | NK cells | 13 | ENSMUSG00000030114 | Klrg1 | 0.35739415 |
| 223 | NK cells | 13 | ENSMUSG00000030707 | Coro1a | 0.128516074 |
| 224 | NK cells | 13 | ENSMUSG00000031880 | Rrad | 0.139268255 |

| | | | | | |
|---|---|---|---|---|---|
| 225 | NK cells | 13 | ENSMUSG00000032035 | Ets1 | 0.134268992 |
| 226 | NK cells | 13 | ENSMUSG00000032238 | Rora | 0.149083918 |
| 227 | NK cells | 13 | ENSMUSG00000034724 | Cnot6l | 0.120214358 |
| 228 | NK cells | 13 | ENSMUSG00000036155 | Mgat5 | 0.119338129 |
| 229 | NK cells | 13 | ENSMUSG00000036478 | Btg1 | 0.178481432 |
| 230 | NK cells | 13 | ENSMUSG00000039264 | Gimap3 | 0.217432048 |
| 231 | NK cells | 13 | ENSMUSG00000037742 | Eef1a1 | 0.133178681 |
| 232 | NK cells | 13 | ENSMUSG00000026069 | Il1rl1 | 0.480124864 |
| 233 | NK cells | 13 | ENSMUSG00000047898 | Ccr4 | 0.123478796 |
| 234 | NK cells | 13 | ENSMUSG00000046908 | Ltb4r1 | 0.230448279 |
| 235 | NK cells | 13 | ENSMUSG00000045817 | Zfp36l2 | 0.148561199 |
| 236 | NK cells | 13 | ENSMUSG00000030108 | Slc6a13 | 0.123092767 |
| 237 | NK cells | 13 | ENSMUSG00000052837 | Junb | 0.204595109 |
| 238 | NK cells | 13 | ENSMUSG00000053310 | Nrgn | 0.240641677 |
| 239 | NK cells | 13 | ENSMUSG00000054408 | Spcs3 | 0.146148708 |
| 240 | NK cells | 13 | ENSMUSG00000057058 | Skap1 | 0.143329313 |
| 241 | NK cells | 13 | ENSMUSG00000079845 | Xlr4a | 0.234275915 |
| 242 | NK cells | 13 | ENSMUSG00000062210 | Tnfaip8 | 0.147988509 |
| 243 | NK cells | 13 | ENSMUSG00000018476 | Kdm6b | 0.154723152 |
| 244 | NK cells | 13 | ENSMUSG00000015619 | Gata3 | 0.462848611 |
| 245 | NK cells | 13 | ENSMUSG00000031438 | Rnf128 | 0.312707591 |
| 246 | NK cells | 13 | ENSMUSG00000022876 | Samsn1 | 0.180165768 |
| 247 | NK cells | 13 | ENSMUSG00000026644 | Acbd7 | 0.237454103 |
| 248 | NK cells | 13 | ENSMUSG00000028266 | Lmo4 | 0.177638154 |
| 249 | NK cells | 13 | ENSMUSG00000082263 | Gm11188 | 0.128205128 |
| 250 | NK cells | 13 | ENSMUSG00000025794 | Rpl14 | 0.135276409 |
| 251 | Macrophage-C | 14 | ENSMUSG00000001525 | Tubb5 | 0.299078242 |
| 252 | Macrophage-C | 14 | ENSMUSG00000005233 | Spc25 | 0.270672679 |
| 253 | Macrophage-C | 14 | ENSMUSG00000005470 | Asf1b | 0.350185519 |
| 254 | Macrophage-C | 14 | ENSMUSG00000006398 | Cdc20 | 0.262906288 |
| 255 | Macrophage-C | 14 | ENSMUSG00000019942 | Cdk1 | 0.349547131 |
| 256 | Macrophage-C | 14 | ENSMUSG00000020330 | Hmmr | 0.336027119 |
| 257 | Macrophage-C | 14 | ENSMUSG00000020649 | Rrm2 | 0.365025177 |
| 258 | Macrophage-C | 14 | ENSMUSG00000020897 | Aurkb | 0.267712456 |
| 259 | Macrophage-C | 14 | ENSMUSG00000022033 | Pbk | 0.301709025 |
| 260 | Macrophage-C | 14 | ENSMUSG00000022034 | Esco2 | 0.278583083 |
| 261 | Macrophage-C | 14 | ENSMUSG00000022322 | Shcbp1 | 0.370129114 |
| 262 | Macrophage-C | 14 | ENSMUSG00000023015 | Racgap1 | 0.218198436 |
| 263 | Macrophage-C | 14 | ENSMUSG00000023505 | Cdca3 | 0.373417443 |
| 264 | Macrophage-C | 14 | ENSMUSG00000024590 | Lmnb1 | 0.238418089 |
| 265 | Macrophage-C | 14 | ENSMUSG00000024660 | Incenp | 0.334095064 |
| 266 | Macrophage-C | 14 | ENSMUSG00000026196 | Bard1 | 0.230135861 |
| 267 | Macrophage-C | 14 | ENSMUSG00000026429 | Ube2t | 0.260541862 |
| 268 | Macrophage-C | 14 | ENSMUSG00000027306 | Nusap1 | 0.31989859 |
| 269 | Macrophage-C | 14 | ENSMUSG00000027496 | Aurka | 0.221035835 |

| | | | | | |
|---|---|---|---|---|---|
| 270 | Macrophage-C | 14 | ENSMUSG00000027715 | Ccna2 | 0.25720012 |
| 271 | Macrophage-C | 14 | ENSMUSG00000028044 | Cks1b | 0.221729311 |
| 272 | Macrophage-C | 14 | ENSMUSG00000028832 | Stmn1 | 0.360955222 |
| 273 | Macrophage-C | 14 | ENSMUSG00000028873 | Cdca8 | 0.391777533 |
| 274 | Macrophage-C | 14 | ENSMUSG00000030677 | Kif22 | 0.242910555 |
| 275 | Macrophage-C | 14 | ENSMUSG00000030978 | Rrm1 | 0.295954426 |
| 276 | Macrophage-C | 14 | ENSMUSG00000031004 | Mki67 | 0.315888022 |
| 277 | Macrophage-C | 14 | ENSMUSG00000032218 | Ccnb2 | 0.439936966 |
| 278 | Macrophage-C | 14 | ENSMUSG00000040084 | Bub1b | 0.237823646 |
| 279 | Macrophage-C | 14 | ENSMUSG00000034349 | Smc4 | 0.255312392 |
| 280 | Macrophage-C | 14 | ENSMUSG00000035683 | Melk | 0.237587856 |
| 281 | Macrophage-C | 14 | ENSMUSG00000038943 | Prc1 | 0.305656591 |
| 282 | Macrophage-C | 14 | ENSMUSG00000042489 | Clspn | 0.22576418 |
| 283 | Macrophage-C | 14 | ENSMUSG00000049932 | H2afx | 0.374259091 |
| 284 | Macrophage-C | 14 | ENSMUSG00000041859 | Mcm3 | 0.224724002 |
| 285 | Macrophage-C | 14 | ENSMUSG00000045328 | Cenpe | 0.220815525 |
| 286 | Macrophage-C | 14 | ENSMUSG00000037628 | Cdkn3 | 0.251248221 |
| 287 | Macrophage-C | 14 | ENSMUSG00000054717 | Hmgb2 | 0.365372277 |
| 288 | Macrophage-C | 14 | ENSMUSG00000020914 | Top2a | 0.461534001 |
| 289 | Macrophage-C | 14 | ENSMUSG00000023004 | Tuba1b | 0.262612473 |
| 290 | Macrophage-C | 14 | ENSMUSG00000094777 | Hist1h2ap | 0.34784987 |
| 291 | Macrophage-C | 14 | ENSMUSG00000017716 | Birc5 | 0.517169608 |
| 292 | Macrophage-C | 14 | ENSMUSG00000068101 | Cenpm | 0.335722626 |
| 293 | Macrophage-C | 14 | ENSMUSG00000024989 | Cep55 | 0.285990578 |
| 294 | Macrophage-C | 14 | ENSMUSG00000056394 | Lig1 | 0.224593801 |
| 295 | Macrophage-C | 14 | ENSMUSG00000074476 | Spc24 | 0.292861791 |
| 296 | Macrophage-C | 14 | ENSMUSG00000029910 | Mad2l1 | 0.244955881 |
| 297 | Macrophage-C | 14 | ENSMUSG00000028312 | Smc2 | 0.230468243 |
| 298 | Macrophage-C | 14 | ENSMUSG00000015880 | Ncapg | 0.246618015 |
| 299 | Macrophage-C | 14 | ENSMUSG00000026605 | Cenpf | 0.288160636 |
| 300 | Macrophage-C | 14 | ENSMUSG00000098318 | Lockd | 0.311924078 |
| 301 | NK cells | 15 | ENSMUSG00000062524 | Ncr1 | 0.223542214 |
| 302 | NK cells | 15 | ENSMUSG00000008668 | Rps18 | 0.143265788 |
| 303 | NK cells | 15 | ENSMUSG00000009687 | Fxyd5 | 0.132072262 |
| 304 | NK cells | 15 | ENSMUSG00000089727 | Klra8 | 0.131578947 |
| 305 | NK cells | 15 | ENSMUSG00000018293 | Pfn1 | 0.132392772 |
| 306 | NK cells | 15 | ENSMUSG00000020009 | Ifngr1 | 0.159861794 |
| 307 | NK cells | 15 | ENSMUSG00000020644 | Id2 | 0.184086979 |
| 308 | NK cells | 15 | ENSMUSG00000024014 | Pim1 | 0.150987217 |
| 309 | NK cells | 15 | ENSMUSG00000024399 | Ltb | 0.146144194 |
| 310 | NK cells | 15 | ENSMUSG00000024910 | Ctsw | 0.153801414 |
| 311 | NK cells | 15 | ENSMUSG00000025647 | Shisa5 | 0.14137411 |
| 312 | NK cells | 15 | ENSMUSG00000026012 | Cd28 | 0.30124175 |
| 313 | NK cells | 15 | ENSMUSG00000027863 | Cd2 | 0.172420334 |
| 314 | NK cells | 15 | ENSMUSG00000030149 | Klrk1 | 0.181419937 |

| | | | | | |
|---|---|---|---|---|---|
| 315 | NK cells | 15 | ENSMUSG00000030167 | Klrc1 | 0.175143648 |
| 316 | NK cells | 15 | ENSMUSG00000030220 | Arhgdib | 0.146799638 |
| 317 | NK cells | 15 | ENSMUSG00000030707 | Coro1a | 0.171510164 |
| 318 | NK cells | 15 | ENSMUSG00000030744 | Rps3 | 0.141660649 |
| 319 | NK cells | 15 | ENSMUSG00000031132 | Cd40lg | 0.157894737 |
| 320 | NK cells | 15 | ENSMUSG00000032035 | Ets1 | 0.134679655 |
| 321 | NK cells | 15 | ENSMUSG00000032094 | Cd3d | 0.177221883 |
| 322 | NK cells | 15 | ENSMUSG00000056290 | Ms4a4b | 0.628441098 |
| 323 | NK cells | 15 | ENSMUSG00000036478 | Btg1 | 0.13978196 |
| 324 | NK cells | 15 | ENSMUSG00000039264 | Gimap3 | 0.208222742 |
| 325 | NK cells | 15 | ENSMUSG00000040747 | Cd53 | 0.163577587 |
| 326 | NK cells | 15 | ENSMUSG00000033220 | Rac2 | 0.14131304 |
| 327 | NK cells | 15 | ENSMUSG00000044199 | S1pr4 | 0.134271778 |
| 328 | NK cells | 15 | ENSMUSG00000050241 | Klre1 | 0.18809632 |
| 329 | NK cells | 15 | ENSMUSG00000050232 | Cxcr3 | 0.20377295 |
| 330 | NK cells | 15 | ENSMUSG00000052837 | Junb | 0.159765839 |
| 331 | NK cells | 15 | ENSMUSG00000054435 | Gimap4 | 0.180466894 |
| 332 | NK cells | 15 | ENSMUSG00000055170 | Ifng | 0.16672972 |
| 333 | NK cells | 15 | ENSMUSG00000004612 | Nkg7 | 0.225710028 |
| 334 | NK cells | 15 | ENSMUSG00000052736 | Klrc2 | 0.203585871 |
| 335 | NK cells | 15 | ENSMUSG00000060550 | H2-Q7 | 0.183898831 |
| 336 | NK cells | 15 | ENSMUSG00000064109 | Hcst | 0.22562366 |
| 337 | NK cells | 15 | ENSMUSG00000057841 | Rpl32 | 0.133985422 |
| 338 | NK cells | 15 | ENSMUSG00000025508 | Rplp2 | 0.139410216 |
| 339 | NK cells | 15 | ENSMUSG00000067274 | Rplp0 | 0.142744081 |
| 340 | NK cells | 15 | ENSMUSG00000068227 | Il2rb | 0.187526529 |
| 341 | NK cells | 15 | ENSMUSG00000075010 | AW112010 | 0.214953315 |
| 342 | NK cells | 15 | ENSMUSG00000048163 | Selplg | 0.161146322 |
| 343 | NK cells | 15 | ENSMUSG00000076490 | Trbc1 | 0.145247786 |
| 344 | NK cells | 15 | ENSMUSG00000076498 | Trbc2 | 0.260249737 |
| 345 | NK cells | 15 | ENSMUSG00000076928 | Trac | 0.153103499 |
| 346 | NK cells | 15 | ENSMUSG00000030830 | Itgal | 0.131326517 |
| 347 | NK cells | 15 | ENSMUSG00000000486 | 1-Sep | 0.205604287 |
| 348 | NK cells | 15 | ENSMUSG00000079523 | Tmsb10 | 0.177305793 |
| 349 | NK cells | 15 | ENSMUSG00000030178 | Klra13-ps | 0.131578947 |
| 350 | NK cells | 15 | ENSMUSG00000030325 | Klrb1c | 0.174768082 |
| 351 | Macrophage-C | 16 | ENSMUSG00000000078 | Klf6 | 0.15917113 |
| 352 | Macrophage-C | 16 | ENSMUSG00000000594 | Gm2a | 0.364234188 |
| 353 | Macrophage-C | 16 | ENSMUSG00000000682 | Cd52 | 0.156408379 |
| 354 | Macrophage-C | 16 | ENSMUSG00000001020 | S100a4 | 0.148212312 |
| 355 | Macrophage-C | 16 | ENSMUSG00000001281 | Itgb7 | 0.206128659 |
| 356 | Macrophage-C | 16 | ENSMUSG00000002204 | Napsa | 0.278308664 |
| 357 | Macrophage-C | 16 | ENSMUSG00000004207 | Psap | 0.202694454 |
| 358 | Macrophage-C | 16 | ENSMUSG00000009687 | Fxyd5 | 0.165086943 |
| 359 | Macrophage-C | 16 | ENSMUSG00000031494 | Cd209a | 0.458196848 |

| | | | | | |
|---|---|---|---|---|---|
| 360 | Macrophage-C | 16 | ENSMUSG00000038642 | Ctss | 0.148490562 |
| 361 | Macrophage-C | 16 | ENSMUSG00000018819 | Lsp1 | 0.224005394 |
| 362 | Macrophage-C | 16 | ENSMUSG00000019122 | Ccl9 | 0.154653396 |
| 363 | Macrophage-C | 16 | ENSMUSG00000022205 | Sub1 | 0.169716045 |
| 364 | Macrophage-C | 16 | ENSMUSG00000061100 | Retnla | 0.188101006 |
| 365 | Macrophage-C | 16 | ENSMUSG00000024014 | Pim1 | 0.148748491 |
| 366 | Macrophage-C | 16 | ENSMUSG00000024521 | Pmaip1 | 0.231664279 |
| 367 | Macrophage-C | 16 | ENSMUSG00000026832 | Cytip | 0.160534469 |
| 368 | Macrophage-C | 16 | ENSMUSG00000027398 | Il1b | 0.234010412 |
| 369 | Macrophage-C | 16 | ENSMUSG00000027447 | Cst3 | 0.209262245 |
| 370 | Macrophage-C | 16 | ENSMUSG00000028581 | Laptm5 | 0.145358145 |
| 371 | Macrophage-C | 16 | ENSMUSG00000028843 | Sh3bgrl3 | 0.159831931 |
| 372 | Macrophage-C | 16 | ENSMUSG00000030214 | Plbd1 | 0.280479126 |
| 373 | Macrophage-C | 16 | ENSMUSG00000030707 | Coro1a | 0.161215097 |
| 374 | Macrophage-C | 16 | ENSMUSG00000031780 | Ccl17 | 0.199976081 |
| 375 | Macrophage-C | 16 | ENSMUSG00000040659 | Efhd2 | 0.158876009 |
| 376 | Macrophage-C | 16 | ENSMUSG00000070868 | Skint3 | 0.195095342 |
| 377 | Macrophage-C | 16 | ENSMUSG00000039783 | Kmo | 0.16964068 |
| 378 | Macrophage-C | 16 | ENSMUSG00000036594 | H2-Aa | 0.21798523 |
| 379 | Macrophage-C | 16 | ENSMUSG00000073421 | H2-Ab1 | 0.211306641 |
| 380 | Macrophage-C | 16 | ENSMUSG00000037649 | H2-DMa | 0.279533731 |
| 381 | Macrophage-C | 16 | ENSMUSG00000042817 | Flt3 | 0.160420333 |
| 382 | Macrophage-C | 16 | ENSMUSG00000044156 | Hepacam2 | 0.273039963 |
| 383 | Macrophage-C | 16 | ENSMUSG00000024610 | Cd74 | 0.219853248 |
| 384 | Macrophage-C | 16 | ENSMUSG00000046908 | Ltb4r1 | 0.199701146 |
| 385 | Macrophage-C | 16 | ENSMUSG00000062825 | Actg1 | 0.173345625 |
| 386 | Macrophage-C | 16 | ENSMUSG00000019876 | Pkib | 0.18911187 |
| 387 | Macrophage-C | 16 | ENSMUSG00000060063 | Alox5ap | 0.249449114 |
| 388 | Macrophage-C | 16 | ENSMUSG00000060586 | H2-Eb1 | 0.195731023 |
| 389 | Macrophage-C | 16 | ENSMUSG00000030147 | Clec4b1 | 0.15019569 |
| 390 | Macrophage-C | 16 | ENSMUSG00000059108 | Ifitm6 | 0.193094089 |
| 391 | Macrophage-C | 16 | ENSMUSG00000063856 | Gpx1 | 0.189091691 |
| 392 | Macrophage-C | 16 | ENSMUSG00000021880 | Rnase6 | 0.269357164 |
| 393 | Macrophage-C | 16 | ENSMUSG00000099974 | Bcl2a1d | 0.211201151 |
| 394 | Macrophage-C | 16 | ENSMUSG00000031097 | Tnni2 | 0.186684779 |
| 395 | Macrophage-C | 16 | ENSMUSG00000018585 | Atox1 | 0.161627372 |
| 396 | Macrophage-C | 16 | ENSMUSG00000038179 | Slamf7 | 0.178075856 |
| 397 | Macrophage-C | 16 | ENSMUSG00000029413 | Naaa | 0.232414278 |
| 398 | Macrophage-C | 16 | ENSMUSG00000079547 | H2-DMb1 | 0.182261503 |
| 399 | Macrophage-C | 16 | ENSMUSG00000044162 | Tnip3 | 0.149664487 |
| 400 | Macrophage-C | 16 | ENSMUSG00000050335 | Lgals3 | 0.183191533 |
| 401 | Smooth muscle cells B | 17 | ENSMUSG00000020676 | Ccl11 | 0.391767103 |
| 402 | Smooth muscle cells B | 17 | ENSMUSG00000000359 | Rem1 | 0.175510544 |
| 403 | Smooth muscle cells B | 17 | ENSMUSG00000004558 | Ndrg2 | 0.172738214 |
| 404 | Smooth muscle cells B | 17 | ENSMUSG00000009876 | Cox4i2 | 0.54700468 |

| | | | | | |
|---|---|---|---|---|---|
| 405 | Smooth muscle cells B | 17 | ENSMUSG00000017344 | Vtn | 0.642336509 |
| 406 | Smooth muscle cells B | 17 | ENSMUSG00000017417 | Plxdc1 | 0.267688969 |
| 407 | Smooth muscle cells B | 17 | ENSMUSG00000020486 | 4-Sep | 0.23718519 |
| 408 | Smooth muscle cells B | 17 | ENSMUSG00000018417 | Myo1b | 0.22037543 |
| 409 | Smooth muscle cells B | 17 | ENSMUSG00000018593 | Sparc | 0.259593601 |
| 410 | Smooth muscle cells B | 17 | ENSMUSG00000019906 | Lin7a | 0.151958431 |
| 411 | Smooth muscle cells B | 17 | ENSMUSG00000020585 | Laptm4a | 0.159551265 |
| 412 | Smooth muscle cells B | 17 | ENSMUSG00000020928 | Higd1b | 0.569348155 |
| 413 | Smooth muscle cells B | 17 | ENSMUSG00000021388 | Aspn | 0.158676457 |
| 414 | Smooth muscle cells B | 17 | ENSMUSG00000022010 | Tsc22d1 | 0.188270697 |
| 415 | Smooth muscle cells B | 17 | ENSMUSG00000024620 | Pdgfrb | 0.418577175 |
| 416 | Smooth muscle cells B | 17 | ENSMUSG00000025351 | Cd63 | 0.150754744 |
| 417 | Smooth muscle cells B | 17 | ENSMUSG00000025491 | Ifitm1 | 0.41061445 |
| 418 | Smooth muscle cells B | 17 | ENSMUSG00000038530 | Rgs4 | 0.279988429 |
| 419 | Smooth muscle cells B | 17 | ENSMUSG00000026678 | Rgs5 | 0.532774491 |
| 420 | Smooth muscle cells B | 17 | ENSMUSG00000028005 | Gucy1b1 | 0.162976633 |
| 421 | Smooth muscle cells B | 17 | ENSMUSG00000028967 | Errfi1 | 0.200043532 |
| 422 | Smooth muscle cells B | 17 | ENSMUSG00000029309 | Sparcl1 | 0.166777992 |
| 423 | Smooth muscle cells B | 17 | ENSMUSG00000029641 | Rasl11a | 0.18581397 |
| 424 | Smooth muscle cells B | 17 | ENSMUSG00000032766 | Gng11 | 0.192648898 |
| 425 | Smooth muscle cells B | 17 | ENSMUSG00000029761 | Cald1 | 0.239258347 |
| 426 | Smooth muscle cells B | 17 | ENSMUSG00000030247 | Kcnj8 | 0.328472071 |
| 427 | Smooth muscle cells B | 17 | ENSMUSG00000030605 | Mfge8 | 0.443328018 |
| 428 | Smooth muscle cells B | 17 | ENSMUSG00000031503 | Col4a2 | 0.202270409 |
| 429 | Smooth muscle cells B | 17 | ENSMUSG00000040280 | Ndufa4l2 | 0.705865573 |
| 430 | Smooth muscle cells B | 17 | ENSMUSG00000036256 | Igfbp7 | 0.225105297 |
| 431 | Smooth muscle cells B | 17 | ENSMUSG00000033910 | Gucy1a1 | 0.156134033 |
| 432 | Smooth muscle cells B | 17 | ENSMUSG00000042155 | Klhl23 | 0.16093222 |
| 433 | Smooth muscle cells B | 17 | ENSMUSG00000050666 | Vstm4 | 0.168479764 |
| 434 | Smooth muscle cells B | 17 | ENSMUSG00000050234 | Gja4 | 0.159378009 |
| 435 | Smooth muscle cells B | 17 | ENSMUSG00000048332 | Lhfp | 0.232725606 |
| 436 | Smooth muscle cells B | 17 | ENSMUSG00000042284 | Itga1 | 0.200501849 |
| 437 | Smooth muscle cells B | 17 | ENSMUSG00000046275 | Trarg1 | 0.271190002 |
| 438 | Smooth muscle cells B | 17 | ENSMUSG00000034164 | Emid1 | 0.206937417 |
| 439 | Smooth muscle cells B | 17 | ENSMUSG00000034520 | Gjc1 | 0.208679356 |
| 440 | Smooth muscle cells B | 17 | ENSMUSG00000030249 | Abcc9 | 0.272040939 |
| 441 | Smooth muscle cells B | 17 | ENSMUSG00000031750 | Il34 | 0.15022105 |
| 442 | Smooth muscle cells B | 17 | ENSMUSG00000057098 | Ebf1 | 0.238966556 |
| 443 | Smooth muscle cells B | 17 | ENSMUSG00000067818 | Myl9 | 0.21894564 |
| 444 | Smooth muscle cells B | 17 | ENSMUSG00000069763 | Tmem100 | 0.264363885 |
| 445 | Smooth muscle cells B | 17 | ENSMUSG00000074676 | Foxs1 | 0.173013666 |
| 446 | Smooth muscle cells B | 17 | ENSMUSG00000028713 | Cyp4b1 | 0.187897263 |
| 447 | Smooth muscle cells B | 17 | ENSMUSG00000032875 | Arhgef17 | 0.162334427 |
| 448 | Smooth muscle cells B | 17 | ENSMUSG00000034842 | Art3 | 0.156425373 |
| 449 | Smooth muscle cells B | 17 | ENSMUSG00000001833 | 7-Sep | 0.173328025 |

| | | | | | |
|---|---|---|---|---|---|
| 450 | Smooth muscle cells B | 17 | ENSMUSG00000012428 | Steap4 | 0.373045846 |
| 451 | Macrophage-B | 18 | ENSMUSG00000029925 | Tbxas1 | 0.164548838 |
| 452 | Macrophage-B | 18 | ENSMUSG00000002985 | Apoe | 0.14383609 |
| 453 | Macrophage-B | 18 | ENSMUSG00000004730 | Adgre1 | 0.151762812 |
| 454 | Macrophage-B | 18 | ENSMUSG00000021939 | Ctsb | 0.153054081 |
| 455 | Macrophage-B | 18 | ENSMUSG00000038642 | Ctss | 0.182994344 |
| 456 | Macrophage-B | 18 | ENSMUSG00000017009 | Sdc4 | 0.146665704 |
| 457 | Macrophage-B | 18 | ENSMUSG00000021190 | Lgmn | 0.194287656 |
| 458 | Macrophage-B | 18 | ENSMUSG00000021423 | Ly86 | 0.156137086 |
| 459 | Macrophage-B | 18 | ENSMUSG00000022324 | Matn2 | 0.232501048 |
| 460 | Macrophage-B | 18 | ENSMUSG00000024397 | Aif1 | 0.163668951 |
| 461 | Macrophage-B | 18 | ENSMUSG00000024621 | Csf1r | 0.162395373 |
| 462 | Macrophage-B | 18 | ENSMUSG00000024672 | Ms4a7 | 0.214881057 |
| 463 | Macrophage-B | 18 | ENSMUSG00000024677 | Ms4a6b | 0.146344818 |
| 464 | Macrophage-B | 18 | ENSMUSG00000025232 | Hexa | 0.157215381 |
| 465 | Macrophage-B | 18 | ENSMUSG00000026385 | Dbi | 0.160012846 |
| 466 | Macrophage-B | 18 | ENSMUSG00000026424 | Gpr37l1 | 0.183694095 |
| 467 | Macrophage-B | 18 | ENSMUSG00000026656 | Fcgr2b | 0.164868717 |
| 468 | Macrophage-B | 18 | ENSMUSG00000026712 | Mrc1 | 0.225283673 |
| 469 | Macrophage-B | 18 | ENSMUSG00000008845 | Cd163 | 0.161822101 |
| 470 | Macrophage-B | 18 | ENSMUSG00000030560 | Ctsc | 0.191000634 |
| 471 | Macrophage-B | 18 | ENSMUSG00000031425 | Plp1 | 0.220256603 |
| 472 | Macrophage-B | 18 | ENSMUSG00000032060 | Cryab | 0.168480717 |
| 473 | Macrophage-B | 18 | ENSMUSG00000032359 | Ctsh | 0.165063947 |
| 474 | Macrophage-B | 18 | ENSMUSG00000032554 | Trf | 0.146674372 |
| 475 | Macrophage-B | 18 | ENSMUSG00000040759 | Cmtm5 | 0.156455222 |
| 476 | Macrophage-B | 18 | ENSMUSG00000036570 | Fxyd1 | 0.159291697 |
| 477 | Macrophage-B | 18 | ENSMUSG00000036594 | H2-Aa | 0.170540864 |
| 478 | Macrophage-B | 18 | ENSMUSG00000073421 | H2-Ab1 | 0.174604657 |
| 479 | Macrophage-B | 18 | ENSMUSG00000036169 | Sostdc1 | 0.162581335 |
| 480 | Macrophage-B | 18 | ENSMUSG00000040950 | Mgl2 | 0.154503174 |
| 481 | Macrophage-B | 18 | ENSMUSG00000034810 | Scn7a | 0.169245394 |
| 482 | Macrophage-B | 18 | ENSMUSG00000036887 | C1qa | 0.194776478 |
| 483 | Macrophage-B | 18 | ENSMUSG00000036896 | C1qc | 0.188564639 |
| 484 | Macrophage-B | 18 | ENSMUSG00000036905 | C1qb | 0.190778828 |
| 485 | Macrophage-B | 18 | ENSMUSG00000024610 | Cd74 | 0.159958856 |
| 486 | Macrophage-B | 18 | ENSMUSG00000049130 | C5ar1 | 0.211260033 |
| 487 | Macrophage-B | 18 | ENSMUSG00000047976 | Kcna1 | 0.184012738 |
| 488 | Macrophage-B | 18 | ENSMUSG00000051439 | Cd14 | 0.151659381 |
| 489 | Macrophage-B | 18 | ENSMUSG00000055541 | Lair1 | 0.210437644 |
| 490 | Macrophage-B | 18 | ENSMUSG00000060586 | H2-Eb1 | 0.175681172 |
| 491 | Macrophage-B | 18 | ENSMUSG00000058715 | Fcer1g | 0.151554004 |
| 492 | Macrophage-B | 18 | ENSMUSG00000067261 | Foxd3 | 0.151962541 |
| 493 | Macrophage-B | 18 | ENSMUSG00000069662 | Marcks | 0.154621904 |
| 494 | Macrophage-B | 18 | ENSMUSG00000008496 | Pou2f2 | 0.165336714 |

| | | | | | |
|---|---|---|---|---|---|
| 495 | Macrophage-B | 18 | ENSMUSG00000074361 | C5ar2 | 0.148111783 |
| 496 | Macrophage-B | 18 | ENSMUSG00000074622 | Mafb | 0.178188104 |
| 497 | Macrophage-B | 18 | ENSMUSG00000039542 | Ncam1 | 0.197077669 |
| 498 | Macrophage-B | 18 | ENSMUSG00000090164 | BC035044 | 0.143062908 |
| 499 | Macrophage-B | 18 | ENSMUSG00000036908 | Unc93b1 | 0.156762111 |
| 500 | Macrophage-B | 18 | ENSMUSG00000059498 | Fcgr3 | 0.150220462 |
| 501 | Unknown | 19 | ENSMUSG00000000308 | Ckmt1 | 0.412513272 |
| 502 | Unknown | 19 | ENSMUSG00000006345 | Ggt1 | 0.404720799 |
| 503 | Unknown | 19 | ENSMUSG00000011034 | Slc5a1 | 0.411815352 |
| 504 | Unknown | 19 | ENSMUSG00000053054 | Adh6a | 0.432574533 |
| 505 | Unknown | 19 | ENSMUSG00000035775 | Krt20 | 0.573007413 |
| 506 | Unknown | 19 | ENSMUSG00000018569 | Cldn7 | 0.523125511 |
| 507 | Unknown | 19 | ENSMUSG00000022555 | Dgat1 | 0.461767941 |
| 508 | Unknown | 19 | ENSMUSG00000022824 | Muc13 | 0.667315253 |
| 509 | Unknown | 19 | ENSMUSG00000023057 | Fabp2 | 0.676126782 |
| 510 | Unknown | 19 | ENSMUSG00000049382 | Krt8 | 0.612199439 |
| 511 | Unknown | 19 | ENSMUSG00000024503 | Spink1 | 0.721401631 |
| 512 | Unknown | 19 | ENSMUSG00000024712 | Rfk | 0.430323386 |
| 513 | Unknown | 19 | ENSMUSG00000025467 | Prap1 | 0.455731333 |
| 514 | Unknown | 19 | ENSMUSG00000026175 | Vil1 | 0.414393112 |
| 515 | Unknown | 19 | ENSMUSG00000026417 | Pigr | 0.421375983 |
| 516 | Unknown | 19 | ENSMUSG00000000544 | Gpa33 | 0.434109373 |
| 517 | Unknown | 19 | ENSMUSG00000028158 | Mttp | 0.385032913 |
| 518 | Unknown | 19 | ENSMUSG00000028307 | Aldob | 0.603625128 |
| 519 | Unknown | 19 | ENSMUSG00000029161 | Cgref1 | 0.38848262 |
| 520 | Unknown | 19 | ENSMUSG00000029269 | Sult1b1 | 0.393202557 |
| 521 | Unknown | 19 | ENSMUSG00000029727 | Cyp3a13 | 0.43048825 |
| 522 | Unknown | 19 | ENSMUSG00000030017 | Reg3g | 0.489237443 |
| 523 | Unknown | 19 | ENSMUSG00000030364 | Clec2h | 0.534644372 |
| 524 | Unknown | 19 | ENSMUSG00000030587 | 2200002D01Rik | 0.670763634 |
| 525 | Unknown | 19 | ENSMUSG00000055730 | Ces2a | 0.394742716 |
| 526 | Unknown | 19 | ENSMUSG00000031886 | Ces2e | 0.451639113 |
| 527 | Unknown | 19 | ENSMUSG00000032081 | Apoc3 | 0.567543454 |
| 528 | Unknown | 19 | ENSMUSG00000032083 | Apoa1 | 0.721611108 |
| 529 | Unknown | 19 | ENSMUSG00000032454 | Rbp2 | 0.753873372 |
| 530 | Unknown | 19 | ENSMUSG00000020609 | Apob | 0.57095057 |
| 531 | Unknown | 19 | ENSMUSG00000032978 | Guca2b | 0.563423448 |
| 532 | Unknown | 19 | ENSMUSG00000054999 | Naaladl1 | 0.394041902 |
| 533 | Unknown | 19 | ENSMUSG00000021208 | Ifi27l2b | 0.380736074 |
| 534 | Unknown | 19 | ENSMUSG00000039062 | Anpep | 0.407501223 |
| 535 | Unknown | 19 | ENSMUSG00000045394 | Epcam | 0.544651655 |
| 536 | Unknown | 19 | ENSMUSG00000050982 | Apol10a | 0.378116156 |
| 537 | Unknown | 19 | ENSMUSG00000046804 | Phgr1 | 0.709692986 |
| 538 | Unknown | 19 | ENSMUSG00000053964 | Lgals4 | 0.749900734 |
| 539 | Unknown | 19 | ENSMUSG00000054422 | Fabp1 | 0.515725926 |

| | | | | | |
|---|---|---|---|---|---|
| 540 | Unknown | 19 | ENSMUSG00000091705 | H2-Q2 | 0.490541394 |
| 541 | Unknown | 19 | ENSMUSG00000003271 | Sult2b1 | 0.401263523 |
| 542 | Unknown | 19 | ENSMUSG00000025401 | Myo1a | 0.462377694 |
| 543 | Unknown | 19 | ENSMUSG00000025497 | Cdhr5 | 0.455279047 |
| 544 | Unknown | 19 | ENSMUSG00000049493 | Pls1 | 0.534009428 |
| 545 | Unknown | 19 | ENSMUSG00000071356 | Reg3b | 0.492624851 |
| 546 | Unknown | 19 | ENSMUSG00000078439 | Smim24 | 0.546895264 |
| 547 | Unknown | 19 | ENSMUSG00000029273 | Sult1d1 | 0.381459086 |
| 548 | Unknown | 19 | ENSMUSG00000035041 | Creb3l3 | 0.445246243 |
| 549 | Unknown | 19 | ENSMUSG00000085042 | Abhd11os | 0.486833814 |
| 550 | Unknown | 19 | ENSMUSG00000096215 | Smim22 | 0.44332427 |
| 551 | MENs | 2 | ENSMUSG00000001025 | S100a6 | 0.289749472 |
| 552 | MENs | 2 | ENSMUSG00000001739 | Cldn15 | 0.218669778 |
| 553 | MENs | 2 | ENSMUSG00000002980 | Bcam | 0.22759976 |
| 554 | MENs | 2 | ENSMUSG00000006360 | Crip1 | 0.261785689 |
| 555 | MENs | 2 | ENSMUSG00000045545 | Krt14 | 0.199939315 |
| 556 | MENs | 2 | ENSMUSG00000020911 | Krt19 | 0.327618161 |
| 557 | MENs | 2 | ENSMUSG00000008540 | Mgst1 | 0.227927609 |
| 558 | MENs | 2 | ENSMUSG00000009281 | Rarres2 | 0.396915172 |
| 559 | MENs | 2 | ENSMUSG00000020467 | Efemp1 | 0.24391868 |
| 560 | MENs | 2 | ENSMUSG00000022037 | Clu | 0.3046635 |
| 561 | MENs | 2 | ENSMUSG00000023043 | Krt18 | 0.205838072 |
| 562 | MENs | 2 | ENSMUSG00000023046 | Igfbp6 | 0.469623182 |
| 563 | MENs | 2 | ENSMUSG00000024164 | C3 | 0.434183264 |
| 564 | MENs | 2 | ENSMUSG00000090231 | Cfb | 0.21623191 |
| 565 | MENs | 2 | ENSMUSG00000024659 | Anxa1 | 0.17927024 |
| 566 | MENs | 2 | ENSMUSG00000025867 | Cplx2 | 0.208119555 |
| 567 | MENs | 2 | ENSMUSG00000053897 | Slc39a8 | 0.245316246 |
| 568 | MENs | 2 | ENSMUSG00000028583 | Pdpn | 0.195254698 |
| 569 | MENs | 2 | ENSMUSG00000028780 | Sema3c | 0.20563504 |
| 570 | MENs | 2 | ENSMUSG00000028871 | Rspo1 | 0.340066877 |
| 571 | MENs | 2 | ENSMUSG00000029718 | Pcolce | 0.245733541 |
| 572 | MENs | 2 | ENSMUSG00000031517 | Gpm6a | 0.383029893 |
| 573 | MENs | 2 | ENSMUSG00000013584 | Aldh1a2 | 0.23886889 |
| 574 | MENs | 2 | ENSMUSG00000032348 | Gsta4 | 0.20392163 |
| 575 | MENs | 2 | ENSMUSG00000035413 | Tmem98 | 0.222123141 |
| 576 | MENs | 2 | ENSMUSG00000060962 | Dmkn | 0.211834864 |
| 577 | MENs | 2 | ENSMUSG00000040170 | Fmo2 | 0.293138471 |
| 578 | MENs | 2 | ENSMUSG00000032902 | Slc16a1 | 0.202786328 |
| 579 | MENs | 2 | ENSMUSG00000042357 | Gjb5 | 0.222578735 |
| 580 | MENs | 2 | ENSMUSG00000040605 | Bace2 | 0.219405763 |
| 581 | MENs | 2 | ENSMUSG00000041559 | Fmod | 0.194981957 |
| 582 | MENs | 2 | ENSMUSG00000043110 | Lrrn4 | 0.305113131 |
| 583 | MENs | 2 | ENSMUSG00000049436 | Upk1b | 0.335803791 |
| 584 | MENs | 2 | ENSMUSG00000042985 | Upk3b | 0.415997736 |

| | | | | | |
|---|---|---|---|---|---|
| 585 | MENs | 2 | ENSMUSG00000021994 | Wnt5a | 0.190898755 |
| 586 | MENs | 2 | ENSMUSG00000052957 | Gas1 | 0.221432763 |
| 587 | MENs | 2 | ENSMUSG00000055172 | C1ra | 0.178873421 |
| 588 | MENs | 2 | ENSMUSG00000023039 | Krt7 | 0.340896185 |
| 589 | MENs | 2 | ENSMUSG00000055653 | Gpc3 | 0.268390939 |
| 590 | MENs | 2 | ENSMUSG00000063011 | Msln | 0.397760502 |
| 591 | MENs | 2 | ENSMUSG00000046056 | Sbsn | 0.1765033 |
| 592 | MENs | 2 | ENSMUSG00000018339 | Gpx3 | 0.249410016 |
| 593 | MENs | 2 | ENSMUSG00000015093 | Clic3 | 0.186720351 |
| 594 | MENs | 2 | ENSMUSG00000020473 | Aebp1 | 0.381803898 |
| 595 | MENs | 2 | ENSMUSG00000027574 | Nkain4 | 0.281806522 |
| 596 | MENs | 2 | ENSMUSG00000019929 | Dcn | 0.273470348 |
| 597 | MENs | 2 | ENSMUSG00000035930 | Chst4 | 0.251966385 |
| 598 | MENs | 2 | ENSMUSG00000017002 | Slpi | 0.431451583 |
| 599 | MENs | 2 | ENSMUSG00000016458 | Wt1 | 0.206402807 |
| 600 | MENs | 2 | ENSMUSG00000078350 | Smim1 | 0.203013948 |
| 601 | NENs | 20 | ENSMUSG00000002108 | Nr1h3 | 0.192370624 |
| 602 | NENs | 20 | ENSMUSG00000031849 | Comp | 0.238905868 |
| 603 | NENs | 20 | ENSMUSG00000003617 | Cp | 0.215225768 |
| 604 | NENs | 20 | ENSMUSG00000003849 | Nqo1 | 0.187978055 |
| 605 | NENs | 20 | ENSMUSG00000004771 | Rab11a | 0.19109985 |
| 606 | NENs | 20 | ENSMUSG00000009097 | Tbx1 | 0.465492021 |
| 607 | NENs | 20 | ENSMUSG00000010175 | Prox1 | 0.500433128 |
| 608 | NENs | 20 | ENSMUSG00000016024 | Lbp | 0.407606408 |
| 609 | NENs | 20 | ENSMUSG00000019890 | Nts | 0.4574443 |
| 610 | NENs | 20 | ENSMUSG00000020357 | Flt4 | 0.322381555 |
| 611 | NENs | 20 | ENSMUSG00000024210 | Ip6k3 | 0.210526316 |
| 612 | NENs | 20 | ENSMUSG00000026185 | Igfbp5 | 0.286050322 |
| 613 | NENs | 20 | ENSMUSG00000056025 | Clca3a1 | 0.288485936 |
| 614 | NENs | 20 | ENSMUSG00000032766 | Gng11 | 0.190673251 |
| 615 | NENs | 20 | ENSMUSG00000030342 | Cd9 | 0.20629862 |
| 616 | NENs | 20 | ENSMUSG00000030638 | Sh3gl3 | 0.449334545 |
| 617 | NENs | 20 | ENSMUSG00000030787 | Lyve1 | 0.44335455 |
| 618 | NENs | 20 | ENSMUSG00000042286 | Stab1 | 0.194418425 |
| 619 | NENs | 20 | ENSMUSG00000040488 | Ltbp4 | 0.255566563 |
| 620 | NENs | 20 | ENSMUSG00000025597 | Klhl4 | 0.284309148 |
| 621 | NENs | 20 | ENSMUSG00000037095 | Lrg1 | 0.254202185 |
| 622 | NENs | 20 | ENSMUSG00000041378 | Cldn5 | 0.436596966 |
| 623 | NENs | 20 | ENSMUSG00000037625 | Cldn11 | 0.324405615 |
| 624 | NENs | 20 | ENSMUSG00000038296 | Galnt18 | 0.276660984 |
| 625 | NENs | 20 | ENSMUSG00000026822 | Lcn2 | 0.513447471 |
| 626 | NENs | 20 | ENSMUSG00000060470 | Adgrg3 | 0.214857423 |
| 627 | NENs | 20 | ENSMUSG00000045954 | Cavin2 | 0.193122385 |
| 628 | NENs | 20 | ENSMUSG00000050447 | Lypd6 | 0.189822234 |
| 629 | NENs | 20 | ENSMUSG00000044921 | Rassf9 | 0.193674441 |

| | | | | | |
|---|---|---|---|---|---|
| 630 | NENs | 20 | ENSMUSG00000001493 | Meox1 | 0.207660507 |
| 631 | NENs | 20 | ENSMUSG00000043342 | Hoxd9 | 0.359381288 |
| 632 | NENs | 20 | ENSMUSG00000047714 | Ppp1r2 | 0.201790096 |
| 633 | NENs | 20 | ENSMUSG00000042453 | Reln | 0.611438603 |
| 634 | NENs | 20 | ENSMUSG00000052861 | Dnah6 | 0.275745339 |
| 635 | NENs | 20 | ENSMUSG00000056214 | Pard6g | 0.200886818 |
| 636 | NENs | 20 | ENSMUSG00000058396 | Gpr182 | 0.273218454 |
| 637 | NENs | 20 | ENSMUSG00000034684 | Sema3f | 0.245351594 |
| 638 | NENs | 20 | ENSMUSG00000066705 | Fxyd6 | 0.216246452 |
| 639 | NENs | 20 | ENSMUSG00000094686 | Ccl21a | 0.194544495 |
| 640 | NENs | 20 | ENSMUSG00000073599 | Ecscr | 0.204168114 |
| 641 | NENs | 20 | ENSMUSG00000074345 | Tnfaip8l3 | 0.203411063 |
| 642 | NENs | 20 | ENSMUSG00000057530 | Ece1 | 0.271284907 |
| 643 | NENs | 20 | ENSMUSG00000028713 | Cyp4b1 | 0.229765867 |
| 644 | NENs | 20 | ENSMUSG00000031377 | Bmx | 0.231382292 |
| 645 | NENs | 20 | ENSMUSG00000062960 | Kdr | 0.199814165 |
| 646 | NENs | 20 | ENSMUSG00000054641 | Mmrn1 | 0.872887801 |
| 647 | NENs | 20 | ENSMUSG00000079654 | Prrt4 | 0.223616907 |
| 648 | NENs | 20 | ENSMUSG00000090698 | Apold1 | 0.297182535 |
| 649 | NENs | 20 | ENSMUSG00000094707 | A830019P07Rik | 0.191723011 |
| 650 | NENs | 20 | ENSMUSG00000098132 | Rassf10 | 0.2544539 |
| 651 | Macrophage-C | 21 | ENSMUSG00000000290 | Itgb2 | 0.384096675 |
| 652 | Macrophage-C | 21 | ENSMUSG00000001029 | Icam2 | 0.294705301 |
| 653 | Macrophage-C | 21 | ENSMUSG00000001128 | Cfp | 0.532730909 |
| 654 | Macrophage-C | 21 | ENSMUSG00000001870 | Ltbp1 | 0.27281908 |
| 655 | Macrophage-C | 21 | ENSMUSG00000004814 | Ccl24 | 0.378723999 |
| 656 | Macrophage-C | 21 | ENSMUSG00000005373 | Mlxipl | 0.24095132 |
| 657 | Macrophage-C | 21 | ENSMUSG00000006014 | Prg4 | 0.778175835 |
| 658 | Macrophage-C | 21 | ENSMUSG00000057729 | Prtn3 | 0.734345385 |
| 659 | Macrophage-C | 21 | ENSMUSG00000040026 | Saa3 | 0.707182673 |
| 660 | Macrophage-C | 21 | ENSMUSG00000010797 | Wnt2 | 0.26372524 |
| 661 | Macrophage-C | 21 | ENSMUSG00000013974 | Mcemp1 | 0.545270394 |
| 662 | Macrophage-C | 21 | ENSMUSG00000015340 | Cybb | 0.267167648 |
| 663 | Macrophage-C | 21 | ENSMUSG00000015854 | Cd5l | 0.89408571 |
| 664 | Macrophage-C | 21 | ENSMUSG00000018924 | Alox15 | 0.767090644 |
| 665 | Macrophage-C | 21 | ENSMUSG00000019122 | Ccl9 | 0.31122662 |
| 666 | Macrophage-C | 21 | ENSMUSG00000019872 | Smpdl3a | 0.241205757 |
| 667 | Macrophage-C | 21 | ENSMUSG00000020120 | Plek | 0.298883962 |
| 668 | Macrophage-C | 21 | ENSMUSG00000022122 | Ednrb | 0.353890352 |
| 669 | Macrophage-C | 21 | ENSMUSG00000024730 | Ms4a8a | 0.248567092 |
| 670 | Macrophage-C | 21 | ENSMUSG00000025330 | Padi4 | 0.682255492 |
| 671 | Macrophage-C | 21 | ENSMUSG00000025534 | Gusb | 0.260385496 |
| 672 | Macrophage-C | 21 | ENSMUSG00000026357 | Rgs18 | 0.302654936 |
| 673 | Macrophage-C | 21 | ENSMUSG00000028108 | Ecm1 | 0.255382941 |
| 674 | Macrophage-C | 21 | ENSMUSG00000040264 | Gbp2b | 0.286089191 |

| | | | | | |
|---|---|---|---|---|---|
| 675 | Macrophage-C | 21 | ENSMUSG00000029373 | Pf4 | 0.306387312 |
| 676 | Macrophage-C | 21 | ENSMUSG00000096630 | Vmn2r26 | 0.42068113 |
| 677 | Macrophage-C | 21 | ENSMUSG00000030142 | Clec4e | 0.59143538 |
| 678 | Macrophage-C | 21 | ENSMUSG00000030144 | Clec4d | 0.349882953 |
| 679 | Macrophage-C | 21 | ENSMUSG00000030793 | Pycard | 0.30273585 |
| 680 | Macrophage-C | 21 | ENSMUSG00000031444 | F10 | 0.723142185 |
| 681 | Macrophage-C | 21 | ENSMUSG00000040152 | Thbs1 | 0.347266669 |
| 682 | Macrophage-C | 21 | ENSMUSG00000038860 | Garnl3 | 0.242501452 |
| 683 | Macrophage-C | 21 | ENSMUSG00000026193 | Fn1 | 0.553745128 |
| 684 | Macrophage-C | 21 | ENSMUSG00000046245 | Pilra | 0.247218805 |
| 685 | Macrophage-C | 21 | ENSMUSG00000030786 | Itgam | 0.333812733 |
| 686 | Macrophage-C | 21 | ENSMUSG00000055546 | Timd4 | 0.498308518 |
| 687 | Macrophage-C | 21 | ENSMUSG00000060063 | Alox5ap | 0.320290682 |
| 688 | Macrophage-C | 21 | ENSMUSG00000021069 | Pygl | 0.340209147 |
| 689 | Macrophage-C | 21 | ENSMUSG00000058427 | Cxcl2 | 0.33517704 |
| 690 | Macrophage-C | 21 | ENSMUSG00000059108 | Ifitm6 | 0.389815558 |
| 691 | Macrophage-C | 21 | ENSMUSG00000026579 | F5 | 0.697376895 |
| 692 | Macrophage-C | 21 | ENSMUSG00000069515 | Lyz1 | 0.456860648 |
| 693 | Macrophage-C | 21 | ENSMUSG00000069516 | Lyz2 | 0.519569054 |
| 694 | Macrophage-C | 21 | ENSMUSG00000069792 | Wfdc17 | 0.568068392 |
| 695 | Macrophage-C | 21 | ENSMUSG00000020377 | Ltc4s | 0.367968673 |
| 696 | Macrophage-C | 21 | ENSMUSG00000063193 | Cd300lb | 0.249642548 |
| 697 | Macrophage-C | 21 | ENSMUSG00000085761 | 4930455G09Rik | 0.267092038 |
| 698 | Macrophage-C | 21 | ENSMUSG00000085327 | Gm16104 | 0.254676284 |
| 699 | Macrophage-C | 21 | ENSMUSG00000092572 | Serpinb10 | 0.451251015 |
| 700 | Macrophage-C | 21 | ENSMUSG00000108371 | Gm38832 | 0.460398311 |
| 701 | Macrophage-B | 3 | ENSMUSG00000035352 | Ccl12 | 0.194408068 |
| 702 | Macrophage-B | 3 | ENSMUSG00000002985 | Apoe | 0.181662025 |
| 703 | Macrophage-B | 3 | ENSMUSG00000004730 | Adgre1 | 0.217694755 |
| 704 | Macrophage-B | 3 | ENSMUSG00000021939 | Ctsb | 0.201575434 |
| 705 | Macrophage-B | 3 | ENSMUSG00000038642 | Ctss | 0.237381174 |
| 706 | Macrophage-B | 3 | ENSMUSG00000017707 | Serinc3 | 0.161902435 |
| 707 | Macrophage-B | 3 | ENSMUSG00000018774 | Cd68 | 0.154352625 |
| 708 | Macrophage-B | 3 | ENSMUSG00000018920 | Cxcl16 | 0.170940051 |
| 709 | Macrophage-B | 3 | ENSMUSG00000021190 | Lgmn | 0.202902376 |
| 710 | Macrophage-B | 3 | ENSMUSG00000021423 | Ly86 | 0.181527318 |
| 711 | Macrophage-B | 3 | ENSMUSG00000024397 | Aif1 | 0.186369545 |
| 712 | Macrophage-B | 3 | ENSMUSG00000024621 | Csf1r | 0.271638136 |
| 713 | Macrophage-B | 3 | ENSMUSG00000024672 | Ms4a7 | 0.243343933 |
| 714 | Macrophage-B | 3 | ENSMUSG00000024677 | Ms4a6b | 0.167931527 |
| 715 | Macrophage-B | 3 | ENSMUSG00000026656 | Fcgr2b | 0.177378166 |
| 716 | Macrophage-B | 3 | ENSMUSG00000026712 | Mrc1 | 0.208656458 |
| 717 | Macrophage-B | 3 | ENSMUSG00000015947 | Fcgr1 | 0.172269399 |
| 718 | Macrophage-B | 3 | ENSMUSG00000029373 | Pf4 | 0.15722968 |
| 719 | Macrophage-B | 3 | ENSMUSG00000030148 | Clec4a2 | 0.178817176 |

| | | | | | |
|---|---|---|---|---|---|
| 720 | Macrophage-B | 3 | ENSMUSG00000030560 | Ctsc | 0.257898118 |
| 721 | Macrophage-B | 3 | ENSMUSG00000032359 | Ctsh | 0.205505075 |
| 722 | Macrophage-B | 3 | ENSMUSG00000032554 | Trf | 0.217323333 |
| 723 | Macrophage-B | 3 | ENSMUSG00000036594 | H2-Aa | 0.235319587 |
| 724 | Macrophage-B | 3 | ENSMUSG00000073421 | H2-Ab1 | 0.243762013 |
| 725 | Macrophage-B | 3 | ENSMUSG00000040950 | Mgl2 | 0.155922689 |
| 726 | Macrophage-B | 3 | ENSMUSG00000040552 | C3ar1 | 0.198882808 |
| 727 | Macrophage-B | 3 | ENSMUSG00000037649 | H2-DMa | 0.195430067 |
| 728 | Macrophage-B | 3 | ENSMUSG00000036887 | C1qa | 0.260865686 |
| 729 | Macrophage-B | 3 | ENSMUSG00000036896 | C1qc | 0.254231965 |
| 730 | Macrophage-B | 3 | ENSMUSG00000036905 | C1qb | 0.24675067 |
| 731 | Macrophage-B | 3 | ENSMUSG00000024610 | Cd74 | 0.231299376 |
| 732 | Macrophage-B | 3 | ENSMUSG00000049130 | C5ar1 | 0.257841714 |
| 733 | Macrophage-B | 3 | ENSMUSG00000049988 | Lrrc25 | 0.162246442 |
| 734 | Macrophage-B | 3 | ENSMUSG00000048779 | P2ry6 | 0.167586125 |
| 735 | Macrophage-B | 3 | ENSMUSG00000049037 | Clec4a1 | 0.191604392 |
| 736 | Macrophage-B | 3 | ENSMUSG00000051439 | Cd14 | 0.170367239 |
| 737 | Macrophage-B | 3 | ENSMUSG00000052160 | Pld4 | 0.19881575 |
| 738 | Macrophage-B | 3 | ENSMUSG00000055541 | Lair1 | 0.183999126 |
| 739 | Macrophage-B | 3 | ENSMUSG00000055435 | Maf | 0.191610136 |
| 740 | Macrophage-B | 3 | ENSMUSG00000060586 | H2-Eb1 | 0.236997372 |
| 741 | Macrophage-B | 3 | ENSMUSG00000022150 | Dab2 | 0.15659219 |
| 742 | Macrophage-B | 3 | ENSMUSG00000059089 | Fcgr4 | 0.154556019 |
| 743 | Macrophage-B | 3 | ENSMUSG00000058715 | Fcer1g | 0.192895349 |
| 744 | Macrophage-B | 3 | ENSMUSG00000015852 | Fcrls | 0.164574129 |
| 745 | Macrophage-B | 3 | ENSMUSG00000073412 | Lst1 | 0.19293467 |
| 746 | Macrophage-B | 3 | ENSMUSG00000074622 | Mafb | 0.198756601 |
| 747 | Macrophage-B | 3 | ENSMUSG00000079547 | H2-DMb1 | 0.218190731 |
| 748 | Macrophage-B | 3 | ENSMUSG00000070873 | Lilra5 | 0.162658575 |
| 749 | Macrophage-B | 3 | ENSMUSG00000036908 | Unc93b1 | 0.227601888 |
| 750 | Macrophage-B | 3 | ENSMUSG00000059498 | Fcgr3 | 0.238575398 |
| 751 | Neuroglia | 4 | ENSMUSG00000014846 | Tppp3 | 0.178878672 |
| 752 | Neuroglia | 4 | ENSMUSG00000018593 | Sparc | 0.20343776 |
| 753 | Neuroglia | 4 | ENSMUSG00000018822 | Sfrp5 | 0.160901706 |
| 754 | Neuroglia | 4 | ENSMUSG00000019874 | Fabp7 | 0.160586549 |
| 755 | Neuroglia | 4 | ENSMUSG00000020774 | Aspa | 0.190709723 |
| 756 | Neuroglia | 4 | ENSMUSG00000022048 | Dpysl2 | 0.11919528 |
| 757 | Neuroglia | 4 | ENSMUSG00000022103 | Gfra2 | 0.14119848 |
| 758 | Neuroglia | 4 | ENSMUSG00000022324 | Matn2 | 0.175082335 |
| 759 | Neuroglia | 4 | ENSMUSG00000022548 | Apod | 0.225511009 |
| 760 | Neuroglia | 4 | ENSMUSG00000024665 | Fads2 | 0.149355531 |
| 761 | Neuroglia | 4 | ENSMUSG00000025203 | Scd2 | 0.118242473 |
| 762 | Neuroglia | 4 | ENSMUSG00000025780 | Itih5 | 0.192077771 |
| 763 | Neuroglia | 4 | ENSMUSG00000026249 | Serpine2 | 0.179903975 |
| 764 | Neuroglia | 4 | ENSMUSG00000026385 | Dbi | 0.30309702 |

| | | | | | |
|---|---|---|---|---|---|
| 765 | Neuroglia | 4 | ENSMUSG00000026424 | Gpr37l1 | 0.281333282 |
| 766 | Neuroglia | 4 | ENSMUSG00000027712 | Anxa5 | 0.118274273 |
| 767 | Neuroglia | 4 | ENSMUSG00000028128 | F3 | 0.128141779 |
| 768 | Neuroglia | 4 | ENSMUSG00000030342 | Cd9 | 0.178142378 |
| 769 | Neuroglia | 4 | ENSMUSG00000031425 | Plp1 | 0.32816092 |
| 770 | Neuroglia | 4 | ENSMUSG00000031548 | Sfrp1 | 0.194323053 |
| 771 | Neuroglia | 4 | ENSMUSG00000032060 | Cryab | 0.261522821 |
| 772 | Neuroglia | 4 | ENSMUSG00000033208 | S100b | 0.248611105 |
| 773 | Neuroglia | 4 | ENSMUSG00000040759 | Cmtm5 | 0.1207736 |
| 774 | Neuroglia | 4 | ENSMUSG00000040724 | Kcna2 | 0.12751356 |
| 775 | Neuroglia | 4 | ENSMUSG00000036570 | Fxyd1 | 0.328367118 |
| 776 | Neuroglia | 4 | ENSMUSG00000033006 | Sox10 | 0.12723249 |
| 777 | Neuroglia | 4 | ENSMUSG00000032679 | Cd59a | 0.199048913 |
| 778 | Neuroglia | 4 | ENSMUSG00000036169 | Sostdc1 | 0.299544231 |
| 779 | Neuroglia | 4 | ENSMUSG00000042116 | Vwa1 | 0.167871027 |
| 780 | Neuroglia | 4 | ENSMUSG00000034810 | Scn7a | 0.370841864 |
| 781 | Neuroglia | 4 | ENSMUSG00000025856 | Pdgfa | 0.173636441 |
| 782 | Neuroglia | 4 | ENSMUSG00000069601 | Ank3 | 0.142232812 |
| 783 | Neuroglia | 4 | ENSMUSG00000041607 | Mbp | 0.173135143 |
| 784 | Neuroglia | 4 | ENSMUSG00000059857 | Ntng1 | 0.12796583 |
| 785 | Neuroglia | 4 | ENSMUSG00000038668 | Lpar1 | 0.168738884 |
| 786 | Neuroglia | 4 | ENSMUSG00000047976 | Kcna1 | 0.275271422 |
| 787 | Neuroglia | 4 | ENSMUSG00000031342 | Gpm6b | 0.165900242 |
| 788 | Neuroglia | 4 | ENSMUSG00000020926 | Adam11 | 0.151689168 |
| 789 | Neuroglia | 4 | ENSMUSG00000032268 | Tmprss5 | 0.194343827 |
| 790 | Neuroglia | 4 | ENSMUSG00000032332 | Col12a1 | 0.124797585 |
| 791 | Neuroglia | 4 | ENSMUSG00000027750 | Postn | 0.13568246 |
| 792 | Neuroglia | 4 | ENSMUSG00000057969 | Sema3b | 0.164929223 |
| 793 | Neuroglia | 4 | ENSMUSG00000030701 | Plekhb1 | 0.159883721 |
| 794 | Neuroglia | 4 | ENSMUSG00000025964 | Adam23 | 0.152554658 |
| 795 | Neuroglia | 4 | ENSMUSG00000053279 | Aldh1a1 | 0.125476038 |
| 796 | Neuroglia | 4 | ENSMUSG00000047216 | Cdh19 | 0.196661807 |
| 797 | Neuroglia | 4 | ENSMUSG00000029838 | Ptn | 0.121150285 |
| 798 | Neuroglia | 4 | ENSMUSG00000034842 | Art3 | 0.21131014 |
| 799 | Neuroglia | 4 | ENSMUSG00000039542 | Ncam1 | 0.281302333 |
| 800 | Neuroglia | 4 | ENSMUSG00000097789 | Gm2115 | 0.122246052 |
| 801 | Vascular endothelium | 5 | ENSMUSG00000001946 | Esam | 0.25855395 |
| 802 | Vascular endothelium | 5 | ENSMUSG00000004655 | Aqp1 | 0.487934165 |
| 803 | Vascular endothelium | 5 | ENSMUSG00000026814 | Eng | 0.204704492 |
| 804 | Vascular endothelium | 5 | ENSMUSG00000017309 | Cd300lg | 0.166309937 |
| 805 | Vascular endothelium | 5 | ENSMUSG00000020577 | Tspan13 | 0.235579382 |
| 806 | Vascular endothelium | 5 | ENSMUSG00000020846 | Rflnb | 0.217862331 |
| 807 | Vascular endothelium | 5 | ENSMUSG00000021367 | Edn1 | 0.210083866 |
| 808 | Vascular endothelium | 5 | ENSMUSG00000044258 | Ctla2a | 0.238431766 |
| 809 | Vascular endothelium | 5 | ENSMUSG00000022018 | Rgcc | 0.235162704 |

| | | | | | |
|---|---|---|---|---|---|
| 810 | Vascular endothelium | 5 | ENSMUSG00000075602 | Ly6a | 0.301746155 |
| 811 | Vascular endothelium | 5 | ENSMUSG00000022661 | Cd200 | 0.211104557 |
| 812 | Vascular endothelium | 5 | ENSMUSG00000024140 | Epas1 | 0.222952964 |
| 813 | Vascular endothelium | 5 | ENSMUSG00000024395 | Lims2 | 0.18712263 |
| 814 | Vascular endothelium | 5 | ENSMUSG00000025608 | Podxl | 0.176370783 |
| 815 | Vascular endothelium | 5 | ENSMUSG00000025902 | Sox17 | 0.196507347 |
| 816 | Vascular endothelium | 5 | ENSMUSG00000062515 | Fabp4 | 0.513452056 |
| 817 | Vascular endothelium | 5 | ENSMUSG00000027800 | Tm4sf1 | 0.17872556 |
| 818 | Vascular endothelium | 5 | ENSMUSG00000028427 | Aqp7 | 0.192231995 |
| 819 | Vascular endothelium | 5 | ENSMUSG00000029309 | Sparcl1 | 0.209174214 |
| 820 | Vascular endothelium | 5 | ENSMUSG00000029648 | Flt1 | 0.32078873 |
| 821 | Vascular endothelium | 5 | ENSMUSG00000032766 | Gng11 | 0.165870138 |
| 822 | Vascular endothelium | 5 | ENSMUSG00000031871 | Cdh5 | 0.370202419 |
| 823 | Vascular endothelium | 5 | ENSMUSG00000036144 | Meox2 | 0.16265381 |
| 824 | Vascular endothelium | 5 | ENSMUSG00000035686 | Thrsp | 0.207126775 |
| 825 | Vascular endothelium | 5 | ENSMUSG00000036256 | Igfbp7 | 0.174176038 |
| 826 | Vascular endothelium | 5 | ENSMUSG00000039167 | Adgrl4 | 0.255447227 |
| 827 | Vascular endothelium | 5 | ENSMUSG00000033191 | Tie1 | 0.162569331 |
| 828 | Vascular endothelium | 5 | ENSMUSG00000034911 | Ushbp1 | 0.212838538 |
| 829 | Vascular endothelium | 5 | ENSMUSG00000045377 | Tmem88 | 0.183503639 |
| 830 | Vascular endothelium | 5 | ENSMUSG00000046916 | Myct1 | 0.213850691 |
| 831 | Vascular endothelium | 5 | ENSMUSG00000045092 | S1pr1 | 0.28356497 |
| 832 | Vascular endothelium | 5 | ENSMUSG00000044562 | Rasip1 | 0.180316254 |
| 833 | Vascular endothelium | 5 | ENSMUSG00000045534 | Kcna5 | 0.215877368 |
| 834 | Vascular endothelium | 5 | ENSMUSG00000020717 | Pecam1 | 0.363026184 |
| 835 | Vascular endothelium | 5 | ENSMUSG00000061353 | Cxcl12 | 0.433800072 |
| 836 | Vascular endothelium | 5 | ENSMUSG00000058254 | Tspan7 | 0.220655186 |
| 837 | Vascular endothelium | 5 | ENSMUSG00000063415 | Cyp26b1 | 0.209469648 |
| 838 | Vascular endothelium | 5 | ENSMUSG00000002944 | Cd36 | 0.35442404 |
| 839 | Vascular endothelium | 5 | ENSMUSG00000037936 | Scarb1 | 0.226349875 |
| 840 | Vascular endothelium | 5 | ENSMUSG00000068079 | Tcf15 | 0.254579352 |
| 841 | Vascular endothelium | 5 | ENSMUSG00000020154 | Ptprb | 0.364337161 |
| 842 | Vascular endothelium | 5 | ENSMUSG00000027435 | Cd93 | 0.232696051 |
| 843 | Vascular endothelium | 5 | ENSMUSG00000026921 | Egfl7 | 0.273986089 |
| 844 | Vascular endothelium | 5 | ENSMUSG00000001930 | Vwf | 0.227933823 |
| 845 | Vascular endothelium | 5 | ENSMUSG00000032125 | Robo4 | 0.183719187 |
| 846 | Vascular endothelium | 5 | ENSMUSG00000041445 | Mmrn2 | 0.162288829 |
| 847 | Vascular endothelium | 5 | ENSMUSG00000062960 | Kdr | 0.153658278 |
| 848 | Vascular endothelium | 5 | ENSMUSG00000056492 | Adgrf5 | 0.30049769 |
| 849 | Vascular endothelium | 5 | ENSMUSG00000054690 | Emcn | 0.220504806 |
| 850 | Vascular endothelium | 5 | ENSMUSG00000068036 | Afdn | 0.187536898 |
| 851 | Smooth muscle cells | 6 | ENSMUSG00000001270 | Ckb | 0.044512355 |
| 852 | Smooth muscle cells | 6 | ENSMUSG00000001349 | Cnn1 | 0.042632764 |
| 853 | Smooth muscle cells | 6 | ENSMUSG00000005611 | Mrvi1 | 0.04686416 |
| 854 | Smooth muscle cells | 6 | ENSMUSG00000005672 | Kit | 0.096315078 |

| | | | | | |
|---|---|---|---|---|---|
| 855 | Smooth muscle cells | 6 | ENSMUSG00000020439 | Smtn | 0.142793709 |
| 856 | Smooth muscle cells | 6 | ENSMUSG00000020719 | Ddx5 | 0.076840728 |
| 857 | Smooth muscle cells | 6 | ENSMUSG00000024661 | Fth1 | 0.049047414 |
| 858 | Smooth muscle cells | 6 | ENSMUSG00000026208 | Des | 0.091448739 |
| 859 | Smooth muscle cells | 6 | ENSMUSG00000026421 | Csrp1 | 0.044586938 |
| 860 | Smooth muscle cells | 6 | ENSMUSG00000028461 | Ccdc107 | 0.065949776 |
| 861 | Smooth muscle cells | 6 | ENSMUSG00000028464 | Tpm2 | 0.125711026 |
| 862 | Smooth muscle cells | 6 | ENSMUSG00000032366 | Tpm1 | 0.086791569 |
| 863 | Smooth muscle cells | 6 | ENSMUSG00000029309 | Sparcl1 | 0.055872942 |
| 864 | Smooth muscle cells | 6 | ENSMUSG00000030409 | Dmpk | 0.108429654 |
| 865 | Smooth muscle cells | 6 | ENSMUSG00000031075 | Ano1 | 0.062515505 |
| 866 | Smooth muscle cells | 6 | ENSMUSG00000031328 | Flna | 0.133841765 |
| 867 | Smooth muscle cells | 6 | ENSMUSG00000031586 | Rbpms | 0.075690891 |
| 868 | Smooth muscle cells | 6 | ENSMUSG00000031842 | Pde4c | 0.060206809 |
| 869 | Smooth muscle cells | 6 | ENSMUSG00000032085 | Tagln | 0.068307546 |
| 870 | Smooth muscle cells | 6 | ENSMUSG00000035967 | Ints6l | 0.090769388 |
| 871 | Smooth muscle cells | 6 | ENSMUSG00000035783 | Acta2 | 0.13278599 |
| 872 | Smooth muscle cells | 6 | ENSMUSG00000037126 | Psd | 0.048024312 |
| 873 | Smooth muscle cells | 6 | ENSMUSG00000041741 | Pde3a | 0.062104617 |
| 874 | Smooth muscle cells | 6 | ENSMUSG00000026463 | Atp2b4 | 0.042966301 |
| 875 | Smooth muscle cells | 6 | ENSMUSG00000037852 | Cpe | 0.081338252 |
| 876 | Smooth muscle cells | 6 | ENSMUSG00000055065 | Ddx17 | 0.057001622 |
| 877 | Smooth muscle cells | 6 | ENSMUSG00000049422 | Chchd10 | 0.042292163 |
| 878 | Smooth muscle cells | 6 | ENSMUSG00000059430 | Actg2 | 0.094062339 |
| 879 | Smooth muscle cells | 6 | ENSMUSG00000064339 | mt-Rnr2 | 0.083584803 |
| 880 | Smooth muscle cells | 6 | ENSMUSG00000064341 | mt-Nd1 | 0.126628605 |
| 881 | Smooth muscle cells | 6 | ENSMUSG00000064345 | mt-Nd2 | 0.114182869 |
| 882 | Smooth muscle cells | 6 | ENSMUSG00000064351 | mt-Co1 | 0.043153331 |
| 883 | Smooth muscle cells | 6 | ENSMUSG00000064357 | mt-Atp6 | 0.084305575 |
| 884 | Smooth muscle cells | 6 | ENSMUSG00000064358 | mt-Co3 | 0.097962996 |
| 885 | Smooth muscle cells | 6 | ENSMUSG00000064363 | mt-Nd4 | 0.134049593 |
| 886 | Smooth muscle cells | 6 | ENSMUSG00000064368 | mt-Nd6 | 0.059089433 |
| 887 | Smooth muscle cells | 6 | ENSMUSG00000064370 | mt-Cytb | 0.133655535 |
| 888 | Smooth muscle cells | 6 | ENSMUSG00000067818 | Myl9 | 0.107251342 |
| 889 | Smooth muscle cells | 6 | ENSMUSG00000018830 | Myh11 | 0.11119696 |
| 890 | Smooth muscle cells | 6 | ENSMUSG00000021134 | Srsf5 | 0.048877983 |
| 891 | Smooth muscle cells | 6 | ENSMUSG00000004151 | Etv1 | 0.054750157 |
| 892 | Smooth muscle cells | 6 | ENSMUSG00000017002 | Slpi | 0.043421157 |
| 893 | Smooth muscle cells | 6 | ENSMUSG00000055632 | Hmcn2 | 0.105138484 |
| 894 | Smooth muscle cells | 6 | ENSMUSG00000034275 | Igsf9b | 0.144568349 |
| 895 | Smooth muscle cells | 6 | ENSMUSG00000024597 | Slc12a2 | 0.049225605 |
| 896 | Smooth muscle cells | 6 | ENSMUSG00000090841 | Myl6 | 0.10085427 |
| 897 | Smooth muscle cells | 6 | ENSMUSG00000092341 | Malat1 | 0.061462557 |
| 898 | Smooth muscle cells | 6 | ENSMUSG00000093674 | Rpl41 | 0.04857493 |
| 899 | Smooth muscle cells | 6 | ENSMUSG00000097971 | Gm26917 | 0.10633018 |

| | | | | | |
|---|---|---|---|---|---|
| 900 | Smooth muscle cells | 6 | ENSMUSG00000101939 | Gm28438 | 0.071258172 |
| 901 | RBC | 7 | ENSMUSG00000006360 | Crip1 | 0.011515177 |
| 902 | RBC | 7 | ENSMUSG00000006574 | Slc4a1 | 0.025688571 |
| 903 | RBC | 7 | ENSMUSG00000019505 | Ubb | 0.011693635 |
| 904 | RBC | 7 | ENSMUSG00000020295 | Hbq1a | 0.057584674 |
| 905 | RBC | 7 | ENSMUSG00000020641 | Rsad2 | 0.007990184 |
| 906 | RBC | 7 | ENSMUSG00000022051 | Bnip3l | 0.031045028 |
| 907 | RBC | 7 | ENSMUSG00000023046 | Igfbp6 | 0.010230757 |
| 908 | RBC | 7 | ENSMUSG00000052305 | Hbb-bs | 0.767469437 |
| 909 | RBC | 7 | ENSMUSG00000023995 | Tspo2 | 0.009374504 |
| 910 | RBC | 7 | ENSMUSG00000024588 | Fech | 0.037919231 |
| 911 | RBC | 7 | ENSMUSG00000024661 | Fth1 | 0.009569772 |
| 912 | RBC | 7 | ENSMUSG00000028906 | Epb41 | 0.009677488 |
| 913 | RBC | 7 | ENSMUSG00000029922 | Mkrn1 | 0.055569706 |
| 914 | RBC | 7 | ENSMUSG00000031950 | Gabarapl2 | 0.00994654 |
| 915 | RBC | 7 | ENSMUSG00000032411 | Tfdp2 | 0.009694653 |
| 916 | RBC | 7 | ENSMUSG00000034248 | Slc25a37 | 0.053734074 |
| 917 | RBC | 7 | ENSMUSG00000039236 | Isg20 | 0.06196803 |
| 918 | RBC | 7 | ENSMUSG00000038871 | Bpgm | 0.151831356 |
| 919 | RBC | 7 | ENSMUSG00000044792 | Isca1 | 0.028407585 |
| 920 | RBC | 7 | ENSMUSG00000047139 | Cd24a | 0.01006716 |
| 921 | RBC | 7 | ENSMUSG00000023572 | Ccndbp1 | 0.012994876 |
| 922 | RBC | 7 | ENSMUSG00000073400 | Trim10 | 0.011935785 |
| 923 | RBC | 7 | ENSMUSG00000035242 | Oaz1 | 0.009833361 |
| 924 | RBC | 7 | ENSMUSG00000051839 | Gypa | 0.070886283 |
| 925 | RBC | 7 | ENSMUSG00000025270 | Alas2 | 0.453144735 |
| 926 | RBC | 7 | ENSMUSG00000064337 | mt-Rnr1 | 0.007781301 |
| 927 | RBC | 7 | ENSMUSG00000064339 | mt-Rnr2 | 0.024780478 |
| 928 | RBC | 7 | ENSMUSG00000064341 | mt-Nd1 | 0.044390826 |
| 929 | RBC | 7 | ENSMUSG00000064345 | mt-Nd2 | 0.036847285 |
| 930 | RBC | 7 | ENSMUSG00000064351 | mt-Co1 | 0.01578484 |
| 931 | RBC | 7 | ENSMUSG00000064357 | mt-Atp6 | 0.024242122 |
| 932 | RBC | 7 | ENSMUSG00000064358 | mt-Co3 | 0.027560643 |
| 933 | RBC | 7 | ENSMUSG00000064363 | mt-Nd4 | 0.047737362 |
| 934 | RBC | 7 | ENSMUSG00000064368 | mt-Nd6 | 0.012628245 |
| 935 | RBC | 7 | ENSMUSG00000064370 | mt-Cytb | 0.047188211 |
| 936 | RBC | 7 | ENSMUSG00000069917 | Hba-a2 | 0.766173317 |
| 937 | RBC | 7 | ENSMUSG00000069919 | Hba-a1 | 0.772964186 |
| 938 | RBC | 7 | ENSMUSG00000050708 | Ftl1 | 0.017848279 |
| 939 | RBC | 7 | ENSMUSG00000073940 | Hbb-bt | 0.709549989 |
| 940 | RBC | 7 | ENSMUSG00000074269 | Rec114 | 0.06148195 |
| 941 | RBC | 7 | ENSMUSG00000029580 | Actb | 0.007638896 |
| 942 | RBC | 7 | ENSMUSG00000073063 | Hbq1b | 0.017500863 |
| 943 | RBC | 7 | ENSMUSG00000017002 | Slpi | 0.014713645 |
| 944 | RBC | 7 | ENSMUSG00000078974 | Sec61g | 0.028728797 |

| | | | | |
|---|---|---|---|---|
| 945 | RBC | 7 | ENSMUSG00000056234 | Ncoa4 | 0.008913816 |
| 946 | RBC | 7 | ENSMUSG00000025889 | Snca | 0.199603939 |
| 947 | RBC | 7 | ENSMUSG00000091694 | Apol11b | 0.026105912 |
| 948 | RBC | 7 | ENSMUSG00000093674 | Rpl41 | 0.014375569 |
| 949 | RBC | 7 | ENSMUSG00000095079 | Igha | 0.009815438 |
| 950 | RBC | 7 | ENSMUSG00000101939 | Gm28438 | 0.016546624 |
| 951 | Pdgfra+ Fibroblasts | 8 | ENSMUSG00000029231 | Pdgfra | 0.421625797 |
| 952 | Pdgfra+ Fibroblasts | 8 | ENSMUSG00000000794 | Kcnn3 | 0.234521908 |
| 953 | Pdgfra+ Fibroblasts | 8 | ENSMUSG00000001506 | Col1a1 | 0.157057604 |
| 954 | Pdgfra+ Fibroblasts | 8 | ENSMUSG00000002847 | Pla1a | 0.146958075 |
| 955 | Pdgfra+ Fibroblasts | 8 | ENSMUSG00000002997 | Prkar2b | 0.157736199 |
| 956 | Pdgfra+ Fibroblasts | 8 | ENSMUSG00000019966 | Kitl | 0.186500033 |
| 957 | Pdgfra+ Fibroblasts | 8 | ENSMUSG00000022860 | Chodl | 0.303736643 |
| 958 | Pdgfra+ Fibroblasts | 8 | ENSMUSG00000022890 | Atp5j | 0.20902947 |
| 959 | Pdgfra+ Fibroblasts | 8 | ENSMUSG00000023224 | Serping1 | 0.155878364 |
| 960 | Pdgfra+ Fibroblasts | 8 | ENSMUSG00000023886 | Smoc2 | 0.240293313 |
| 961 | Pdgfra+ Fibroblasts | 8 | ENSMUSG00000023913 | Pla2g7 | 0.154503513 |
| 962 | Pdgfra+ Fibroblasts | 8 | ENSMUSG00000025488 | Cox8b | 0.199222527 |
| 963 | Pdgfra+ Fibroblasts | 8 | ENSMUSG00000025491 | Ifitm1 | 0.157008198 |
| 964 | Pdgfra+ Fibroblasts | 8 | ENSMUSG00000026574 | Dpt | 0.2061035 |
| 965 | Pdgfra+ Fibroblasts | 8 | ENSMUSG00000027210 | Meis2 | 0.18596615 |
| 966 | Pdgfra+ Fibroblasts | 8 | ENSMUSG00000028005 | Gucy1b1 | 0.226268112 |
| 967 | Pdgfra+ Fibroblasts | 8 | ENSMUSG00000028364 | Tnc | 0.232239058 |
| 968 | Pdgfra+ Fibroblasts | 8 | ENSMUSG00000029163 | Emilin1 | 0.228113592 |
| 969 | Pdgfra+ Fibroblasts | 8 | ENSMUSG00000029467 | Atp2a2 | 0.157102041 |
| 970 | Pdgfra+ Fibroblasts | 8 | ENSMUSG00000029661 | Col1a2 | 0.161634708 |
| 971 | Pdgfra+ Fibroblasts | 8 | ENSMUSG00000030218 | Mgp | 0.330195957 |
| 972 | Pdgfra+ Fibroblasts | 8 | ENSMUSG00000056973 | Ces1d | 0.335992721 |
| 973 | Pdgfra+ Fibroblasts | 8 | ENSMUSG00000031740 | Mmp2 | 0.154647209 |
| 974 | Pdgfra+ Fibroblasts | 8 | ENSMUSG00000032024 | Clmp | 0.199171332 |
| 975 | Pdgfra+ Fibroblasts | 8 | ENSMUSG00000027859 | Ngf | 0.153951611 |
| 976 | Pdgfra+ Fibroblasts | 8 | ENSMUSG00000036446 | Lum | 0.434627489 |
| 977 | Pdgfra+ Fibroblasts | 8 | ENSMUSG00000039252 | Lgi2 | 0.270556078 |
| 978 | Pdgfra+ Fibroblasts | 8 | ENSMUSG00000042436 | Mfap4 | 0.367178192 |
| 979 | Pdgfra+ Fibroblasts | 8 | ENSMUSG00000033910 | Gucy1a1 | 0.195125829 |
| 980 | Pdgfra+ Fibroblasts | 8 | ENSMUSG00000036766 | Dner | 0.188677324 |
| 981 | Pdgfra+ Fibroblasts | 8 | ENSMUSG00000006369 | Fbln1 | 0.35053298 |
| 982 | Pdgfra+ Fibroblasts | 8 | ENSMUSG00000046491 | C1qtnf2 | 0.145172438 |
| 983 | Pdgfra+ Fibroblasts | 8 | ENSMUSG00000048376 | F2r | 0.157757167 |
| 984 | Pdgfra+ Fibroblasts | 8 | ENSMUSG00000041261 | Car8 | 0.156933005 |
| 985 | Pdgfra+ Fibroblasts | 8 | ENSMUSG00000064179 | Tnnt1 | 0.158294574 |
| 986 | Pdgfra+ Fibroblasts | 8 | ENSMUSG00000058620 | Adra2b | 0.183565926 |
| 987 | Pdgfra+ Fibroblasts | 8 | ENSMUSG00000031673 | Cdh11 | 0.168847712 |
| 988 | Pdgfra+ Fibroblasts | 8 | ENSMUSG00000057280 | Musk | 0.27898356 |
| 989 | Pdgfra+ Fibroblasts | 8 | ENSMUSG00000066705 | Fxyd6 | 0.199668831 |

| | | | | | |
|---|---|---|---|---|---|
| 990 | Pdgfra+ Fibroblasts | 8 | ENSMUSG00000070576 | Mn1 | 0.187930888 |
| 991 | Pdgfra+ Fibroblasts | 8 | ENSMUSG00000053062 | Jam2 | 0.241732855 |
| 992 | Pdgfra+ Fibroblasts | 8 | ENSMUSG00000074971 | Fibin | 0.462272865 |
| 993 | Pdgfra+ Fibroblasts | 8 | ENSMUSG00000029838 | Ptn | 0.299532525 |
| 994 | Pdgfra+ Fibroblasts | 8 | ENSMUSG00000019929 | Dcn | 0.181427493 |
| 995 | Pdgfra+ Fibroblasts | 8 | ENSMUSG00000055632 | Hmcn2 | 0.182716047 |
| 996 | Pdgfra+ Fibroblasts | 8 | ENSMUSG00000052516 | Robo2 | 0.14701159 |
| 997 | Pdgfra+ Fibroblasts | 8 | ENSMUSG00000048022 | Tmem229a | 0.151045304 |
| 998 | Pdgfra+ Fibroblasts | 8 | ENSMUSG00000045658 | Pid1 | 0.161482713 |
| 999 | Pdgfra+ Fibroblasts | 8 | ENSMUSG00000021943 | Gdf10 | 0.214852841 |
| 1000 | Pdgfra+ Fibroblasts | 8 | ENSMUSG00000023885 | Thbs2 | 0.449965378 |
| 1001 | B Lymphocytes | 9 | ENSMUSG00000000903 | Vpreb3 | 0.218782885 |
| 1002 | B Lymphocytes | 9 | ENSMUSG00000003379 | Cd79a | 0.673138729 |
| 1003 | B Lymphocytes | 9 | ENSMUSG00000003970 | Rpl8 | 0.129340216 |
| 1004 | B Lymphocytes | 9 | ENSMUSG00000012848 | Rps5 | 0.133624786 |
| 1005 | B Lymphocytes | 9 | ENSMUSG00000005540 | Fcer2a | 0.135134819 |
| 1006 | B Lymphocytes | 9 | ENSMUSG00000005583 | Mef2c | 0.137426096 |
| 1007 | B Lymphocytes | 9 | ENSMUSG00000008668 | Rps18 | 0.148030432 |
| 1008 | B Lymphocytes | 9 | ENSMUSG00000017404 | Rpl19 | 0.130265285 |
| 1009 | B Lymphocytes | 9 | ENSMUSG00000024334 | H2-Oa | 0.128764061 |
| 1010 | B Lymphocytes | 9 | ENSMUSG00000025362 | Rps26 | 0.127154298 |
| 1011 | B Lymphocytes | 9 | ENSMUSG00000027368 | Dusp2 | 0.339040433 |
| 1012 | B Lymphocytes | 9 | ENSMUSG00000028081 | Rps3a1 | 0.137308631 |
| 1013 | B Lymphocytes | 9 | ENSMUSG00000030707 | Coro1a | 0.151828568 |
| 1014 | B Lymphocytes | 9 | ENSMUSG00000030724 | Cd19 | 0.186531445 |
| 1015 | B Lymphocytes | 9 | ENSMUSG00000030744 | Rps3 | 0.134127741 |
| 1016 | B Lymphocytes | 9 | ENSMUSG00000030798 | Cd37 | 0.227398939 |
| 1017 | B Lymphocytes | 9 | ENSMUSG00000032399 | Rpl4 | 0.134862711 |
| 1018 | B Lymphocytes | 9 | ENSMUSG00000032518 | Rpsa | 0.126736003 |
| 1019 | B Lymphocytes | 9 | ENSMUSG00000008193 | Spib | 0.158789469 |
| 1020 | B Lymphocytes | 9 | ENSMUSG00000034892 | Rps29 | 0.132151127 |
| 1021 | B Lymphocytes | 9 | ENSMUSG00000036478 | Btg1 | 0.159004464 |
| 1022 | B Lymphocytes | 9 | ENSMUSG00000042474 | Fcmr | 0.402390769 |
| 1023 | B Lymphocytes | 9 | ENSMUSG00000034634 | Ly6d | 0.631436981 |
| 1024 | B Lymphocytes | 9 | ENSMUSG00000037548 | H2-DMb2 | 0.358234576 |
| 1025 | B Lymphocytes | 9 | ENSMUSG00000038274 | Fau | 0.152528463 |
| 1026 | B Lymphocytes | 9 | ENSMUSG00000040592 | Cd79b | 0.480994885 |
| 1027 | B Lymphocytes | 9 | ENSMUSG00000038421 | Fcrla | 0.193555676 |
| 1028 | B Lymphocytes | 9 | ENSMUSG00000024610 | Cd74 | 0.148190422 |
| 1029 | B Lymphocytes | 9 | ENSMUSG00000045128 | Rpl18a | 0.156315222 |
| 1030 | B Lymphocytes | 9 | ENSMUSG00000062006 | Rpl34 | 0.136866727 |
| 1031 | B Lymphocytes | 9 | ENSMUSG00000063457 | Rps15 | 0.126298109 |
| 1032 | B Lymphocytes | 9 | ENSMUSG00000052837 | Junb | 0.159731551 |
| 1033 | B Lymphocytes | 9 | ENSMUSG00000057098 | Ebf1 | 0.212683637 |
| 1034 | B Lymphocytes | 9 | ENSMUSG00000060036 | Rpl3 | 0.130752325 |

| | | | | | |
|---|---|---|---|---|---|
| 1035 | B Lymphocytes | 9 | ENSMUSG00000057841 | Rpl32 | 0.140723695 |
| 1036 | B Lymphocytes | 9 | ENSMUSG00000067274 | Rplp0 | 0.132988198 |
| 1037 | B Lymphocytes | 9 | ENSMUSG00000067288 | Rps28 | 0.14846752 |
| 1038 | B Lymphocytes | 9 | ENSMUSG00000068105 | Tnfrsf13c | 0.2693664 |
| 1039 | B Lymphocytes | 9 | ENSMUSG00000028234 | Rps20 | 0.135724679 |
| 1040 | B Lymphocytes | 9 | ENSMUSG00000037944 | Ccr7 | 0.173605768 |
| 1041 | B Lymphocytes | 9 | ENSMUSG00000076617 | Ighm | 0.356513702 |
| 1042 | B Lymphocytes | 9 | ENSMUSG00000076937 | Iglc2 | 0.460852786 |
| 1043 | B Lymphocytes | 9 | ENSMUSG00000008683 | Rps15a | 0.13678476 |
| 1044 | B Lymphocytes | 9 | ENSMUSG00000040952 | Rps19 | 0.160995277 |
| 1045 | B Lymphocytes | 9 | ENSMUSG00000021754 | Map3k1 | 0.129247702 |
| 1046 | B Lymphocytes | 9 | ENSMUSG00000025290 | Rps24 | 0.14307302 |
| 1047 | B Lymphocytes | 9 | ENSMUSG00000079641 | Rpl39 | 0.131732759 |
| 1048 | B Lymphocytes | 9 | ENSMUSG00000024673 | Ms4a1 | 0.127638535 |
| 1049 | B Lymphocytes | 9 | ENSMUSG00000090862 | Rps13 | 0.145913739 |
| 1050 | B Lymphocytes | 9 | ENSMUSG00000104213 | Ighd | 0.263925032 |

| mean_expression | fraction_expressing | specificity | pseudo_R2 | marker_test_p_value |
|---|---|---|---|---|
| 3.181609816 | 0.638790036 | 0.096048737 | 0.024652417 | 4.96E-40 |
| 2.038975798 | 0.480427046 | 0.127087097 | 0.002692142 | 1.30E-05 |
| 3.42059983 | 0.660587189 | 0.099249818 | 0.031813612 | 2.92E-51 |
| 4.806937795 | 0.771352313 | 0.111910639 | 0.049751321 | 1.39E-79 |
| 1.351879534 | 0.387010676 | 0.149520013 | 0.009963662 | 4.49E-17 |
| 4.990066223 | 0.737099644 | 0.113916354 | 0.005389042 | 6.72E-10 |
| 2.136804119 | 0.50088968 | 0.296571484 | 0.172997515 | 3.09E-283 |
| 4.580151243 | 0.743772242 | 0.086994198 | 0.014956997 | 7.11E-25 |
| 15.76517747 | 0.96975089 | 0.147542353 | 0.1781753 | 3.57E-292 |
| 8.043152531 | 0.862099644 | 0.140363936 | 0.103715955 | 1.07E-166 |
| 3.783295727 | 0.591637011 | 0.198049646 | 0.193077463 | 4.41273743451818e-318 |
| 3.135481314 | 0.639234875 | 0.194667432 | 0.221959377 | 0 |
| 3.271225812 | 0.640569395 | 0.096290316 | 0.015401382 | 1.44E-25 |
| 1.372294441 | 0.373220641 | 0.160445665 | 0.014335486 | 6.65E-24 |
| 7.618103377 | 0.862544484 | 0.125582165 | 0.164378344 | 1.97E-268 |
| 10.58717566 | 0.870106762 | 0.172866403 | 0.285862533 | 0 |
| 11.38454497 | 0.861209964 | 0.157181138 | 0.234348632 | 0 |
| 3.147221295 | 0.659697509 | 0.114996032 | 0.072032437 | 3.48E-115 |
| 6.72300203 | 0.808274021 | 0.193076682 | 0.258152734 | 0 |
| 4.076045158 | 0.682829181 | 0.174787844 | 0.171039814 | 7.25E-280 |
| 6.165325631 | 0.799377224 | 0.192576128 | 0.261329177 | 0 |
| 43.52878319 | 0.980871886 | 0.187658064 | 0.402879453 | 0 |
| 3.897342204 | 0.704181495 | 0.089916267 | 0.023184329 | 9.86E-38 |
| 2.365075497 | 0.503113879 | 0.151373879 | 0.004414077 | 2.33E-08 |
| 6.149446652 | 0.820284698 | 0.095293697 | 0.045601828 | 5.27E-73 |
| 2.625151442 | 0.590302491 | 0.09903018 | 0.012043544 | 2.52E-20 |
| 2.805655592 | 0.614323843 | 0.100860202 | 0.012036246 | 2.59E-20 |
| 11.7169106 | 0.904359431 | 0.189298451 | 0.335148708 | 0 |
| 2.848013267 | 0.616548043 | 0.094890932 | 0.008049223 | 4.45E-14 |
| 3.588491131 | 0.708185053 | 0.109835404 | 0.0461633 | 6.80E-74 |
| 4.223415936 | 0.711298932 | 0.212723276 | 0.250886124 | 0 |
| 2.191224511 | 0.550711744 | 0.10783745 | 0.012069441 | 2.30E-20 |
| 2.860904467 | 0.62588968 | 0.095170138 | 0.008894965 | 2.11E-15 |
| 3.947282982 | 0.693505338 | 0.092848402 | 0.027114426 | 6.90E-44 |
| 9.365162335 | 0.87544484 | 0.112702979 | 0.016366357 | 4.48E-27 |
| 20.12379207 | 0.989323843 | 0.110019261 | 0.126004693 | 1.38E-203 |
| 5.210061539 | 0.798487544 | 0.115637425 | 0.024079841 | 3.91E-39 |
| 3.058945434 | 0.613879004 | 0.112702554 | 0.02127537 | 9.55E-35 |
| 4.850345875 | 0.775355872 | 0.114338524 | 0.063591339 | 1.22E-101 |
| 12.14856253 | 0.957295374 | 0.116364061 | 0.100856218 | 5.30E-162 |
| 23.47165238 | 0.995551601 | 0.11040295 | 0.14647601 | 5.78E-238 |
| 2.714752142 | 0.619661922 | 0.093566119 | 0.01265983 | 2.75E-21 |
| 10.04000671 | 0.84430605 | 0.111565751 | 0.297183217 | 0 |
| 16.89305372 | 0.96841637 | 0.175600627 | 0.374005293 | 0 |

| | | | | |
|---|---|---|---|---|
| 1.181194898 | 0.351423488 | 0.288479791 | 0.083375289 | 1.67E-133 |
| 14.06037974 | 0.816725979 | 0.099593661 | 0.030024904 | 1.88E-48 |
| 37.16138509 | 0.993772242 | 0.161435911 | 0.520989108 | 0 |
| 3.01557762 | 0.629003559 | 0.092926068 | 0.017199932 | 2.23E-28 |
| 3.505573311 | 0.667704626 | 0.095395725 | 0.015736409 | 4.31E-26 |
| 5.110974159 | 0.791370107 | 0.084476871 | 0.003517923 | 6.17E-07 |
| 0.637152455 | 0.584158416 | 0.48252115 | 0.178098794 | 1.25E-44 |
| 1.39169468 | 0.772277228 | 0.321744724 | 0.191557339 | 6.72E-48 |
| 0.784429912 | 0.663366337 | 0.326970102 | 0.174084614 | 1.18E-43 |
| 2.09792247 | 0.821782178 | 0.375082662 | 0.273534203 | 6.81E-68 |
| 0.702436093 | 0.425742574 | 0.925369715 | 0.333308561 | 1.45E-82 |
| 1.595600235 | 0.752475248 | 0.274023196 | 0.13951753 | 2.85E-35 |
| 1.301084825 | 0.752475248 | 0.464686885 | 0.284597938 | 1.33E-70 |
| 0.657904986 | 0.603960396 | 0.895397082 | 0.46026098 | 5.36E-114 |
| 1.743334568 | 0.762376238 | 0.28204773 | 0.121631011 | 6.11E-31 |
| 0.419960801 | 0.445544554 | 0.677471174 | 0.190686341 | 1.09E-47 |
| 1.26093524 | 0.742574257 | 0.409998016 | 0.281946673 | 5.93E-70 |
| 0.297394075 | 0.376237624 | 0.629457084 | 0.136372364 | 1.65E-34 |
| 0.63664264 | 0.574257426 | 0.357291611 | 0.152385866 | 2.17E-38 |
| 1.315460694 | 0.772277228 | 0.289663348 | 0.112149947 | 1.20E-28 |
| 0.576575304 | 0.633663366 | 0.896277443 | 0.507905581 | 6.78E-126 |
| 0.838001037 | 0.613861386 | 0.51852312 | 0.161001594 | 1.77E-40 |
| 2.067432143 | 0.762376238 | 0.296073571 | 0.186037008 | 1.48E-46 |
| 0.523391518 | 0.445544554 | 0.517990912 | 0.176154616 | 3.71E-44 |
| 2.363813563 | 0.821782178 | 0.24490547 | 0.137949933 | 6.83E-35 |
| 0.355795733 | 0.425742574 | 0.722182313 | 0.299925462 | 2.32E-74 |
| 3.960654551 | 0.851485149 | 0.600495643 | 0.555530347 | 7.61E-138 |
| 0.66459585 | 0.653465347 | 0.407876238 | 0.191342556 | 7.58E-48 |
| 0.567747014 | 0.544554455 | 0.44692307 | 0.142578725 | 5.17E-36 |
| 40.74068455 | 1 | 0.662758607 | 0.854689678 | 2.94E-214 |
| 1.620250991 | 0.643564356 | 0.651491234 | 0.338890486 | 6.15E-84 |
| 0.329833332 | 0.485148515 | 0.414955845 | 0.170437201 | 9.07E-43 |
| 2.690456606 | 0.861386139 | 0.408752449 | 0.340976472 | 1.88E-84 |
| 1.90350951 | 0.762376238 | 0.715780808 | 0.468397699 | 5.03E-116 |
| 0.978130899 | 0.683168317 | 0.429548579 | 0.183414922 | 6.40E-46 |
| 1.375392503 | 0.742574257 | 0.299026075 | 0.145367247 | 1.09E-36 |
| 0.145923234 | 0.287128713 | 0.852441847 | 0.179583153 | 5.46E-45 |
| 0.379024695 | 0.425742574 | 0.468123856 | 0.154497335 | 6.68E-39 |
| 0.310771443 | 0.405940594 | 0.567468443 | 0.214250584 | 2.02E-53 |
| 0.706738713 | 0.584158416 | 0.41264738 | 0.17322953 | 1.90E-43 |
| 0.575512934 | 0.584158416 | 0.496983868 | 0.170699285 | 7.83E-43 |
| 0.917758488 | 0.653465347 | 0.355736045 | 0.161876944 | 1.08E-40 |
| 0.943878984 | 0.603960396 | 0.627732453 | 0.35688444 | 2.26E-88 |
| 1.161751821 | 0.584158416 | 0.69102793 | 0.27608739 | 1.61E-68 |
| 1.922034935 | 0.792079208 | 0.253743671 | 0.301364286 | 1.03E-74 |

| | | | | |
|---|---|---|---|---|
| 1.828501102 | 0.772277228 | 0.447424916 | 0.250423512 | 3.05E-62 |
| 1.106433799 | 0.683168317 | 0.645018997 | 0.405742895 | 1.88E-100 |
| 5.458944697 | 0.95049505 | 0.398409221 | 0.410038047 | 1.62E-101 |
| 0.793611671 | 0.633663366 | 0.378286211 | 0.133590072 | 7.77E-34 |
| 0.234769124 | 0.356435644 | 0.577469119 | 0.09485258 | 1.84E-24 |
| 1.891145564 | 0.782178218 | 0.247923734 | 0.136357595 | 1.66E-34 |
| 0.68044076 | 0.504950495 | 0.542864099 | 0.201443561 | 2.65E-50 |
| 16.37256898 | 0.96039604 | 0.333112911 | 0.215637754 | 9.29E-54 |
| 0.96884293 | 0.435643564 | 0.686180738 | 0.29754713 | 8.89E-74 |
| 1.62361147 | 0.821782178 | 0.36701037 | 0.266397155 | 3.80E-66 |
| 1.042333137 | 0.772277228 | 0.325700949 | 0.17235845 | 3.10E-43 |
| 3.539364275 | 0.8875 | 0.668715874 | 0.720468647 | 6.50E-149 |
| 0.715196758 | 0.6 | 0.846894824 | 0.468884615 | 1.39E-96 |
| 7.499119616 | 0.75 | 0.776120193 | 0.60794592 | 2.25E-125 |
| 0.417333633 | 0.5 | 0.410072119 | 0.151761977 | 4.61E-32 |
| 33.41909158 | 0.85 | 0.790364297 | 0.70810523 | 2.57E-146 |
| 0.884192255 | 0.6875 | 0.54458517 | 0.508338622 | 1.02E-104 |
| 0.315507972 | 0.4 | 0.711902469 | 0.321217284 | 2.34E-66 |
| 1.577624062 | 0.7375 | 0.201011043 | 0.163674342 | 1.83E-34 |
| 0.28792459 | 0.3625 | 0.442248296 | 0.244076897 | 1.06E-50 |
| 1.177911932 | 0.5875 | 0.658806128 | 0.381022998 | 1.48E-78 |
| 0.136468369 | 0.1875 | 0.860563908 | 0.1029658 | 3.07E-22 |
| 0.388728519 | 0.45 | 0.442867594 | 0.257833301 | 1.74E-53 |
| 0.265003602 | 0.2875 | 0.498196064 | 0.188147769 | 2.13E-39 |
| 0.2971785 | 0.3625 | 0.39947513 | 0.190029602 | 8.89E-40 |
| 0.488224431 | 0.4875 | 0.471858511 | 0.174943543 | 9.81E-37 |
| 1.669801049 | 0.6375 | 0.504533172 | 0.444823862 | 1.24E-91 |
| 1.356300062 | 0.7625 | 0.555007534 | 0.577385399 | 5.11E-119 |
| 0.571937987 | 0.4875 | 0.386025452 | 0.235184442 | 6.70E-49 |
| 48.95590245 | 0.9625 | 0.682922062 | 0.78592941 | 1.03E-162 |
| 3.572536573 | 0.8875 | 0.206235549 | 0.284166126 | 7.94E-59 |
| 1.222391384 | 0.6625 | 0.373324312 | 0.256636629 | 3.04E-53 |
| 0.297909436 | 0.325 | 0.467408618 | 0.104314359 | 1.64E-22 |
| 0.317425372 | 0.3125 | 0.772966572 | 0.218518131 | 1.57E-45 |
| 0.851524197 | 0.6 | 0.514367662 | 0.296435662 | 2.56E-61 |
| 0.338359002 | 0.4 | 0.475173919 | 0.238173914 | 1.66E-49 |
| 5.146903693 | 0.925 | 0.166202786 | 0.253682147 | 1.21E-52 |
| 1.960209999 | 0.775 | 0.892288394 | 0.668763381 | 4.54E-138 |
| 0.663362474 | 0.6375 | 0.524278191 | 0.413262552 | 3.71E-85 |
| 0.676215254 | 0.575 | 0.278851689 | 0.216131455 | 4.77E-45 |
| 3.790232357 | 0.8 | 0.510517924 | 0.592799104 | 3.20E-122 |
| 0.713125461 | 0.55 | 0.278168577 | 0.182394459 | 3.09E-38 |
| 0.296483748 | 0.375 | 0.485974387 | 0.254059023 | 1.01E-52 |
| 0.605219506 | 0.45 | 0.456937618 | 0.297064384 | 1.91E-61 |
| 0.417102547 | 0.4 | 0.671835748 | 0.241134122 | 4.19E-50 |

| | | | | |
|---|---|---|---|---|
| 4.474610619 | 0.85 | 0.304077421 | 0.249897172 | 7.04E-52 |
| 0.154575541 | 0.1875 | 0.862985391 | 0.132408303 | 3.64E-28 |
| 1.220219917 | 0.7125 | 0.675365464 | 0.575596537 | 1.20E-118 |
| 0.945017878 | 0.3375 | 0.473307497 | 0.219458285 | 1.01E-45 |
| 1.835970983 | 0.7125 | 0.424744921 | 0.483141057 | 1.61E-99 |
| 1.205351334 | 0.5125 | 0.587332478 | 0.35884882 | 5.00E-74 |
| 0.514211202 | 0.4125 | 0.749782851 | 0.284296981 | 7.47E-59 |
| 1.005364095 | 0.65 | 0.472204469 | 0.442412046 | 3.87E-91 |
| 0.763782332 | 0.5875 | 0.245684938 | 0.149570213 | 1.27E-31 |
| 0.259779765 | 0.325 | 0.579725582 | 0.157351572 | 3.45E-33 |
| 1.591858919 | 0.6875 | 0.536787382 | 0.342818029 | 9.31E-71 |
| 0.220403168 | 0.2625 | 0.559717496 | 0.178551585 | 1.84E-37 |
| 0.499802433 | 0.5 | 0.463284084 | 0.333649637 | 6.87E-69 |
| 0.1840478 | 0.2 | 0.851145987 | 0.13712787 | 4.08E-29 |
| 0.475245827 | 0.5125 | 0.560170395 | 0.363349904 | 6.02E-75 |
| 0.469944922 | 0.3125 | 0.973781508 | 0.27754853 | 1.75E-57 |
| 0.477077741 | 0.413793103 | 0.777087631 | 0.195206105 | 7.51E-32 |
| 1.449534426 | 0.465517241 | 0.695007137 | 0.223722388 | 2.75E-36 |
| 0.850642745 | 0.25862069 | 0.769612891 | 0.130726572 | 7.88E-22 |
| 1.967753188 | 0.689655172 | 0.715414235 | 0.40270093 | 2.93E-64 |
| 0.616746013 | 0.431034483 | 0.644121998 | 0.196858165 | 4.15E-32 |
| 0.151092969 | 0.224137931 | 0.90142157 | 0.102088278 | 2.23E-17 |
| 0.606909319 | 0.413793103 | 0.789169142 | 0.213848412 | 9.45E-35 |
| 0.95994963 | 0.482758621 | 0.719932097 | 0.27607278 | 1.92E-44 |
| 1.458647973 | 0.5 | 0.613106316 | 0.224111718 | 2.39E-36 |
| 0.82091344 | 0.327586207 | 0.776605077 | 0.175960872 | 7.37E-29 |
| 0.735596832 | 0.413793103 | 0.769038182 | 0.192151294 | 2.24E-31 |
| 0.319519963 | 0.275862069 | 0.786541802 | 0.11084824 | 9.70E-19 |
| 1.073814157 | 0.482758621 | 0.53678251 | 0.199588638 | 1.56E-32 |
| 0.146872925 | 0.224137931 | 0.88821551 | 0.12582805 | 4.55E-21 |
| 0.239488208 | 0.275862069 | 0.735552921 | 0.116428788 | 1.32E-19 |
| 0.24313487 | 0.310344828 | 0.782401857 | 0.130669239 | 8.05E-22 |
| 0.442414756 | 0.293103448 | 0.720688325 | 0.101879499 | 2.41E-17 |
| 0.169675282 | 0.25862069 | 0.786410706 | 0.12280966 | 1.34E-20 |
| 0.763720677 | 0.327586207 | 0.696838862 | 0.154584834 | 1.55E-25 |
| 3.470624467 | 0.706896552 | 0.624779023 | 0.345106684 | 3.13E-55 |
| 0.289822417 | 0.275862069 | 0.832317359 | 0.120233397 | 3.37E-20 |
| 0.344769932 | 0.327586207 | 0.89903825 | 0.206785774 | 1.19E-33 |
| 0.816158351 | 0.465517241 | 0.590137995 | 0.204590674 | 2.61E-33 |
| 0.560883589 | 0.344827586 | 0.61389823 | 0.153001833 | 2.73E-25 |
| 0.740628563 | 0.448275862 | 0.556736212 | 0.156193643 | 8.70E-26 |
| 0.676121157 | 0.344827586 | 0.907484068 | 0.223817126 | 2.66E-36 |
| 0.376854608 | 0.293103448 | 0.778237613 | 0.136580784 | 9.70E-23 |
| 0.198325668 | 0.275862069 | 0.735634709 | 0.127451081 | 2.55E-21 |
| 0.238367019 | 0.275862069 | 0.829427297 | 0.118025102 | 7.43E-20 |

| | | | | |
|---|---|---|---|---|
| 0.366451224 | 0.275862069 | 0.740433159 | 0.121584227 | 2.08E-20 |
| 1.458595644 | 0.586206897 | 0.407824464 | 0.152198603 | 3.63E-25 |
| 0.357731716 | 0.275862069 | 0.794001138 | 0.1321336 | 4.76E-22 |
| 0.644266888 | 0.25862069 | 0.858521175 | 0.122161635 | 1.69E-20 |
| 0.419249803 | 0.327586207 | 0.880788197 | 0.165759023 | 2.84E-27 |
| 0.269871926 | 0.275862069 | 0.714839723 | 0.112436008 | 5.49E-19 |
| 0.796852542 | 0.293103448 | 0.829142889 | 0.198751765 | 2.11E-32 |
| 0.462897687 | 0.362068966 | 0.736324434 | 0.197062183 | 3.86E-32 |
| 0.204925801 | 0.24137931 | 0.855001372 | 0.107927249 | 2.76E-18 |
| 0.194235645 | 0.275862069 | 0.827609062 | 0.135173985 | 1.61E-22 |
| 0.52921691 | 0.362068966 | 0.579702057 | 0.116721974 | 1.18E-19 |
| 0.889082143 | 0.534482759 | 0.806741549 | 0.313087655 | 3.19E-50 |
| 1.870979518 | 0.482758621 | 0.565114711 | 0.168324599 | 1.13E-27 |
| 0.860195459 | 0.413793103 | 0.651470791 | 0.21087142 | 2.75E-34 |
| 0.428842312 | 0.344827586 | 0.658357312 | 0.133178141 | 3.28E-22 |
| 1.04023738 | 0.517241379 | 0.587279694 | 0.243203115 | 2.55E-39 |
| 0.652921487 | 0.448275862 | 0.489291129 | 0.138426351 | 5.01E-23 |
| 5.134037472 | 0.75862069 | 0.854218035 | 0.546572075 | 6.05E-87 |
| 0.419128245 | 0.275862069 | 0.904282989 | 0.154108904 | 1.83E-25 |
| 0.368531921 | 0.310344828 | 0.709120638 | 0.128842385 | 1.55E-21 |
| 1.855264696 | 0.655172414 | 0.577368274 | 0.215579399 | 5.08E-35 |
| 0.611019604 | 0.564102564 | 0.252561357 | 0.249816045 | 1.55E-29 |
| 0.931778131 | 0.666666667 | 0.574020422 | 0.43410804 | 2.91E-50 |
| 6.184612438 | 0.897435897 | 0.13588562 | 0.060070874 | 3.35E-08 |
| 6.639658803 | 0.897435897 | 0.145453904 | 0.117975915 | 9.66E-15 |
| 1.369834634 | 0.58974359 | 0.246954045 | 0.127616456 | 7.97E-16 |
| 4.629667084 | 0.948717949 | 0.149432546 | 0.116097067 | 1.57E-14 |
| 3.998378163 | 0.871794872 | 0.153943059 | 0.118881172 | 7.64E-15 |
| 0.551378841 | 0.512820513 | 0.413425864 | 0.173072095 | 6.29E-21 |
| 0.510212585 | 0.487179487 | 0.556608046 | 0.380145909 | 3.51E-44 |
| 0.235852278 | 0.307692308 | 0.525573154 | 0.193715266 | 3.04E-23 |
| 1.895649487 | 0.794871795 | 0.174151492 | 0.108997858 | 9.89E-14 |
| 0.773277766 | 0.615384615 | 0.197020978 | 0.069538332 | 2.82E-09 |
| 2.512271131 | 0.923076923 | 0.289603017 | 0.347460895 | 1.67E-40 |
| 3.026880143 | 0.769230769 | 0.179512768 | 0.095942158 | 2.92E-12 |
| 0.435219051 | 0.41025641 | 0.45083342 | 0.211495851 | 3.08E-25 |
| 0.58512937 | 0.538461538 | 0.342683254 | 0.203673462 | 2.33E-24 |
| 0.168863245 | 0.230769231 | 0.679754602 | 0.104347214 | 3.30E-13 |
| 1.509958026 | 0.743589744 | 0.426397905 | 0.391744903 | 1.73E-45 |
| 1.173004021 | 0.820512821 | 0.160516246 | 0.132057098 | 2.53E-16 |
| 1.237103541 | 0.717948718 | 0.303851164 | 0.205825164 | 1.33E-24 |
| 0.19109144 | 0.307692308 | 0.591782773 | 0.199015907 | 7.74E-24 |
| 0.373438096 | 0.435897436 | 0.819904226 | 0.373035688 | 2.22E-43 |
| 1.344917519 | 0.743589744 | 0.172831961 | 0.133127791 | 1.92E-16 |
| 0.601307622 | 0.358974359 | 0.387961568 | 0.124030303 | 2.02E-15 |

| | | | | |
|---|---|---|---|---|
| 0.840923855 | 0.692307692 | 0.193944099 | 0.14221464 | 1.83E-17 |
| 0.502308038 | 0.512820513 | 0.29071364 | 0.145551346 | 7.71E-18 |
| 0.501040782 | 0.41025641 | 0.293022497 | 0.076257467 | 4.89E-10 |
| 0.296876231 | 0.384615385 | 0.310279136 | 0.095475107 | 3.30E-12 |
| 3.280748121 | 0.923076923 | 0.193354884 | 0.250081638 | 1.45E-29 |
| 0.766525973 | 0.615384615 | 0.353327079 | 0.330878596 | 1.22E-38 |
| 6.207681097 | 0.948717949 | 0.140377529 | 0.141177601 | 2.39E-17 |
| 0.727064748 | 0.58974359 | 0.814124769 | 0.36762948 | 9.00E-43 |
| 0.072631004 | 0.128205128 | 0.963134605 | 0.107115077 | 1.61E-13 |
| 0.469767672 | 0.461538462 | 0.499304604 | 0.255963207 | 3.17E-30 |
| 0.636558661 | 0.538461538 | 0.275899369 | 0.130897273 | 3.41E-16 |
| 0.16191324 | 0.153846154 | 0.800102987 | 0.090568938 | 1.18E-11 |
| 8.443888702 | 0.846153846 | 0.24179422 | 0.272106107 | 4.89E-32 |
| 0.249720714 | 0.358974359 | 0.670358958 | 0.178882739 | 1.40E-21 |
| 0.47924263 | 0.41025641 | 0.356237475 | 0.080653074 | 1.56E-10 |
| 0.35766824 | 0.384615385 | 0.372656214 | 0.215283803 | 1.16E-25 |
| 0.298796432 | 0.435897436 | 0.53745651 | 0.266524872 | 2.07E-31 |
| 0.269315018 | 0.307692308 | 0.480962654 | 0.120425666 | 5.13E-15 |
| 1.760159407 | 0.769230769 | 0.201140098 | 0.13382262 | 1.60E-16 |
| 1.476848685 | 0.666666667 | 0.694272917 | 0.524497404 | 1.79E-60 |
| 0.699775309 | 0.538461538 | 0.580742669 | 0.320312539 | 1.88E-37 |
| 0.451695778 | 0.487179487 | 0.369813945 | 0.193601843 | 3.13E-23 |
| 0.219168816 | 0.256410256 | 0.926071003 | 0.223165093 | 1.51E-26 |
| 0.962927594 | 0.615384615 | 0.288662 | 0.1380146 | 5.41E-17 |
| 0.161992686 | 0.128205128 | 1 | 0.112553493 | 3.94E-14 |
| 4.768782978 | 0.948717949 | 0.142588647 | 0.131120135 | 3.22E-16 |
| 7.562876599 | 0.973684211 | 0.307161438 | 0.372095785 | 2.21E-42 |
| 0.185882707 | 0.447368421 | 0.605033048 | 0.133829825 | 3.34E-16 |
| 0.225354971 | 0.605263158 | 0.578567379 | 0.179725308 | 3.04E-21 |
| 0.329152499 | 0.421052632 | 0.624402433 | 0.158365111 | 6.73E-19 |
| 0.274469356 | 0.578947368 | 0.603763226 | 0.218165198 | 1.83E-25 |
| 0.178952503 | 0.421052632 | 0.798064407 | 0.25240415 | 3.19E-29 |
| 0.379796852 | 0.605263158 | 0.603085075 | 0.23832788 | 1.12E-27 |
| 0.161376778 | 0.394736842 | 0.678204889 | 0.167194503 | 7.22E-20 |
| 0.19809464 | 0.5 | 0.60341805 | 0.196037312 | 4.92E-23 |
| 0.14996815 | 0.342105263 | 0.814319782 | 0.229688446 | 9.96E-27 |
| 0.193748175 | 0.5 | 0.740258227 | 0.270105258 | 3.64E-31 |
| 0.219633919 | 0.526315789 | 0.414577028 | 0.165789888 | 1.03E-19 |
| 0.273330589 | 0.5 | 0.746834887 | 0.201974702 | 1.10E-23 |
| 0.421660612 | 0.684210526 | 0.348457207 | 0.202054757 | 1.08E-23 |
| 0.328400612 | 0.605263158 | 0.551983149 | 0.180706601 | 2.37E-21 |
| 0.142329408 | 0.421052632 | 0.546572671 | 0.126105265 | 2.36E-15 |
| 0.228647645 | 0.394736842 | 0.660039385 | 0.145948992 | 1.56E-17 |
| 0.308784768 | 0.526315789 | 0.60780732 | 0.211344257 | 1.03E-24 |
| 0.126076955 | 0.342105263 | 0.646104748 | 0.13853651 | 1.02E-16 |

| | | | | |
|---|---|---|---|---|
| 0.074855521 | 0.315789474 | 0.814467048 | 0.106735476 | 3.21E-13 |
| 0.333026644 | 0.578947368 | 0.382986992 | 0.153703112 | 2.19E-18 |
| 2.417834316 | 0.921052632 | 0.391894241 | 0.306770411 | 3.40E-35 |
| 0.333960361 | 0.605263158 | 0.647284619 | 0.237366134 | 1.43E-27 |
| 0.129668281 | 0.394736842 | 0.615373406 | 0.125533484 | 2.73E-15 |
| 0.292496449 | 0.578947368 | 0.511194008 | 0.144613973 | 2.18E-17 |
| 0.199386437 | 0.526315789 | 0.600187243 | 0.241145971 | 5.50E-28 |
| 0.374776741 | 0.578947368 | 0.759891123 | 0.293347263 | 1.02E-33 |
| 0.11845175 | 0.342105263 | 0.695176811 | 0.100426337 | 1.59E-12 |
| 0.539375056 | 0.763157895 | 0.334547272 | 0.203233199 | 7.98E-24 |
| 0.11287721 | 0.315789474 | 0.752361544 | 0.141520497 | 4.77E-17 |
| 0.118228842 | 0.394736842 | 0.774330031 | 0.210832507 | 1.17E-24 |
| 0.129790927 | 0.394736842 | 0.571935924 | 0.166756751 | 8.06E-20 |
| 0.873695395 | 0.736842105 | 0.507923052 | 0.280791504 | 2.44E-32 |
| 0.291337145 | 0.605263158 | 0.371283134 | 0.134869848 | 2.57E-16 |
| 0.205098064 | 0.368421053 | 0.599356425 | 0.160311968 | 4.11E-19 |
| 0.153013006 | 0.342105263 | 0.734417878 | 0.151070048 | 4.26E-18 |
| 2.495491964 | 0.947368421 | 0.385670737 | 0.402663402 | 9.49E-46 |
| 0.649924712 | 0.710526316 | 0.649566372 | 0.379084496 | 3.76E-43 |
| 3.250533707 | 0.868421053 | 0.302402241 | 0.250267883 | 5.48E-29 |
| 0.557640815 | 0.473684211 | 0.734349726 | 0.232361343 | 5.07E-27 |
| 0.719001855 | 0.736842105 | 0.70187304 | 0.40502687 | 5.21E-46 |
| 0.221221843 | 0.578947368 | 0.579884536 | 0.222708423 | 5.81E-26 |
| 0.140167898 | 0.315789474 | 0.90563683 | 0.153487183 | 2.31E-18 |
| 0.296282919 | 0.552631579 | 0.40640783 | 0.126031115 | 2.41E-15 |
| 0.18673075 | 0.5 | 0.585723581 | 0.170432252 | 3.18E-20 |
| 0.236976665 | 0.5 | 0.489911762 | 0.134564199 | 2.78E-16 |
| 0.337395998 | 0.605263158 | 0.380773618 | 0.149504218 | 6.33E-18 |
| 0.130321146 | 0.342105263 | 0.72088343 | 0.132792055 | 4.35E-16 |
| 0.203700632 | 0.421052632 | 0.684381512 | 0.185653924 | 6.79E-22 |
| 0.340816993 | 0.684210526 | 0.455889037 | 0.209659207 | 1.57E-24 |
| 0.380538294 | 0.236842105 | 0.943844902 | 0.2061085 | 3.86E-24 |
| 6.740366213 | 0.973684211 | 0.147137836 | 0.178548916 | 4.09E-21 |
| 0.834412415 | 0.763157895 | 0.173060206 | 0.130031053 | 8.75E-16 |
| 0.234339789 | 0.131578947 | 1 | 0.115565173 | 3.42E-14 |
| 2.812354902 | 0.947368421 | 0.139747926 | 0.103298483 | 7.68E-13 |
| 1.206002807 | 0.684210526 | 0.233644161 | 0.095955663 | 4.96E-12 |
| 2.066618904 | 0.736842105 | 0.249832329 | 0.22006981 | 1.13E-25 |
| 1.207759979 | 0.684210526 | 0.220673624 | 0.153951139 | 2.06E-18 |
| 0.719126157 | 0.605263158 | 0.241455626 | 0.243419419 | 3.10E-28 |
| 0.463396798 | 0.473684211 | 0.324691874 | 0.208600638 | 2.06E-24 |
| 1.354505509 | 0.789473684 | 0.179073872 | 0.150916931 | 4.43E-18 |
| 0.433583375 | 0.368421053 | 0.817656178 | 0.25481235 | 1.74E-29 |
| 0.315620285 | 0.368421053 | 0.467998049 | 0.182576592 | 1.48E-21 |
| 0.368047086 | 0.289473684 | 0.626723419 | 0.151787649 | 3.55E-18 |

| | | | | |
|---|---|---|---|---|
| 0.361395787 | 0.184210526 | 0.950779806 | 0.14096314 | 5.49E-17 |
| 1.274139257 | 0.763157895 | 0.192358146 | 0.128388559 | 1.33E-15 |
| 1.603560533 | 0.868421053 | 0.197496553 | 0.207765642 | 2.54E-24 |
| 4.358513794 | 0.947368421 | 0.149530686 | 0.136545674 | 1.68E-16 |
| 0.140476454 | 0.157894737 | 1 | 0.138956146 | 9.13E-17 |
| 1.008468216 | 0.605263158 | 0.222514212 | 0.147659974 | 1.01E-17 |
| 0.684481391 | 0.552631579 | 0.320687217 | 0.254851715 | 1.72E-29 |
| 2.302056982 | 0.842105263 | 0.746273804 | 0.580708863 | 1.94E-65 |
| 2.299321792 | 0.947368421 | 0.147547625 | 0.204843979 | 5.31E-24 |
| 0.700337722 | 0.631578947 | 0.329686008 | 0.326666737 | 2.21E-37 |
| 1.488621402 | 0.736842105 | 0.221998154 | 0.119361431 | 1.31E-14 |
| 1.26275962 | 0.710526316 | 0.198885019 | 0.146498315 | 1.35E-17 |
| 0.239487146 | 0.263157895 | 0.510232758 | 0.100354882 | 1.62E-12 |
| 0.398973569 | 0.315789474 | 0.595638347 | 0.188865873 | 3.01E-22 |
| 0.31766326 | 0.289473684 | 0.703942917 | 0.135900525 | 1.98E-16 |
| 5.444478976 | 0.921052632 | 0.173460054 | 0.234252116 | 3.14E-27 |
| 0.887221625 | 0.526315789 | 0.342887099 | 0.190292135 | 2.10E-22 |
| 0.278830315 | 0.236842105 | 0.703969928 | 0.157322699 | 8.76E-19 |
| 2.426436467 | 0.631578947 | 0.357374211 | 0.313604112 | 6.04E-36 |
| 0.169779842 | 0.210526316 | 0.96703289 | 0.177206725 | 5.74E-21 |
| 1.128901086 | 0.736842105 | 0.249576985 | 0.160036199 | 4.41E-19 |
| 0.938902797 | 0.657894737 | 0.342947964 | 0.236719044 | 1.68E-27 |
| 6.340716338 | 1 | 0.133985422 | 0.151912809 | 3.44E-18 |
| 4.547404594 | 1 | 0.139410216 | 0.142011931 | 4.21E-17 |
| 5.313034799 | 1 | 0.142744081 | 0.157728277 | 7.91E-19 |
| 0.441260769 | 0.526315789 | 0.356300405 | 0.255829968 | 1.34E-29 |
| 3.703679222 | 0.815789474 | 0.263491161 | 0.154070719 | 1.99E-18 |
| 0.766947532 | 0.605263158 | 0.26624175 | 0.189690089 | 2.45E-22 |
| 0.632744356 | 0.421052632 | 0.344963493 | 0.1986244 | 2.56E-23 |
| 1.384853939 | 0.763157895 | 0.341016897 | 0.364160799 | 1.65E-41 |
| 0.785540969 | 0.394736842 | 0.387862198 | 0.186477395 | 5.51E-22 |
| 0.247808054 | 0.263157895 | 0.499040764 | 0.112958059 | 6.62E-14 |
| 0.851872209 | 0.631578947 | 0.325540121 | 0.255102265 | 1.62E-29 |
| 7.486845678 | 1 | 0.177305793 | 0.22512794 | 3.15E-26 |
| 0.226924106 | 0.131578947 | 1 | 0.115565172 | 3.42E-14 |
| 0.23014769 | 0.184210526 | 0.948741015 | 0.139001836 | 9.03E-17 |
| 1.063928756 | 0.941176471 | 0.169119326 | 0.108483278 | 2.35E-12 |
| 3.120851139 | 1 | 0.364234188 | 0.297496368 | 2.77E-31 |
| 0.662460068 | 0.794117647 | 0.1969587 | 0.104892707 | 5.41E-12 |
| 0.589449468 | 0.617647059 | 0.239962791 | 0.129292253 | 1.91E-14 |
| 0.633556294 | 0.794117647 | 0.259569423 | 0.31521613 | 4.66E-33 |
| 0.377451257 | 0.588235294 | 0.473124729 | 0.241209978 | 1.19E-25 |
| 3.698530186 | 1 | 0.202694454 | 0.169248825 | 1.89E-18 |
| 0.885595389 | 0.911764706 | 0.181063099 | 0.156593444 | 3.51E-17 |
| 1.086007816 | 0.529411765 | 0.865482936 | 0.39203178 | 9.30E-41 |

| | | | | |
|---|---|---|---|---|
| 1.568282789 | 0.941176471 | 0.157771222 | 0.078506797 | 2.48E-09 |
| 0.298679839 | 0.735294118 | 0.304647336 | 0.190695891 | 1.35E-20 |
| 0.746007991 | 0.588235294 | 0.262910773 | 0.124109831 | 6.34E-14 |
| 1.484015684 | 0.970588235 | 0.174858956 | 0.15468556 | 5.44E-17 |
| 10.95376507 | 0.264705882 | 0.7106038 | 0.107083849 | 3.26E-12 |
| 0.810835305 | 0.911764706 | 0.163143507 | 0.167581036 | 2.78E-18 |
| 0.268621614 | 0.558823529 | 0.414557131 | 0.135950982 | 4.11E-15 |
| 0.308499683 | 0.676470588 | 0.237311823 | 0.176220547 | 3.80E-19 |
| 0.457301277 | 0.470588235 | 0.497272126 | 0.135950814 | 4.11E-15 |
| 13.59876301 | 1 | 0.209262245 | 0.101883199 | 1.09E-11 |
| 0.914577384 | 1 | 0.145358145 | 0.122710775 | 8.76E-14 |
| 1.80024258 | 0.941176471 | 0.169821427 | 0.14304716 | 7.98E-16 |
| 0.644772321 | 0.735294118 | 0.381451612 | 0.157079727 | 3.13E-17 |
| 1.276459136 | 0.970588235 | 0.166100403 | 0.202225062 | 9.48E-22 |
| 0.182986384 | 0.235294118 | 0.849898345 | 0.117941315 | 2.64E-13 |
| 0.529709753 | 0.823529412 | 0.192920868 | 0.09519455 | 5.12E-11 |
| 0.130666064 | 0.294117647 | 0.663324164 | 0.097250224 | 3.18E-11 |
| 0.100524857 | 0.352941176 | 0.480648593 | 0.084183962 | 6.61E-10 |
| 14.95860157 | 0.970588235 | 0.224590843 | 0.119862605 | 1.69E-13 |
| 18.21795288 | 0.941176471 | 0.224513306 | 0.115983883 | 4.15E-13 |
| 2.094609564 | 1 | 0.279533731 | 0.184480174 | 5.66E-20 |
| 0.093405764 | 0.323529412 | 0.495844667 | 0.083123338 | 8.46E-10 |
| 0.206404418 | 0.382352941 | 0.714104519 | 0.137699203 | 2.74E-15 |
| 53.66017558 | 1 | 0.219853248 | 0.118732297 | 2.20E-13 |
| 0.308750974 | 0.558823529 | 0.357359946 | 0.216829258 | 3.28E-23 |
| 4.671533697 | 0.970588235 | 0.178598523 | 0.227106726 | 3.07E-24 |
| 0.206544176 | 0.470588235 | 0.401862725 | 0.104408053 | 6.05E-12 |
| 1.132186165 | 0.911764706 | 0.27358935 | 0.209013447 | 1.98E-22 |
| 12.73789397 | 0.970588235 | 0.201662266 | 0.101117282 | 1.30E-11 |
| 0.291923262 | 0.411764706 | 0.364760961 | 0.08571329 | 4.63E-10 |
| 0.300298943 | 0.529411765 | 0.36473328 | 0.174947938 | 5.09E-19 |
| 1.870694956 | 0.970588235 | 0.194821743 | 0.14298046 | 8.11E-16 |
| 0.380504231 | 0.676470588 | 0.398180155 | 0.138757406 | 2.15E-15 |
| 0.268286463 | 0.558823529 | 0.377938901 | 0.151295653 | 1.19E-16 |
| 0.150073373 | 0.323529412 | 0.577025681 | 0.078943465 | 2.24E-09 |
| 1.331716465 | 0.970588235 | 0.166525172 | 0.101534979 | 1.18E-11 |
| 0.186201308 | 0.470588235 | 0.378411193 | 0.105285377 | 4.94E-12 |
| 0.617713971 | 0.705882353 | 0.329253561 | 0.121658272 | 1.12E-13 |
| 0.796855707 | 0.911764706 | 0.199899713 | 0.090183327 | 1.64E-10 |
| 0.067333605 | 0.264705882 | 0.565399175 | 0.105082991 | 5.17E-12 |
| 1.462659123 | 0.882352941 | 0.207617071 | 0.147768463 | 2.69E-16 |
| 1.923450025 | 0.709677419 | 0.552035464 | 0.405128594 | 4.75E-39 |
| 0.467862919 | 0.35483871 | 0.494620624 | 0.135521038 | 4.66E-14 |
| 0.862399429 | 0.516129032 | 0.33468029 | 0.098401561 | 1.33E-10 |
| 1.559490912 | 0.64516129 | 0.847857254 | 0.400372462 | 1.31E-38 |

| | | | | |
|---|---|---|---|---|
| 2.903637078 | 0.774193548 | 0.829684657 | 0.595984305 | 8.19E-57 |
| 0.472596517 | 0.322580645 | 0.829835802 | 0.228954048 | 1.01E-22 |
| 0.804209512 | 0.548387097 | 0.432514171 | 0.172214351 | 1.83E-17 |
| 0.614352721 | 0.419354839 | 0.525510642 | 0.15038002 | 1.94E-15 |
| 7.549845186 | 1 | 0.259593601 | 0.15919196 | 2.96E-16 |
| 0.216116165 | 0.193548387 | 0.785118562 | 0.122660356 | 7.31E-13 |
| 3.323401843 | 1 | 0.159551265 | 0.072332664 | 3.67E-08 |
| 1.096023516 | 0.64516129 | 0.88248964 | 0.470358827 | 4.14E-45 |
| 0.491793845 | 0.387096774 | 0.409914179 | 0.145684549 | 5.30E-15 |
| 1.388538599 | 0.774193548 | 0.243182983 | 0.10101843 | 7.58E-11 |
| 1.298089346 | 0.612903226 | 0.682941707 | 0.321623733 | 2.64E-31 |
| 1.738043723 | 0.774193548 | 0.194724878 | 0.083414512 | 3.35E-09 |
| 1.821968435 | 0.806451613 | 0.509161918 | 0.351076083 | 4.92E-34 |
| 0.857779494 | 0.516129032 | 0.542477581 | 0.262726224 | 7.55E-26 |
| 8.203648453 | 1 | 0.532774491 | 0.560519988 | 1.67E-53 |
| 0.776343998 | 0.483870968 | 0.336818375 | 0.162308474 | 1.52E-16 |
| 1.007687485 | 0.516129032 | 0.387584343 | 0.203334198 | 2.39E-20 |
| 1.679473838 | 0.64516129 | 0.258505888 | 0.095916723 | 2.27E-10 |
| 0.386531552 | 0.290322581 | 0.640025896 | 0.122526153 | 7.52E-13 |
| 1.795858865 | 0.741935484 | 0.259657211 | 0.151429715 | 1.55E-15 |
| 2.504506067 | 0.806451613 | 0.29668035 | 0.146198956 | 4.75E-15 |
| 1.603915711 | 0.580645161 | 0.5657019 | 0.215373223 | 1.84E-21 |
| 2.742191106 | 0.838709677 | 0.528583406 | 0.402235609 | 8.82E-39 |
| 1.051768712 | 0.580645161 | 0.348354594 | 0.132592768 | 8.71E-14 |
| 2.826208179 | 0.903225806 | 0.781494027 | 0.636376942 | 1.38E-60 |
| 6.220087632 | 0.967741935 | 0.232608807 | 0.134816905 | 5.41E-14 |
| 0.846874907 | 0.483870968 | 0.322677002 | 0.138175221 | 2.64E-14 |
| 0.593669407 | 0.387096774 | 0.415741569 | 0.105399052 | 2.96E-11 |
| 0.382566688 | 0.290322581 | 0.580319186 | 0.09209188 | 5.17E-10 |
| 0.459800342 | 0.290322581 | 0.548968697 | 0.133266434 | 7.54E-14 |
| 0.898971994 | 0.612903226 | 0.379710199 | 0.206731792 | 1.16E-20 |
| 1.154132866 | 0.580645161 | 0.345308741 | 0.129724162 | 1.61E-13 |
| 0.537781885 | 0.322580645 | 0.840689006 | 0.188349294 | 5.85E-19 |
| 0.573554755 | 0.322580645 | 0.641505993 | 0.183757155 | 1.56E-18 |
| 0.605052446 | 0.419354839 | 0.497620004 | 0.1320903 | 9.70E-14 |
| 0.948055045 | 0.580645161 | 0.46851495 | 0.193712348 | 1.86E-19 |
| 0.268967297 | 0.258064516 | 0.582106569 | 0.089157264 | 9.72E-10 |
| 2.489419955 | 0.774193548 | 0.308665135 | 0.292199555 | 1.41E-28 |
| 2.626085601 | 0.741935484 | 0.295100645 | 0.130667298 | 1.32E-13 |
| 0.620423741 | 0.483870968 | 0.546352029 | 0.2168426 | 1.34E-21 |
| 0.179654503 | 0.193548387 | 0.893903942 | 0.126940638 | 2.92E-13 |
| 0.867050382 | 0.483870968 | 0.388321009 | 0.211902555 | 3.85E-21 |
| 0.570162852 | 0.290322581 | 0.559151915 | 0.111227298 | 8.47E-12 |
| 0.802201144 | 0.516129032 | 0.303074159 | 0.065388691 | 1.65E-07 |
| 1.581938861 | 0.677419355 | 0.25586518 | 0.12314508 | 6.58E-13 |

| | | | | |
|---|---|---|---|---|
| 1.01786398 | 0.677419355 | 0.550686726 | 0.232124387 | 5.16E-23 |
| 0.260381708 | 0.681818182 | 0.241338295 | 0.073751228 | 1.51E-06 |
| 0.554725226 | 0.863636364 | 0.166547051 | 0.043400807 | 0.000225025 |
| 0.70462508 | 0.727272727 | 0.208673867 | 0.056024751 | 2.77E-05 |
| 2.762424489 | 0.954545455 | 0.160342371 | 0.066066769 | 5.31E-06 |
| 2.026654741 | 0.954545455 | 0.19170836 | 0.086015829 | 2.05E-07 |
| 2.109143431 | 1 | 0.146665704 | 0.033715836 | 0.001148637 |
| 1.545608361 | 1 | 0.194287656 | 0.09512432 | 4.67E-08 |
| 0.706517608 | 0.818181818 | 0.190834216 | 0.066883219 | 4.64E-06 |
| 0.689620795 | 0.727272727 | 0.319688941 | 0.097201024 | 3.33E-08 |
| 1.104680081 | 0.909090909 | 0.180035846 | 0.050817969 | 6.55E-05 |
| 0.827311737 | 0.863636364 | 0.188036747 | 0.072999997 | 1.71E-06 |
| 0.919761384 | 0.863636364 | 0.248809645 | 0.065983876 | 5.38E-06 |
| 0.697877888 | 0.727272727 | 0.201224125 | 0.064164937 | 7.25E-06 |
| 0.895344005 | 0.863636364 | 0.182038862 | 0.075079616 | 1.21E-06 |
| 2.751385579 | 1 | 0.160012846 | 0.078111985 | 7.41E-07 |
| 0.456295724 | 0.772727273 | 0.237721769 | 0.062203098 | 1.00E-05 |
| 0.577390123 | 0.772727273 | 0.213359516 | 0.059470281 | 1.57E-05 |
| 0.705078482 | 0.863636364 | 0.260854779 | 0.099245794 | 2.39E-08 |
| 0.336806517 | 0.636363636 | 0.254291873 | 0.051795639 | 5.57E-05 |
| 1.417344261 | 0.954545455 | 0.200095902 | 0.098879743 | 2.54E-08 |
| 0.74248033 | 0.863636364 | 0.255033962 | 0.086803377 | 1.80E-07 |
| 0.98869659 | 0.909090909 | 0.185328789 | 0.063924721 | 7.54E-06 |
| 0.883915044 | 0.909090909 | 0.181570342 | 0.079058185 | 6.35E-07 |
| 1.382364207 | 0.909090909 | 0.161341809 | 0.056646949 | 2.50E-05 |
| 0.167870491 | 0.5 | 0.312910444 | 0.048002902 | 0.000104492 |
| 1.439853597 | 0.909090909 | 0.175220867 | 0.07124177 | 2.27E-06 |
| 10.40019401 | 1 | 0.170540864 | 0.091485589 | 8.42E-08 |
| 13.07091778 | 1 | 0.174604657 | 0.096698506 | 3.62E-08 |
| 0.371249838 | 0.727272727 | 0.223549336 | 0.043939651 | 0.000205651 |
| 0.642604474 | 0.818181818 | 0.188837213 | 0.072063895 | 1.99E-06 |
| 0.74781608 | 0.863636364 | 0.195968351 | 0.061511283 | 1.12E-05 |
| 6.801256918 | 1 | 0.194776478 | 0.103074058 | 1.29E-08 |
| 4.504669221 | 1 | 0.188564639 | 0.10719521 | 6.62E-09 |
| 6.089490958 | 1 | 0.190778828 | 0.101480682 | 1.67E-08 |
| 35.27807779 | 1 | 0.159958856 | 0.085816107 | 2.11E-07 |
| 0.611567251 | 0.909090909 | 0.232386036 | 0.0804115 | 5.09E-07 |
| 0.478770093 | 0.636363636 | 0.289162874 | 0.061832581 | 1.06E-05 |
| 0.529025545 | 0.636363636 | 0.238321884 | 0.063054376 | 8.70E-06 |
| 0.411123243 | 0.681818182 | 0.308641878 | 0.080786371 | 4.79E-07 |
| 10.61887035 | 1 | 0.175681172 | 0.091053093 | 9.03E-08 |
| 2.872181131 | 0.954545455 | 0.158770861 | 0.071026839 | 2.35E-06 |
| 0.134349667 | 0.454545455 | 0.33431759 | 0.046696701 | 0.000129857 |
| 1.447460324 | 0.954545455 | 0.161984852 | 0.087139896 | 1.70E-07 |
| 0.388342792 | 0.681818182 | 0.242493848 | 0.088278352 | 1.42E-07 |

| | | | | |
|---|---|---|---|---|
| 0.259597598 | 0.454545455 | 0.325845923 | 0.051105933 | 6.24E-05 |
| 0.63612727 | 0.772727273 | 0.23059637 | 0.088954542 | 1.27E-07 |
| 0.760748016 | 0.863636364 | 0.228195195 | 0.074723802 | 1.29E-06 |
| 0.222994331 | 0.454545455 | 0.314738397 | 0.060548583 | 1.31E-05 |
| 0.801546111 | 0.909090909 | 0.172438322 | 0.063918309 | 7.55E-06 |
| 0.859613869 | 0.909090909 | 0.165242508 | 0.066840526 | 4.67E-06 |
| 1.194212123 | 0.590909091 | 0.698099383 | 0.241037775 | 3.15E-18 |
| 0.780101532 | 0.5 | 0.809441598 | 0.249400095 | 8.27E-19 |
| 1.07389859 | 0.5 | 0.823630703 | 0.323222115 | 6.27E-24 |
| 1.028525037 | 0.454545455 | 0.951663972 | 0.327102838 | 3.38E-24 |
| 2.920038276 | 0.636363636 | 0.90044022 | 0.452911208 | 6.34E-33 |
| 1.525986474 | 0.590909091 | 0.885289327 | 0.373723305 | 1.98E-27 |
| 1.71765679 | 0.727272727 | 0.634930919 | 0.337544548 | 6.37E-25 |
| 2.601236411 | 0.772727273 | 0.863584445 | 0.495104288 | 7.46E-36 |
| 16.25517598 | 0.727272727 | 0.929674326 | 0.586030608 | 3.57E-42 |
| 4.575565555 | 0.863636364 | 0.708862509 | 0.400157885 | 2.90E-29 |
| 6.201096211 | 0.818181818 | 0.881713105 | 0.617882837 | 2.17E-44 |
| 2.17626625 | 0.681818182 | 0.631140967 | 0.272851736 | 1.95E-20 |
| 1.593251896 | 0.590909091 | 0.77123764 | 0.340516685 | 3.97E-25 |
| 0.924705861 | 0.5 | 0.828786224 | 0.301526051 | 2.00E-22 |
| 1.121597324 | 0.5 | 0.842751966 | 0.282280802 | 4.33E-21 |
| 0.316028426 | 0.454545455 | 0.955040621 | 0.342101108 | 3.08E-25 |
| 0.523598262 | 0.5 | 0.770065825 | 0.256621561 | 2.61E-19 |
| 1.582222181 | 0.681818182 | 0.885316854 | 0.403351414 | 1.74E-29 |
| 0.648127855 | 0.5 | 0.776965241 | 0.320473771 | 9.73E-24 |
| 0.546388767 | 0.454545455 | 0.865045626 | 0.21560318 | 1.83E-16 |
| 0.761599025 | 0.545454545 | 0.789228458 | 0.223771668 | 4.97E-17 |
| 4.007900956 | 0.545454545 | 0.896935313 | 0.34478807 | 2.00E-25 |
| 1.038948994 | 0.590909091 | 0.904782784 | 0.372123113 | 2.55E-27 |
| 6.359548297 | 0.818181818 | 0.81982222 | 0.602992439 | 2.36E-43 |
| 0.583721848 | 0.409090909 | 0.964926639 | 0.314481723 | 2.53E-23 |
| 0.5199903 | 0.5 | 0.903278226 | 0.300864735 | 2.23E-22 |
| 3.307362903 | 0.681818182 | 0.832397066 | 0.413092294 | 3.67E-30 |
| 34.54072314 | 0.818181818 | 0.881969133 | 0.557228001 | 3.59E-40 |
| 12.03036507 | 0.863636364 | 0.87290601 | 0.662693319 | 1.64E-47 |
| 1.892240326 | 0.636363636 | 0.897208039 | 0.423926669 | 6.50E-31 |
| 3.702184919 | 0.636363636 | 0.885379703 | 0.418568848 | 1.53E-30 |
| 0.420098886 | 0.409090909 | 0.963213538 | 0.34947144 | 9.49E-26 |
| 0.817580618 | 0.5 | 0.761472147 | 0.228593057 | 2.30E-17 |
| 2.772037078 | 0.545454545 | 0.747085576 | 0.36981323 | 3.69E-27 |
| 2.913868222 | 0.727272727 | 0.748896026 | 0.476312263 | 1.50E-34 |
| 0.695865259 | 0.409090909 | 0.924283936 | 0.226421189 | 3.25E-17 |
| 2.055936279 | 0.772727273 | 0.918426217 | 0.583893446 | 5.03E-42 |
| 4.345932357 | 0.863636364 | 0.868306113 | 0.64756319 | 1.86E-46 |
| 3.448950717 | 0.590909091 | 0.872766952 | 0.343897697 | 2.31E-25 |

| | | | | |
|---|---|---|---|---|
| 1.017717942 | 0.545454545 | 0.899325889 | 0.353695374 | 4.84E-26 |
| 0.760063702 | 0.454545455 | 0.88277975 | 0.323282389 | 6.21E-24 |
| 0.909748577 | 0.5 | 0.924755388 | 0.354820784 | 4.04E-26 |
| 1.278702985 | 0.545454545 | 0.834678252 | 0.361373943 | 1.42E-26 |
| 1.370605264 | 0.590909091 | 0.903708262 | 0.430801039 | 2.17E-31 |
| 4.43544547 | 0.545454545 | 0.90314556 | 0.347592273 | 1.28E-25 |
| 2.879305059 | 0.727272727 | 0.751980988 | 0.421702978 | 9.28E-31 |
| 0.572380209 | 0.409090909 | 0.932455545 | 0.276012965 | 1.18E-20 |
| 0.750437109 | 0.5 | 0.890492485 | 0.286996021 | 2.04E-21 |
| 0.93557858 | 0.636363636 | 0.765024565 | 0.369184291 | 4.08E-27 |
| 1.507530264 | 0.636363636 | 0.696652425 | 0.260566935 | 1.39E-19 |
| 13.00654659 | 0.974358974 | 0.297374459 | 0.700163771 | 0 |
| 0.5672978 | 0.554206028 | 0.394564056 | 0.406056833 | 0 |
| 1.228494235 | 0.663517769 | 0.343019841 | 0.382748516 | 0 |
| 29.19317249 | 0.993702204 | 0.26344481 | 0.841904824 | 0 |
| 0.263733161 | 0.324786325 | 0.615602627 | 0.230604755 | 0 |
| 2.244809604 | 0.743589744 | 0.44058994 | 0.547373051 | 0 |
| 1.353363818 | 0.671614935 | 0.339372455 | 0.472281987 | 0 |
| 13.35459779 | 0.968960864 | 0.409629724 | 0.728940202 | 0 |
| 1.648036834 | 0.709401709 | 0.343837175 | 0.443906941 | 0 |
| 1.556613594 | 0.707602339 | 0.430557509 | 0.460136865 | 0 |
| 0.536045886 | 0.512820513 | 0.40138424 | 0.374808977 | 0 |
| 15.34147683 | 0.976608187 | 0.480871641 | 0.811892736 | 0 |
| 9.51695892 | 0.892487629 | 0.486486591 | 0.667029635 | 0 |
| 0.431565791 | 0.494826811 | 0.436985033 | 0.35021845 | 0 |
| 0.494878928 | 0.518218623 | 0.345935541 | 0.349349727 | 0 |
| 0.833114563 | 0.560953666 | 0.371010241 | 0.354481067 | 0 |
| 0.31013219 | 0.446243815 | 0.549735902 | 0.334475302 | 0 |
| 0.718969619 | 0.561853351 | 0.347518969 | 0.366288673 | 0 |
| 0.50786087 | 0.529464687 | 0.388382918 | 0.357844194 | 0 |
| 1.207053997 | 0.643274854 | 0.528649418 | 0.504349613 | 0 |
| 2.042364173 | 0.743139901 | 0.330669287 | 0.499909536 | 0 |
| 2.190270618 | 0.75888439 | 0.504727594 | 0.591282161 | 0 |
| 0.581665365 | 0.547008547 | 0.436682189 | 0.366374169 | 0 |
| 0.436989422 | 0.504723347 | 0.404026544 | 0.287330718 | 0 |
| 0.416593759 | 0.513720198 | 0.43238156 | 0.325943908 | 0 |
| 0.52523277 | 0.448043185 | 0.472800104 | 0.310955758 | 0 |
| 1.545659885 | 0.668466037 | 0.438524106 | 0.46552623 | 0 |
| 0.447216388 | 0.521367521 | 0.388950826 | 0.339596054 | 0 |
| 0.224553074 | 0.401709402 | 0.554078978 | 0.299454905 | 0 |
| 0.41361939 | 0.48582996 | 0.451610196 | 0.337613687 | 0 |
| 0.206093316 | 0.358974359 | 0.543164023 | 0.257194137 | 0 |
| 0.507650387 | 0.524966262 | 0.581205218 | 0.416251293 | 0 |
| 0.850141335 | 0.611336032 | 0.549294943 | 0.469978581 | 0 |
| 2.658434164 | 0.788573999 | 0.527531642 | 0.607656158 | 0 |

| | | | | |
|---|---|---|---|---|
| 0.440693498 | 0.516419253 | 0.369658477 | 0.348065591 | 0 |
| 1.37372863 | 0.65677013 | 0.337154132 | 0.421820057 | 0 |
| 0.534576643 | 0.55105713 | 0.324600502 | 0.349460875 | 0 |
| 1.148975589 | 0.644174539 | 0.529198478 | 0.480703988 | 0 |
| 1.439163404 | 0.69545659 | 0.385920477 | 0.475481242 | 0 |
| 3.096610282 | 0.805218174 | 0.493978546 | 0.588844002 | 0 |
| 0.176640079 | 0.32568601 | 0.541943143 | 0.234843569 | 0 |
| 3.036510966 | 0.808367072 | 0.308535596 | 0.50975983 | 0 |
| 0.14068288 | 0.316239316 | 0.590440028 | 0.24708908 | 0 |
| 3.51321288 | 0.79577148 | 0.479790879 | 0.600275346 | 0 |
| 1.136026165 | 0.647773279 | 0.435038819 | 0.401012881 | 0 |
| 13.7445282 | 0.937921727 | 0.291570543 | 0.671509334 | 0 |
| 0.395944742 | 0.499325236 | 0.504613759 | 0.366931416 | 0 |
| 36.22002748 | 0.992352677 | 0.434776459 | 0.836517062 | 0 |
| 0.199041151 | 0.388663968 | 0.531057222 | 0.291627567 | 0 |
| 0.448881829 | 0.520017994 | 0.390397929 | 0.341735982 | 0 |
| 0.518877368 | 0.315789474 | 0.609173644 | 0.071350755 | 8.81E-06 |
| 0.159937756 | 0.263157895 | 0.907842298 | 0.123428386 | 4.99E-09 |
| 0.386070883 | 0.526315789 | 0.408928958 | 0.115721019 | 1.50E-08 |
| 0.135680422 | 0.315789474 | 0.595263842 | 0.073767213 | 6.20E-06 |
| 0.900193599 | 0.684210526 | 0.279299781 | 0.113664404 | 2.01E-08 |
| 0.354369416 | 0.473684211 | 0.982705377 | 0.396304121 | 9.00E-26 |
| 0.543590924 | 0.578947368 | 0.864384493 | 0.303548283 | 4.28E-20 |
| 0.621938111 | 0.578947368 | 0.704047432 | 0.265734368 | 8.86E-18 |
| 1.018906358 | 0.526315789 | 0.86914417 | 0.305849927 | 3.10E-20 |
| 0.349110245 | 0.421052632 | 0.765656193 | 0.161041742 | 2.37E-11 |
| 0.050385333 | 0.210526316 | 1 | 0.187134054 | 5.88E-13 |
| 3.448029448 | 0.526315789 | 0.543495611 | 0.18433188 | 8.75E-13 |
| 0.208969421 | 0.421052632 | 0.685154097 | 0.125987911 | 3.46E-09 |
| 1.971789799 | 0.684210526 | 0.27867629 | 0.122827246 | 5.43E-09 |
| 4.064461473 | 0.842105263 | 0.244979611 | 0.098925982 | 1.66E-07 |
| 0.361734654 | 0.526315789 | 0.853735636 | 0.280288898 | 1.14E-18 |
| 2.074635486 | 0.578947368 | 0.765794222 | 0.305953557 | 3.05E-20 |
| 0.959017083 | 0.473684211 | 0.410438898 | 0.066574666 | 1.76E-05 |
| 2.249981213 | 0.789473684 | 0.323717646 | 0.123736164 | 4.77E-09 |
| 0.254561837 | 0.421052632 | 0.675234227 | 0.118252075 | 1.04E-08 |
| 0.894566646 | 0.473684211 | 0.536649056 | 0.200955431 | 8.33E-14 |
| 3.144944548 | 0.684210526 | 0.638103258 | 0.282486465 | 8.34E-19 |
| 0.347957437 | 0.368421053 | 0.880529526 | 0.260146691 | 1.95E-17 |
| 0.455460583 | 0.526315789 | 0.525655871 | 0.146596332 | 1.84E-10 |
| 0.935656553 | 0.578947368 | 0.886863813 | 0.419028966 | 3.66E-27 |
| 0.06994209 | 0.263157895 | 0.816458208 | 0.07796462 | 3.38E-06 |
| 0.384196621 | 0.421052632 | 0.458665663 | 0.064353012 | 2.43E-05 |
| 0.10146658 | 0.263157895 | 0.72132449 | 0.065531082 | 2.05E-05 |
| 0.166899624 | 0.368421053 | 0.525687769 | 0.06406285 | 2.54E-05 |

| | | | | |
|---|---|---|---|---|
| 0.299709079 | 0.315789474 | 0.657591604 | 0.097180553 | 2.13E-07 |
| 0.238755726 | 0.421052632 | 0.85353056 | 0.20332282 | 5.96E-14 |
| 1.171719172 | 0.736842105 | 0.273857987 | 0.144705851 | 2.41E-10 |
| 1.350092643 | 0.631578947 | 0.968111121 | 0.467369613 | 4.02E-30 |
| 0.290896759 | 0.368421053 | 0.748451634 | 0.137755926 | 6.48E-10 |
| 0.430198928 | 0.368421053 | 0.545264219 | 0.155137916 | 5.48E-11 |
| 0.197724196 | 0.421052632 | 0.648893827 | 0.111669074 | 2.67E-08 |
| 0.263802983 | 0.421052632 | 0.582710036 | 0.07976095 | 2.61E-06 |
| 1.754263052 | 0.631578947 | 0.342390215 | 0.164400877 | 1.47E-11 |
| 0.124454389 | 0.210526316 | 0.92408635 | 0.183723355 | 9.53E-13 |
| 0.341340367 | 0.473684211 | 0.431021575 | 0.082912506 | 1.66E-06 |
| 0.101354109 | 0.263157895 | 0.772962039 | 0.082422468 | 1.78E-06 |
| 1.521952739 | 0.736842105 | 0.368172373 | 0.15822457 | 3.54E-11 |
| 1.005645489 | 0.526315789 | 0.436555147 | 0.184596421 | 8.43E-13 |
| 0.109018505 | 0.315789474 | 0.732710591 | 0.125299724 | 3.82E-09 |
| 0.476266031 | 0.473684211 | 0.421829903 | 0.100765422 | 1.27E-07 |
| 3.993060027 | 0.894736842 | 0.975580484 | 0.791890712 | 4.86E-50 |
| 0.092837532 | 0.315789474 | 0.708120205 | 0.08191776 | 1.91E-06 |
| 0.470538745 | 0.526315789 | 0.564646816 | 0.154117346 | 6.34E-11 |
| 0.087786645 | 0.263157895 | 0.728547442 | 0.057407787 | 6.71E-05 |
| 0.108046327 | 0.315789474 | 0.805770683 | 0.077725019 | 3.50E-06 |
| 1.149223923 | 0.941176471 | 0.408102717 | 0.355480803 | 2.67E-21 |
| 0.856000945 | 0.764705882 | 0.385383855 | 0.236374503 | 1.15E-14 |
| 3.531254397 | 0.941176471 | 0.56602659 | 0.465344709 | 2.05E-27 |
| 1.323538053 | 0.705882353 | 0.386493697 | 0.162170057 | 1.63E-10 |
| 0.414257795 | 0.764705882 | 0.49525446 | 0.184516267 | 9.12E-12 |
| 0.057925327 | 0.411764706 | 0.585167492 | 0.082042227 | 5.50E-06 |
| 3.284733062 | 0.882352941 | 0.881932613 | 0.685689467 | 1.08E-39 |
| 0.552246882 | 0.764705882 | 0.960297811 | 0.623887096 | 3.03E-36 |
| 3.643666896 | 0.823529412 | 0.858721817 | 0.506061311 | 1.11E-29 |
| 0.234295552 | 0.411764706 | 0.640475583 | 0.321285063 | 2.14E-19 |
| 0.497571222 | 0.823529412 | 0.66211405 | 0.394715063 | 1.75E-23 |
| 0.817472029 | 0.882352941 | 0.302790001 | 0.151744317 | 6.28E-10 |
| 1.697137231 | 0.941176471 | 0.949966066 | 0.80135274 | 3.76E-46 |
| 2.444516289 | 0.823529412 | 0.931467211 | 0.645378085 | 1.92E-37 |
| 1.097670939 | 0.882352941 | 0.352723503 | 0.247839841 | 2.65E-15 |
| 1.193143522 | 0.882352941 | 0.273366525 | 0.151516697 | 6.46E-10 |
| 0.461793356 | 0.764705882 | 0.390848258 | 0.14684322 | 1.18E-09 |
| 0.837091766 | 0.941176471 | 0.376008499 | 0.235748124 | 1.25E-14 |
| 0.155978182 | 0.588235294 | 0.422564057 | 0.22674957 | 3.98E-14 |
| 0.415968812 | 0.764705882 | 0.892180259 | 0.50336232 | 1.57E-29 |
| 0.478306637 | 0.882352941 | 0.295103563 | 0.127459413 | 1.46E-08 |
| 0.240469107 | 0.705882353 | 0.428761159 | 0.16676787 | 9.01E-11 |
| 1.034188126 | 0.823529412 | 0.310107857 | 0.165800379 | 1.02E-10 |
| 0.213549011 | 0.529411765 | 0.540390695 | 0.112863506 | 9.74E-08 |

| | | | | |
|---|---|---|---|---|
| 6.735567228 | 1 | 0.306387312 | 0.140933398 | 2.54E-09 |
| 0.163126554 | 0.470588235 | 0.893947402 | 0.295020902 | 6.21E-18 |
| 0.25712133 | 0.764705882 | 0.773415497 | 0.418223627 | 8.60E-25 |
| 0.329489866 | 0.705882353 | 0.495667516 | 0.259018684 | 6.30E-16 |
| 0.989376375 | 0.882352941 | 0.34310063 | 0.22776688 | 3.49E-14 |
| 0.425722816 | 0.823529412 | 0.878101224 | 0.676548635 | 3.51E-39 |
| 0.910807561 | 0.941176471 | 0.368970835 | 0.210506022 | 3.21E-13 |
| 0.348476026 | 0.588235294 | 0.412252468 | 0.144264526 | 1.65E-09 |
| 7.371557221 | 0.941176471 | 0.588354199 | 0.614468341 | 1.01E-35 |
| 0.167721356 | 0.705882353 | 0.350226641 | 0.104986128 | 2.72E-07 |
| 0.078289358 | 0.470588235 | 0.709352057 | 0.125170184 | 1.97E-08 |
| 0.748437132 | 0.647058824 | 0.770113164 | 0.338181239 | 2.45E-20 |
| 1.508421728 | 0.941176471 | 0.34030885 | 0.235921418 | 1.22E-14 |
| 0.211821619 | 0.764705882 | 0.444888885 | 0.151796463 | 6.23E-10 |
| 4.36622954 | 0.705882353 | 0.47483414 | 0.199572012 | 1.31E-12 |
| 0.417238102 | 0.823529412 | 0.473347463 | 0.289822982 | 1.21E-17 |
| 0.556453592 | 0.705882353 | 0.987950601 | 0.628774351 | 1.62E-36 |
| 2.127278674 | 0.764705882 | 0.597433155 | 0.388010758 | 4.13E-23 |
| 73.84489486 | 1 | 0.519569054 | 0.646000135 | 1.77E-37 |
| 12.6726714 | 0.941176471 | 0.603572667 | 0.457468088 | 5.63E-27 |
| 1.146669399 | 0.882352941 | 0.417031162 | 0.202762768 | 8.70E-13 |
| 0.082227285 | 0.470588235 | 0.530490415 | 0.11030072 | 1.36E-07 |
| 0.136321505 | 0.470588235 | 0.56757058 | 0.094955527 | 1.01E-06 |
| 0.054801122 | 0.294117647 | 0.865899366 | 0.073990089 | 1.59E-05 |
| 0.197345982 | 0.588235294 | 0.767126725 | 0.381458511 | 9.56E-23 |
| 0.129320947 | 0.529411765 | 0.869641254 | 0.359043745 | 1.69E-21 |
| 1.723968876 | 0.623847926 | 0.311627336 | 0.292143489 | 0 |
| 0.852048277 | 0.788018433 | 0.230530172 | 0.29647969 | 0 |
| 0.94312469 | 0.836981567 | 0.260095041 | 0.491870438 | 0 |
| 3.794956849 | 0.987903226 | 0.204043705 | 0.430984007 | 0 |
| 2.777139567 | 0.975806452 | 0.243266657 | 0.554025593 | 0 |
| 2.39807119 | 0.969470046 | 0.167000967 | 0.430536667 | 0 |
| 0.591851143 | 0.709677419 | 0.217496881 | 0.342956351 | 0 |
| 0.552051074 | 0.665898618 | 0.25670582 | 0.306898504 | 0 |
| 1.770362004 | 0.942396313 | 0.215304722 | 0.427567728 | 0 |
| 0.853753812 | 0.824308756 | 0.220217627 | 0.404556291 | 0 |
| 1.324767631 | 0.902073733 | 0.206601233 | 0.301280071 | 0 |
| 1.457769004 | 0.94124424 | 0.28859474 | 0.571259202 | 0 |
| 1.126943622 | 0.838709677 | 0.290140843 | 0.43295029 | 0 |
| 0.764681673 | 0.778801843 | 0.215628056 | 0.462946999 | 0 |
| 0.681030948 | 0.733870968 | 0.241702116 | 0.311105267 | 0 |
| 0.777990456 | 0.742511521 | 0.281014439 | 0.45952395 | 0 |
| 0.35731296 | 0.542626728 | 0.317473117 | 0.329326121 | 0 |
| 3.191351754 | 0.903225806 | 0.174075717 | 0.330675355 | 0 |
| 0.549565759 | 0.680299539 | 0.26285065 | 0.313616882 | 0 |

| | | | | |
|---|---|---|---|---|
| 2.088907536 | 0.96140553 | 0.268251129 | 0.611405531 | 0 |
| 1.191927498 | 0.90264977 | 0.227668673 | 0.485483392 | 0 |
| 2.243700322 | 0.933179724 | 0.232884757 | 0.405765487 | 0 |
| 16.12046553 | 0.990207373 | 0.237646773 | 0.548946498 | 0 |
| 20.46984299 | 0.994239631 | 0.245174307 | 0.584753734 | 0 |
| 0.836707707 | 0.676267281 | 0.230563704 | 0.336650955 | 0 |
| 0.446108457 | 0.60656682 | 0.327882768 | 0.376306248 | 0 |
| 1.477513028 | 0.910138249 | 0.214725694 | 0.439241775 | 0 |
| 10.03336144 | 0.998271889 | 0.261317271 | 0.586993994 | 0 |
| 6.723959964 | 0.995967742 | 0.255261244 | 0.598138763 | 0 |
| 8.583945719 | 0.997695853 | 0.247320533 | 0.566176447 | 0 |
| 57.43258045 | 0.999423963 | 0.231432689 | 0.5384189 | 0 |
| 0.956073381 | 0.790898618 | 0.326011083 | 0.503201121 | 0 |
| 0.398964383 | 0.59735023 | 0.271610244 | 0.3723101 | 0 |
| 0.315997457 | 0.52937788 | 0.316571831 | 0.322125799 | 0 |
| 0.515222098 | 0.631336406 | 0.303490168 | 0.326786157 | 0 |
| 0.573883173 | 0.672235023 | 0.253434042 | 0.428407659 | 0 |
| 0.738429068 | 0.785714286 | 0.253038227 | 0.445210628 | 0 |
| 0.416899814 | 0.589861751 | 0.311936019 | 0.391657135 | 0 |
| 1.083042971 | 0.843317972 | 0.227209833 | 0.474188992 | 0 |
| 15.95361387 | 0.991359447 | 0.239063008 | 0.520250294 | 0 |
| 0.719069134 | 0.745391705 | 0.210080403 | 0.234550715 | 0 |
| 0.370274782 | 0.516705069 | 0.29911845 | 0.25792354 | 0 |
| 3.753905093 | 0.991359447 | 0.194576599 | 0.431180262 | 0 |
| 0.720024156 | 0.683179724 | 0.240894341 | 0.322234973 | 0 |
| 0.909292207 | 0.815092166 | 0.236702889 | 0.284270438 | 0 |
| 0.765493186 | 0.749423963 | 0.265212497 | 0.516913779 | 0 |
| 1.125112805 | 0.841013825 | 0.259437746 | 0.454391611 | 0 |
| 0.420377545 | 0.582949309 | 0.279026963 | 0.317270866 | 0 |
| 1.291505263 | 0.919930876 | 0.247411946 | 0.510786469 | 0 |
| 1.528638844 | 0.93375576 | 0.255500858 | 0.514872969 | 0 |
| 2.537722705 | 0.740963855 | 0.241413492 | 0.455187725 | 0 |
| 6.422219875 | 0.885542169 | 0.229732436 | 0.380877613 | 0 |
| 0.371581474 | 0.289759036 | 0.555294868 | 0.233742903 | 0 |
| 0.700063838 | 0.437349398 | 0.367181367 | 0.312190112 | 0 |
| 0.650112374 | 0.436746988 | 0.436659505 | 0.331976737 | 0 |
| 2.014314393 | 0.670481928 | 0.17777553 | 0.168416909 | 6.99E-246 |
| 0.458777916 | 0.363855422 | 0.388062048 | 0.265493623 | 0 |
| 0.834927521 | 0.472891566 | 0.370237804 | 0.367006533 | 0 |
| 1.176798399 | 0.462650602 | 0.487432651 | 0.336023793 | 0 |
| 0.628777195 | 0.41686747 | 0.358280609 | 0.220769833 | 0 |
| 0.715163798 | 0.427108434 | 0.276844154 | 0.188820872 | 3.75E-277 |
| 0.960163125 | 0.526506024 | 0.364815904 | 0.382291729 | 0 |
| 1.174408572 | 0.58253012 | 0.308832056 | 0.330994347 | 0 |
| 7.272932082 | 0.906626506 | 0.334312992 | 0.575005627 | 0 |

| | | | | |
|---|---|---|---|---|
| 1.149219599 | 0.578915663 | 0.48596592 | 0.461331218 | 0 |
| 1.637263727 | 0.645783133 | 0.183148595 | 0.144978494 | 1.89E-210 |
| 0.546255041 | 0.38373494 | 0.33393305 | 0.218317694 | 4.44659081257122e-323 |
| 3.560701289 | 0.803614458 | 0.221676423 | 0.356755934 | 0 |
| 1.725967966 | 0.668072289 | 0.491205706 | 0.536603034 | 0 |
| 1.034811459 | 0.521686747 | 0.372489917 | 0.332995187 | 0 |
| 2.329258163 | 0.736746988 | 0.354969651 | 0.414396852 | 0 |
| 0.835799885 | 0.513855422 | 0.48381528 | 0.374540172 | 0 |
| 0.283684888 | 0.256024096 | 0.471727473 | 0.209305487 | 0.00E+00 |
| 0.393616487 | 0.309036145 | 0.412616979 | 0.225723884 | 0 |
| 4.118751119 | 0.842771084 | 0.389627888 | 0.606586065 | 0 |
| 0.315180001 | 0.270481928 | 0.470391834 | 0.208762522 | 4.16E-308 |
| 0.827290564 | 0.509638554 | 0.390568789 | 0.302013737 | 0 |
| 1.180166475 | 0.541566265 | 0.553107257 | 0.437328449 | 0 |
| 0.718932704 | 0.412650602 | 0.406811541 | 0.253560893 | 0 |
| 2.485232761 | 0.74939759 | 0.494853292 | 0.591270066 | 0 |
| 0.967979652 | 0.521686747 | 0.332836596 | 0.283196856 | 0 |
| 0.413521224 | 0.343373494 | 0.414221873 | 0.251736946 | 0 |
| 0.541279728 | 0.374698795 | 0.462064852 | 0.263284166 | 0 |
| 0.265606538 | 0.221686747 | 0.577237166 | 0.194208649 | 1.78E-285 |
| 0.732963211 | 0.471686747 | 0.357735053 | 0.318444354 | 0 |
| 1.015013914 | 0.526506024 | 0.522826728 | 0.446918627 | 0 |
| 0.500116583 | 0.371686747 | 0.446344248 | 0.265050582 | 0 |
| 0.530556985 | 0.345180723 | 0.43944855 | 0.239491176 | 0 |
| 0.541500849 | 0.370481928 | 0.524570331 | 0.288266022 | 0 |
| 0.476213848 | 0.293975904 | 0.424516375 | 0.194929936 | 1.36E-286 |
| 0.740496757 | 0.437951807 | 0.309811393 | 0.264015924 | 0 |
| 0.525741288 | 0.371084337 | 0.444452127 | 0.260351713 | 0 |
| 0.560679793 | 0.403614458 | 0.396129817 | 0.264342293 | 0 |
| 0.362263705 | 0.286746988 | 0.532018344 | 0.209786443 | 0.00E+00 |
| 0.636860062 | 0.394578313 | 0.318000341 | 0.199388736 | 1.66E-293 |
| 0.470504205 | 0.359036145 | 0.547749327 | 0.288725272 | 0 |
| 1.219260586 | 0.579518072 | 0.209053507 | 0.238046003 | 0 |
| 1.050981769 | 0.56686747 | 0.372768153 | 0.428539591 | 0 |
| 1.729574411 | 0.657228916 | 0.428012715 | 0.500874557 | 0 |
| 0.211394186 | 0.215060241 | 0.568427019 | 0.164917188 | 1.47E-240 |
| 1.210970699 | 0.607929515 | 0.425302512 | 0.31189567 | 0 |
| 7.464440156 | 0.794052863 | 0.614485745 | 0.50812815 | 0 |
| 1.017581296 | 0.577092511 | 0.354716944 | 0.305827268 | 0 |
| 0.388666334 | 0.348017621 | 0.47787792 | 0.262373733 | 6.89E-293 |
| 1.671804534 | 0.666299559 | 0.353563767 | 0.386508184 | 0 |
| 0.564740176 | 0.422907489 | 0.515153638 | 0.269362309 | 3.94E-301 |
| 0.553040564 | 0.373348018 | 0.562702509 | 0.248264647 | 2.48E-276 |
| 1.103724902 | 0.546255507 | 0.436483959 | 0.401542524 | 0 |
| 1.452582627 | 0.558370044 | 0.42115924 | 0.341489481 | 0 |

| | | | | | |
|---|---|---|---|---|---|
| 2.963153834 | | 0.773127753 | 0.390292748 | 0.536299685 | 0 |
| 1.466664144 | | 0.625550661 | 0.33746996 | 0.20505613 | 2.55E-226 |
| 0.957679358 | | 0.558370044 | 0.399292487 | 0.367831223 | 0 |
| 0.917247823 | | 0.539647577 | 0.34674969 | 0.238201812 | 1.36E-264 |
| 0.590475667 | | 0.428414097 | 0.411682958 | 0.262369595 | 6.97E-293 |
| 0.300745242 | | 0.296255507 | 0.663303608 | 0.205866185 | 3.01E-227 |
| 10.21006848 | | 0.899779736 | 0.570641943 | 0.67706517 | 0 |
| 1.637263666 | | 0.657488987 | 0.2718305 | 0.192296974 | 9.57E-212 |
| 0.433641601 | | 0.375550661 | 0.511867013 | 0.280448652 | 2.90273802489717e-314 |
| 1.974590115 | | 0.715859031 | 0.292200287 | 0.348684613 | 0 |
| 0.703550847 | | 0.471365639 | 0.680551791 | 0.366125136 | 0 |
| 1.759775701 | | 0.648678414 | 0.255704729 | 0.347121293 | 0 |
| 1.400173831 | | 0.65969163 | 0.561174952 | 0.510929196 | 0 |
| 0.310775701 | | 0.332599119 | 0.489038608 | 0.230569775 | 9.99E-256 |
| 0.497503327 | | 0.378854626 | 0.546718349 | 0.267323515 | 1.01E-298 |
| 5.453036773 | | 0.827092511 | 0.210588339 | 0.197919662 | 3.71E-218 |
| 0.456473464 | | 0.386563877 | 0.660815049 | 0.277055486 | 3.0896611260693e-310 |
| 0.443392176 | | 0.407488987 | 0.398953925 | 0.292578452 | 0 |
| 0.329046274 | | 0.310572687 | 0.685309901 | 0.213904225 | 1.77E-236 |
| 0.429197623 | | 0.385462555 | 0.476060869 | 0.24199427 | 5.20E-269 |
| 0.377076022 | | 0.368942731 | 0.579631127 | 0.259399207 | 2.18E-289 |
| 0.870017038 | | 0.546255507 | 0.519106841 | 0.386181633 | 0 |
| 0.428480854 | | 0.372246696 | 0.484399878 | 0.257459343 | 4.15E-287 |
| 0.328836993 | | 0.341409692 | 0.632311776 | 0.251349871 | 6.07E-280 |
| 1.395267857 | | 0.672907489 | 0.539488994 | 0.518590223 | 0 |
| 2.177191151 | | 0.628854626 | 0.689825683 | 0.483669746 | 0 |
| 0.849803172 | | 0.513215859 | 0.429946156 | 0.225024845 | 2.63E-249 |
| 0.98570625 | | 0.487885463 | 0.429341852 | 0.319611509 | 0 |
| 2.883763788 | | 0.796255507 | 0.445113456 | 0.47180183 | 0 |
| 0.682441732 | | 0.465859031 | 0.485876326 | 0.285949679 | 8.31512481950818e-321 |
| 0.586331213 | | 0.43061674 | 0.591197064 | 0.314351509 | 0 |
| 0.951065394 | | 0.544052863 | 0.669672352 | 0.422873418 | 0 |
| 0.807835422 | | 0.502202643 | 0.463350908 | 0.250002791 | 2.29E-278 |
| 0.968178595 | | 0.581497797 | 0.471173047 | 0.411772113 | 0 |
| 0.508051495 | | 0.356828194 | 0.638777504 | 0.222594716 | 1.70E-246 |
| 0.35464523 | | 0.333700441 | 0.550551227 | 0.245414515 | 5.30E-273 |
| 0.488260224 | | 0.400881057 | 0.404830375 | 0.278523744 | 5.59965127401604e-312 |
| 0.449843559 | | 0.381057269 | 0.403241955 | 0.269802995 | 1.19E-301 |
| 0.867085773 | | 0.531938326 | 0.564910772 | 0.356086561 | 0 |
| 0.759001113 | | 0.501101322 | 0.44004036 | 0.37075396 | 0 |
| 0.577987015 | | 0.45814978 | 0.409335345 | 0.23430855 | 4.57E-260 |
| 1.11729689 | | 0.287371134 | 0.154895011 | 0.006055539 | 2.28E-07 |
| 0.417958708 | | 0.113402062 | 0.375943466 | 0.037809527 | 1.75E-38 |
| 0.266412086 | | 0.108247423 | 0.432935569 | 0.038881404 | 1.56E-39 |
| 0.858582 | | 0.192010309 | 0.501614098 | 0.109502787 | 1.46E-109 |

| | | | | |
|---|---|---|---|---|
| 1.628480386 | 0.328608247 | 0.434540856 | 0.162771631 | 1.14E-163 |
| 2.680771863 | 0.550257732 | 0.139644976 | 0.043612645 | 3.59E-44 |
| 5.519213465 | 0.75257732 | 0.065172591 | 0.050910325 | 2.44E-51 |
| 1.502739041 | 0.322164948 | 0.283856884 | 0.072640518 | 8.74E-73 |
| 1.033403408 | 0.24742268 | 0.180205543 | 0.007405952 | 1.04E-08 |
| 0.663870143 | 0.207474227 | 0.317869728 | 0.044515672 | 4.67E-45 |
| 2.417870562 | 0.434278351 | 0.289471088 | 0.106576915 | 1.27E-106 |
| 1.882859593 | 0.353092784 | 0.24580386 | 0.066435686 | 1.21E-66 |
| 1.125411852 | 0.292525773 | 0.191001777 | 0.040873319 | 1.74E-41 |
| 0.969117914 | 0.266752577 | 0.40648025 | 0.126356129 | 1.49E-126 |
| 0.5861946 | 0.154639175 | 0.404266932 | 0.034874998 | 1.31E-35 |
| 2.370403462 | 0.414948454 | 0.322550341 | 0.148403696 | 5.87E-149 |
| 0.879586141 | 0.255154639 | 0.29664713 | 0.088965915 | 5.23E-89 |
| 0.324572884 | 0.092783505 | 0.648895607 | 0.0649701 | 3.39E-65 |
| 0.957491808 | 0.221649485 | 0.30817823 | 0.076998211 | 4.18E-77 |
| 0.680460344 | 0.197164948 | 0.460372846 | 0.102878524 | 6.50E-103 |
| 2.95658206 | 0.420103093 | 0.316079535 | 0.130153698 | 2.13E-130 |
| 0.236608663 | 0.086340206 | 0.556221878 | 0.042621314 | 3.37E-43 |
| 0.762579692 | 0.217783505 | 0.285166762 | 0.090323065 | 2.32E-90 |
| 0.443548195 | 0.144329897 | 0.297695087 | 0.035361799 | 4.37E-36 |
| 1.37328478 | 0.300257732 | 0.27089478 | 0.059382449 | 1.11E-59 |
| 0.875106408 | 0.233247423 | 0.244382645 | 0.064345459 | 1.40E-64 |
| 0.696638231 | 0.204896907 | 0.206407033 | 0.033137415 | 6.58E-34 |
| 1.556451209 | 0.274484536 | 0.342687207 | 0.079853082 | 6.12E-80 |
| 7.788775571 | 0.855670103 | 0.097683445 | 0.001819418 | 0.004575824 |
| 24.51408367 | 0.988402062 | 0.128114469 | 0.115135047 | 3.16E-115 |
| 6.692538119 | 0.81443299 | 0.140199218 | 0.037835885 | 1.65E-38 |
| 1.929760593 | 0.445876289 | 0.096783193 | 0.000810646 | 0.058427283 |
| 3.655250434 | 0.652061856 | 0.129290764 | 0.027023068 | 6.28E-28 |
| 5.75203556 | 0.75128866 | 0.130393285 | 0.038236284 | 6.70E-39 |
| 15.42394913 | 0.958762887 | 0.139815167 | 0.113605208 | 1.10E-113 |
| 1.686773333 | 0.398195876 | 0.148392881 | 0.014885425 | 4.63E-16 |
| 30.38263996 | 0.992268041 | 0.134697006 | 0.152264458 | 6.67E-153 |
| 2.368219175 | 0.393041237 | 0.272875545 | 0.103688158 | 1.00E-103 |
| 1.378560581 | 0.282216495 | 0.394012971 | 0.117204178 | 2.60E-117 |
| 0.959625775 | 0.270618557 | 0.180615784 | 0.025598588 | 1.55E-26 |
| 0.709954676 | 0.162371134 | 0.337191443 | 0.048631307 | 4.24E-49 |
| 3.679676958 | 0.578608247 | 0.075044138 | 0.003269493 | 0.000143789 |
| 1.268908295 | 0.319587629 | 0.328981709 | 0.154057273 | 9.78E-155 |
| 1.556360425 | 0.297680412 | 0.48564952 | 0.149962449 | 1.50E-150 |
| 0.565788435 | 0.163659794 | 0.300780076 | 0.053013952 | 2.08E-53 |
| 4.358089838 | 0.639175258 | 0.157788132 | 0.026442352 | 2.32E-27 |
| 41.92893051 | 0.882731959 | 0.069627656 | 0.003179397 | 0.000177697 |
| 4.077970132 | 0.68685567 | 0.070720723 | 0.008467352 | 9.30E-10 |
| 23.92155477 | 0.858247423 | 0.123892222 | 0.175754312 | 4.90E-177 |

| | | | | |
|---|---|---|---|---|
| 2.243209738 | 0.467783505 | 0.152331519 | 0.012465154 | 1.09E-13 |
| 1.343599868 | 0.492537313 | 0.023379299 | 0.130069373 | 6.67E-120 |
| 0.005234472 | 0.026865672 | 0.956185684 | 0.024378349 | 1.86E-23 |
| 0.674524737 | 0.520895522 | 0.022449098 | 0.12884445 | 9.16E-119 |
| 0.048983853 | 0.059701493 | 0.964543296 | 0.052719651 | 5.14E-49 |
| 0.043307091 | 0.07761194 | 0.102950453 | 0.002234093 | 0.00256836 |
| 0.240703416 | 0.343283582 | 0.090435515 | 0.008517905 | 3.84E-09 |
| 0.614985449 | 0.259701493 | 0.039394296 | 0.055404649 | 1.91E-51 |
| 362.3285574 | 0.974626866 | 0.787449499 | 0.838826591 | 0 |
| 0.007035734 | 0.019402985 | 0.483147528 | 0.005730692 | 1.36E-06 |
| 0.101811949 | 0.237313433 | 0.159785437 | 0.032147594 | 1.89E-30 |
| 1.080458171 | 0.555223881 | 0.01723588 | 0.349294919 | 0 |
| 0.051398946 | 0.119402985 | 0.081048965 | 0.010479678 | 6.35E-11 |
| 0.286763367 | 0.36119403 | 0.153850014 | 0.069411063 | 3.73E-64 |
| 0.09214893 | 0.213432836 | 0.046602672 | 3.72E-05 | 0.697465883 |
| 0.04026027 | 0.097014925 | 0.099929504 | 0.003081402 | 0.000398084 |
| 0.119111365 | 0.197014925 | 0.272741133 | 0.066212001 | 3.01E-61 |
| 0.085688604 | 0.210447761 | 0.294458015 | 0.059393394 | 4.65E-55 |
| 0.347416277 | 0.389552239 | 0.389758653 | 0.199344567 | 4.11E-185 |
| 0.083088471 | 0.207462687 | 0.136928645 | 0.029804019 | 2.43E-28 |
| 0.113062767 | 0.191044776 | 0.052695289 | 0.024301367 | 2.18E-23 |
| 0.06312497 | 0.156716418 | 0.082919682 | 0.006491215 | 2.72E-07 |
| 0.002975957 | 0.017910448 | 0.666414646 | 0.009325448 | 7.08E-10 |
| 0.252367334 | 0.334328358 | 0.029412285 | 0.03765258 | 2.07E-35 |
| 0.0583665 | 0.07761194 | 0.913342491 | 0.065683509 | 9.10E-61 |
| 0.658556839 | 0.562686567 | 0.805323534 | 0.465470809 | 0 |
| 0.265698712 | 0.21641791 | 0.035954976 | 0.033235643 | 1.98E-31 |
| 2.054641052 | 0.734328358 | 0.033745773 | 0.161067617 | 7.19E-149 |
| 6.745527202 | 0.956716418 | 0.046399148 | 0.190122416 | 2.48E-176 |
| 1.910930142 | 0.701492537 | 0.052526981 | 0.069377814 | 3.99E-64 |
| 0.657565145 | 0.38358209 | 0.04115114 | 0.036359028 | 3.03E-34 |
| 0.953398098 | 0.540298507 | 0.044868016 | 0.028031789 | 9.57E-27 |
| 1.401351659 | 0.643283582 | 0.042843691 | 0.042550118 | 7.93E-40 |
| 4.455479987 | 0.902985075 | 0.05286617 | 0.098884515 | 4.30E-91 |
| 0.412054179 | 0.256716418 | 0.049191418 | 0.020020789 | 1.55E-19 |
| 8.226513797 | 0.97761194 | 0.048268857 | 0.154216523 | 1.94E-142 |
| 190.0457239 | 0.96119403 | 0.79710578 | 0.836151995 | 0 |
| 211.2911271 | 0.968656716 | 0.797975354 | 0.840870677 | 0 |
| 1.379487329 | 0.740298507 | 0.02410957 | 0.099386858 | 1.49E-91 |
| 42.31930417 | 0.886567164 | 0.800334162 | 0.710319177 | 0 |
| 0.062939319 | 0.114925373 | 0.534972811 | 0.054826733 | 6.37E-51 |
| 2.082046468 | 0.356716418 | 0.021414478 | 0.237458347 | 9.68E-222 |
| 0.004808354 | 0.019402985 | 0.901967559 | 0.016111538 | 5.17E-16 |
| 1.253893994 | 0.465671642 | 0.031596609 | 0.072646038 | 4.24E-67 |
| 0.185140319 | 0.3 | 0.095762658 | 0.008850716 | 1.91E-09 |

| | | | | |
|---|---|---|---|---|
| 0.072172886 | 0.16119403 | 0.055298671 | 0.00241054 | 0.001736049 |
| 0.286269417 | 0.368656716 | 0.541435787 | 0.250225337 | 3.92E-234 |
| 0.014971665 | 0.040298507 | 0.647813382 | 0.027403407 | 3.52E-26 |
| 1.073341745 | 0.598507463 | 0.024019031 | 0.153443604 | 1.03E-141 |
| 3.59548791 | 0.013432836 | 0.730704802 | 0.006360609 | 3.59E-07 |
| 0.536521164 | 0.332835821 | 0.049714072 | 0.030440905 | 6.49E-29 |
| 1.874331605 | 0.708333333 | 0.595236419 | 0.458966967 | 2.98E-220 |
| 0.613059566 | 0.391666667 | 0.598779339 | 0.199951783 | 1.13E-94 |
| 1.321297628 | 0.595833333 | 0.263593181 | 0.161591828 | 1.34E-76 |
| 0.330211201 | 0.3125 | 0.47026584 | 0.185706391 | 6.02E-88 |
| 0.315589621 | 0.341666667 | 0.461666923 | 0.140381525 | 1.18E-66 |
| 0.734426651 | 0.4875 | 0.382564171 | 0.130502069 | 4.89E-62 |
| 0.711188192 | 0.479166667 | 0.633885167 | 0.263475165 | 6.85E-125 |
| 4.560934703 | 0.7625 | 0.27413701 | 0.200688917 | 5.05E-95 |
| 2.186913546 | 0.675 | 0.230930909 | 0.107997661 | 1.52E-51 |
| 1.338786853 | 0.5875 | 0.409009894 | 0.201541233 | 2.00E-95 |
| 0.663018684 | 0.504166667 | 0.306453248 | 0.119631005 | 5.79E-57 |
| 0.429354209 | 0.4 | 0.498056318 | 0.201954793 | 1.27E-95 |
| 0.946506415 | 0.516666667 | 0.303886835 | 0.256134332 | 2.22E-121 |
| 0.383242752 | 0.391666667 | 0.526221701 | 0.258727509 | 1.28E-122 |
| 0.724856327 | 0.495833333 | 0.375057782 | 0.154565565 | 2.66E-73 |
| 0.940669156 | 0.579166667 | 0.390678755 | 0.342114776 | 8.26E-163 |
| 0.767399877 | 0.475 | 0.488924333 | 0.188909027 | 1.85E-89 |
| 0.870043764 | 0.570833333 | 0.399615051 | 0.233639233 | 1.18E-110 |
| 0.55965574 | 0.433333333 | 0.362543172 | 0.133063726 | 3.11E-63 |
| 1.381815834 | 0.658333333 | 0.245521075 | 0.185365762 | 8.71E-88 |
| 3.181855134 | 0.766666667 | 0.430690379 | 0.349904439 | 1.35E-166 |
| 1.097511932 | 0.458333333 | 0.733075028 | 0.283301241 | 2.14E-134 |
| 0.385178084 | 0.420833333 | 0.367478518 | 0.14591726 | 3.01E-69 |
| 0.526294833 | 0.475 | 0.419308067 | 0.178908418 | 9.63E-85 |
| 0.300215968 | 0.2875 | 0.535483865 | 0.155345723 | 1.15E-73 |
| 7.353545487 | 0.775 | 0.560809664 | 0.4701244 | 8.27E-226 |
| 0.580595214 | 0.441666667 | 0.6125798 | 0.235621093 | 1.35E-111 |
| 1.810015503 | 0.575 | 0.638570769 | 0.374872927 | 8.88E-179 |
| 0.930840447 | 0.5625 | 0.346890362 | 0.27691387 | 2.49E-131 |
| 0.404126651 | 0.329166667 | 0.573196934 | 0.178540887 | 1.43E-84 |
| 1.526138425 | 0.654166667 | 0.535846594 | 0.386412049 | 1.99E-184 |
| 0.427739393 | 0.408333333 | 0.355524337 | 0.115939033 | 3.05E-55 |
| 0.538557053 | 0.441666667 | 0.357186039 | 0.192217715 | 5.09E-91 |
| 0.304957488 | 0.329166667 | 0.476758497 | 0.114120077 | 2.14E-54 |
| 0.248347039 | 0.241666667 | 0.655012029 | 0.115268044 | 6.26E-55 |
| 0.808627453 | 0.5 | 0.367131851 | 0.226606122 | 2.61E-107 |
| 0.412850737 | 0.383333333 | 0.440472291 | 0.094303339 | 3.55E-45 |
| 0.547230612 | 0.404166667 | 0.690268602 | 0.239124297 | 2.89E-113 |
| 1.5218339 | 0.65 | 0.307182817 | 0.262700898 | 1.61E-124 |

| | | | | |
|---|---|---|---|---|
| 0.612704884 | 0.416666667 | 0.45103413 | 0.124645804 | 2.65E-59 |
| 0.759789415 | 0.525 | 0.460443533 | 0.187561847 | 8.02E-89 |
| 2.478288286 | 0.704166667 | 0.656482175 | 0.432900673 | 2.54E-207 |
| 3.043180622 | 0.7125 | 0.420396527 | 0.244519229 | 7.77E-116 |
| 9.592761557 | 0.816666667 | 0.222156114 | 0.112724572 | 9.58E-54 |
| 1.224150651 | 0.570833333 | 0.320086505 | 0.155948472 | 5.98E-74 |
| 0.128020091 | 0.2125 | 0.691819249 | 0.135512821 | 2.23E-64 |
| 0.184322863 | 0.2 | 0.755226522 | 0.116307901 | 2.05E-55 |
| 0.62831889 | 0.5 | 0.322965425 | 0.138881304 | 5.92E-66 |
| 0.987418787 | 0.5125 | 0.419225055 | 0.29272977 | 6.26E-139 |
| 1.589179565 | 0.675 | 0.666615375 | 0.415019403 | 1.72E-198 |
| 0.408620278 | 0.319018405 | 0.685800198 | 0.245890776 | 8.17E-88 |
| 2.552712515 | 0.865030675 | 0.778167467 | 0.771378631 | 4.51E-282 |
| 4.110341958 | 0.938650307 | 0.137793825 | 0.16299593 | 1.70E-58 |
| 5.758336042 | 0.969325153 | 0.137853419 | 0.203792453 | 6.94E-73 |
| 0.159588809 | 0.165644172 | 0.815813905 | 0.138512568 | 6.80E-50 |
| 0.726854599 | 0.570552147 | 0.240865092 | 0.092095746 | 1.19E-33 |
| 7.076762606 | 0.969325153 | 0.15271494 | 0.253562162 | 1.52E-90 |
| 3.46176473 | 0.944785276 | 0.137878191 | 0.165702328 | 1.90E-59 |
| 0.287351955 | 0.294478528 | 0.437261292 | 0.15688701 | 2.40E-56 |
| 2.704342349 | 0.883435583 | 0.143931602 | 0.112939485 | 6.18E-41 |
| 1.489398175 | 0.680981595 | 0.497870186 | 0.428852718 | 9.91E-154 |
| 4.267520409 | 0.938650307 | 0.146283051 | 0.162190801 | 3.27E-58 |
| 1.608795314 | 0.766871166 | 0.197984452 | 0.244421654 | 2.72E-87 |
| 0.256854437 | 0.245398773 | 0.760115637 | 0.200702817 | 8.59E-72 |
| 4.038199507 | 0.950920245 | 0.141050463 | 0.16397404 | 7.71E-59 |
| 0.912273372 | 0.656441718 | 0.346411467 | 0.340788635 | 8.68E-122 |
| 3.259602482 | 0.932515337 | 0.144622512 | 0.178385671 | 6.48E-64 |
| 2.79680065 | 0.877300613 | 0.144461318 | 0.162757962 | 2.07E-58 |
| 0.23680939 | 0.239263804 | 0.663658549 | 0.190747436 | 2.82E-68 |
| 8.339886222 | 0.981595092 | 0.134628961 | 0.148679495 | 1.84E-53 |
| 3.012413744 | 0.877300613 | 0.18124285 | 0.283443698 | 3.33E-101 |
| 0.780251453 | 0.484662577 | 0.830249308 | 0.392588686 | 1.58E-140 |
| 2.155997481 | 0.72392638 | 0.872239219 | 0.616299502 | 3.95E-223 |
| 1.099596469 | 0.63803681 | 0.561463806 | 0.442445861 | 1.07E-158 |
| 4.735969011 | 0.969325153 | 0.157355313 | 0.19007573 | 4.88E-68 |
| 1.555218846 | 0.791411043 | 0.607768731 | 0.491429381 | 1.11E-176 |
| 0.321135929 | 0.282208589 | 0.685860329 | 0.223911096 | 5.22E-80 |
| 31.91637598 | 1 | 0.148190422 | 0.113291634 | 4.66E-41 |
| 8.312131569 | 0.969325153 | 0.161261906 | 0.261678768 | 1.96E-93 |
| 4.156956719 | 0.969325153 | 0.141197952 | 0.151130359 | 2.53E-54 |
| 4.044299567 | 0.938650307 | 0.134552888 | 0.125124259 | 3.35E-45 |
| 5.912011437 | 0.865030675 | 0.184654205 | 0.292738175 | 1.58E-104 |
| 2.052655804 | 0.797546012 | 0.26667256 | 0.399774178 | 3.88E-143 |
| 2.926108391 | 0.883435583 | 0.148004368 | 0.160338651 | 1.47E-57 |

| | | | | |
|---|---|---|---|---|
| 6.869685842 | 0.987730061 | 0.142471815 | 0.206178465 | 9.95E-74 |
| 5.00376718 | 0.975460123 | 0.136333813 | 0.178849647 | 4.45E-64 |
| 5.617532902 | 0.957055215 | 0.155129524 | 0.222089822 | 2.31E-79 |
| 0.469433112 | 0.411042945 | 0.655324227 | 0.309625622 | 1.40E-110 |
| 2.82230843 | 0.889570552 | 0.15257326 | 0.137019241 | 2.27E-49 |
| 0.362794808 | 0.27607362 | 0.628838672 | 0.220295821 | 9.98E-79 |
| 2.545669211 | 0.766871166 | 0.464893867 | 0.344489889 | 4.02E-123 |
| 1.519445471 | 0.582822086 | 0.790726359 | 0.450168495 | 1.59E-161 |
| 4.244799113 | 0.932515337 | 0.146683657 | 0.172934628 | 5.40E-62 |
| 10.51279309 | 0.981595092 | 0.164013939 | 0.322016967 | 4.96E-115 |
| 1.390256623 | 0.705521472 | 0.183194569 | 0.136323685 | 3.98E-49 |
| 5.536453725 | 0.963190184 | 0.148540778 | 0.181756108 | 4.20E-65 |
| 5.586780851 | 0.963190184 | 0.136767131 | 0.148639046 | 1.90E-53 |
| 0.168043734 | 0.17791411 | 0.717416591 | 0.14827831 | 2.54E-53 |
| 2.047725042 | 0.877300613 | 0.166321254 | 0.167662512 | 3.88E-60 |
| 0.441206916 | 0.306748466 | 0.860395605 | 0.232018655 | 6.93E-83 |

| marker_test_q_value |
| ---: |
| 2.38E-34 |
| 1 |
| 1.40E-45 |
| 6.68E-74 |
| 2.15E-11 |
| 0.000321514 |
| 1.48E-277 |
| 3.40E-19 |
| 1.71E-286 |
| 5.14E-161 |
| 2.11226267873055e-312 |
| 0 |
| 6.89E-20 |
| 3.18E-18 |
| 9.45E-263 |
| 0 |
| 0 |
| 1.66E-109 |
| 0 |
| 3.47E-274 |
| 0 |
| 0 |
| 4.72E-32 |
| 0.011150931 |
| 2.52E-67 |
| 1.21E-14 |
| 1.24E-14 |
| 0 |
| 2.13E-08 |
| 3.26E-68 |
| 0 |
| 1.10E-14 |
| 1.01E-09 |
| 3.30E-38 |
| 2.14E-21 |
| 6.62E-198 |
| 1.87E-33 |
| 4.57E-29 |
| 5.85E-96 |
| 2.54E-156 |
| 2.77E-232 |
| 1.32E-15 |
| 0 |
| 0 |

| |
|---|
| 7.99E-128 |
| 8.99E-43 |
| 0 |
| 1.07E-22 |
| 2.07E-20 |
| 0.295495973 |
| 5.99E-39 |
| 3.22E-42 |
| 5.65E-38 |
| 3.26E-62 |
| 6.96E-77 |
| 1.36E-29 |
| 6.36E-65 |
| 2.57E-108 |
| 2.92E-25 |
| 5.24E-42 |
| 2.84E-64 |
| 7.88E-29 |
| 1.04E-32 |
| 5.76E-23 |
| 3.25E-120 |
| 8.46E-35 |
| 7.07E-41 |
| 1.78E-38 |
| 3.27E-29 |
| 1.11E-68 |
| 3.64E-132 |
| 3.63E-42 |
| 2.47E-30 |
| 1.41E-208 |
| 2.94E-78 |
| 4.34E-37 |
| 9.02E-79 |
| 2.41E-110 |
| 3.06E-40 |
| 5.22E-31 |
| 2.61E-39 |
| 3.20E-33 |
| 9.68E-48 |
| 9.12E-38 |
| 3.75E-37 |
| 5.19E-35 |
| 1.08E-82 |
| 7.73E-63 |
| 4.93E-69 |

| |
|---|
| 1.46E-56 |
| 9.02E-95 |
| 7.77E-96 |
| 3.72E-28 |
| 8.81E-19 |
| 7.95E-29 |
| 1.27E-44 |
| 4.45E-48 |
| 4.26E-68 |
| 1.82E-60 |
| 1.48E-37 |
| 3.11E-143 |
| 6.67E-91 |
| 1.08E-119 |
| 2.21E-26 |
| 1.23E-140 |
| 4.88E-99 |
| 1.12E-60 |
| 8.78E-29 |
| 5.08E-45 |
| 7.07E-73 |
| 1.47E-16 |
| 8.32E-48 |
| 1.02E-33 |
| 4.26E-34 |
| 4.70E-31 |
| 5.92E-86 |
| 2.45E-113 |
| 3.21E-43 |
| 4.95E-157 |
| 3.80E-53 |
| 1.45E-47 |
| 7.85E-17 |
| 7.52E-40 |
| 1.22E-55 |
| 7.96E-44 |
| 5.77E-47 |
| 2.17E-132 |
| 1.78E-79 |
| 2.28E-39 |
| 1.53E-116 |
| 1.48E-32 |
| 4.84E-47 |
| 9.13E-56 |
| 2.00E-44 |

| |
|---|
| 3.37E-46 |
| 1.74E-22 |
| 5.75E-113 |
| 4.85E-40 |
| 7.71E-94 |
| 2.39E-68 |
| 3.57E-53 |
| 1.85E-85 |
| 6.09E-26 |
| 1.65E-27 |
| 4.45E-65 |
| 8.80E-32 |
| 3.29E-63 |
| 1.95E-23 |
| 2.88E-69 |
| 8.37E-52 |
| 3.59E-26 |
| 1.32E-30 |
| 3.77E-16 |
| 1.40E-58 |
| 1.99E-26 |
| 1.07E-11 |
| 4.53E-29 |
| 9.17E-39 |
| 1.14E-30 |
| 3.53E-23 |
| 1.07E-25 |
| 4.64E-13 |
| 7.48E-27 |
| 2.18E-15 |
| 6.29E-14 |
| 3.85E-16 |
| 1.15E-11 |
| 6.41E-15 |
| 7.40E-20 |
| 1.50E-49 |
| 1.61E-14 |
| 5.68E-28 |
| 1.25E-27 |
| 1.30E-19 |
| 4.16E-20 |
| 1.27E-30 |
| 4.65E-17 |
| 1.22E-15 |
| 3.55E-14 |

| |
|---|
| 9.94E-15 |
| 1.74E-19 |
| 2.28E-16 |
| 8.09E-15 |
| 1.36E-21 |
| 2.63E-13 |
| 1.01E-26 |
| 1.85E-26 |
| 1.32E-12 |
| 7.68E-17 |
| 5.67E-14 |
| 1.52E-44 |
| 5.42E-22 |
| 1.31E-28 |
| 1.57E-16 |
| 1.22E-33 |
| 2.40E-17 |
| 2.89E-81 |
| 8.78E-20 |
| 7.41E-16 |
| 2.43E-29 |
| 7.42E-24 |
| 1.39E-44 |
| 0.016050125 |
| 4.63E-09 |
| 3.81E-10 |
| 7.53E-09 |
| 3.66E-09 |
| 3.01E-15 |
| 1.68E-38 |
| 1.46E-17 |
| 4.73E-08 |
| 0.001349464 |
| 8.00E-35 |
| 1.40E-06 |
| 1.48E-19 |
| 1.11E-18 |
| 1.58E-07 |
| 8.30E-40 |
| 1.21E-10 |
| 6.39E-19 |
| 3.71E-18 |
| 1.06E-37 |
| 9.17E-11 |
| 9.65E-10 |

| |
|---|
| 8.74E-12 |
| 3.69E-12 |
| 0.000233888 |
| 1.58E-06 |
| 6.93E-24 |
| 5.86E-33 |
| 1.14E-11 |
| 4.31E-37 |
| 7.71E-08 |
| 1.52E-24 |
| 1.63E-10 |
| 5.65E-06 |
| 2.34E-26 |
| 6.71E-16 |
| 7.44E-05 |
| 5.55E-20 |
| 9.90E-26 |
| 2.45E-09 |
| 7.66E-11 |
| 8.55E-55 |
| 9.02E-32 |
| 1.50E-17 |
| 7.25E-21 |
| 2.59E-11 |
| 1.88E-08 |
| 1.54E-10 |
| 1.06E-36 |
| 1.60E-10 |
| 1.45E-15 |
| 3.22E-13 |
| 8.77E-20 |
| 1.53E-23 |
| 5.37E-22 |
| 3.45E-14 |
| 2.36E-17 |
| 4.77E-21 |
| 1.74E-25 |
| 4.93E-14 |
| 5.25E-18 |
| 5.15E-18 |
| 1.13E-15 |
| 1.13E-09 |
| 7.45E-12 |
| 4.92E-19 |
| 4.86E-11 |

| |
|---|
| 1.54E-07 |
| 1.05E-12 |
| 1.63E-29 |
| 6.85E-22 |
| 1.31E-09 |
| 1.04E-11 |
| 2.63E-22 |
| 4.87E-28 |
| 7.62E-07 |
| 3.82E-18 |
| 2.28E-11 |
| 5.60E-19 |
| 3.86E-14 |
| 1.17E-26 |
| 1.23E-10 |
| 1.97E-13 |
| 2.04E-12 |
| 4.54E-40 |
| 1.80E-37 |
| 2.62E-23 |
| 2.43E-21 |
| 2.49E-40 |
| 2.78E-20 |
| 1.11E-12 |
| 1.15E-09 |
| 1.52E-14 |
| 1.33E-10 |
| 3.03E-12 |
| 2.08E-10 |
| 3.25E-16 |
| 7.53E-19 |
| 1.85E-18 |
| 1.96E-15 |
| 4.19E-10 |
| 1.64E-08 |
| 3.67E-07 |
| 2.37E-06 |
| 5.42E-20 |
| 9.84E-13 |
| 1.48E-22 |
| 9.84E-19 |
| 2.12E-12 |
| 8.32E-24 |
| 7.07E-16 |
| 1.70E-12 |

| |
|---|
| 2.63E-11 |
| 6.35E-10 |
| 1.22E-18 |
| 8.05E-11 |
| 4.37E-11 |
| 4.83E-12 |
| 8.24E-24 |
| 9.28E-60 |
| 2.54E-18 |
| 1.06E-31 |
| 6.25E-09 |
| 6.48E-12 |
| 7.76E-07 |
| 1.44E-16 |
| 9.47E-11 |
| 1.50E-21 |
| 1.01E-16 |
| 4.19E-13 |
| 2.89E-30 |
| 2.75E-15 |
| 2.11E-13 |
| 8.06E-22 |
| 1.65E-12 |
| 2.02E-11 |
| 3.78E-13 |
| 6.43E-24 |
| 9.55E-13 |
| 1.17E-16 |
| 1.22E-17 |
| 7.92E-36 |
| 2.64E-16 |
| 3.17E-08 |
| 7.73E-24 |
| 1.51E-20 |
| 1.64E-08 |
| 4.32E-11 |
| 1.13E-06 |
| 1.33E-25 |
| 2.59E-06 |
| 9.16E-09 |
| 2.23E-27 |
| 5.70E-20 |
| 9.06E-13 |
| 1.68E-11 |
| 4.45E-35 |

| |
|---|
| 0.001185556 |
| 6.46E-15 |
| 3.03E-08 |
| 2.61E-11 |
| 1.56E-06 |
| 1.33E-12 |
| 1.97E-09 |
| 1.82E-13 |
| 1.97E-09 |
| 5.20E-06 |
| 4.19E-08 |
| 3.82E-10 |
| 1.50E-11 |
| 4.54E-16 |
| 1.26E-07 |
| 2.45E-05 |
| 1.52E-05 |
| 0.000316344 |
| 8.10E-08 |
| 1.99E-07 |
| 2.71E-14 |
| 0.000404846 |
| 1.31E-09 |
| 1.05E-07 |
| 1.57E-17 |
| 1.47E-18 |
| 2.90E-06 |
| 9.50E-17 |
| 6.21E-06 |
| 0.000221685 |
| 2.44E-13 |
| 3.88E-10 |
| 1.03E-09 |
| 5.70E-11 |
| 0.001070927 |
| 5.64E-06 |
| 2.36E-06 |
| 5.35E-08 |
| 7.85E-05 |
| 2.48E-06 |
| 1.29E-10 |
| 2.28E-33 |
| 2.23E-08 |
| 6.37E-05 |
| 6.29E-33 |

| |
|---|
| 3.92E-51 |
| 4.85E-17 |
| 8.77E-12 |
| 9.30E-10 |
| 1.41E-10 |
| 3.50E-07 |
| 0.017559075 |
| 1.98E-39 |
| 2.54E-09 |
| 3.63E-05 |
| 1.26E-25 |
| 0.001603133 |
| 2.35E-28 |
| 3.62E-20 |
| 7.97E-48 |
| 7.27E-11 |
| 1.15E-14 |
| 0.000108605 |
| 3.60E-07 |
| 7.43E-10 |
| 2.27E-09 |
| 8.79E-16 |
| 4.22E-33 |
| 4.17E-08 |
| 6.61E-55 |
| 2.59E-08 |
| 1.26E-08 |
| 1.42E-05 |
| 0.000247273 |
| 3.61E-08 |
| 5.55E-15 |
| 7.71E-08 |
| 2.80E-13 |
| 7.46E-13 |
| 4.64E-08 |
| 8.92E-14 |
| 0.000465087 |
| 6.73E-23 |
| 6.30E-08 |
| 6.42E-16 |
| 1.40E-07 |
| 1.84E-15 |
| 4.05E-06 |
| 0.07905291 |
| 3.15E-07 |

| |
|---|
| 2.47E-17 |
| 0.722176855 |
| 1 |
| 1 |
| 1 |
| 0.097938315 |
| 1 |
| 0.02233366 |
| 1 |
| 0.015952415 |
| 1 |
| 0.816462688 |
| 1 |
| 1 |
| 0.581369027 |
| 0.354520932 |
| 1 |
| 1 |
| 0.011455619 |
| 1 |
| 0.012155064 |
| 0.086172995 |
| 1 |
| 0.303857435 |
| 1 |
| 1 |
| 1 |
| 0.040291633 |
| 0.01730534 |
| 1 |
| 0.951415869 |
| 1 |
| 0.00616561 |
| 0.003166786 |
| 0.007978566 |
| 0.101169686 |
| 0.243739776 |
| 1 |
| 1 |
| 0.229305375 |
| 0.043220645 |
| 1 |
| 1 |
| 0.081587881 |
| 0.067813064 |

| |
|---|
| 1 |
| 0.060761093 |
| 0.6161351 |
| 1 |
| 1 |
| 1 |
| 1.51E-12 |
| 3.96E-13 |
| 3.00E-18 |
| 1.62E-18 |
| 3.03E-27 |
| 9.45E-22 |
| 3.05E-19 |
| 3.57E-30 |
| 1.71E-36 |
| 1.39E-23 |
| 1.04E-38 |
| 9.34E-15 |
| 1.90E-19 |
| 9.59E-17 |
| 2.07E-15 |
| 1.47E-19 |
| 1.25E-13 |
| 8.33E-24 |
| 4.66E-18 |
| 8.78E-11 |
| 2.38E-11 |
| 9.60E-20 |
| 1.22E-21 |
| 1.13E-37 |
| 1.21E-17 |
| 1.07E-16 |
| 1.76E-24 |
| 1.72E-34 |
| 7.86E-42 |
| 3.11E-25 |
| 7.33E-25 |
| 4.54E-20 |
| 1.10E-11 |
| 1.77E-21 |
| 7.20E-29 |
| 1.56E-11 |
| 2.41E-36 |
| 8.90E-41 |
| 1.11E-19 |

| |
|---|
| 2.31E-20 |
| 2.97E-18 |
| 1.93E-20 |
| 6.79E-21 |
| 1.04E-25 |
| 6.13E-20 |
| 4.44E-25 |
| 5.64E-15 |
| 9.76E-16 |
| 1.95E-21 |
| 6.65E-14 |
| 0 |
| 0 |
| 0 |
| 0 |
| 0 |
| 0 |
| 0 |
| 0 |
| 0 |
| 0 |
| 0 |
| 0 |
| 0 |
| 0 |
| 0 |
| 0 |
| 0 |
| 0 |
| 0 |
| 0 |
| 0 |
| 0 |
| 0 |
| 0 |
| 0 |
| 0 |
| 0 |
| 0 |
| 0 |
| 0 |
| 0 |
| 0 |
| 0 |
| 0 |

| |
|---|
| 0 |
| 0 |
| 0 |
| 0 |
| 0 |
| 0 |
| 0 |
| 0 |
| 0 |
| 0 |
| 0 |
| 0 |
| 0 |
| 0 |
| 0 |
| 0 |
| 1 |
| 0.002386434 |
| 0.007167517 |
| 1 |
| 0.009614844 |
| 4.31E-20 |
| 2.05E-14 |
| 4.24E-12 |
| 1.48E-14 |
| 1.14E-05 |
| 2.82E-07 |
| 4.19E-07 |
| 0.001656883 |
| 0.002600016 |
| 0.079250631 |
| 5.45E-13 |
| 1.46E-14 |
| 1 |
| 0.002283975 |
| 0.004993911 |
| 3.99E-08 |
| 3.99E-13 |
| 9.33E-12 |
| 8.83E-05 |
| 1.75E-21 |
| 1 |
| 1 |
| 1 |
| 1 |

| |
|---|
| 0.101795472 |
| 2.85E-08 |
| 0.000115469 |
| 1.92E-24 |
| 0.000310183 |
| 2.62E-05 |
| 0.012787008 |
| 1 |
| 7.05E-06 |
| 4.56E-07 |
| 0.792256352 |
| 0.850268957 |
| 1.69E-05 |
| 4.03E-07 |
| 0.001827636 |
| 0.060880825 |
| 2.33E-44 |
| 0.914475817 |
| 3.03E-05 |
| 1 |
| 1 |
| 1.28E-15 |
| 5.52E-09 |
| 9.82E-22 |
| 7.81E-05 |
| 4.37E-06 |
| 1 |
| 5.19E-34 |
| 1.45E-30 |
| 5.31E-24 |
| 1.02E-13 |
| 8.37E-18 |
| 0.000300444 |
| 1.80E-40 |
| 9.19E-32 |
| 1.27E-09 |
| 0.000309421 |
| 0.000566455 |
| 5.99E-09 |
| 1.90E-08 |
| 7.51E-24 |
| 0.006988585 |
| 4.31E-05 |
| 4.89E-05 |
| 0.046633344 |

| |
|---|
| 0.001217595 |
| 2.97E-12 |
| 4.12E-19 |
| 3.01E-10 |
| 1.67E-08 |
| 1.68E-33 |
| 1.54E-07 |
| 0.000790937 |
| 4.86E-30 |
| 0.13024427 |
| 0.009408209 |
| 1.17E-14 |
| 5.86E-09 |
| 0.000298425 |
| 6.28E-07 |
| 5.79E-12 |
| 7.74E-31 |
| 1.98E-17 |
| 8.48E-32 |
| 2.69E-21 |
| 4.16E-07 |
| 0.065120126 |
| 0.483303861 |
| 1 |
| 4.58E-17 |
| 8.09E-16 |
| 0 |
| 0 |
| 0 |
| 0 |
| 0 |
| 0 |
| 0 |
| 0 |
| 0 |
| 0 |
| 0 |
| 0 |
| 0 |
| 0 |
| 0 |
| 0 |
| 0 |
| 0 |
| 0 |

| |
|---|
| 0 |
| 0 |
| 0 |
| 0 |
| 0 |
| 0 |
| 0 |
| 0 |
| 0 |
| 0 |
| 0 |
| 0 |
| 0 |
| 0 |
| 0 |
| 0 |
| 0 |
| 0 |
| 0 |
| 0 |
| 0 |
| 0 |
| 0 |
| 0 |
| 0 |
| 0 |
| 0 |
| 0 |
| 0 |
| 0 |
| 0 |
| 0 |
| 0 |
| 0 |
| 0 |
| 0 |
| 3.35E-240 |
| 0 |
| 0 |
| 0 |
| 0 |
| 1.80E-271 |
| 0 |
| 0 |
| 0 |

| |
|---|
| 0 |
| 9.07E-205 |
| 2.12846741061672e-317 |
| 0 |
| 0 |
| 0 |
| 0 |
| 0 |
| 2.82E-303 |
| 0 |
| 0 |
| 1.99E-302 |
| 0 |
| 0 |
| 0 |
| 0 |
| 0 |
| 0 |
| 0 |
| 8.50E-280 |
| 0 |
| 0 |
| 0 |
| 0 |
| 0 |
| 6.50E-281 |
| 0 |
| 0 |
| 0 |
| 5.00E-304 |
| 7.92E-288 |
| 0 |
| 0 |
| 0 |
| 0 |
| 7.05E-235 |
| 0 |
| 0 |
| 0 |
| 3.30E-287 |
| 0 |
| 1.89E-295 |
| 1.19E-270 |
| 0 |
| 0 |

| |
|---|
| 0 |
| 1.22E-220 |
| 0 |
| 6.51E-259 |
| 3.33E-287 |
| 1.44E-221 |
| 0 |
| 4.58E-206 |
| 0.00E+00 |
| 0 |
| 0 |
| 0 |
| 0 |
| 4.78E-250 |
| 4.82E-293 |
| 1.77E-212 |
| 1.48E-304 |
| 0 |
| 8.48E-231 |
| 2.49E-263 |
| 1.04E-283 |
| 0 |
| 1.98E-281 |
| 2.91E-274 |
| 0 |
| 0 |
| 1.26E-243 |
| 0 |
| 0 |
| 3.98023405785326e-315 |
| 0 |
| 0 |
| 1.10E-272 |
| 0 |
| 8.12E-241 |
| 2.54E-267 |
| 2.68E-306 |
| 5.69E-296 |
| 0 |
| 0 |
| 2.19E-254 |
| 0.109040498 |
| 8.39E-33 |
| 7.48E-34 |
| 7.00E-104 |

| |
|---|
| 5.47E-158 |
| 1.72E-38 |
| 1.17E-45 |
| 4.19E-67 |
| 0.004983841 |
| 2.23E-39 |
| 6.07E-101 |
| 5.78E-61 |
| 8.35E-36 |
| 7.13E-121 |
| 6.27E-30 |
| 2.81E-143 |
| 2.50E-83 |
| 1.62E-59 |
| 2.00E-71 |
| 3.11E-97 |
| 1.02E-124 |
| 1.61E-37 |
| 1.11E-84 |
| 2.09E-30 |
| 5.32E-54 |
| 6.72E-59 |
| 3.15E-28 |
| 2.93E-74 |
| 1 |
| 1.51E-109 |
| 7.91E-33 |
| 1 |
| 3.00E-22 |
| 3.21E-33 |
| 5.24E-108 |
| 2.21E-10 |
| 3.19E-147 |
| 4.80E-98 |
| 1.24E-111 |
| 7.42E-21 |
| 2.03E-43 |
| 1 |
| 4.68E-149 |
| 7.19E-145 |
| 9.96E-48 |
| 1.11E-21 |
| 1 |
| 0.000445218 |
| 2.35E-171 |

| |
|---|
| 5.21E-08 |
| 3.19E-114 |
| 8.89E-18 |
| 4.38E-113 |
| 2.46E-43 |
| 1 |
| 0.001839221 |
| 9.14E-46 |
| 0 |
| 0.651488951 |
| 9.03E-25 |
| 0 |
| 3.04E-05 |
| 1.78E-58 |
| 1 |
| 1 |
| 1.44E-55 |
| 2.22E-49 |
| 1.97E-179 |
| 1.16E-22 |
| 1.04E-17 |
| 0.130435764 |
| 0.000339056 |
| 9.91E-30 |
| 4.36E-55 |
| 0 |
| 9.46E-26 |
| 3.44E-143 |
| 1.19E-170 |
| 1.91E-58 |
| 1.45E-28 |
| 4.58E-21 |
| 3.80E-34 |
| 2.06E-85 |
| 7.44E-14 |
| 9.29E-137 |
| 0 |
| 0 |
| 7.11E-86 |
| 0 |
| 3.05E-45 |
| 4.63E-216 |
| 2.48E-10 |
| 2.03E-61 |
| 0.000915823 |

| |
|---|
| 1 |
| 1.88E-228 |
| 1.68E-20 |
| 4.92E-136 |
| 0.171855121 |
| 3.11E-23 |
| 1.42E-214 |
| 5.39E-89 |
| 6.43E-71 |
| 2.88E-82 |
| 5.63E-61 |
| 2.34E-56 |
| 3.28E-119 |
| 2.42E-89 |
| 7.28E-46 |
| 9.55E-90 |
| 2.77E-51 |
| 6.09E-90 |
| 1.06E-115 |
| 6.11E-117 |
| 1.28E-67 |
| 3.95E-157 |
| 8.88E-84 |
| 5.66E-105 |
| 1.49E-57 |
| 4.17E-82 |
| 6.44E-161 |
| 1.02E-128 |
| 1.44E-63 |
| 4.61E-79 |
| 5.49E-68 |
| 3.96E-220 |
| 6.45E-106 |
| 4.25E-173 |
| 1.19E-125 |
| 6.86E-79 |
| 9.52E-179 |
| 1.46E-49 |
| 2.43E-85 |
| 1.03E-48 |
| 2.99E-49 |
| 1.25E-101 |
| 1.70E-39 |
| 1.38E-107 |
| 7.69E-119 |

| |
|---|
| 1.27E-53 |
| 3.84E-83 |
| 1.22E-201 |
| 3.72E-110 |
| 4.58E-48 |
| 2.86E-68 |
| 1.07E-58 |
| 9.81E-50 |
| 2.83E-60 |
| 3.00E-133 |
| 8.24E-193 |
| 3.91E-82 |
| 2.16E-276 |
| 8.15E-53 |
| 3.32E-67 |
| 3.25E-44 |
| 5.72E-28 |
| 7.29E-85 |
| 9.10E-54 |
| 1.15E-50 |
| 2.96E-35 |
| 4.74E-148 |
| 1.57E-52 |
| 1.30E-81 |
| 4.11E-66 |
| 3.69E-53 |
| 4.16E-116 |
| 3.10E-58 |
| 9.89E-53 |
| 1.35E-62 |
| 8.78E-48 |
| 1.59E-95 |
| 7.57E-135 |
| 1.89E-217 |
| 5.11E-153 |
| 2.33E-62 |
| 5.31E-171 |
| 2.50E-74 |
| 2.23E-35 |
| 9.36E-88 |
| 1.21E-48 |
| 1.61E-39 |
| 7.54E-99 |
| 1.86E-137 |
| 7.01E-52 |

| |
|---|
| 4.76E-68 |
| 2.13E-58 |
| 1.10E-73 |
| 6.69E-105 |
| 1.09E-43 |
| 4.78E-73 |
| 1.92E-117 |
| 7.62E-156 |
| 2.59E-56 |
| 2.37E-109 |
| 1.91E-43 |
| 2.01E-59 |
| 9.08E-48 |
| 1.22E-47 |
| 1.86E-54 |
| 3.32E-77 |

| | cell_type | cell_group | gene_id | gene_short_name | marker_score |
|---|---|---|---|---|---|
| 1 | SMC | 1 | ENSMUSG00000001270 | Ckb | 0.14606208 |
| 2 | SMC | 1 | ENSMUSG00000001349 | Cnn1 | 0.153536771 |
| 3 | SMC | 1 | ENSMUSG00000004951 | Hspb1 | 0.08393632 |
| 4 | SMC | 1 | ENSMUSG00000007892 | Rplp1 | 0.114572584 |
| 5 | SMC | 1 | ENSMUSG00000020439 | Smtn | 0.108457127 |
| 6 | SMC | 1 | ENSMUSG00000015143 | Actn1 | 0.07980899 |
| 7 | SMC | 1 | ENSMUSG00000000555 | Itga5 | 0.080407726 |
| 8 | SMC | 1 | ENSMUSG00000022836 | Mylk | 0.103104152 |
| 9 | SMC | 1 | ENSMUSG00000024661 | Fth1 | 0.085296498 |
| 10 | SMC | 1 | ENSMUSG00000025393 | Atp5b | 0.077896596 |
| 11 | SMC | 1 | ENSMUSG00000026208 | Des | 0.178257681 |
| 12 | SMC | 1 | ENSMUSG00000026414 | Tnnt2 | 0.077375233 |
| 13 | SMC | 1 | ENSMUSG00000026421 | Csrp1 | 0.145727595 |
| 14 | SMC | 1 | ENSMUSG00000028464 | Tpm2 | 0.160938052 |
| 15 | SMC | 1 | ENSMUSG00000032366 | Tpm1 | 0.147916927 |
| 16 | SMC | 1 | ENSMUSG00000029761 | Cald1 | 0.115109225 |
| 17 | SMC | 1 | ENSMUSG00000030409 | Dmpk | 0.129999237 |
| 18 | SMC | 1 | ENSMUSG00000030695 | Aldoa | 0.084921834 |
| 19 | SMC | 1 | ENSMUSG00000031328 | Flna | 0.141815551 |
| 20 | SMC | 1 | ENSMUSG00000031633 | Slc25a4 | 0.077478068 |
| 21 | SMC | 1 | ENSMUSG00000031636 | Pdlim3 | 0.087962845 |
| 22 | SMC | 1 | ENSMUSG00000032085 | Tagln | 0.154284964 |
| 23 | SMC | 1 | ENSMUSG00000035202 | Lars2 | 0.099915011 |
| 24 | SMC | 1 | ENSMUSG00000035783 | Acta2 | 0.146894745 |
| 25 | SMC | 1 | ENSMUSG00000037742 | Eef1a1 | 0.086457734 |
| 26 | SMC | 1 | ENSMUSG00000021493 | Pdlim7 | 0.09881733 |
| 27 | SMC | 1 | ENSMUSG00000050856 | Atp5k | 0.084040678 |
| 28 | SMC | 1 | ENSMUSG00000046330 | Rpl37a | 0.097586945 |
| 29 | SMC | 1 | ENSMUSG00000059430 | Actg2 | 0.153226055 |
| 30 | SMC | 1 | ENSMUSG00000057322 | Rpl38 | 0.088040184 |
| 31 | SMC | 1 | ENSMUSG00000064337 | mt-Rnr1 | 0.114793898 |
| 32 | SMC | 1 | ENSMUSG00000064339 | mt-Rnr2 | 0.107467211 |
| 33 | SMC | 1 | ENSMUSG00000064341 | mt-Nd1 | 0.117679698 |
| 34 | SMC | 1 | ENSMUSG00000064345 | mt-Nd2 | 0.124898882 |
| 35 | SMC | 1 | ENSMUSG00000064356 | mt-Atp8 | 0.091488434 |
| 36 | SMC | 1 | ENSMUSG00000064363 | mt-Nd4 | 0.112435415 |
| 37 | SMC | 1 | ENSMUSG00000064370 | mt-Cytb | 0.115641045 |
| 38 | SMC | 1 | ENSMUSG00000067818 | Myl9 | 0.1529935 |
| 39 | SMC | 1 | ENSMUSG00000068220 | Lgals1 | 0.079175265 |
| 40 | SMC | 1 | ENSMUSG00000018830 | Myh11 | 0.156708784 |
| 41 | SMC | 1 | ENSMUSG00000071356 | Reg3b | 0.085134109 |
| 42 | SMC | 1 | ENSMUSG00000029580 | Actb | 0.113329994 |
| 43 | SMC | 1 | ENSMUSG00000015932 | Dstn | 0.083632239 |
| 44 | SMC | 1 | ENSMUSG00000060126 | Tpt1 | 0.089301956 |

| | | | | | |
|---|---|---|---|---|---|
| 45 | SMC | 1 | ENSMUSG00000025290 | Rps24 | 0.078884664 |
| 46 | SMC | 1 | ENSMUSG00000090841 | Myl6 | 0.14298616 |
| 47 | SMC | 1 | ENSMUSG00000093674 | Rpl41 | 0.089889924 |
| 48 | SMC | 1 | ENSMUSG00000098973 | Mir6236 | 0.11564415 |
| 49 | SMC | 1 | ENSMUSG00000101939 | Gm28438 | 0.085449719 |
| 50 | SMC | 1 | ENSMUSG00000106106 | CT010467.1 | 0.154644599 |
| 51 | Fibroblasts | 10 | ENSMUSG00000000359 | Rem1 | 0.138547604 |
| 52 | Fibroblasts | 10 | ENSMUSG00000029231 | Pdgfra | 0.403483895 |
| 53 | Fibroblasts | 10 | ENSMUSG00000000794 | Kcnn3 | 0.29539374 |
| 54 | Fibroblasts | 10 | ENSMUSG00000002900 | Lamb1 | 0.131497369 |
| 55 | Fibroblasts | 10 | ENSMUSG00000002997 | Prkar2b | 0.147658547 |
| 56 | Fibroblasts | 10 | ENSMUSG00000015134 | Aldh1a3 | 0.142794827 |
| 57 | Fibroblasts | 10 | ENSMUSG00000019966 | Kitl | 0.133447933 |
| 58 | Fibroblasts | 10 | ENSMUSG00000020719 | Ddx5 | 0.151553936 |
| 59 | Fibroblasts | 10 | ENSMUSG00000020866 | Cacna1g | 0.13574336 |
| 60 | Fibroblasts | 10 | ENSMUSG00000022114 | Spry2 | 0.18589787 |
| 61 | Fibroblasts | 10 | ENSMUSG00000022860 | Chodl | 0.207719078 |
| 62 | Fibroblasts | 10 | ENSMUSG00000025969 | Nrp2 | 0.127075005 |
| 63 | Fibroblasts | 10 | ENSMUSG00000027210 | Meis2 | 0.245260706 |
| 64 | Fibroblasts | 10 | ENSMUSG00000028005 | Gucy1b1 | 0.254290586 |
| 65 | Fibroblasts | 10 | ENSMUSG00000028364 | Tnc | 0.148766703 |
| 66 | Fibroblasts | 10 | ENSMUSG00000029465 | Arpc3 | 0.175629174 |
| 67 | Fibroblasts | 10 | ENSMUSG00000029467 | Atp2a2 | 0.137755884 |
| 68 | Fibroblasts | 10 | ENSMUSG00000030218 | Mgp | 0.203210652 |
| 69 | Fibroblasts | 10 | ENSMUSG00000032035 | Ets1 | 0.135502354 |
| 70 | Fibroblasts | 10 | ENSMUSG00000035356 | Nfkbiz | 0.243663872 |
| 71 | Fibroblasts | 10 | ENSMUSG00000036446 | Lum | 0.204137141 |
| 72 | Fibroblasts | 10 | ENSMUSG00000041757 | Plekha6 | 0.197020705 |
| 73 | Fibroblasts | 10 | ENSMUSG00000039252 | Lgi2 | 0.165418373 |
| 74 | Fibroblasts | 10 | ENSMUSG00000042436 | Mfap4 | 0.141139996 |
| 75 | Fibroblasts | 10 | ENSMUSG00000028369 | Svep1 | 0.207496717 |
| 76 | Fibroblasts | 10 | ENSMUSG00000037379 | Spon2 | 0.125492539 |
| 77 | Fibroblasts | 10 | ENSMUSG00000041482 | Piezo2 | 0.141599085 |
| 78 | Fibroblasts | 10 | ENSMUSG00000033910 | Gucy1a1 | 0.30076372 |
| 79 | Fibroblasts | 10 | ENSMUSG00000050212 | Eva1b | 0.1383244 |
| 80 | Fibroblasts | 10 | ENSMUSG00000045875 | Adra1a | 0.137514912 |
| 81 | Fibroblasts | 10 | ENSMUSG00000048126 | Col6a3 | 0.171831303 |
| 82 | Fibroblasts | 10 | ENSMUSG00000041261 | Car8 | 0.205657784 |
| 83 | Fibroblasts | 10 | ENSMUSG00000038855 | Itpkb | 0.147658404 |
| 84 | Fibroblasts | 10 | ENSMUSG00000029673 | Auts2 | 0.12989721 |
| 85 | Fibroblasts | 10 | ENSMUSG00000025936 | Gm4956 | 0.211007169 |
| 86 | Fibroblasts | 10 | ENSMUSG00000070576 | Mn1 | 0.130404227 |
| 87 | Fibroblasts | 10 | ENSMUSG00000021134 | Srsf5 | 0.127658776 |
| 88 | Fibroblasts | 10 | ENSMUSG00000038886 | Man2a2 | 0.141332316 |
| 89 | Fibroblasts | 10 | ENSMUSG00000053062 | Jam2 | 0.177941296 |

| | | | | | |
|---|---|---|---|---|---|
| 90 | Fibroblasts | 10 | ENSMUSG00000074971 | Fibin | 0.217203906 |
| 91 | Fibroblasts | 10 | ENSMUSG00000019929 | Dcn | 0.125925014 |
| 92 | Fibroblasts | 10 | ENSMUSG00000047907 | Tshz2 | 0.271080201 |
| 93 | Fibroblasts | 10 | ENSMUSG00000055632 | Hmcn2 | 0.221179399 |
| 94 | Fibroblasts | 10 | ENSMUSG00000022961 | Son | 0.18119264 |
| 95 | Fibroblasts | 10 | ENSMUSG00000052516 | Robo2 | 0.254135404 |
| 96 | Fibroblasts | 10 | ENSMUSG00000045658 | Pid1 | 0.422136222 |
| 97 | Fibroblasts | 10 | ENSMUSG00000023885 | Thbs2 | 0.189712434 |
| 98 | Fibroblasts | 10 | ENSMUSG00000092341 | Malat1 | 0.162224034 |
| 99 | Fibroblasts | 10 | ENSMUSG00000097324 | Carmn | 0.126668063 |
| 100 | Fibroblasts | 10 | ENSMUSG00000097971 | Gm26917 | 0.192157109 |
| 101 | SMC | 11 | ENSMUSG00000001270 | Ckb | 0.19982487 |
| 102 | SMC | 11 | ENSMUSG00000001349 | Cnn1 | 0.224307365 |
| 103 | SMC | 11 | ENSMUSG00000002379 | Ndufa11 | 0.135174673 |
| 104 | SMC | 11 | ENSMUSG00000004951 | Hspb1 | 0.150987106 |
| 105 | SMC | 11 | ENSMUSG00000009013 | Dynll1 | 0.143269237 |
| 106 | SMC | 11 | ENSMUSG00000016252 | Atp5e | 0.132231142 |
| 107 | SMC | 11 | ENSMUSG00000020163 | Uqcr11 | 0.134512517 |
| 108 | SMC | 11 | ENSMUSG00000020439 | Smtn | 0.13297887 |
| 109 | SMC | 11 | ENSMUSG00000022354 | Ndufb9 | 0.122613826 |
| 110 | SMC | 11 | ENSMUSG00000022836 | Mylk | 0.131553721 |
| 111 | SMC | 11 | ENSMUSG00000024608 | Rps14 | 0.126784692 |
| 112 | SMC | 11 | ENSMUSG00000024661 | Fth1 | 0.132363379 |
| 113 | SMC | 11 | ENSMUSG00000025393 | Atp5b | 0.128390179 |
| 114 | SMC | 11 | ENSMUSG00000026208 | Des | 0.177687598 |
| 115 | SMC | 11 | ENSMUSG00000026414 | Tnnt2 | 0.178293975 |
| 116 | SMC | 11 | ENSMUSG00000026421 | Csrp1 | 0.187417142 |
| 117 | SMC | 11 | ENSMUSG00000026678 | Rgs5 | 0.16638294 |
| 118 | SMC | 11 | ENSMUSG00000028464 | Tpm2 | 0.248809377 |
| 119 | SMC | 11 | ENSMUSG00000032366 | Tpm1 | 0.238990372 |
| 120 | SMC | 11 | ENSMUSG00000029632 | Ndufa4 | 0.126656116 |
| 121 | SMC | 11 | ENSMUSG00000029761 | Cald1 | 0.223141265 |
| 122 | SMC | 11 | ENSMUSG00000031328 | Flna | 0.197557317 |
| 123 | SMC | 11 | ENSMUSG00000031633 | Slc25a4 | 0.145593757 |
| 124 | SMC | 11 | ENSMUSG00000058056 | Palld | 0.156171916 |
| 125 | SMC | 11 | ENSMUSG00000032330 | Cox7a2 | 0.131848291 |
| 126 | SMC | 11 | ENSMUSG00000038690 | Atp5j2 | 0.120537055 |
| 127 | SMC | 11 | ENSMUSG00000033306 | Lpp | 0.133134286 |
| 128 | SMC | 11 | ENSMUSG00000035783 | Acta2 | 0.231643138 |
| 129 | SMC | 11 | ENSMUSG00000035885 | Cox8a | 0.180290737 |
| 130 | SMC | 11 | ENSMUSG00000036438 | Calm2 | 0.132778026 |
| 131 | SMC | 11 | ENSMUSG00000038387 | Rras | 0.123097547 |
| 132 | SMC | 11 | ENSMUSG00000037166 | Ppp1r14a | 0.20939098 |
| 133 | SMC | 11 | ENSMUSG00000040666 | Sh3bgr | 0.159743447 |
| 134 | SMC | 11 | ENSMUSG00000050856 | Atp5k | 0.153219447 |

| | | | | | |
|---|---|---|---|---|---|
| 135 | SMC | 11 | ENSMUSG00000063882 | Uqcrh | 0.130497297 |
| 136 | SMC | 11 | ENSMUSG00000050503 | Fbxl22 | 0.125878242 |
| 137 | SMC | 11 | ENSMUSG00000059534 | Uqcr10 | 0.116273724 |
| 138 | SMC | 11 | ENSMUSG00000049422 | Chchd10 | 0.117592603 |
| 139 | SMC | 11 | ENSMUSG00000039001 | Rps21 | 0.119700457 |
| 140 | SMC | 11 | ENSMUSG00000048096 | Lmod1 | 0.172882781 |
| 141 | SMC | 11 | ENSMUSG00000090223 | Pcp4 | 0.160611714 |
| 142 | SMC | 11 | ENSMUSG00000019907 | Ppp1r12a | 0.168882722 |
| 143 | SMC | 11 | ENSMUSG00000059430 | Actg2 | 0.198139906 |
| 144 | SMC | 11 | ENSMUSG00000067818 | Myl9 | 0.247456996 |
| 145 | SMC | 11 | ENSMUSG00000018830 | Myh11 | 0.232074842 |
| 146 | SMC | 11 | ENSMUSG00000075702 | Selenom | 0.143170688 |
| 147 | SMC | 11 | ENSMUSG00000029580 | Actb | 0.15933606 |
| 148 | SMC | 11 | ENSMUSG00000015932 | Dstn | 0.145730887 |
| 149 | SMC | 11 | ENSMUSG00000090841 | Myl6 | 0.191652341 |
| 150 | SMC | 11 | ENSMUSG00000097324 | Carmn | 0.13149949 |
| 151 | RBC | 12 | ENSMUSG00000022283 | Pabpc1 | 0.061881862 |
| 152 | RBC | 12 | ENSMUSG00000005161 | Prdx2 | 0.045217929 |
| 153 | RBC | 12 | ENSMUSG00000006574 | Slc4a1 | 0.114497169 |
| 154 | RBC | 12 | ENSMUSG00000007659 | Bcl2l1 | 0.057009613 |
| 155 | RBC | 12 | ENSMUSG00000018677 | Slc25a39 | 0.068314493 |
| 156 | RBC | 12 | ENSMUSG00000019505 | Ubb | 0.053325895 |
| 157 | RBC | 12 | ENSMUSG00000020641 | Rsad2 | 0.080882952 |
| 158 | RBC | 12 | ENSMUSG00000021792 | Prxl2a | 0.056061355 |
| 159 | RBC | 12 | ENSMUSG00000022051 | Bnip3l | 0.204192042 |
| 160 | RBC | 12 | ENSMUSG00000052305 | Hbb-bs | 0.674474679 |
| 161 | RBC | 12 | ENSMUSG00000024158 | Hagh | 0.061621178 |
| 162 | RBC | 12 | ENSMUSG00000024588 | Fech | 0.243922464 |
| 163 | RBC | 12 | ENSMUSG00000026208 | Des | 0.051595384 |
| 164 | RBC | 12 | ENSMUSG00000028436 | Dcaf12 | 0.043433972 |
| 165 | RBC | 12 | ENSMUSG00000028464 | Tpm2 | 0.04961238 |
| 166 | RBC | 12 | ENSMUSG00000032366 | Tpm1 | 0.043355706 |
| 167 | RBC | 12 | ENSMUSG00000028906 | Epb41 | 0.18754585 |
| 168 | RBC | 12 | ENSMUSG00000029922 | Mkrn1 | 0.324583244 |
| 169 | RBC | 12 | ENSMUSG00000030878 | Cdr2 | 0.05381744 |
| 170 | RBC | 12 | ENSMUSG00000031812 | Map1lc3b | 0.050239975 |
| 171 | RBC | 12 | ENSMUSG00000031950 | Gabarapl2 | 0.104177413 |
| 172 | RBC | 12 | ENSMUSG00000034248 | Slc25a37 | 0.309095985 |
| 173 | RBC | 12 | ENSMUSG00000039236 | Isg20 | 0.184561486 |
| 174 | RBC | 12 | ENSMUSG00000035783 | Acta2 | 0.044087488 |
| 175 | RBC | 12 | ENSMUSG00000038871 | Bpgm | 0.376450672 |
| 176 | RBC | 12 | ENSMUSG00000044792 | Isca1 | 0.14993053 |
| 177 | RBC | 12 | ENSMUSG00000047139 | Cd24a | 0.051621943 |
| 178 | RBC | 12 | ENSMUSG00000023572 | Ccndbp1 | 0.098886303 |
| 179 | RBC | 12 | ENSMUSG00000073400 | Trim10 | 0.048618194 |

| | | | | | | |
|---|---|---|---|---|---|---|
| 180 | RBC | 12 | ENSMUSG00000035242 | Oaz1 | | 0.046437097 |
| 181 | RBC | 12 | ENSMUSG00000044468 | Tent5c | | 0.37953239 |
| 182 | RBC | 12 | ENSMUSG00000051839 | Gypa | | 0.095743496 |
| 183 | RBC | 12 | ENSMUSG00000079557 | | 2-Mar | 0.124952393 |
| 184 | RBC | 12 | ENSMUSG00000025270 | Alas2 | | 0.600955307 |
| 185 | RBC | 12 | ENSMUSG00000059430 | Actg2 | | 0.045342852 |
| 186 | RBC | 12 | ENSMUSG00000020802 | Ube2o | | 0.087376349 |
| 187 | RBC | 12 | ENSMUSG00000018830 | Myh11 | | 0.044280481 |
| 188 | RBC | 12 | ENSMUSG00000069917 | Hba-a2 | | 0.678135084 |
| 189 | RBC | 12 | ENSMUSG00000069919 | Hba-a1 | | 0.677549921 |
| 190 | RBC | 12 | ENSMUSG00000050708 | Ftl1 | | 0.097280507 |
| 191 | RBC | 12 | ENSMUSG00000073940 | Hbb-bt | | 0.656987743 |
| 192 | RBC | 12 | ENSMUSG00000074269 | Rec114 | | 0.133433722 |
| 193 | RBC | 12 | ENSMUSG00000027078 | Ube2l6 | | 0.341947132 |
| 194 | RBC | 12 | ENSMUSG00000078974 | Sec61g | | 0.048243929 |
| 195 | RBC | 12 | ENSMUSG00000060126 | Tpt1 | | 0.050878839 |
| 196 | RBC | 12 | ENSMUSG00000056234 | Ncoa4 | | 0.047757476 |
| 197 | RBC | 12 | ENSMUSG00000025889 | Snca | | 0.418236419 |
| 198 | RBC | 12 | ENSMUSG00000033685 | Ucp2 | | 0.060671697 |
| 199 | RBC | 12 | ENSMUSG00000091694 | Apol11b | | 0.053004804 |
| 200 | RBC | 12 | ENSMUSG00000106106 | CT010467.1 | | 0.069884974 |
| 201 | Endothelial | 14 | ENSMUSG00000001029 | Icam2 | | 0.297878601 |
| 202 | Endothelial | 14 | ENSMUSG00000001946 | Esam | | 0.397173175 |
| 203 | Endothelial | 14 | ENSMUSG00000022220 | Adcy4 | | 0.333093914 |
| 204 | Endothelial | 14 | ENSMUSG00000004655 | Aqp1 | | 0.390362979 |
| 205 | Endothelial | 14 | ENSMUSG00000026814 | Eng | | 0.296539367 |
| 206 | Endothelial | 14 | ENSMUSG00000030123 | Plxnd1 | | 0.277241292 |
| 207 | Endothelial | 14 | ENSMUSG00000015468 | Notch4 | | 0.283156408 |
| 208 | Endothelial | 14 | ENSMUSG00000017309 | Cd300lg | | 0.30819892 |
| 209 | Endothelial | 14 | ENSMUSG00000020427 | Igfbp3 | | 0.297516547 |
| 210 | Endothelial | 14 | ENSMUSG00000020846 | Rflnb | | 0.289286817 |
| 211 | Endothelial | 14 | ENSMUSG00000021367 | Edn1 | | 0.286465593 |
| 212 | Endothelial | 14 | ENSMUSG00000044258 | Ctla2a | | 0.405209972 |
| 213 | Endothelial | 14 | ENSMUSG00000022579 | Gpihbp1 | | 0.536559844 |
| 214 | Endothelial | 14 | ENSMUSG00000024140 | Epas1 | | 0.388613427 |
| 215 | Endothelial | 14 | ENSMUSG00000025608 | Podxl | | 0.291842218 |
| 216 | Endothelial | 14 | ENSMUSG00000025902 | Sox17 | | 0.31695711 |
| 217 | Endothelial | 14 | ENSMUSG00000062515 | Fabp4 | | 0.574836302 |
| 218 | Endothelial | 14 | ENSMUSG00000029648 | Flt1 | | 0.620936132 |
| 219 | Endothelial | 14 | ENSMUSG00000031871 | Cdh5 | | 0.589857422 |
| 220 | Endothelial | 14 | ENSMUSG00000024451 | Arap3 | | 0.390474066 |
| 221 | Endothelial | 14 | ENSMUSG00000041378 | Cldn5 | | 0.338245866 |
| 222 | Endothelial | 14 | ENSMUSG00000039167 | Adgrl4 | | 0.455678548 |
| 223 | Endothelial | 14 | ENSMUSG00000033191 | Tie1 | | 0.412433036 |
| 224 | Endothelial | 14 | ENSMUSG00000034911 | Ushbp1 | | 0.466234436 |

| | | | | | |
|---|---|---|---|---|---|
| 225 | Endothelial | 14 | ENSMUSG00000045377 | Tmem88 | 0.319185368 |
| 226 | Endothelial | 14 | ENSMUSG00000046916 | Myct1 | 0.377889139 |
| 227 | Endothelial | 14 | ENSMUSG00000047867 | Gimap6 | 0.278852407 |
| 228 | Endothelial | 14 | ENSMUSG00000045092 | S1pr1 | 0.349834076 |
| 229 | Endothelial | 14 | ENSMUSG00000044338 | Aplnr | 0.292118334 |
| 230 | Endothelial | 14 | ENSMUSG00000044562 | Rasip1 | 0.400601891 |
| 231 | Endothelial | 14 | ENSMUSG00000027684 | Mecom | 0.392509415 |
| 232 | Endothelial | 14 | ENSMUSG00000052921 | Arhgef15 | 0.330641719 |
| 233 | Endothelial | 14 | ENSMUSG00000079018 | Ly6c1 | 0.412797735 |
| 234 | Endothelial | 14 | ENSMUSG00000020717 | Pecam1 | 0.654472124 |
| 235 | Endothelial | 14 | ENSMUSG00000061353 | Cxcl12 | 0.305317963 |
| 236 | Endothelial | 14 | ENSMUSG00000002944 | Cd36 | 0.439373446 |
| 237 | Endothelial | 14 | ENSMUSG00000037936 | Scarb1 | 0.277573916 |
| 238 | Endothelial | 14 | ENSMUSG00000068079 | Tcf15 | 0.303925165 |
| 239 | Endothelial | 14 | ENSMUSG00000020154 | Ptprb | 0.688434229 |
| 240 | Endothelial | 14 | ENSMUSG00000031442 | Mcf2l | 0.325534339 |
| 241 | Endothelial | 14 | ENSMUSG00000027435 | Cd93 | 0.543705618 |
| 242 | Endothelial | 14 | ENSMUSG00000026921 | Egfl7 | 0.451413138 |
| 243 | Endothelial | 14 | ENSMUSG00000001930 | Vwf | 0.322714795 |
| 244 | Endothelial | 14 | ENSMUSG00000032125 | Robo4 | 0.414186804 |
| 245 | Endothelial | 14 | ENSMUSG00000001240 | Ramp2 | 0.34214761 |
| 246 | Endothelial | 14 | ENSMUSG00000041445 | Mmrn2 | 0.493304134 |
| 247 | Endothelial | 14 | ENSMUSG00000062960 | Kdr | 0.39208845 |
| 248 | Endothelial | 14 | ENSMUSG00000056492 | Adgrf5 | 0.427795755 |
| 249 | Endothelial | 14 | ENSMUSG00000041134 | Cyyr1 | 0.408176843 |
| 250 | Endothelial | 14 | ENSMUSG00000054690 | Emcn | 0.462698365 |
| 251 | ICC | 15 | ENSMUSG00000005087 | Cd44 | 0.105741256 |
| 252 | ICC | 15 | ENSMUSG00000005672 | Kit | 0.472435862 |
| 253 | ICC | 15 | ENSMUSG00000019943 | Atp2b1 | 0.115707369 |
| 254 | ICC | 15 | ENSMUSG00000020788 | Atp2a3 | 0.192845644 |
| 255 | ICC | 15 | ENSMUSG00000020902 | Ntn1 | 0.113943939 |
| 256 | ICC | 15 | ENSMUSG00000025203 | Scd2 | 0.145023309 |
| 257 | ICC | 15 | ENSMUSG00000025790 | Slco3a1 | 0.187650571 |
| 258 | ICC | 15 | ENSMUSG00000025969 | Nrp2 | 0.104587845 |
| 259 | ICC | 15 | ENSMUSG00000026223 | Itm2c | 0.110285076 |
| 260 | ICC | 15 | ENSMUSG00000026235 | Epha4 | 0.130356864 |
| 261 | ICC | 15 | ENSMUSG00000026721 | Rabgap1l | 0.117413003 |
| 262 | ICC | 15 | ENSMUSG00000026778 | Prkcq | 0.153962718 |
| 263 | ICC | 15 | ENSMUSG00000027217 | Tspan18 | 0.112502469 |
| 264 | ICC | 15 | ENSMUSG00000027524 | Edn3 | 0.218951782 |
| 265 | ICC | 15 | ENSMUSG00000028031 | Dkk2 | 0.291345987 |
| 266 | ICC | 15 | ENSMUSG00000029426 | Scarb2 | 0.127791103 |
| 267 | ICC | 15 | ENSMUSG00000029563 | Foxp2 | 0.107439058 |
| 268 | ICC | 15 | ENSMUSG00000031075 | Ano1 | 0.388303183 |
| 269 | ICC | 15 | ENSMUSG00000031842 | Pde4c | 0.279933998 |

| | | | | | |
|---|---|---|---|---|---|
| 270 | ICC | 15 | ENSMUSG00000038633 | Degs1 | 0.104907798 |
| 271 | ICC | 15 | ENSMUSG00000033161 | Atp1a1 | 0.120639876 |
| 272 | ICC | 15 | ENSMUSG00000042613 | Pbxip1 | 0.139793791 |
| 273 | ICC | 15 | ENSMUSG00000035967 | Ints6l | 0.204367493 |
| 274 | ICC | 15 | ENSMUSG00000025558 | Dock9 | 0.10429933 |
| 275 | ICC | 15 | ENSMUSG00000026110 | Mgat4a | 0.127721258 |
| 276 | ICC | 15 | ENSMUSG00000041741 | Pde3a | 0.242445625 |
| 277 | ICC | 15 | ENSMUSG00000021276 | Cinp | 0.10292136 |
| 278 | ICC | 15 | ENSMUSG00000037852 | Cpe | 0.156779729 |
| 279 | ICC | 15 | ENSMUSG00000050069 | Grem2 | 0.289432629 |
| 280 | ICC | 15 | ENSMUSG00000044017 | Adgrd1 | 0.40180573 |
| 281 | ICC | 15 | ENSMUSG00000045281 | Gpr20 | 0.294045075 |
| 282 | ICC | 15 | ENSMUSG00000049265 | Kcnk3 | 0.33609358 |
| 283 | ICC | 15 | ENSMUSG00000050953 | Gja1 | 0.400979029 |
| 284 | ICC | 15 | ENSMUSG00000056427 | Slit3 | 0.121536206 |
| 285 | ICC | 15 | ENSMUSG00000035566 | Pcdh17 | 0.332873664 |
| 286 | ICC | 15 | ENSMUSG00000041220 | Elovl6 | 0.256593828 |
| 287 | ICC | 15 | ENSMUSG00000020105 | Lrig3 | 0.116367819 |
| 288 | ICC | 15 | ENSMUSG00000064272 | Gpbar1 | 0.196463059 |
| 289 | ICC | 15 | ENSMUSG00000024112 | Cacna1h | 0.108176935 |
| 290 | ICC | 15 | ENSMUSG00000064345 | mt-Nd2 | 0.10657517 |
| 291 | ICC | 15 | ENSMUSG00000064370 | mt-Cytb | 0.108868598 |
| 292 | ICC | 15 | ENSMUSG00000007097 | Atp1a2 | 0.108901431 |
| 293 | ICC | 15 | ENSMUSG00000004151 | Etv1 | 0.371834482 |
| 294 | ICC | 15 | ENSMUSG00000057530 | Ece1 | 0.105754923 |
| 295 | ICC | 15 | ENSMUSG00000055632 | Hmcn2 | 0.109652063 |
| 296 | ICC | 15 | ENSMUSG00000034275 | Igsf9b | 0.262611672 |
| 297 | ICC | 15 | ENSMUSG00000024597 | Slc12a2 | 0.269467427 |
| 298 | ICC | 15 | ENSMUSG00000034472 | Rasd2 | 0.12319255 |
| 299 | ICC | 15 | ENSMUSG00000092341 | Malat1 | 0.14155268 |
| 300 | ICC | 15 | ENSMUSG00000097971 | Gm26917 | 0.138875279 |
| 301 | Macrophage | 16 | ENSMUSG00000035352 | Ccl12 | 0.408965857 |
| 302 | Macrophage | 16 | ENSMUSG00000000682 | Cd52 | 0.51558667 |
| 303 | Macrophage | 16 | ENSMUSG00000001128 | Cfp | 0.389930967 |
| 304 | Macrophage | 16 | ENSMUSG00000002111 | Spi1 | 0.367605196 |
| 305 | Macrophage | 16 | ENSMUSG00000002985 | Apoe | 0.356446159 |
| 306 | Macrophage | 16 | ENSMUSG00000004730 | Adgre1 | 0.417077553 |
| 307 | Macrophage | 16 | ENSMUSG00000015340 | Cybb | 0.438517351 |
| 308 | Macrophage | 16 | ENSMUSG00000018774 | Cd68 | 0.433539811 |
| 309 | Macrophage | 16 | ENSMUSG00000021423 | Ly86 | 0.46373745 |
| 310 | Macrophage | 16 | ENSMUSG00000024621 | Csf1r | 0.492080288 |
| 311 | Macrophage | 16 | ENSMUSG00000024672 | Ms4a7 | 0.36677246 |
| 312 | Macrophage | 16 | ENSMUSG00000024677 | Ms4a6b | 0.405135214 |
| 313 | Macrophage | 16 | ENSMUSG00000025150 | Cbr2 | 0.350743814 |
| 314 | Macrophage | 16 | ENSMUSG00000026126 | Ptpn18 | 0.409737014 |

| | | | | | |
|---|---|---|---|---|---|
| 315 | Macrophage | 16 | ENSMUSG00000026712 | Mrc1 | 0.5999411 |
| 316 | Macrophage | 16 | ENSMUSG00000028581 | Laptm5 | 0.597457244 |
| 317 | Macrophage | 16 | ENSMUSG00000029373 | Pf4 | 0.540841172 |
| 318 | Macrophage | 16 | ENSMUSG00000008845 | Cd163 | 0.433896409 |
| 319 | Macrophage | 16 | ENSMUSG00000030148 | Clec4a2 | 0.38782934 |
| 320 | Macrophage | 16 | ENSMUSG00000030560 | Ctsc | 0.447669341 |
| 321 | Macrophage | 16 | ENSMUSG00000030579 | Tyrobp | 0.492371848 |
| 322 | Macrophage | 16 | ENSMUSG00000030707 | Coro1a | 0.414652566 |
| 323 | Macrophage | 16 | ENSMUSG00000030787 | Lyve1 | 0.374207146 |
| 324 | Macrophage | 16 | ENSMUSG00000042286 | Stab1 | 0.484397353 |
| 325 | Macrophage | 16 | ENSMUSG00000040747 | Cd53 | 0.449093177 |
| 326 | Macrophage | 16 | ENSMUSG00000037443 | Cep85 | 0.367068616 |
| 327 | Macrophage | 16 | ENSMUSG00000024965 | Fermt3 | 0.360851439 |
| 328 | Macrophage | 16 | ENSMUSG00000040950 | Mgl2 | 0.438695992 |
| 329 | Macrophage | 16 | ENSMUSG00000040552 | C3ar1 | 0.369785159 |
| 330 | Macrophage | 16 | ENSMUSG00000018008 | Cyth4 | 0.400050982 |
| 331 | Macrophage | 16 | ENSMUSG00000033220 | Rac2 | 0.380658655 |
| 332 | Macrophage | 16 | ENSMUSG00000036887 | C1qa | 0.598041293 |
| 333 | Macrophage | 16 | ENSMUSG00000036896 | C1qc | 0.574313065 |
| 334 | Macrophage | 16 | ENSMUSG00000036905 | C1qb | 0.574915126 |
| 335 | Macrophage | 16 | ENSMUSG00000049130 | C5ar1 | 0.356942095 |
| 336 | Macrophage | 16 | ENSMUSG00000049988 | Lrrc25 | 0.376960236 |
| 337 | Macrophage | 16 | ENSMUSG00000048865 | Arhgap30 | 0.36467383 |
| 338 | Macrophage | 16 | ENSMUSG00000052160 | Pld4 | 0.476517227 |
| 339 | Macrophage | 16 | ENSMUSG00000052336 | Cx3cr1 | 0.396656424 |
| 340 | Macrophage | 16 | ENSMUSG00000060063 | Alox5ap | 0.376909731 |
| 341 | Macrophage | 16 | ENSMUSG00000060791 | Gmfg | 0.352313927 |
| 342 | Macrophage | 16 | ENSMUSG00000058715 | Fcer1g | 0.544770709 |
| 343 | Macrophage | 16 | ENSMUSG00000015852 | Fcrls | 0.430199505 |
| 344 | Macrophage | 16 | ENSMUSG00000069516 | Lyz2 | 0.517208469 |
| 345 | Macrophage | 16 | ENSMUSG00000020377 | Ltc4s | 0.383629486 |
| 346 | Macrophage | 16 | ENSMUSG00000026395 | Ptprc | 0.492522778 |
| 347 | Macrophage | 16 | ENSMUSG00000021998 | Lcp1 | 0.418018421 |
| 348 | Macrophage | 16 | ENSMUSG00000036908 | Unc93b1 | 0.356273805 |
| 349 | Macrophage | 16 | ENSMUSG00000059498 | Fcgr3 | 0.49076908 |
| 350 | Macrophage | 16 | ENSMUSG00000079419 | Ms4a6c | 0.370695231 |
| 351 | Fibroblasts | 17 | ENSMUSG00000020676 | Ccl11 | 0.254896789 |
| 352 | Fibroblasts | 17 | ENSMUSG00000000753 | Serpinf1 | 0.393992824 |
| 353 | Fibroblasts | 17 | ENSMUSG00000001506 | Col1a1 | 0.33317856 |
| 354 | Fibroblasts | 17 | ENSMUSG00000007805 | Twist2 | 0.22595677 |
| 355 | Fibroblasts | 17 | ENSMUSG00000001131 | Timp1 | 0.229656531 |
| 356 | Fibroblasts | 17 | ENSMUSG00000029675 | Eln | 0.210502468 |
| 357 | Fibroblasts | 17 | ENSMUSG00000015354 | Pcolce2 | 0.283959888 |
| 358 | Fibroblasts | 17 | ENSMUSG00000015568 | Lpl | 0.411083264 |
| 359 | Fibroblasts | 17 | ENSMUSG00000020363 | Gfpt2 | 0.217654329 |

| | | | | | |
|---|---|---|---|---|---|
| 360 | Fibroblasts | 17 | ENSMUSG00000020810 | Cygb | 0.328995558 |
| 361 | Fibroblasts | 17 | ENSMUSG00000021186 | Fbln5 | 0.278941411 |
| 362 | Fibroblasts | 17 | ENSMUSG00000022032 | Scara5 | 0.232381557 |
| 363 | Fibroblasts | 17 | ENSMUSG00000022371 | Col14a1 | 0.441264878 |
| 364 | Fibroblasts | 17 | ENSMUSG00000075602 | Ly6a | 0.290629976 |
| 365 | Fibroblasts | 17 | ENSMUSG00000024053 | Emilin2 | 0.222076871 |
| 366 | Fibroblasts | 17 | ENSMUSG00000024076 | Vit | 0.287002397 |
| 367 | Fibroblasts | 17 | ENSMUSG00000024713 | Pcsk5 | 0.226902707 |
| 368 | Fibroblasts | 17 | ENSMUSG00000025784 | Clec3b | 0.415983374 |
| 369 | Fibroblasts | 17 | ENSMUSG00000026586 | Prrx1 | 0.377848895 |
| 370 | Fibroblasts | 17 | ENSMUSG00000026748 | Plxdc2 | 0.255558082 |
| 371 | Fibroblasts | 17 | ENSMUSG00000026879 | Gsn | 0.477978999 |
| 372 | Fibroblasts | 17 | ENSMUSG00000026837 | Col5a1 | 0.238568665 |
| 373 | Fibroblasts | 17 | ENSMUSG00000027204 | Fbn1 | 0.322971228 |
| 374 | Fibroblasts | 17 | ENSMUSG00000029061 | Mmp23 | 0.259758332 |
| 375 | Fibroblasts | 17 | ENSMUSG00000029661 | Col1a2 | 0.334214462 |
| 376 | Fibroblasts | 17 | ENSMUSG00000029718 | Pcolce | 0.250302816 |
| 377 | Fibroblasts | 17 | ENSMUSG00000030116 | Mfap5 | 0.416966538 |
| 378 | Fibroblasts | 17 | ENSMUSG00000031239 | Itm2a | 0.224281513 |
| 379 | Fibroblasts | 17 | ENSMUSG00000031375 | Bgn | 0.253947839 |
| 380 | Fibroblasts | 17 | ENSMUSG00000036545 | Adamts2 | 0.32256485 |
| 381 | Fibroblasts | 17 | ENSMUSG00000064080 | Fbln2 | 0.256896209 |
| 382 | Fibroblasts | 17 | ENSMUSG00000040181 | Fmo1 | 0.271994727 |
| 383 | Fibroblasts | 17 | ENSMUSG00000038463 | Olfml2b | 0.220273534 |
| 384 | Fibroblasts | 17 | ENSMUSG00000006369 | Fbln1 | 0.222488828 |
| 385 | Fibroblasts | 17 | ENSMUSG00000032334 | Loxl1 | 0.238516338 |
| 386 | Fibroblasts | 17 | ENSMUSG00000020053 | Igf1 | 0.300057478 |
| 387 | Fibroblasts | 17 | ENSMUSG00000055172 | C1ra | 0.219758258 |
| 388 | Fibroblasts | 17 | ENSMUSG00000055653 | Gpc3 | 0.210899648 |
| 389 | Fibroblasts | 17 | ENSMUSG00000056481 | Cd248 | 0.335723328 |
| 390 | Fibroblasts | 17 | ENSMUSG00000060572 | Mfap2 | 0.214635455 |
| 391 | Fibroblasts | 17 | ENSMUSG00000057098 | Ebf1 | 0.337462396 |
| 392 | Fibroblasts | 17 | ENSMUSG00000028339 | Col15a1 | 0.214821631 |
| 393 | Fibroblasts | 17 | ENSMUSG00000033327 | Tnxb | 0.238282268 |
| 394 | Fibroblasts | 17 | ENSMUSG00000029096 | Htra3 | 0.303883512 |
| 395 | Fibroblasts | 17 | ENSMUSG00000026043 | Col3a1 | 0.319257988 |
| 396 | Fibroblasts | 17 | ENSMUSG00000019929 | Dcn | 0.221233148 |
| 397 | Fibroblasts | 17 | ENSMUSG00000021614 | Vcan | 0.307545233 |
| 398 | Fibroblasts | 17 | ENSMUSG00000024011 | Pi16 | 0.301079229 |
| 399 | Fibroblasts | 17 | ENSMUSG00000022816 | Fstl1 | 0.211162152 |
| 400 | Fibroblasts | 17 | ENSMUSG00000038521 | C1s1 | 0.223295679 |
| 401 | Unknown | 19 | ENSMUSG00000020447 | Npc1l1 | 0.623999698 |
| 402 | Unknown | 19 | ENSMUSG00000006345 | Ggt1 | 0.581916406 |
| 403 | Unknown | 19 | ENSMUSG00000011034 | Slc5a1 | 0.674186105 |
| 404 | Unknown | 19 | ENSMUSG00000018569 | Cldn7 | 0.529704662 |

| | | | | | |
|---|---|---|---|---|---|
| 405 | Unknown | 19 | ENSMUSG00000021278 | Amn | 0.627629799 |
| 406 | Unknown | 19 | ENSMUSG00000021565 | Slc6a19 | 0.661764165 |
| 407 | Unknown | 19 | ENSMUSG00000022824 | Muc13 | 0.49865082 |
| 408 | Unknown | 19 | ENSMUSG00000023057 | Fabp2 | 0.585735951 |
| 409 | Unknown | 19 | ENSMUSG00000023914 | Mep1a | 0.635602779 |
| 410 | Unknown | 19 | ENSMUSG00000024503 | Spink1 | 0.571252053 |
| 411 | Unknown | 19 | ENSMUSG00000025467 | Prap1 | 0.573255747 |
| 412 | Unknown | 19 | ENSMUSG00000025528 | 2010106E10Rik | 0.554349464 |
| 413 | Unknown | 19 | ENSMUSG00000028158 | Mttp | 0.523960003 |
| 414 | Unknown | 19 | ENSMUSG00000068547 | Clca4a | 0.499220086 |
| 415 | Unknown | 19 | ENSMUSG00000029700 | Slc13a1 | 0.482241986 |
| 416 | Unknown | 19 | ENSMUSG00000029727 | Cyp3a13 | 0.584006418 |
| 417 | Unknown | 19 | ENSMUSG00000030364 | Clec2h | 0.515271237 |
| 418 | Unknown | 19 | ENSMUSG00000030492 | Slc7a9 | 0.521230352 |
| 419 | Unknown | 19 | ENSMUSG00000031886 | Ces2e | 0.554649701 |
| 420 | Unknown | 19 | ENSMUSG00000032038 | St3gal4 | 0.503924086 |
| 421 | Unknown | 19 | ENSMUSG00000032081 | Apoc3 | 0.583276513 |
| 422 | Unknown | 19 | ENSMUSG00000032083 | Apoa1 | 0.681391787 |
| 423 | Unknown | 19 | ENSMUSG00000032098 | Treh | 0.485054434 |
| 424 | Unknown | 19 | ENSMUSG00000034918 | Cdhr2 | 0.57195295 |
| 425 | Unknown | 19 | ENSMUSG00000020609 | Apob | 0.693136887 |
| 426 | Unknown | 19 | ENSMUSG00000034427 | Myo15b | 0.4795872 |
| 427 | Unknown | 19 | ENSMUSG00000037390 | Muc3 | 0.685119043 |
| 428 | Unknown | 19 | ENSMUSG00000035699 | Slc51a | 0.48998425 |
| 429 | Unknown | 19 | ENSMUSG00000054999 | Naaladl1 | 0.61910795 |
| 430 | Unknown | 19 | ENSMUSG00000035000 | Dpp4 | 0.521175714 |
| 431 | Unknown | 19 | ENSMUSG00000039062 | Anpep | 0.725501757 |
| 432 | Unknown | 19 | ENSMUSG00000045394 | Epcam | 0.559586194 |
| 433 | Unknown | 19 | ENSMUSG00000024292 | Cyp4f14 | 0.631950878 |
| 434 | Unknown | 19 | ENSMUSG00000068587 | Mgam | 0.610538087 |
| 435 | Unknown | 19 | ENSMUSG00000020839 | Tmigd1 | 0.548948925 |
| 436 | Unknown | 19 | ENSMUSG00000015405 | Ace2 | 0.654489795 |
| 437 | Unknown | 19 | ENSMUSG00000091705 | H2-Q2 | 0.53259916 |
| 438 | Unknown | 19 | ENSMUSG00000025497 | Cdhr5 | 0.514434097 |
| 439 | Unknown | 19 | ENSMUSG00000024313 | Mep1b | 0.577410771 |
| 440 | Unknown | 19 | ENSMUSG00000025557 | Slc15a1 | 0.650916594 |
| 441 | Unknown | 19 | ENSMUSG00000046697 | Enpp7 | 0.515824093 |
| 442 | Unknown | 19 | ENSMUSG00000027790 | Sis | 0.572528964 |
| 443 | Unknown | 19 | ENSMUSG00000070777 | Ceacam20 | 0.584633903 |
| 444 | Unknown | 19 | ENSMUSG00000029188 | Slc34a2 | 0.598995514 |
| 445 | Unknown | 19 | ENSMUSG00000024887 | Asah2 | 0.519687281 |
| 446 | Unknown | 19 | ENSMUSG00000074195 | Clca4b | 0.664518943 |
| 447 | Unknown | 19 | ENSMUSG00000029134 | Plb1 | 0.568961012 |
| 448 | Unknown | 19 | ENSMUSG00000078439 | Smim24 | 0.529432023 |
| 449 | Unknown | 19 | ENSMUSG00000079440 | Alpi | 0.612222827 |

| | | | | | |
|---|---|---|---|---|---|
| 450 | Unknown | 19 | ENSMUSG00000035041 | Creb3l3 | 0.491323104 |
| 451 | ICC | 22 | ENSMUSG00000000088 | Cox5a | 0.220525005 |
| 452 | ICC | 22 | ENSMUSG00000003955 | Fam162a | 0.268638235 |
| 453 | ICC | 22 | ENSMUSG00000005672 | Kit | 0.230627145 |
| 454 | ICC | 22 | ENSMUSG00000006941 | Eif1b | 0.31478041 |
| 455 | ICC | 22 | ENSMUSG00000021396 | Nxnl2 | 0.229072451 |
| 456 | ICC | 22 | ENSMUSG00000024556 | Me2 | 0.273172472 |
| 457 | ICC | 22 | ENSMUSG00000024601 | Isoc1 | 0.271853699 |
| 458 | ICC | 22 | ENSMUSG00000025190 | Got1 | 0.22437491 |
| 459 | ICC | 22 | ENSMUSG00000025488 | Cox8b | 0.221778594 |
| 460 | ICC | 22 | ENSMUSG00000025790 | Slco3a1 | 0.217712218 |
| 461 | ICC | 22 | ENSMUSG00000026721 | Rabgap1l | 0.231308512 |
| 462 | ICC | 22 | ENSMUSG00000026778 | Prkcq | 0.447880904 |
| 463 | ICC | 22 | ENSMUSG00000027524 | Edn3 | 0.354548092 |
| 464 | ICC | 22 | ENSMUSG00000027562 | Car2 | 0.404318935 |
| 465 | ICC | 22 | ENSMUSG00000028031 | Dkk2 | 0.305574235 |
| 466 | ICC | 22 | ENSMUSG00000028278 | Rragd | 0.335758054 |
| 467 | ICC | 22 | ENSMUSG00000029128 | Rab28 | 0.24716687 |
| 468 | ICC | 22 | ENSMUSG00000029167 | Ppargc1a | 0.258167065 |
| 469 | ICC | 22 | ENSMUSG00000029563 | Foxp2 | 0.25682556 |
| 470 | ICC | 22 | ENSMUSG00000030246 | Ldhb | 0.305540549 |
| 471 | ICC | 22 | ENSMUSG00000030844 | Rgs10 | 0.427945941 |
| 472 | ICC | 22 | ENSMUSG00000031075 | Ano1 | 0.232312451 |
| 473 | ICC | 22 | ENSMUSG00000031367 | Ap1s2 | 0.240238663 |
| 474 | ICC | 22 | ENSMUSG00000031613 | Hpgd | 0.227938002 |
| 475 | ICC | 22 | ENSMUSG00000031842 | Pde4c | 0.303708549 |
| 476 | ICC | 22 | ENSMUSG00000037335 | Hand1 | 0.229273165 |
| 477 | ICC | 22 | ENSMUSG00000041741 | Pde3a | 0.294507701 |
| 478 | ICC | 22 | ENSMUSG00000035561 | Aldh1b1 | 0.369134262 |
| 479 | ICC | 22 | ENSMUSG00000035615 | Frmpd1 | 0.288525071 |
| 480 | ICC | 22 | ENSMUSG00000041012 | Cmtm8 | 0.225260824 |
| 481 | ICC | 22 | ENSMUSG00000050069 | Grem2 | 0.257664102 |
| 482 | ICC | 22 | ENSMUSG00000048387 | Osr1 | 0.254718653 |
| 483 | ICC | 22 | ENSMUSG00000049422 | Chchd10 | 0.279638757 |
| 484 | ICC | 22 | ENSMUSG00000046093 | Hpcal4 | 0.255025541 |
| 485 | ICC | 22 | ENSMUSG00000045281 | Gpr20 | 0.274019383 |
| 486 | ICC | 22 | ENSMUSG00000053199 | Arhgap20 | 0.224775243 |
| 487 | ICC | 22 | ENSMUSG00000049265 | Kcnk3 | 0.295853919 |
| 488 | ICC | 22 | ENSMUSG00000050953 | Gja1 | 0.218278818 |
| 489 | ICC | 22 | ENSMUSG00000035566 | Pcdh17 | 0.341348514 |
| 490 | ICC | 22 | ENSMUSG00000041220 | Elovl6 | 0.308519991 |
| 491 | ICC | 22 | ENSMUSG00000064272 | Gpbar1 | 0.233238497 |
| 492 | ICC | 22 | ENSMUSG00000053279 | Aldh1a1 | 0.266555014 |
| 493 | ICC | 22 | ENSMUSG00000059173 | Pde1a | 0.22006835 |
| 494 | ICC | 22 | ENSMUSG00000004151 | Etv1 | 0.252816753 |

| | | | | | |
|---|---|---|---|---|---|
| 495 | ICC | 22 | ENSMUSG00000074218 | Cox7a1 | 0.283431022 |
| 496 | ICC | 22 | ENSMUSG00000041390 | Mdfic | 0.297375058 |
| 497 | ICC | 22 | ENSMUSG00000034275 | Igsf9b | 0.237463009 |
| 498 | ICC | 22 | ENSMUSG00000024597 | Slc12a2 | 0.224965626 |
| 499 | ICC | 22 | ENSMUSG00000034472 | Rasd2 | 0.320783513 |
| 500 | ICC | 22 | ENSMUSG00000087075 | Lbhd2 | 0.374702156 |
| 501 | Unknown | 25 | ENSMUSG00000001247 | Lsr | 0.239630862 |
| 502 | Unknown | 25 | ENSMUSG00000013643 | Lypd8 | 0.666808534 |
| 503 | Unknown | 25 | ENSMUSG00000018919 | Tm4sf5 | 0.253695503 |
| 504 | Unknown | 25 | ENSMUSG00000020581 | Agr2 | 0.738440613 |
| 505 | Unknown | 25 | ENSMUSG00000021263 | Degs2 | 0.235054601 |
| 506 | Unknown | 25 | ENSMUSG00000022683 | Pla2g10 | 0.30496775 |
| 507 | Unknown | 25 | ENSMUSG00000022824 | Muc13 | 0.253227937 |
| 508 | Unknown | 25 | ENSMUSG00000023247 | Guca2a | 0.467291904 |
| 509 | Unknown | 25 | ENSMUSG00000024029 | Tff3 | 0.820250041 |
| 510 | Unknown | 25 | ENSMUSG00000033200 | Tpsg1 | 0.27305461 |
| 511 | Unknown | 25 | ENSMUSG00000025515 | Muc2 | 0.883511318 |
| 512 | Unknown | 25 | ENSMUSG00000026417 | Pigr | 0.397406873 |
| 513 | Unknown | 25 | ENSMUSG00000026961 | Lrrc26 | 0.415436002 |
| 514 | Unknown | 25 | ENSMUSG00000027801 | Tm4sf4 | 0.267918251 |
| 515 | Unknown | 25 | ENSMUSG00000028255 | Clca1 | 0.815852977 |
| 516 | Unknown | 25 | ENSMUSG00000028415 | Spink4 | 0.6261236 |
| 517 | Unknown | 25 | ENSMUSG00000028699 | Tspan1 | 0.402492835 |
| 518 | Unknown | 25 | ENSMUSG00000029102 | Hgfac | 0.335562959 |
| 519 | Unknown | 25 | ENSMUSG00000030866 | Ern2 | 0.441827275 |
| 520 | Unknown | 25 | ENSMUSG00000031608 | Galnt7 | 0.233474193 |
| 521 | Unknown | 25 | ENSMUSG00000032226 | Gcnt3 | 0.494388125 |
| 522 | Unknown | 25 | ENSMUSG00000034127 | Tspan8 | 0.317465472 |
| 523 | Unknown | 25 | ENSMUSG00000038991 | Txndc5 | 0.25995462 |
| 524 | Unknown | 25 | ENSMUSG00000040121 | Rep15 | 0.231265632 |
| 525 | Unknown | 25 | ENSMUSG00000043501 | Lgals2 | 0.220803029 |
| 526 | Unknown | 25 | ENSMUSG00000039774 | Galnt12 | 0.361223351 |
| 527 | Unknown | 25 | ENSMUSG00000038580 | Sct | 0.365945715 |
| 528 | Unknown | 25 | ENSMUSG00000040412 | 5330417C22Rik | 0.3652569 |
| 529 | Unknown | 25 | ENSMUSG00000037145 | 2210407C18Rik | 0.218763881 |
| 530 | Unknown | 25 | ENSMUSG00000044156 | Hepacam2 | 0.600224555 |
| 531 | Unknown | 25 | ENSMUSG00000049350 | Zg16 | 0.815604671 |
| 532 | Unknown | 25 | ENSMUSG00000047501 | Cldn4 | 0.217355054 |
| 533 | Unknown | 25 | ENSMUSG00000034107 | Ano7 | 0.417235158 |
| 534 | Unknown | 25 | ENSMUSG00000035506 | Slc12a8 | 0.525025084 |
| 535 | Unknown | 25 | ENSMUSG00000044626 | Liph | 0.256004611 |
| 536 | Unknown | 25 | ENSMUSG00000053964 | Lgals4 | 0.252886636 |
| 537 | Unknown | 25 | ENSMUSG00000054200 | Ffar4 | 0.242799381 |
| 538 | Unknown | 25 | ENSMUSG00000063903 | Klk1 | 0.49605056 |
| 539 | Unknown | 25 | ENSMUSG00000047730 | Fcgbp | 0.893426925 |

| | | | | | |
|---|---|---|---|---|---|
| 540 | Unknown | 25 | ENSMUSG00000044860 | Gm1123 | 0.544072647 |
| 541 | Unknown | 25 | ENSMUSG00000064213 | Defa24 | 0.279582031 |
| 542 | Unknown | 25 | ENSMUSG00000046410 | Kcnk6 | 0.232948278 |
| 543 | Unknown | 25 | ENSMUSG00000070473 | Cldn3 | 0.384512516 |
| 544 | Unknown | 25 | ENSMUSG00000034112 | Atp2c2 | 0.297395408 |
| 545 | Unknown | 25 | ENSMUSG00000071178 | Serpina1b | 0.469340014 |
| 546 | Unknown | 25 | ENSMUSG00000074227 | Spint2 | 0.254438868 |
| 547 | Unknown | 25 | ENSMUSG00000075010 | AW112010 | 0.283169901 |
| 548 | Unknown | 25 | ENSMUSG00000026828 | Galnt5 | 0.245765529 |
| 549 | Unknown | 25 | ENSMUSG00000079445 | B3gnt7 | 0.371032225 |
| 550 | Unknown | 25 | ENSMUSG00000094840 | Muc3a | 0.518697965 |
| 551 | SMC | 3 | ENSMUSG00000001506 | Col1a1 | 0.155838939 |
| 552 | SMC | 3 | ENSMUSG00000017009 | Sdc4 | 0.146894126 |
| 553 | SMC | 3 | ENSMUSG00000019326 | Aoc3 | 0.104363334 |
| 554 | SMC | 3 | ENSMUSG00000018566 | Slc2a4 | 0.14469887 |
| 555 | SMC | 3 | ENSMUSG00000000555 | Itga5 | 0.201215745 |
| 556 | SMC | 3 | ENSMUSG00000022836 | Mylk | 0.112132358 |
| 557 | SMC | 3 | ENSMUSG00000025780 | Itih5 | 0.125249817 |
| 558 | SMC | 3 | ENSMUSG00000026208 | Des | 0.123815636 |
| 559 | SMC | 3 | ENSMUSG00000026421 | Csrp1 | 0.119510053 |
| 560 | SMC | 3 | ENSMUSG00000027861 | Casq2 | 0.132181067 |
| 561 | SMC | 3 | ENSMUSG00000028108 | Ecm1 | 0.118747574 |
| 562 | SMC | 3 | ENSMUSG00000028461 | Ccdc107 | 0.129716458 |
| 563 | SMC | 3 | ENSMUSG00000028464 | Tpm2 | 0.120838122 |
| 564 | SMC | 3 | ENSMUSG00000032366 | Tpm1 | 0.114302651 |
| 565 | SMC | 3 | ENSMUSG00000028763 | Hspg2 | 0.135872886 |
| 566 | SMC | 3 | ENSMUSG00000028776 | Tinagl1 | 0.228001967 |
| 567 | SMC | 3 | ENSMUSG00000029309 | Sparcl1 | 0.22944687 |
| 568 | SMC | 3 | ENSMUSG00000029661 | Col1a2 | 0.141487004 |
| 569 | SMC | 3 | ENSMUSG00000031502 | Col4a1 | 0.168249955 |
| 570 | SMC | 3 | ENSMUSG00000031503 | Col4a2 | 0.234041848 |
| 571 | SMC | 3 | ENSMUSG00000032085 | Tagln | 0.129025769 |
| 572 | SMC | 3 | ENSMUSG00000033306 | Lpp | 0.123376304 |
| 573 | SMC | 3 | ENSMUSG00000042613 | Pbxip1 | 0.159116762 |
| 574 | SMC | 3 | ENSMUSG00000035783 | Acta2 | 0.160139894 |
| 575 | SMC | 3 | ENSMUSG00000032679 | Cd59a | 0.203298328 |
| 576 | SMC | 3 | ENSMUSG00000026463 | Atp2b4 | 0.26714529 |
| 577 | SMC | 3 | ENSMUSG00000037852 | Cpe | 0.121018046 |
| 578 | SMC | 3 | ENSMUSG00000026193 | Fn1 | 0.290930591 |
| 579 | SMC | 3 | ENSMUSG00000042284 | Itga1 | 0.267416492 |
| 580 | SMC | 3 | ENSMUSG00000023175 | Bsg | 0.109378925 |
| 581 | SMC | 3 | ENSMUSG00000031996 | Aplp2 | 0.11616325 |
| 582 | SMC | 3 | ENSMUSG00000063275 | Hacd1 | 0.117679663 |
| 583 | SMC | 3 | ENSMUSG00000059430 | Actg2 | 0.195039509 |
| 584 | SMC | 3 | ENSMUSG00000063873 | Slc24a3 | 0.120150063 |

| | | | | | |
|---|---|---|---|---|---|
| 585 | SMC | 3 | ENSMUSG00000064337 | mt-Rnr1 | 0.120149738 |
| 586 | SMC | 3 | ENSMUSG00000064339 | mt-Rnr2 | 0.122317996 |
| 587 | SMC | 3 | ENSMUSG00000064341 | mt-Nd1 | 0.164526188 |
| 588 | SMC | 3 | ENSMUSG00000064345 | mt-Nd2 | 0.165214395 |
| 589 | SMC | 3 | ENSMUSG00000064351 | mt-Co1 | 0.137779721 |
| 590 | SMC | 3 | ENSMUSG00000064356 | mt-Atp8 | 0.142925238 |
| 591 | SMC | 3 | ENSMUSG00000064363 | mt-Nd4 | 0.162819792 |
| 592 | SMC | 3 | ENSMUSG00000064367 | mt-Nd5 | 0.142646655 |
| 593 | SMC | 3 | ENSMUSG00000064368 | mt-Nd6 | 0.125797115 |
| 594 | SMC | 3 | ENSMUSG00000064370 | mt-Cytb | 0.157439418 |
| 595 | SMC | 3 | ENSMUSG00000067818 | Myl9 | 0.111981819 |
| 596 | SMC | 3 | ENSMUSG00000025809 | Itgb1 | 0.255088275 |
| 597 | SMC | 3 | ENSMUSG00000018830 | Myh11 | 0.13562463 |
| 598 | SMC | 3 | ENSMUSG00000070436 | Serpinh1 | 0.147836564 |
| 599 | SMC | 3 | ENSMUSG00000022816 | Fstl1 | 0.105874501 |
| 600 | SMC | 3 | ENSMUSG00000101939 | Gm28438 | 0.151073526 |
| 601 | Fibroblasts | 4 | ENSMUSG00000029231 | Pdgfra | 0.315891623 |
| 602 | Fibroblasts | 4 | ENSMUSG00000000794 | Kcnn3 | 0.253105294 |
| 603 | Fibroblasts | 4 | ENSMUSG00000001119 | Col6a1 | 0.188662982 |
| 604 | Fibroblasts | 4 | ENSMUSG00000020241 | Col6a2 | 0.207194419 |
| 605 | Fibroblasts | 4 | ENSMUSG00000001506 | Col1a1 | 0.171940725 |
| 606 | Fibroblasts | 4 | ENSMUSG00000002847 | Pla1a | 0.144942238 |
| 607 | Fibroblasts | 4 | ENSMUSG00000002900 | Lamb1 | 0.181419402 |
| 608 | Fibroblasts | 4 | ENSMUSG00000029675 | Eln | 0.239438918 |
| 609 | Fibroblasts | 4 | ENSMUSG00000018593 | Sparc | 0.142851594 |
| 610 | Fibroblasts | 4 | ENSMUSG00000019966 | Kitl | 0.13188295 |
| 611 | Fibroblasts | 4 | ENSMUSG00000020695 | Mrc2 | 0.176966086 |
| 612 | Fibroblasts | 4 | ENSMUSG00000022860 | Chodl | 0.163482825 |
| 613 | Fibroblasts | 4 | ENSMUSG00000023224 | Serping1 | 0.128914663 |
| 614 | Fibroblasts | 4 | ENSMUSG00000023886 | Smoc2 | 0.167009349 |
| 615 | Fibroblasts | 4 | ENSMUSG00000023913 | Pla2g7 | 0.157840213 |
| 616 | Fibroblasts | 4 | ENSMUSG00000025969 | Nrp2 | 0.145246819 |
| 617 | Fibroblasts | 4 | ENSMUSG00000026478 | Lamc1 | 0.128899984 |
| 618 | Fibroblasts | 4 | ENSMUSG00000028364 | Tnc | 0.348307908 |
| 619 | Fibroblasts | 4 | ENSMUSG00000029163 | Emilin1 | 0.215766673 |
| 620 | Fibroblasts | 4 | ENSMUSG00000029467 | Atp2a2 | 0.184583371 |
| 621 | Fibroblasts | 4 | ENSMUSG00000029661 | Col1a2 | 0.174960208 |
| 622 | Fibroblasts | 4 | ENSMUSG00000030218 | Mgp | 0.359322673 |
| 623 | Fibroblasts | 4 | ENSMUSG00000030605 | Mfge8 | 0.142195237 |
| 624 | Fibroblasts | 4 | ENSMUSG00000031375 | Bgn | 0.21385974 |
| 625 | Fibroblasts | 4 | ENSMUSG00000031740 | Mmp2 | 0.160459421 |
| 626 | Fibroblasts | 4 | ENSMUSG00000036446 | Lum | 0.287985331 |
| 627 | Fibroblasts | 4 | ENSMUSG00000039252 | Lgi2 | 0.269808236 |
| 628 | Fibroblasts | 4 | ENSMUSG00000042436 | Mfap4 | 0.209660533 |
| 629 | Fibroblasts | 4 | ENSMUSG00000028369 | Svep1 | 0.146802315 |

| | | | | | |
|---|---|---|---|---|---|
| 630 | Fibroblasts | 4 | ENSMUSG00000033880 | Lgals3bp | 0.15440402 |
| 631 | Fibroblasts | 4 | ENSMUSG00000037379 | Spon2 | 0.204063632 |
| 632 | Fibroblasts | 4 | ENSMUSG00000042254 | Cilp | 0.129890347 |
| 633 | Fibroblasts | 4 | ENSMUSG00000040249 | Lrp1 | 0.19055727 |
| 634 | Fibroblasts | 4 | ENSMUSG00000048126 | Col6a3 | 0.32521217 |
| 635 | Fibroblasts | 4 | ENSMUSG00000006369 | Fbln1 | 0.242422 |
| 636 | Fibroblasts | 4 | ENSMUSG00000048376 | F2r | 0.246899715 |
| 637 | Fibroblasts | 4 | ENSMUSG00000066705 | Fxyd6 | 0.144840649 |
| 638 | Fibroblasts | 4 | ENSMUSG00000026042 | Col5a2 | 0.130403846 |
| 639 | Fibroblasts | 4 | ENSMUSG00000026043 | Col3a1 | 0.140933198 |
| 640 | Fibroblasts | 4 | ENSMUSG00000067586 | S1pr3 | 0.140673718 |
| 641 | Fibroblasts | 4 | ENSMUSG00000025809 | Itgb1 | 0.133910016 |
| 642 | Fibroblasts | 4 | ENSMUSG00000019899 | Lama2 | 0.142732701 |
| 643 | Fibroblasts | 4 | ENSMUSG00000070436 | Serpinh1 | 0.132793543 |
| 644 | Fibroblasts | 4 | ENSMUSG00000053062 | Jam2 | 0.198178922 |
| 645 | Fibroblasts | 4 | ENSMUSG00000074971 | Fibin | 0.282975321 |
| 646 | Fibroblasts | 4 | ENSMUSG00000019929 | Dcn | 0.201966203 |
| 647 | Fibroblasts | 4 | ENSMUSG00000055632 | Hmcn2 | 0.176125509 |
| 648 | Fibroblasts | 4 | ENSMUSG00000058297 | Spock2 | 0.146083501 |
| 649 | Fibroblasts | 4 | ENSMUSG00000021943 | Gdf10 | 0.178366401 |
| 650 | Fibroblasts | 4 | ENSMUSG00000023885 | Thbs2 | 0.354766374 |
| 651 | Neuroglia | 5 | ENSMUSG00000000489 | Pdgfb | 0.256088954 |
| 652 | Neuroglia | 5 | ENSMUSG00000002985 | Apoe | 0.263595076 |
| 653 | Neuroglia | 5 | ENSMUSG00000018822 | Sfrp5 | 0.26534908 |
| 654 | Neuroglia | 5 | ENSMUSG00000022103 | Gfra2 | 0.300661247 |
| 655 | Neuroglia | 5 | ENSMUSG00000022324 | Matn2 | 0.334333396 |
| 656 | Neuroglia | 5 | ENSMUSG00000025089 | Gfra1 | 0.242709071 |
| 657 | Neuroglia | 5 | ENSMUSG00000025780 | Itih5 | 0.268824361 |
| 658 | Neuroglia | 5 | ENSMUSG00000026249 | Serpine2 | 0.274455048 |
| 659 | Neuroglia | 5 | ENSMUSG00000026424 | Gpr37l1 | 0.569089614 |
| 660 | Neuroglia | 5 | ENSMUSG00000027375 | Mal | 0.283049013 |
| 661 | Neuroglia | 5 | ENSMUSG00000030342 | Cd9 | 0.236413895 |
| 662 | Neuroglia | 5 | ENSMUSG00000031425 | Plp1 | 0.544622159 |
| 663 | Neuroglia | 5 | ENSMUSG00000031610 | Scrg1 | 0.226619533 |
| 664 | Neuroglia | 5 | ENSMUSG00000038400 | Pmepa1 | 0.424641834 |
| 665 | Neuroglia | 5 | ENSMUSG00000033208 | S100b | 0.265798115 |
| 666 | Neuroglia | 5 | ENSMUSG00000040759 | Cmtm5 | 0.355696931 |
| 667 | Neuroglia | 5 | ENSMUSG00000040724 | Kcna2 | 0.295425281 |
| 668 | Neuroglia | 5 | ENSMUSG00000036570 | Fxyd1 | 0.233989367 |
| 669 | Neuroglia | 5 | ENSMUSG00000033006 | Sox10 | 0.397429567 |
| 670 | Neuroglia | 5 | ENSMUSG00000036169 | Sostdc1 | 0.35731447 |
| 671 | Neuroglia | 5 | ENSMUSG00000034810 | Scn7a | 0.579019759 |
| 672 | Neuroglia | 5 | ENSMUSG00000040690 | Col16a1 | 0.245140911 |
| 673 | Neuroglia | 5 | ENSMUSG00000039239 | Tgfb2 | 0.287471193 |
| 674 | Neuroglia | 5 | ENSMUSG00000025856 | Pdgfa | 0.266692768 |

| | | | | | |
|---|---|---|---|---|---|
| 675 | Neuroglia | 5 | ENSMUSG00000042401 | Crtac1 | 0.242819975 |
| 676 | Neuroglia | 5 | ENSMUSG00000047976 | Kcna1 | 0.434627484 |
| 677 | Neuroglia | 5 | ENSMUSG00000044708 | Kcnj10 | 0.225861994 |
| 678 | Neuroglia | 5 | ENSMUSG00000046618 | Olfml2a | 0.318692929 |
| 679 | Neuroglia | 5 | ENSMUSG00000031342 | Gpm6b | 0.414368595 |
| 680 | Neuroglia | 5 | ENSMUSG00000031391 | L1cam | 0.255759231 |
| 681 | Neuroglia | 5 | ENSMUSG00000030077 | Chl1 | 0.227656196 |
| 682 | Neuroglia | 5 | ENSMUSG00000054793 | Cadm4 | 0.23481436 |
| 683 | Neuroglia | 5 | ENSMUSG00000020926 | Adam11 | 0.426588573 |
| 684 | Neuroglia | 5 | ENSMUSG00000073418 | C4b | 0.23945178 |
| 685 | Neuroglia | 5 | ENSMUSG00000032268 | Tmprss5 | 0.379107985 |
| 686 | Neuroglia | 5 | ENSMUSG00000032332 | Col12a1 | 0.349840108 |
| 687 | Neuroglia | 5 | ENSMUSG00000032373 | Car12 | 0.241434707 |
| 688 | Neuroglia | 5 | ENSMUSG00000001435 | Col18a1 | 0.344960915 |
| 689 | Neuroglia | 5 | ENSMUSG00000027750 | Postn | 0.227561429 |
| 690 | Neuroglia | 5 | ENSMUSG00000057969 | Sema3b | 0.233734736 |
| 691 | Neuroglia | 5 | ENSMUSG00000056966 | Gjc3 | 0.268802288 |
| 692 | Neuroglia | 5 | ENSMUSG00000067261 | Foxd3 | 0.346907725 |
| 693 | Neuroglia | 5 | ENSMUSG00000025964 | Adam23 | 0.378889129 |
| 694 | Neuroglia | 5 | ENSMUSG00000068748 | Ptprz1 | 0.403589511 |
| 695 | Neuroglia | 5 | ENSMUSG00000027966 | Col11a1 | 0.291658525 |
| 696 | Neuroglia | 5 | ENSMUSG00000047216 | Cdh19 | 0.425446519 |
| 697 | Neuroglia | 5 | ENSMUSG00000039546 | Ajap1 | 0.290391156 |
| 698 | Neuroglia | 5 | ENSMUSG00000016356 | Col20a1 | 0.357022675 |
| 699 | Neuroglia | 5 | ENSMUSG00000039542 | Ncam1 | 0.362473285 |
| 700 | Neuroglia | 5 | ENSMUSG00000024998 | Plce1 | 0.228835893 |
| 701 | Fibroblasts | 7 | ENSMUSG00000000901 | Mmp11 | 0.156034459 |
| 702 | Fibroblasts | 7 | ENSMUSG00000001349 | Cnn1 | 0.10312321 |
| 703 | Fibroblasts | 7 | ENSMUSG00000003534 | Ddr1 | 0.097050585 |
| 704 | Fibroblasts | 7 | ENSMUSG00000007892 | Rplp1 | 0.101096666 |
| 705 | Fibroblasts | 7 | ENSMUSG00000020599 | Rgs9 | 0.162164475 |
| 706 | Fibroblasts | 7 | ENSMUSG00000021094 | Dhrs7 | 0.118073655 |
| 707 | Fibroblasts | 7 | ENSMUSG00000021508 | Cxcl14 | 0.464821624 |
| 708 | Fibroblasts | 7 | ENSMUSG00000021702 | Thbs4 | 0.255061556 |
| 709 | Fibroblasts | 7 | ENSMUSG00000023886 | Smoc2 | 0.177426209 |
| 710 | Fibroblasts | 7 | ENSMUSG00000024526 | Cidea | 0.171153544 |
| 711 | Fibroblasts | 7 | ENSMUSG00000026208 | Des | 0.117077263 |
| 712 | Fibroblasts | 7 | ENSMUSG00000026421 | Csrp1 | 0.120420143 |
| 713 | Fibroblasts | 7 | ENSMUSG00000028464 | Tpm2 | 0.102276602 |
| 714 | Fibroblasts | 7 | ENSMUSG00000032366 | Tpm1 | 0.122017284 |
| 715 | Fibroblasts | 7 | ENSMUSG00000029309 | Sparcl1 | 0.180012985 |
| 716 | Fibroblasts | 7 | ENSMUSG00000029669 | Tspan12 | 0.256711685 |
| 717 | Fibroblasts | 7 | ENSMUSG00000030029 | Lrig1 | 0.260313548 |
| 718 | Fibroblasts | 7 | ENSMUSG00000030043 | Tacr1 | 0.113256828 |
| 719 | Fibroblasts | 7 | ENSMUSG00000030096 | Slc6a6 | 0.144934101 |

| | | | | | |
|---|---|---|---|---|---|
| 720 | Fibroblasts | 7 | ENSMUSG00000031328 | Flna | 0.125149814 |
| 721 | Fibroblasts | 7 | ENSMUSG00000031548 | Sfrp1 | 0.162291119 |
| 722 | Fibroblasts | 7 | ENSMUSG00000031586 | Rbpms | 0.127021467 |
| 723 | Fibroblasts | 7 | ENSMUSG00000032085 | Tagln | 0.131907948 |
| 724 | Fibroblasts | 7 | ENSMUSG00000032338 | Hcn4 | 0.147590835 |
| 725 | Fibroblasts | 7 | ENSMUSG00000038319 | Kcnh2 | 0.114662895 |
| 726 | Fibroblasts | 7 | ENSMUSG00000038591 | Colec10 | 0.114162461 |
| 727 | Fibroblasts | 7 | ENSMUSG00000036264 | Fstl4 | 0.13709877 |
| 728 | Fibroblasts | 7 | ENSMUSG00000035783 | Acta2 | 0.110215746 |
| 729 | Fibroblasts | 7 | ENSMUSG00000035493 | Tgfbi | 0.193651245 |
| 730 | Fibroblasts | 7 | ENSMUSG00000037852 | Cpe | 0.115277983 |
| 731 | Fibroblasts | 7 | ENSMUSG00000053907 | Mat2a | 0.149110302 |
| 732 | Fibroblasts | 7 | ENSMUSG00000048616 | Nog | 0.326680609 |
| 733 | Fibroblasts | 7 | ENSMUSG00000031283 | Chrdl1 | 0.311208603 |
| 734 | Fibroblasts | 7 | ENSMUSG00000030077 | Chl1 | 0.103759009 |
| 735 | Fibroblasts | 7 | ENSMUSG00000026387 | Sctr | 0.2309947 |
| 736 | Fibroblasts | 7 | ENSMUSG00000059430 | Actg2 | 0.109492896 |
| 737 | Fibroblasts | 7 | ENSMUSG00000031673 | Cdh11 | 0.105771398 |
| 738 | Fibroblasts | 7 | ENSMUSG00000063727 | Tnfrsf11b | 0.207872509 |
| 739 | Fibroblasts | 7 | ENSMUSG00000067818 | Myl9 | 0.113974587 |
| 740 | Fibroblasts | 7 | ENSMUSG00000018830 | Myh11 | 0.113333654 |
| 741 | Fibroblasts | 7 | ENSMUSG00000074934 | Grem1 | 0.308227433 |
| 742 | Fibroblasts | 7 | ENSMUSG00000087006 | Gm13889 | 0.113079742 |
| 743 | Fibroblasts | 7 | ENSMUSG00000057530 | Ece1 | 0.271550726 |
| 744 | Fibroblasts | 7 | ENSMUSG00000063564 | Col23a1 | 0.129003708 |
| 745 | Fibroblasts | 7 | ENSMUSG00000055632 | Hmcn2 | 0.109339341 |
| 746 | Fibroblasts | 7 | ENSMUSG00000062980 | Cped1 | 0.151897675 |
| 747 | Fibroblasts | 7 | ENSMUSG00000032572 | Col6a4 | 0.114972444 |
| 748 | Fibroblasts | 7 | ENSMUSG00000090841 | Myl6 | 0.116919605 |
| 749 | Fibroblasts | 7 | ENSMUSG00000070469 | Adamtsl3 | 0.114264039 |
| 750 | Fibroblasts | 7 | ENSMUSG00000092341 | Malat1 | 0.10130088 |
| 751 | NENs | 8 | ENSMUSG00000055022 | Cntn1 | 0.299539797 |
| 752 | NENs | 8 | ENSMUSG00000003410 | Elavl3 | 0.295101612 |
| 753 | NENs | 8 | ENSMUSG00000019796 | Lrp11 | 0.290948425 |
| 754 | NENs | 8 | ENSMUSG00000020297 | Nsg2 | 0.38027727 |
| 755 | NENs | 8 | ENSMUSG00000021087 | Rtn1 | 0.64887941 |
| 756 | NENs | 8 | ENSMUSG00000022577 | Ly6h | 0.302948392 |
| 757 | NENs | 8 | ENSMUSG00000023011 | Faim2 | 0.322310628 |
| 758 | NENs | 8 | ENSMUSG00000023484 | Prph | 0.36804432 |
| 759 | NENs | 8 | ENSMUSG00000024261 | Syt4 | 0.312764693 |
| 760 | NENs | 8 | ENSMUSG00000024907 | Gal | 0.346487259 |
| 761 | NENs | 8 | ENSMUSG00000026204 | Ptprn | 0.535497229 |
| 762 | NENs | 8 | ENSMUSG00000027273 | Snap25 | 0.382277374 |
| 763 | NENs | 8 | ENSMUSG00000027350 | Chgb | 0.369335867 |
| 764 | NENs | 8 | ENSMUSG00000027419 | Pcsk2 | 0.301097269 |

| | | | | | |
|---|---|---|---|---|---|
| 765 | NENs | 8 | ENSMUSG00000027500 | Stmn2 | 0.407681502 |
| 766 | NENs | 8 | ENSMUSG00000028137 | Celf3 | 0.389315233 |
| 767 | NENs | 8 | ENSMUSG00000029126 | Nsg1 | 0.315575873 |
| 768 | NENs | 8 | ENSMUSG00000029219 | Slc10a4 | 0.321701544 |
| 769 | NENs | 8 | ENSMUSG00000030110 | Ret | 0.33536641 |
| 770 | NENs | 8 | ENSMUSG00000032076 | Cadm1 | 0.349098213 |
| 771 | NENs | 8 | ENSMUSG00000032181 | Scg3 | 0.329529879 |
| 772 | NENs | 8 | ENSMUSG00000032303 | Chrna3 | 0.399998204 |
| 773 | NENs | 8 | ENSMUSG00000033061 | Resp18 | 0.457848693 |
| 774 | NENs | 8 | ENSMUSG00000039278 | Pcsk1n | 0.546501933 |
| 775 | NENs | 8 | ENSMUSG00000004113 | Cacna1b | 0.333358567 |
| 776 | NENs | 8 | ENSMUSG00000037428 | Vgf | 0.37623764 |
| 777 | NENs | 8 | ENSMUSG00000070695 | Cntnap5a | 0.298051712 |
| 778 | NENs | 8 | ENSMUSG00000035964 | Tmem59l | 0.36390734 |
| 779 | NENs | 8 | ENSMUSG00000041608 | Entpd3 | 0.305067601 |
| 780 | NENs | 8 | ENSMUSG00000050711 | Scg2 | 0.628591489 |
| 781 | NENs | 8 | ENSMUSG00000024109 | Nrxn1 | 0.363563914 |
| 782 | NENs | 8 | ENSMUSG00000044288 | Cnr1 | 0.357050602 |
| 783 | NENs | 8 | ENSMUSG00000055567 | Unc80 | 0.316831235 |
| 784 | NENs | 8 | ENSMUSG00000035864 | Syt1 | 0.480525917 |
| 785 | NENs | 8 | ENSMUSG00000004961 | Syt5 | 0.315001808 |
| 786 | NENs | 8 | ENSMUSG00000057182 | Scn3a | 0.373733997 |
| 787 | NENs | 8 | ENSMUSG00000054013 | Tmem179 | 0.320108549 |
| 788 | NENs | 8 | ENSMUSG00000061576 | Dpp6 | 0.369334274 |
| 789 | NENs | 8 | ENSMUSG00000033981 | Gria2 | 0.328984809 |
| 790 | NENs | 8 | ENSMUSG00000061762 | Tac1 | 0.301445314 |
| 791 | NENs | 8 | ENSMUSG00000068923 | Syt11 | 0.357455318 |
| 792 | NENs | 8 | ENSMUSG00000069072 | Slc7a14 | 0.392168956 |
| 793 | NENs | 8 | ENSMUSG00000023945 | Slc5a7 | 0.371101233 |
| 794 | NENs | 8 | ENSMUSG00000018411 | Mapt | 0.321667807 |
| 795 | NENs | 8 | ENSMUSG00000028546 | Elavl4 | 0.396577617 |
| 796 | NENs | 8 | ENSMUSG00000027581 | Stmn3 | 0.364941262 |
| 797 | NENs | 8 | ENSMUSG00000044349 | Snhg11 | 0.531561955 |
| 798 | NENs | 8 | ENSMUSG00000021268 | Meg3 | 0.295118744 |
| 799 | NENs | 8 | ENSMUSG00000021700 | Rab3c | 0.30188109 |
| 800 | NENs | 8 | ENSMUSG00000097311 | Gm26871 | 0.364572063 |
| 801 | MENs | 9 | ENSMUSG00000001025 | S100a6 | 0.262503174 |
| 802 | MENs | 9 | ENSMUSG00000002980 | Bcam | 0.233576279 |
| 803 | MENs | 9 | ENSMUSG00000006360 | Crip1 | 0.26940636 |
| 804 | MENs | 9 | ENSMUSG00000020911 | Krt19 | 0.298327534 |
| 805 | MENs | 9 | ENSMUSG00000009281 | Rarres2 | 0.342753636 |
| 806 | MENs | 9 | ENSMUSG00000017969 | Ptgis | 0.219217982 |
| 807 | MENs | 9 | ENSMUSG00000020186 | Csrp2 | 0.240573473 |
| 808 | MENs | 9 | ENSMUSG00000020467 | Efemp1 | 0.338016146 |
| 809 | MENs | 9 | ENSMUSG00000023046 | Igfbp6 | 0.54482707 |

| | | | | | |
|---|---|---|---|---|---|
| 810 | MENs | 9 | ENSMUSG00000023224 | Serping1 | 0.215587423 |
| 811 | MENs | 9 | ENSMUSG00000024164 | C3 | 0.545390358 |
| 812 | MENs | 9 | ENSMUSG00000026185 | Igfbp5 | 0.278488869 |
| 813 | MENs | 9 | ENSMUSG00000027357 | Crls1 | 0.239019825 |
| 814 | MENs | 9 | ENSMUSG00000053897 | Slc39a8 | 0.226657682 |
| 815 | MENs | 9 | ENSMUSG00000028583 | Pdpn | 0.285918915 |
| 816 | MENs | 9 | ENSMUSG00000028871 | Rspo1 | 0.337464481 |
| 817 | MENs | 9 | ENSMUSG00000031112 | Stk26 | 0.184220241 |
| 818 | MENs | 9 | ENSMUSG00000031517 | Gpm6a | 0.451723341 |
| 819 | MENs | 9 | ENSMUSG00000109564 | Muc16 | 0.344017365 |
| 820 | MENs | 9 | ENSMUSG00000032231 | Anxa2 | 0.204954172 |
| 821 | MENs | 9 | ENSMUSG00000039457 | Ppl | 0.230079209 |
| 822 | MENs | 9 | ENSMUSG00000039831 | Arhgap29 | 0.208332126 |
| 823 | MENs | 9 | ENSMUSG00000038725 | Pkhd1l1 | 0.314950067 |
| 824 | MENs | 9 | ENSMUSG00000040713 | Creg1 | 0.187683688 |
| 825 | MENs | 9 | ENSMUSG00000064080 | Fbln2 | 0.194905177 |
| 826 | MENs | 9 | ENSMUSG00000022425 | Enpp2 | 0.19093109 |
| 827 | MENs | 9 | ENSMUSG00000060962 | Dmkn | 0.267305225 |
| 828 | MENs | 9 | ENSMUSG00000040703 | Cyp2s1 | 0.215164272 |
| 829 | MENs | 9 | ENSMUSG00000040170 | Fmo2 | 0.24539683 |
| 830 | MENs | 9 | ENSMUSG00000042357 | Gjb5 | 0.293710381 |
| 831 | MENs | 9 | ENSMUSG00000041559 | Fmod | 0.208106954 |
| 832 | MENs | 9 | ENSMUSG00000043110 | Lrrn4 | 0.514390443 |
| 833 | MENs | 9 | ENSMUSG00000040856 | Dlk1 | 0.266488222 |
| 834 | MENs | 9 | ENSMUSG00000049436 | Upk1b | 0.450818946 |
| 835 | MENs | 9 | ENSMUSG00000022665 | Ccdc80 | 0.225868281 |
| 836 | MENs | 9 | ENSMUSG00000042985 | Upk3b | 0.565948874 |
| 837 | MENs | 9 | ENSMUSG00000052957 | Gas1 | 0.209179581 |
| 838 | MENs | 9 | ENSMUSG00000023039 | Krt7 | 0.304450314 |
| 839 | MENs | 9 | ENSMUSG00000063011 | Msln | 0.488152429 |
| 840 | MENs | 9 | ENSMUSG00000061451 | Tmem151a | 0.234403411 |
| 841 | MENs | 9 | ENSMUSG00000053522 | Lgals7 | 0.234922891 |
| 842 | MENs | 9 | ENSMUSG00000036098 | Myrf | 0.225383131 |
| 843 | MENs | 9 | ENSMUSG00000016918 | Sulf1 | 0.281158033 |
| 844 | MENs | 9 | ENSMUSG00000068220 | Lgals1 | 0.231426298 |
| 845 | MENs | 9 | ENSMUSG00000020473 | Aebp1 | 0.437471375 |
| 846 | MENs | 9 | ENSMUSG00000027574 | Nkain4 | 0.3147652 |
| 847 | MENs | 9 | ENSMUSG00000019929 | Dcn | 0.300216657 |
| 848 | MENs | 9 | ENSMUSG00000017002 | Slpi | 0.542461782 |
| 849 | MENs | 9 | ENSMUSG00000016458 | Wt1 | 0.293277401 |
| 850 | MENs | 9 | ENSMUSG00000040612 | Ildr2 | 0.244345184 |

| mean_expression | fraction_expressing | specificity | pseudo_R2 | marker_test_p_value |
|---|---|---|---|---|
| 5.499081378 | 0.924185168 | 0.158044172 | 0.185072334 | 0 |
| 6.081814569 | 0.941898914 | 0.163007696 | 0.23799343 | 0 |
| 1.812158131 | 0.583136514 | 0.143939401 | 0.040598069 | 4.43E-76 |
| 5.33365418 | 0.908833255 | 0.126065572 | 0.07285064 | 1.26E-136 |
| 2.536295289 | 0.707368918 | 0.153324699 | 0.063642354 | 3.40E-119 |
| 1.732123685 | 0.565658951 | 0.141090298 | 0.020831327 | 1.24E-39 |
| 1.631506588 | 0.537789325 | 0.149515288 | 0.019679969 | 1.60E-37 |
| 2.347990495 | 0.661549362 | 0.155852546 | 0.054124626 | 2.66E-101 |
| 4.509980321 | 0.866084081 | 0.098485239 | 0.010430008 | 1.36E-20 |
| 2.173880632 | 0.643127067 | 0.121121626 | 0.017316898 | 3.42E-33 |
| 18.90668949 | 0.998110534 | 0.178595131 | 0.493465288 | 0 |
| 1.412369763 | 0.485356637 | 0.159419336 | 0.055409604 | 1.04E-103 |
| 6.488837663 | 0.951346245 | 0.153180397 | 0.177887028 | 0 |
| 25.64453935 | 0.999527633 | 0.16101411 | 0.457145316 | 0 |
| 16.89699197 | 0.998346717 | 0.148161881 | 0.348481317 | 0 |
| 4.151102986 | 0.840340104 | 0.136979331 | 0.075510313 | 1.10E-141 |
| 2.639668518 | 0.723193198 | 0.179757273 | 0.11930439 | 1.82E-226 |
| 1.926344374 | 0.623051488 | 0.136299866 | 0.034677946 | 4.07E-65 |
| 7.068618802 | 0.962919225 | 0.147276684 | 0.190521229 | 0 |
| 2.202557307 | 0.650448748 | 0.119114793 | 0.011586225 | 1.04E-22 |
| 1.563094423 | 0.543693906 | 0.16178744 | 0.052288771 | 7.29E-98 |
| 6.229623579 | 0.932687766 | 0.165419736 | 0.181437142 | 0 |
| 2.413839193 | 0.571799717 | 0.174737777 | 0.061102197 | 2.09E-114 |
| 27.23184672 | 0.999763817 | 0.146929447 | 0.333230712 | 0 |
| 3.615978413 | 0.820264525 | 0.105402259 | 0.016671753 | 5.18E-32 |
| 1.935764222 | 0.60014171 | 0.16465666 | 0.050914392 | 2.71E-95 |
| 2.044780244 | 0.619036372 | 0.135760484 | 0.030490874 | 2.18E-57 |
| 3.789199691 | 0.831601323 | 0.117348232 | 0.034867579 | 1.82E-65 |
| 29.65805144 | 0.999763817 | 0.153262253 | 0.345482918 | 0 |
| 2.844992267 | 0.741143127 | 0.118789719 | 0.023852726 | 3.56E-45 |
| 13.71470322 | 0.967170524 | 0.118690443 | 0.058559443 | 1.27E-109 |
| 50.873854 | 1 | 0.107467211 | 0.082028323 | 4.02E-154 |
| 14.38502083 | 0.985356637 | 0.119428533 | 0.068339971 | 4.51E-128 |
| 17.41225149 | 0.995512518 | 0.12546189 | 0.132519506 | 1.25E-252 |
| 3.41332044 | 0.754369391 | 0.121278031 | 0.012951664 | 3.31E-25 |
| 13.55364915 | 0.985829003 | 0.11405164 | 0.06203638 | 3.63E-116 |
| 28.4216941 | 0.99929145 | 0.115723041 | 0.090642516 | 1.18E-170 |
| 14.04290752 | 0.998110534 | 0.153283123 | 0.376011126 | 0 |
| 3.125201788 | 0.76948512 | 0.102893822 | 0.03956113 | 3.71E-74 |
| 16.28068284 | 0.998110534 | 0.157005441 | 0.361954771 | 0 |
| 4.259915055 | 0.850259802 | 0.100127172 | 0.236859838 | 0 |
| 12.89231392 | 0.995984884 | 0.113786861 | 0.171253745 | 0 |
| 1.864920696 | 0.588332546 | 0.142151304 | 0.03102217 | 2.29E-58 |
| 4.358061918 | 0.860888049 | 0.103732369 | 0.015431637 | 9.64E-30 |

| | | | | |
|---|---|---|---|---|
| 3.022882137 | 0.754841757 | 0.104504902 | 0.010131857 | 4.77E-20 |
| 10.79981243 | 0.993386868 | 0.143938041 | 0.287917358 | 0 |
| 3.926299986 | 0.842701937 | 0.106668704 | 0.0133869 | 5.30E-26 |
| 3.991595123 | 0.718233349 | 0.161011946 | 0.055572854 | 5.14E-104 |
| 4.645012622 | 0.75720359 | 0.112849068 | 0.000641165 | 0.021303168 |
| 33.82427369 | 0.999527633 | 0.154717682 | 0.369454679 | 0 |
| 0.299734743 | 0.383333333 | 0.361428533 | 0.123162188 | 1.09E-85 |
| 5.574472781 | 0.95 | 0.42471989 | 0.440896096 | 9.42687454720685e-316 |
| 1.421238405 | 0.742857143 | 0.397645419 | 0.277486329 | 8.86E-195 |
| 1.148436159 | 0.683333333 | 0.192435174 | 0.105119072 | 3.14E-73 |
| 0.397154824 | 0.438095238 | 0.337046684 | 0.140237417 | 1.57E-97 |
| 0.42875304 | 0.423809524 | 0.336931614 | 0.1175335 | 8.55E-82 |
| 1.030850992 | 0.621428571 | 0.2147438 | 0.098394201 | 1.33E-68 |
| 2.744705938 | 0.926190476 | 0.163631499 | 0.132660293 | 2.87E-92 |
| 0.321615999 | 0.383333333 | 0.354113112 | 0.103820144 | 2.46E-72 |
| 0.659821377 | 0.557142857 | 0.333662844 | 0.17472305 | 1.29E-121 |
| 0.828105044 | 0.542857143 | 0.382640406 | 0.164993241 | 8.50E-115 |
| 1.59061247 | 0.730952381 | 0.173848541 | 0.106022686 | 7.49E-74 |
| 2.431682167 | 0.852380952 | 0.287736024 | 0.321939311 | 4.18E-227 |
| 1.102346563 | 0.635714286 | 0.400007663 | 0.236725705 | 1.67E-165 |
| 2.269749514 | 0.75 | 0.198355604 | 0.115717887 | 1.54E-80 |
| 1.581575684 | 0.683333333 | 0.257018303 | 0.175792715 | 2.29E-122 |
| 2.396940472 | 0.835714286 | 0.1648361 | 0.083391078 | 2.65E-58 |
| 7.220542347 | 0.907142857 | 0.224011742 | 0.185036364 | 7.30E-129 |
| 0.49537864 | 0.480952381 | 0.281737568 | 0.142804337 | 2.58E-99 |
| 0.833322606 | 0.583333333 | 0.417709495 | 0.227950118 | 3.03E-159 |
| 3.96160375 | 0.854761905 | 0.238823396 | 0.182981035 | 2.04E-127 |
| 0.539994399 | 0.507142857 | 0.388491531 | 0.158026796 | 6.31E-110 |
| 1.131106149 | 0.664285714 | 0.249016906 | 0.146085585 | 1.34E-101 |
| 1.194732157 | 0.628571429 | 0.224540902 | 0.102768962 | 1.30E-71 |
| 1.001389186 | 0.633333333 | 0.327626396 | 0.195347652 | 3.95E-136 |
| 1.140701931 | 0.592857143 | 0.211674163 | 0.09383642 | 1.81E-65 |
| 0.561041096 | 0.488095238 | 0.290105442 | 0.109479806 | 3.10E-76 |
| 1.618653677 | 0.719047619 | 0.41828067 | 0.297539764 | 2.61E-209 |
| 0.870133514 | 0.611904762 | 0.22605544 | 0.111603153 | 1.06E-77 |
| 0.162840483 | 0.24047619 | 0.571844188 | 0.096843075 | 1.56E-67 |
| 4.08190487 | 0.919047619 | 0.186966703 | 0.151109696 | 4.24E-105 |
| 0.52085529 | 0.452380952 | 0.454611944 | 0.187430314 | 1.51E-130 |
| 0.738722656 | 0.602380952 | 0.245124624 | 0.116224556 | 6.86E-81 |
| 0.6358832 | 0.530952381 | 0.244649455 | 0.135225561 | 4.76E-94 |
| 0.312477929 | 0.371428571 | 0.568096224 | 0.168764643 | 1.94E-117 |
| 0.360492129 | 0.383333333 | 0.340184941 | 0.112189228 | 4.19E-78 |
| 1.051032662 | 0.716666667 | 0.178128524 | 0.100412952 | 5.45E-70 |
| 0.366496425 | 0.388095238 | 0.364169158 | 0.121732593 | 1.07E-84 |
| 0.926081211 | 0.611904762 | 0.290799005 | 0.173108743 | 1.75E-120 |

| | | | | |
|---|---|---|---|---|
| 1.677168101 | 0.702380952 | 0.309239459 | 0.198240535 | 3.58E-138 |
| 6.145072731 | 0.916666667 | 0.137372743 | 0.091097377 | 1.38E-63 |
| 2.842145747 | 0.907142857 | 0.298828568 | 0.32526455 | 1.53E-229 |
| 2.701325982 | 0.892857143 | 0.247720926 | 0.22056265 | 5.48E-154 |
| 2.883250374 | 0.930952381 | 0.19463148 | 0.205491484 | 2.68E-143 |
| 0.93647835 | 0.635714286 | 0.399763557 | 0.235764472 | 8.13E-165 |
| 2.874855784 | 0.852380952 | 0.495243613 | 0.48658242 | 0 |
| 1.500930544 | 0.697619048 | 0.271942738 | 0.176413863 | 8.40E-123 |
| 69.06741851 | 0.997619048 | 0.162611203 | 0.302443467 | 7.08E-213 |
| 1.138055155 | 0.676190476 | 0.187326009 | 0.093693036 | 2.27E-65 |
| 16.66496047 | 0.935714286 | 0.205358742 | 0.141467094 | 2.19E-98 |
| 7.781805292 | 0.971428571 | 0.205702072 | 0.140435032 | 1.14E-97 |
| 9.692454861 | 0.966666667 | 0.232042102 | 0.159076811 | 1.17E-110 |
| 1.901391214 | 0.628571429 | 0.215050616 | 0.084941225 | 2.30E-59 |
| 3.113984865 | 0.695238095 | 0.217173235 | 0.047361021 | 1.04E-33 |
| 3.184265618 | 0.802380952 | 0.178555132 | 0.068741973 | 2.79E-48 |
| 2.588582409 | 0.771428571 | 0.171410739 | 0.06026125 | 1.71E-42 |
| 2.414794653 | 0.723809524 | 0.185839661 | 0.068923375 | 2.09E-48 |
| 2.887493309 | 0.785714286 | 0.169245834 | 0.03180473 | 3.92E-23 |
| 1.745572979 | 0.614285714 | 0.199603903 | 0.065594648 | 3.92E-46 |
| 2.719674521 | 0.754761905 | 0.174298305 | 0.030760904 | 2.01E-22 |
| 3.223620421 | 0.819047619 | 0.154795263 | 0.060777829 | 7.58E-43 |
| 7.138426727 | 0.942857143 | 0.140385402 | 0.0621288 | 9.09E-44 |
| 3.031693782 | 0.821428571 | 0.156301087 | 0.053054313 | 1.39E-37 |
| 19.20071326 | 0.983333333 | 0.180699252 | 0.061147713 | 4.24E-43 |
| 2.809982501 | 0.664285714 | 0.268399532 | 0.080850416 | 1.46E-56 |
| 8.761349713 | 0.973809524 | 0.192457701 | 0.109001834 | 6.62E-76 |
| 2.560671553 | 0.676190476 | 0.246059278 | 0.087643147 | 3.23E-61 |
| 45.55396137 | 1 | 0.248809377 | 0.424078139 | 5.04E-303 |
| 31.73510058 | 1 | 0.238990372 | 0.372652979 | 1.65E-264 |
| 2.689582574 | 0.766666667 | 0.165203629 | 0.058960462 | 1.31E-41 |
| 8.196549985 | 0.971428571 | 0.229704244 | 0.220597684 | 5.17E-154 |
| 10.50015319 | 0.992857143 | 0.198978593 | 0.176052619 | 1.51E-122 |
| 3.63351527 | 0.833333333 | 0.174712508 | 0.068541751 | 3.82E-48 |
| 1.88140741 | 0.611904762 | 0.255222586 | 0.089717773 | 1.22E-62 |
| 2.482227686 | 0.742857143 | 0.177488084 | 0.073018143 | 3.33E-51 |
| 2.144174842 | 0.695238095 | 0.173375216 | 0.05919144 | 9.15E-42 |
| 2.355064572 | 0.70952381 | 0.187638927 | 0.045527842 | 1.84E-32 |
| 49.6134915 | 1 | 0.231643138 | 0.43660268 | 1.69784655712915e-312 |
| 5.0505005 | 0.923809524 | 0.195160076 | 0.169608326 | 4.98E-118 |
| 2.472329632 | 0.769047619 | 0.172652542 | 0.055371985 | 3.66E-39 |
| 1.422155996 | 0.545238095 | 0.225768427 | 0.06418662 | 3.59E-45 |
| 2.64767197 | 0.69047619 | 0.303255902 | 0.151566025 | 2.04E-105 |
| 1.474634809 | 0.54047619 | 0.295560563 | 0.087959014 | 1.96E-61 |
| 3.102015724 | 0.821428571 | 0.186528022 | 0.072937404 | 3.79E-51 |

| | | | | |
|---|---|---|---|---|
| 2.328284738 | 0.711904762 | 0.18330724 | 0.062864539 | 2.86E-44 |
| 1.415702036 | 0.528571429 | 0.238148026 | 0.058104466 | 5.04E-41 |
| 1.921937772 | 0.647619048 | 0.179540308 | 0.060708167 | 8.46E-43 |
| 1.655509912 | 0.59047619 | 0.199148763 | 0.0994227 | 2.62E-69 |
| 3.724320995 | 0.869047619 | 0.137737512 | 0.048096983 | 3.29E-34 |
| 1.490795836 | 0.566666667 | 0.30508726 | 0.110525084 | 5.89E-77 |
| 2.351585599 | 0.657142857 | 0.24440913 | 0.072906395 | 3.98E-51 |
| 2.769455595 | 0.728571429 | 0.231799814 | 0.094124408 | 1.15E-65 |
| 41.58431835 | 1 | 0.198139906 | 0.119663907 | 2.88E-83 |
| 26.50030454 | 0.997619048 | 0.248047585 | 0.441687516 | 2.36480015503223e-316 |
| 27.26070563 | 1 | 0.232074842 | 0.320140192 | 8.65E-226 |
| 2.531475423 | 0.719047619 | 0.199111553 | 0.085934892 | 4.79E-60 |
| 19.99305003 | 1 | 0.15933606 | 0.235627567 | 1.02E-164 |
| 2.709304918 | 0.771428571 | 0.188910409 | 0.058076013 | 5.27E-41 |
| 15.78159699 | 0.997619048 | 0.192109745 | 0.243676456 | 1.80E-170 |
| 1.608122268 | 0.54047619 | 0.243303021 | 0.103906096 | 2.15E-72 |
| 0.754821089 | 0.48757764 | 0.12691694 | 0.011028119 | 9.68E-08 |
| 0.477502546 | 0.425465839 | 0.106278636 | 0.011812086 | 3.41E-08 |
| 0.103545184 | 0.152173913 | 0.75240997 | 0.107060414 | 1.15E-62 |
| 0.323217706 | 0.332298137 | 0.17156164 | 0.037432451 | 7.61E-23 |
| 0.333272324 | 0.450310559 | 0.151705288 | 0.042909573 | 5.77E-26 |
| 1.108959663 | 0.627329193 | 0.085004644 | 0.003702995 | 0.002010012 |
| 0.177918738 | 0.217391304 | 0.37206158 | 0.078203271 | 4.14E-46 |
| 0.248186087 | 0.304347826 | 0.184201596 | 0.031736087 | 1.34E-19 |
| 0.837895282 | 0.586956522 | 0.347882739 | 0.210507117 | 7.62E-123 |
| 599.0340179 | 1 | 0.674474679 | 0.928431896 | 0 |
| 0.202809006 | 0.295031056 | 0.208863362 | 0.046895656 | 3.09E-28 |
| 0.939364725 | 0.599378882 | 0.406958722 | 0.209897254 | 1.74E-122 |
| 4.045996692 | 0.959627329 | 0.053766064 | 0.074213484 | 7.93E-44 |
| 0.143535463 | 0.229813665 | 0.188996475 | 0.023549062 | 6.29E-15 |
| 5.823862709 | 0.98136646 | 0.050554387 | 0.092171925 | 4.12E-54 |
| 3.781909223 | 0.947204969 | 0.045772253 | 0.127744167 | 1.37E-74 |
| 0.81980744 | 0.562111801 | 0.333645103 | 0.172188356 | 2.16E-100 |
| 2.118599262 | 0.645962733 | 0.50247983 | 0.316113081 | 8.87E-186 |
| 0.222522623 | 0.276397516 | 0.194710288 | 0.027508133 | 3.46E-17 |
| 0.09893921 | 0.226708075 | 0.221606467 | 0.040725272 | 1.01E-24 |
| 0.265281193 | 0.400621118 | 0.260039743 | 0.096640689 | 1.12E-56 |
| 1.233369885 | 0.680124224 | 0.454469896 | 0.297177822 | 2.23E-174 |
| 0.292372186 | 0.416149068 | 0.443498496 | 0.158790678 | 1.38E-92 |
| 6.04452786 | 0.97826087 | 0.04506721 | 0.123844211 | 2.46E-72 |
| 2.489221086 | 0.664596273 | 0.566435123 | 0.347131079 | 1.44E-204 |
| 0.310245499 | 0.428571429 | 0.349837902 | 0.121844435 | 3.50E-71 |
| 0.365898075 | 0.391304348 | 0.131922744 | 0.02005207 | 6.28E-13 |
| 0.243670048 | 0.316770186 | 0.312170486 | 0.081266985 | 7.31E-48 |
| 0.039188056 | 0.068322981 | 0.711593572 | 0.048626097 | 3.19E-29 |

| | | | | |
|---|---|---|---|---|
| 0.34664381 | 0.48447205 | 0.09585093 | 0.008976133 | 1.51E-06 |
| 1.088498913 | 0.602484472 | 0.629945513 | 0.33563025 | 1.38E-197 |
| 0.107999149 | 0.161490683 | 0.592873186 | 0.083115225 | 6.39E-49 |
| 0.637284347 | 0.562111801 | 0.222290998 | 0.096421832 | 1.50E-56 |
| 2.370841149 | 0.801242236 | 0.750029492 | 0.571939214 | 0 |
| 6.424748387 | 0.98136646 | 0.046203792 | 0.105190413 | 1.37E-61 |
| 0.266425414 | 0.326086957 | 0.267954137 | 0.065889355 | 4.53E-39 |
| 3.467128654 | 0.947204969 | 0.046748573 | 0.098973645 | 5.14E-58 |
| 172.8252947 | 1 | 0.678135084 | 0.936270745 | 0 |
| 239.3292083 | 1 | 0.677549921 | 0.932040479 | 0 |
| 2.044306432 | 0.860248447 | 0.1130842 | 0.063028738 | 1.95E-37 |
| 104.4331369 | 0.99378882 | 0.661093916 | 0.795046934 | 0 |
| 0.120335333 | 0.226708075 | 0.588570665 | 0.107965303 | 3.48E-63 |
| 0.93572215 | 0.664596273 | 0.514518582 | 0.352132109 | 1.30E-207 |
| 0.292310715 | 0.394409938 | 0.122319252 | 0.021517873 | 9.10E-14 |
| 2.031932708 | 0.894409938 | 0.056885369 | 0.015260594 | 3.51E-10 |
| 0.209953539 | 0.279503106 | 0.170865636 | 0.024605491 | 1.57E-15 |
| 1.307687351 | 0.602484472 | 0.694186221 | 0.375279071 | 9.59E-222 |
| 0.376505495 | 0.459627329 | 0.132001935 | 0.049101346 | 1.71E-29 |
| 0.059390344 | 0.074534161 | 0.711147788 | 0.040163437 | 2.12E-24 |
| 12.39570471 | 0.98447205 | 0.070987261 | 0.190143251 | 6.90E-111 |
| 0.444554062 | 0.440433213 | 0.676330923 | 0.250128209 | 1.11E-131 |
| 1.376203333 | 0.714801444 | 0.555641259 | 0.402022632 | 6.90E-214 |
| 0.528200569 | 0.505415162 | 0.659050101 | 0.277265777 | 3.60E-146 |
| 4.309770589 | 0.649819495 | 0.600725251 | 0.314266936 | 4.60E-166 |
| 1.95571876 | 0.790613718 | 0.375074907 | 0.345267425 | 7.47E-183 |
| 0.759790022 | 0.570397112 | 0.486049607 | 0.256249858 | 6.10E-135 |
| 0.35053452 | 0.411552347 | 0.688020395 | 0.22179429 | 1.22E-116 |
| 0.559546285 | 0.418772563 | 0.735957767 | 0.252914477 | 3.65E-133 |
| 1.370641376 | 0.458483755 | 0.648914043 | 0.247935921 | 1.64E-130 |
| 1.184357173 | 0.592057762 | 0.488612489 | 0.279385352 | 2.63E-147 |
| 0.764896757 | 0.429602888 | 0.666814867 | 0.236306683 | 2.48E-124 |
| 1.576725761 | 0.620938628 | 0.652576525 | 0.383439548 | 1.10E-203 |
| 2.789002169 | 0.685920578 | 0.782247773 | 0.473487524 | 1.58E-253 |
| 1.210793361 | 0.667870036 | 0.581869834 | 0.39711433 | 3.45E-211 |
| 1.244340993 | 0.63898917 | 0.456724827 | 0.271559659 | 4.07E-143 |
| 0.326943335 | 0.389891697 | 0.812936292 | 0.248123226 | 1.30E-130 |
| 10.49030114 | 0.848375451 | 0.677573003 | 0.47354159 | 1.48E-253 |
| 1.93492967 | 0.736462094 | 0.843133865 | 0.565340204 | 1.98E-305 |
| 1.792799074 | 0.736462094 | 0.800933852 | 0.570017002 | 4.22E-308 |
| 0.671147082 | 0.588447653 | 0.663566358 | 0.360734704 | 2.85E-191 |
| 0.782286658 | 0.440433213 | 0.767984466 | 0.246741931 | 7.06E-130 |
| 0.80670186 | 0.555956679 | 0.819629596 | 0.398753005 | 4.34E-212 |
| 0.640272249 | 0.57400722 | 0.718515415 | 0.344413911 | 2.17E-182 |
| 0.778191 | 0.57400722 | 0.812244899 | 0.38071778 | 3.42E-202 |

| | | | | |
|---|---|---|---|---|
| 0.429117115 | 0.444043321 | 0.718815829 | 0.268053322 | 3.05E-141 |
| 0.434746597 | 0.465703971 | 0.811436369 | 0.315295761 | 1.28E-166 |
| 0.459403932 | 0.469314079 | 0.594170128 | 0.236858473 | 1.26E-124 |
| 0.672592872 | 0.530685921 | 0.65921115 | 0.340265928 | 3.89E-180 |
| 0.443680996 | 0.379061372 | 0.770635987 | 0.246187672 | 1.39E-129 |
| 0.628458643 | 0.519855596 | 0.770602248 | 0.313829121 | 7.93E-166 |
| 0.544779802 | 0.483754513 | 0.811381402 | 0.33338966 | 2.09E-176 |
| 0.514927016 | 0.483754513 | 0.683490717 | 0.261550368 | 9.07E-138 |
| 2.183524562 | 0.664259928 | 0.621440069 | 0.385781551 | 5.75E-205 |
| 2.158136415 | 0.801444043 | 0.816616118 | 0.626026154 | 0 |
| 2.590460718 | 0.667870036 | 0.45715176 | 0.268538985 | 1.68E-141 |
| 2.147644231 | 0.711191336 | 0.617799211 | 0.454608905 | 5.40E-243 |
| 0.922673438 | 0.584837545 | 0.474617128 | 0.261477736 | 9.92E-138 |
| 0.410565272 | 0.411552347 | 0.73848483 | 0.251494962 | 2.08E-132 |
| 2.995897209 | 0.833935018 | 0.825525028 | 0.64724457 | 0 |
| 0.915861229 | 0.555956679 | 0.585539039 | 0.309367898 | 2.02E-163 |
| 2.316863351 | 0.80866426 | 0.672350251 | 0.584408287 | 2.42577245053954e-316 |
| 1.255983185 | 0.660649819 | 0.683286554 | 0.444298098 | 2.93E-237 |
| 0.824836057 | 0.42599278 | 0.757559307 | 0.240506068 | 1.46E-126 |
| 0.707596614 | 0.563176895 | 0.735447081 | 0.387144105 | 1.03E-205 |
| 1.331073477 | 0.657039711 | 0.520741143 | 0.323597667 | 4.18E-171 |
| 0.981020242 | 0.635379061 | 0.776393438 | 0.430913681 | 7.76E-230 |
| 1.137917529 | 0.624548736 | 0.627794801 | 0.354594344 | 6.31E-188 |
| 1.315034729 | 0.657039711 | 0.651095737 | 0.393920556 | 1.96E-209 |
| 0.607907139 | 0.505415162 | 0.807607039 | 0.3413594 | 9.92E-181 |
| 0.920357493 | 0.59566787 | 0.776772406 | 0.388207079 | 2.69E-206 |
| 0.662761953 | 0.43071161 | 0.245503612 | 0.100873952 | 4.42E-52 |
| 6.924121015 | 0.947565543 | 0.498578558 | 0.664926117 | 0 |
| 1.815086258 | 0.74906367 | 0.154469338 | 0.092922685 | 4.44E-48 |
| 1.451322645 | 0.666666667 | 0.289268466 | 0.220551007 | 5.09E-113 |
| 0.710116795 | 0.441947566 | 0.257822302 | 0.102443054 | 7.16E-53 |
| 1.700207572 | 0.68164794 | 0.212753976 | 0.106129416 | 9.94E-55 |
| 1.573311696 | 0.696629213 | 0.269369369 | 0.193028944 | 6.99E-99 |
| 1.476323241 | 0.63670412 | 0.16426444 | 0.06247767 | 8.51E-33 |
| 1.367325358 | 0.588014981 | 0.187554875 | 0.07182834 | 1.75E-37 |
| 0.552880378 | 0.397003745 | 0.328351724 | 0.130986444 | 2.75E-67 |
| 0.719586197 | 0.415730337 | 0.282425872 | 0.128700328 | 3.96E-66 |
| 0.899713808 | 0.531835206 | 0.28949328 | 0.258111912 | 1.92E-132 |
| 0.581198031 | 0.370786517 | 0.303415751 | 0.089013499 | 4.11E-46 |
| 1.268843538 | 0.56928839 | 0.384606091 | 0.328450421 | 3.20E-169 |
| 1.58788741 | 0.65917603 | 0.441985106 | 0.413698457 | 1.56E-214 |
| 1.284949558 | 0.595505618 | 0.214592606 | 0.105580007 | 1.88E-54 |
| 0.610248749 | 0.411985019 | 0.260783895 | 0.105843968 | 1.38E-54 |
| 4.619189675 | 0.891385768 | 0.435617436 | 0.529855649 | 1.21E-277 |
| 1.487877134 | 0.610486891 | 0.458542193 | 0.39075972 | 2.93E-202 |

| | | | | |
|---|---|---|---|---|
| 1.014329075 | 0.520599251 | 0.20151354 | 0.06076943 | 6.11E-32 |
| 4.452167972 | 0.880149813 | 0.137067434 | 0.08060078 | 6.92E-42 |
| 2.918672846 | 0.823970037 | 0.169658828 | 0.068489379 | 8.26E-36 |
| 1.339529369 | 0.565543071 | 0.361365037 | 0.206860154 | 5.61E-106 |
| 0.51795671 | 0.367041199 | 0.284162461 | 0.121048317 | 2.93E-62 |
| 0.824815086 | 0.483146067 | 0.264353301 | 0.158955419 | 1.75E-81 |
| 2.621417417 | 0.801498127 | 0.30249057 | 0.312557055 | 7.29E-161 |
| 0.35280767 | 0.29588015 | 0.347848141 | 0.092090033 | 1.17E-47 |
| 3.581827369 | 0.883895131 | 0.177373676 | 0.089573946 | 2.15E-46 |
| 1.883314901 | 0.711610487 | 0.40672901 | 0.352607288 | 5.60E-182 |
| 3.193417396 | 0.838951311 | 0.47893808 | 0.494009428 | 5.39E-258 |
| 2.612371456 | 0.72659176 | 0.404690902 | 0.371691927 | 4.20E-192 |
| 3.052155469 | 0.823970037 | 0.40789539 | 0.435796828 | 2.04E-226 |
| 6.226610261 | 0.947565543 | 0.423167592 | 0.514483211 | 3.38E-269 |
| 0.843609835 | 0.494382022 | 0.245834598 | 0.125438186 | 1.77E-64 |
| 3.857648369 | 0.857677903 | 0.388110342 | 0.535784576 | 6.58E-281 |
| 2.254173937 | 0.767790262 | 0.334197815 | 0.340356717 | 1.68E-175 |
| 0.400357887 | 0.314606742 | 0.369883425 | 0.109009679 | 3.51E-56 |
| 0.694007755 | 0.43071161 | 0.456135972 | 0.219706241 | 1.38E-112 |
| 0.577226861 | 0.449438202 | 0.240693681 | 0.080761999 | 5.75E-42 |
| 14.51677274 | 0.97752809 | 0.109025174 | 7.41E-05 | 0.682230238 |
| 26.52079644 | 0.992509363 | 0.109690247 | 0.000451683 | 0.312024872 |
| 0.571546483 | 0.397003745 | 0.274308321 | 0.086873876 | 4.89E-45 |
| 3.910136578 | 0.838951311 | 0.443213422 | 0.491838593 | 8.24E-257 |
| 1.199698702 | 0.554307116 | 0.190787598 | 0.101136964 | 3.26E-52 |
| 1.443871608 | 0.711610487 | 0.154090004 | 0.073022731 | 4.40E-38 |
| 2.805223894 | 0.805243446 | 0.326127053 | 0.293910916 | 4.29E-151 |
| 4.097461835 | 0.868913858 | 0.310119841 | 0.313608346 | 2.05E-161 |
| 0.55436866 | 0.411985019 | 0.299021916 | 0.15117101 | 1.59E-77 |
| 58.16395901 | 0.992509363 | 0.142621002 | 0.065711109 | 2.04E-34 |
| 12.32493515 | 0.850187266 | 0.163346693 | 0.093030515 | 3.92E-48 |
| 2.222828363 | 0.523809524 | 0.780753 | 0.315019083 | 4.58E-146 |
| 1.20655841 | 0.653679654 | 0.788745171 | 0.41167299 | 5.25E-192 |
| 0.911653645 | 0.670995671 | 0.581122926 | 0.304519177 | 3.99E-141 |
| 0.509262838 | 0.493506494 | 0.744884213 | 0.282862341 | 5.76E-131 |
| 18.54331871 | 0.887445887 | 0.401653964 | 0.250563605 | 6.86E-116 |
| 0.671388452 | 0.54978355 | 0.758621376 | 0.302519733 | 3.47E-140 |
| 0.566153203 | 0.532467532 | 0.823556977 | 0.32427232 | 1.99E-150 |
| 0.880320604 | 0.614718615 | 0.705265467 | 0.349905118 | 1.50E-162 |
| 0.63605404 | 0.554112554 | 0.83690118 | 0.352576828 | 8.12E-164 |
| 1.551470428 | 0.731601732 | 0.672606785 | 0.428765495 | 3.21E-200 |
| 0.454403092 | 0.467532468 | 0.784485539 | 0.270785544 | 2.56E-125 |
| 0.78840881 | 0.567099567 | 0.714398736 | 0.359410338 | 4.63E-167 |
| 0.880793491 | 0.489177489 | 0.717007266 | 0.256667023 | 9.88E-119 |
| 0.70435513 | 0.601731602 | 0.680929857 | 0.343381451 | 1.85E-159 |

| | | | | |
|---|---|---|---|---|
| 4.154764885 | 0.757575758 | 0.791922252 | 0.537367427 | 4.75E-253 |
| 1.85830161 | 0.800865801 | 0.74601418 | 0.520870697 | 5.95E-245 |
| 4.75540728 | 0.714285714 | 0.757177641 | 0.469128515 | 1.01E-219 |
| 0.920041866 | 0.545454545 | 0.795476749 | 0.357263907 | 4.84E-166 |
| 0.316559053 | 0.467532468 | 0.829523866 | 0.292676824 | 1.45E-135 |
| 1.847281266 | 0.727272727 | 0.615545344 | 0.447385104 | 3.38E-209 |
| 1.652277867 | 0.692640693 | 0.710861855 | 0.429608508 | 1.26E-200 |
| 0.914777611 | 0.632034632 | 0.656059881 | 0.328933364 | 1.26E-152 |
| 1.500217382 | 0.536796537 | 0.697111699 | 0.264217862 | 2.98E-122 |
| 3.145435313 | 0.731601732 | 0.662105258 | 0.43196671 | 9.24E-202 |
| 0.426566355 | 0.575757576 | 0.780003938 | 0.327038965 | 9.85E-152 |
| 1.776598667 | 0.748917749 | 0.490132083 | 0.394858514 | 5.93E-184 |
| 0.375458127 | 0.523809524 | 0.688898202 | 0.262244802 | 2.48E-121 |
| 1.164610641 | 0.58008658 | 0.756259508 | 0.339959491 | 7.71E-158 |
| 0.434931484 | 0.510822511 | 0.723901455 | 0.296881616 | 1.54E-137 |
| 0.597229906 | 0.597402597 | 0.669650556 | 0.33468917 | 2.40E-155 |
| 0.486188086 | 0.523809524 | 0.726711978 | 0.277665015 | 1.56E-128 |
| 8.838615446 | 0.792207792 | 0.754904583 | 0.527853459 | 2.24E-248 |
| 5.721761896 | 0.748917749 | 0.766857329 | 0.516916785 | 5.15E-243 |
| 7.046420163 | 0.783549784 | 0.73373146 | 0.511513939 | 2.27E-240 |
| 0.590839397 | 0.476190476 | 0.7495784 | 0.301044529 | 1.71E-139 |
| 0.307561746 | 0.493506494 | 0.763840478 | 0.281378207 | 2.85E-130 |
| 0.295153803 | 0.528138528 | 0.690488973 | 0.321929436 | 2.54E-149 |
| 0.583935523 | 0.575757576 | 0.827635184 | 0.340607179 | 3.81E-158 |
| 1.084676524 | 0.519480519 | 0.763563616 | 0.3220979 | 2.11E-149 |
| 0.606857525 | 0.545454545 | 0.691001174 | 0.300431894 | 3.32E-139 |
| 0.36497206 | 0.489177489 | 0.720216965 | 0.273169714 | 1.97E-126 |
| 2.708288064 | 0.766233766 | 0.710971943 | 0.484351209 | 4.05E-227 |
| 1.146383637 | 0.593073593 | 0.725372888 | 0.350555937 | 7.38E-163 |
| 4.410401484 | 0.748917749 | 0.69060784 | 0.412941676 | 1.29E-192 |
| 0.716481717 | 0.532467532 | 0.720474888 | 0.285414425 | 3.68E-132 |
| 0.616542184 | 0.614718615 | 0.801216632 | 0.356714728 | 8.83E-166 |
| 0.85257274 | 0.614718615 | 0.680015882 | 0.345967091 | 1.10E-160 |
| 0.829415506 | 0.696969697 | 0.511175459 | 0.34121665 | 1.96E-158 |
| 0.959518731 | 0.632034632 | 0.776490804 | 0.390571474 | 6.64E-182 |
| 0.308181894 | 0.441558442 | 0.83951567 | 0.265853038 | 5.14E-123 |
| 0.243541325 | 0.430434783 | 0.592184459 | 0.17118292 | 5.78E-79 |
| 1.504292623 | 0.860869565 | 0.457668432 | 0.369989612 | 1.52E-171 |
| 33.24602752 | 0.986956522 | 0.337581801 | 0.382700432 | 1.40E-177 |
| 0.090703492 | 0.408695652 | 0.552872948 | 0.128432156 | 1.77E-59 |
| 0.296826842 | 0.613043478 | 0.374617036 | 0.160419319 | 4.80E-74 |
| 5.621851457 | 0.930434783 | 0.22624097 | 0.118022996 | 9.39E-55 |
| 0.301251556 | 0.582608696 | 0.487393838 | 0.244300223 | 1.27E-112 |
| 1.724565391 | 0.830434783 | 0.495021732 | 0.319804407 | 7.48E-148 |
| 0.178194532 | 0.456521739 | 0.476766625 | 0.136466355 | 3.95E-63 |

| | | | | |
|---|---|---|---|---|
| 2.032231176 | 0.830434783 | 0.396172662 | 0.315420134 | 8.57E-146 |
| 0.661693942 | 0.756521739 | 0.368715658 | 0.237927746 | 1.13E-109 |
| 0.15618921 | 0.430434783 | 0.539876344 | 0.153173464 | 9.72E-71 |
| 2.070083753 | 0.886956522 | 0.497504519 | 0.444805398 | 2.76E-207 |
| 1.323947578 | 0.760869565 | 0.381970825 | 0.303425723 | 3.61E-140 |
| 0.218004671 | 0.595652174 | 0.372829784 | 0.180416424 | 3.43E-83 |
| 0.146597624 | 0.491304348 | 0.584164171 | 0.191842732 | 1.97E-88 |
| 0.421492953 | 0.67826087 | 0.334536042 | 0.204844365 | 2.09E-94 |
| 2.121797011 | 0.860869565 | 0.48321301 | 0.372583853 | 8.95E-173 |
| 0.226745292 | 0.569565217 | 0.663398824 | 0.251342616 | 6.87E-116 |
| 0.381701049 | 0.630434783 | 0.405367992 | 0.169741984 | 2.64E-78 |
| 16.39900218 | 0.97826087 | 0.488600755 | 0.660990298 | 1.4596281880392e-313 |
| 2.201859284 | 0.926086957 | 0.257609356 | 0.193954708 | 2.12E-89 |
| 3.420856616 | 0.943478261 | 0.342319735 | 0.305980984 | 2.29E-141 |
| 0.719351248 | 0.813043478 | 0.319488858 | 0.203106135 | 1.32E-93 |
| 21.04594125 | 0.995652174 | 0.335673914 | 0.387301285 | 9.08E-180 |
| 2.885979461 | 0.904347826 | 0.276777152 | 0.254780922 | 1.74E-117 |
| 2.914303322 | 0.860869565 | 0.48435507 | 0.364096429 | 9.47E-169 |
| 0.874788516 | 0.747826087 | 0.299911326 | 0.18393528 | 8.37E-85 |
| 3.316190026 | 0.939130435 | 0.270407422 | 0.216251494 | 1.16E-99 |
| 0.944909847 | 0.82173913 | 0.392539235 | 0.28657384 | 2.75E-132 |
| 1.489340129 | 0.847826087 | 0.303005785 | 0.255978721 | 4.84E-118 |
| 0.215625626 | 0.569565217 | 0.477547994 | 0.171361487 | 4.79E-79 |
| 0.587260474 | 0.795652174 | 0.276846518 | 0.180486113 | 3.19E-83 |
| 2.509576936 | 0.9 | 0.247209809 | 0.18389517 | 8.73E-85 |
| 1.187097249 | 0.865217391 | 0.275672149 | 0.208143602 | 6.33E-96 |
| 1.793959647 | 0.82173913 | 0.365149312 | 0.262158483 | 6.52E-121 |
| 0.449412554 | 0.72173913 | 0.304484334 | 0.17363908 | 4.35E-80 |
| 0.615508767 | 0.808695652 | 0.260789888 | 0.190403806 | 9.03E-88 |
| 1.424044634 | 0.847826087 | 0.395981361 | 0.273635789 | 2.97E-126 |
| 0.598989868 | 0.713043478 | 0.301013138 | 0.175043842 | 9.90E-81 |
| 1.03452945 | 0.852173913 | 0.396001791 | 0.360400396 | 5.33E-167 |
| 1.344993314 | 0.830434783 | 0.258685733 | 0.154722623 | 1.91E-71 |
| 2.623182946 | 0.939130435 | 0.253726489 | 0.188617947 | 5.96E-87 |
| 0.630212475 | 0.734782609 | 0.413569277 | 0.220878407 | 8.55E-102 |
| 23.97137945 | 0.995652174 | 0.320652128 | 0.466943132 | 5.83E-218 |
| 11.70351792 | 0.986956522 | 0.224156934 | 0.221342274 | 5.22E-102 |
| 1.115231778 | 0.87826087 | 0.350175265 | 0.25181047 | 4.17E-116 |
| 0.748141805 | 0.504347826 | 0.596967438 | 0.16856618 | 9.10E-78 |
| 4.029111312 | 0.943478261 | 0.223812419 | 0.168262984 | 1.25E-77 |
| 0.613353253 | 0.765217391 | 0.291806854 | 0.188175867 | 9.50E-87 |
| 3.297643818 | 0.862068966 | 0.72383965 | 0.603603181 | 2.71E-230 |
| 2.755855366 | 0.885057471 | 0.657489965 | 0.538978715 | 8.04E-205 |
| 7.702609371 | 0.908045977 | 0.742458115 | 0.636603794 | 2.10E-243 |
| 6.800816038 | 0.948275862 | 0.558597643 | 0.657235778 | 1.23E-251 |

| | | | | |
|---|---|---|---|---|
| 2.424878163 | 0.856321839 | 0.732936812 | 0.547695016 | 3.05E-208 |
| 4.41604235 | 0.856321839 | 0.772798421 | 0.59779135 | 5.42E-228 |
| 12.69780054 | 0.971264368 | 0.513403803 | 0.698133602 | 4.93E-268 |
| 8.542407613 | 0.931034483 | 0.629123799 | 0.510843572 | 8.20E-194 |
| 3.107932848 | 0.862068966 | 0.737299224 | 0.561650077 | 9.97E-214 |
| 6.831231758 | 0.936781609 | 0.609802805 | 0.599042587 | 1.73E-228 |
| 1.432976006 | 0.787356322 | 0.728076643 | 0.48390993 | 2.55E-183 |
| 1.719941744 | 0.793103448 | 0.698962367 | 0.478137464 | 4.46E-181 |
| 2.008082116 | 0.775862069 | 0.675326227 | 0.45957339 | 7.08E-174 |
| 1.425712455 | 0.597701149 | 0.835233605 | 0.407834759 | 6.43E-154 |
| 0.855799454 | 0.643678161 | 0.749197371 | 0.37425396 | 4.75E-141 |
| 1.471915445 | 0.827586207 | 0.705674422 | 0.490740659 | 5.62E-186 |
| 3.864758715 | 0.91954023 | 0.56035747 | 0.616078257 | 3.05E-235 |
| 1.164146674 | 0.764367816 | 0.681910385 | 0.42497748 | 1.63E-160 |
| 2.681042086 | 0.83908046 | 0.661020877 | 0.486808437 | 1.90E-184 |
| 3.330873547 | 0.91954023 | 0.548017443 | 0.578339714 | 2.64E-220 |
| 4.555351452 | 0.867816092 | 0.672119955 | 0.510114795 | 1.58E-193 |
| 64.04631261 | 0.994252874 | 0.685330468 | 0.86568832 | 0 |
| 0.723951948 | 0.666666667 | 0.727581651 | 0.386101172 | 1.39E-145 |
| 3.859575245 | 0.879310345 | 0.650456296 | 0.577215833 | 7.33E-220 |
| 5.759189102 | 0.91954023 | 0.753786364 | 0.645124773 | 8.45E-247 |
| 1.716231882 | 0.850574713 | 0.563839006 | 0.437454876 | 2.52E-165 |
| 11.0764378 | 0.948275862 | 0.722489172 | 0.712223145 | 1.04E-273 |
| 3.458911885 | 0.804597701 | 0.608980426 | 0.398781828 | 1.92E-150 |
| 2.567378417 | 0.844827586 | 0.732821655 | 0.530514348 | 1.67E-201 |
| 3.14290345 | 0.856321839 | 0.608621303 | 0.50757531 | 1.55E-192 |
| 14.94558551 | 0.948275862 | 0.76507458 | 0.749397554 | 9.61E-289 |
| 8.822709976 | 0.936781609 | 0.597349679 | 0.699401855 | 1.52E-268 |
| 3.093131437 | 0.885057471 | 0.714022421 | 0.549416063 | 6.44E-209 |
| 5.604432349 | 0.862068966 | 0.708224181 | 0.522957258 | 1.51E-198 |
| 2.294108774 | 0.787356322 | 0.697205204 | 0.469706127 | 8.36E-178 |
| 6.430666233 | 0.879310345 | 0.744321728 | 0.614640139 | 1.14E-234 |
| 1.223726117 | 0.75862069 | 0.702062529 | 0.446118889 | 1.14E-168 |
| 4.569751118 | 0.936781609 | 0.549150508 | 0.651173724 | 3.26E-249 |
| 2.460366346 | 0.83908046 | 0.688147084 | 0.537690764 | 2.57E-204 |
| 3.742426098 | 0.862068966 | 0.755063248 | 0.603694134 | 2.49E-230 |
| 1.63050767 | 0.735632184 | 0.701198376 | 0.418459909 | 5.28E-158 |
| 3.115612702 | 0.810344828 | 0.706525105 | 0.475948086 | 3.16E-180 |
| 0.94690041 | 0.724137931 | 0.80735158 | 0.464703697 | 7.28E-176 |
| 2.337298261 | 0.752873563 | 0.795612363 | 0.506237069 | 5.14E-192 |
| 2.015996893 | 0.781609195 | 0.664894021 | 0.465927411 | 2.44E-176 |
| 7.287798943 | 0.902298851 | 0.736473223 | 0.640406866 | 6.42E-245 |
| 3.60552085 | 0.804597701 | 0.707137258 | 0.480779279 | 4.20E-182 |
| 7.069827266 | 0.908045977 | 0.583045393 | 0.572110865 | 7.56E-218 |
| 1.612647061 | 0.764367816 | 0.800953173 | 0.526477421 | 6.35E-200 |

| | | | | | |
|---|---|---|---|---|---|
| 1.115913763 | 0.695402299 | 0.706530744 | 0.389659966 | | 6.04E-147 |
| 1.932210257 | 0.911111111 | 0.242039639 | 0.164016648 | | 2.88E-22 |
| 0.97068704 | 0.888888889 | 0.302218015 | 0.18759624 | | 2.99E-25 |
| 2.897069386 | 0.911111111 | 0.253127354 | 0.272403432 | | 5.50E-36 |
| 1.810981592 | 0.911111111 | 0.345490694 | 0.247659619 | | 7.49E-33 |
| 0.147441631 | 0.488888889 | 0.468557287 | 0.120270338 | | 9.92E-17 |
| 0.734491902 | 0.822222222 | 0.33223679 | 0.219948098 | | 2.41E-29 |
| 0.928764218 | 0.866666667 | 0.313677345 | 0.211415352 | | 2.90E-28 |
| 0.69540587 | 0.755555556 | 0.296966792 | 0.144343472 | | 8.88E-20 |
| 0.228413063 | 0.555555556 | 0.399201469 | 0.111079207 | | 1.45E-15 |
| 1.434514656 | 0.866666667 | 0.251206405 | 0.175397057 | | 1.04E-23 |
| 0.770315971 | 0.777777778 | 0.297396658 | 0.168676056 | | 7.40E-23 |
| 1.770523764 | 0.911111111 | 0.491576602 | 0.437824157 | | 5.24E-57 |
| 1.372865444 | 0.866666667 | 0.409093952 | 0.357250862 | | 9.43E-47 |
| 1.274172484 | 0.8 | 0.505398669 | 0.328578127 | | 4.13E-43 |
| 1.319865091 | 0.8 | 0.381967793 | 0.286326502 | | 9.46E-38 |
| 0.33127572 | 0.688888889 | 0.487390723 | 0.188668804 | | 2.19E-25 |
| 0.534848309 | 0.733333333 | 0.337045732 | 0.142766745 | | 1.41E-19 |
| 0.31619562 | 0.711111111 | 0.363047436 | 0.158964153 | | 1.25E-21 |
| 0.802632424 | 0.8 | 0.32103195 | 0.17833454 | | 4.44E-24 |
| 1.887765706 | 0.866666667 | 0.352546787 | 0.223631895 | | 8.24E-30 |
| 3.545900362 | 0.933333333 | 0.458513509 | 0.440399094 | | 2.46E-57 |
| 2.378348498 | 0.888888889 | 0.261351507 | 0.241775446 | | 4.16E-32 |
| 0.66327609 | 0.755555556 | 0.317962936 | 0.16609789 | | 1.57E-22 |
| 0.381914112 | 0.755555556 | 0.30168265 | 0.150824736 | | 1.34E-20 |
| 1.093333556 | 0.844444444 | 0.35965486 | 0.289976059 | | 3.26E-38 |
| 0.575623074 | 0.755555556 | 0.303449777 | 0.137792436 | | 5.99E-19 |
| 2.860115388 | 0.911111111 | 0.323240159 | 0.285920401 | | 1.06E-37 |
| 1.148330018 | 0.933333333 | 0.395500995 | 0.257469229 | | 4.29E-34 |
| 0.186034016 | 0.555555556 | 0.519345128 | 0.131028588 | | 4.30E-18 |
| 0.192076096 | 0.6 | 0.375434707 | 0.125847102 | | 1.95E-17 |
| 1.297106883 | 0.844444444 | 0.305128541 | 0.235143324 | | 2.88E-31 |
| 0.896604373 | 0.844444444 | 0.301640511 | 0.201573655 | | 5.10E-27 |
| 3.131635929 | 0.866666667 | 0.322660104 | 0.212607225 | | 2.05E-28 |
| 0.303576803 | 0.688888889 | 0.370198365 | 0.152364514 | | 8.57E-21 |
| 1.835332367 | 0.888888889 | 0.308271805 | 0.277182203 | | 1.36E-36 |
| 0.158863337 | 0.533333333 | 0.421453581 | 0.116770418 | | 2.75E-16 |
| 2.34743716 | 0.888888889 | 0.332835658 | 0.286820654 | | 8.19E-38 |
| 3.061586771 | 0.888888889 | 0.24556367 | 0.211850457 | | 2.55E-28 |
| 3.805532728 | 0.888888889 | 0.384017078 | 0.378186089 | | 2.06E-49 |
| 2.293201832 | 0.911111111 | 0.338619502 | 0.30788299 | | 1.75E-40 |
| 0.420408103 | 0.755555556 | 0.30869801 | 0.169980093 | | 5.06E-23 |
| 1.185995334 | 0.777777778 | 0.342713589 | 0.175942123 | | 8.91E-24 |
| 0.409279715 | 0.777777778 | 0.282945021 | 0.140611003 | | 2.63E-19 |
| 2.278211885 | 0.866666667 | 0.291711638 | 0.270923838 | | 8.47E-36 |

| | | | | |
|---|---|---|---|---|
| 0.588086021 | 0.711111111 | 0.398574874 | 0.179739057 | 2.95E-24 |
| 0.744767383 | 0.8 | 0.371718823 | 0.241216954 | 4.90E-32 |
| 2.390932666 | 0.822222222 | 0.288806362 | 0.191415881 | 9.83E-26 |
| 2.937331436 | 0.933333333 | 0.241034599 | 0.203070339 | 3.30E-27 |
| 0.811777398 | 0.8 | 0.400979391 | 0.243522708 | 2.50E-32 |
| 0.425396978 | 0.644444444 | 0.581434381 | 0.212676753 | 2.01E-28 |
| 0.396384641 | 0.527777778 | 0.454037422 | 0.159045601 | 3.07E-18 |
| 9.162648085 | 0.916666667 | 0.727427492 | 0.616602159 | 1.54E-66 |
| 1.451395508 | 0.694444444 | 0.365321525 | 0.221449125 | 8.40E-25 |
| 15.27355958 | 0.833333333 | 0.886128735 | 0.648548708 | 6.14E-70 |
| 0.540061172 | 0.5 | 0.470109201 | 0.171752024 | 1.41E-19 |
| 0.213517878 | 0.333333333 | 0.914903251 | 0.191795516 | 1.10E-21 |
| 5.999193068 | 0.888888889 | 0.28488143 | 0.281979029 | 3.62E-31 |
| 4.869742379 | 0.722222222 | 0.647019559 | 0.345496082 | 7.44E-38 |
| 60.56476746 | 0.944444444 | 0.868500043 | 0.784595707 | 1.85E-84 |
| 0.231326834 | 0.305555556 | 0.89363327 | 0.163939811 | 9.38E-19 |
| 56.41005011 | 1 | 0.883511318 | 0.919350271 | 6.42E-99 |
| 2.305146297 | 0.777777778 | 0.510951694 | 0.300713462 | 3.86E-33 |
| 0.597356065 | 0.5 | 0.830872003 | 0.24764834 | 1.48E-27 |
| 1.624319685 | 0.722222222 | 0.370963732 | 0.204082517 | 5.62E-23 |
| 33.58592619 | 0.888888889 | 0.917834599 | 0.743888875 | 4.18E-80 |
| 18.66777496 | 0.722222222 | 0.866940369 | 0.465154911 | 1.75E-50 |
| 0.547595153 | 0.527777778 | 0.762618003 | 0.262403668 | 4.14E-29 |
| 0.432091686 | 0.388888889 | 0.86287618 | 0.210273012 | 1.26E-23 |
| 0.539334108 | 0.472222222 | 0.93563423 | 0.338910496 | 3.67E-37 |
| 0.531006078 | 0.527777778 | 0.442372154 | 0.154641701 | 8.92E-18 |
| 1.433179666 | 0.75 | 0.659184166 | 0.411647611 | 7.86E-45 |
| 1.261420452 | 0.666666667 | 0.476198208 | 0.292908114 | 2.56E-32 |
| 1.874771583 | 0.75 | 0.34660616 | 0.176361191 | 4.63E-20 |
| 0.23799128 | 0.361111111 | 0.640427904 | 0.139419431 | 3.58E-16 |
| 0.503923743 | 0.5 | 0.441606057 | 0.113434899 | 1.96E-13 |
| 0.614240488 | 0.583333333 | 0.619240031 | 0.232179348 | 6.25E-26 |
| 0.495990378 | 0.5 | 0.73189143 | 0.245997392 | 2.20E-27 |
| 0.36139369 | 0.527777778 | 0.692065705 | 0.244589864 | 3.10E-27 |
| 0.193309489 | 0.25 | 0.875055526 | 0.114241086 | 1.61E-13 |
| 0.948774653 | 0.694444444 | 0.864323359 | 0.485438708 | 1.25E-52 |
| 222.0699575 | 0.916666667 | 0.88975055 | 0.750685959 | 7.84E-81 |
| 0.319807407 | 0.333333333 | 0.652065161 | 0.142285868 | 1.78E-16 |
| 0.412997734 | 0.5 | 0.834470315 | 0.278432022 | 8.54E-31 |
| 0.724304545 | 0.611111111 | 0.859131955 | 0.389891615 | 1.55E-42 |
| 0.387211458 | 0.5 | 0.512009221 | 0.174537171 | 7.20E-20 |
| 1.455845317 | 0.75 | 0.337182181 | 0.185332836 | 5.27E-21 |
| 0.228383759 | 0.333333333 | 0.728398144 | 0.154861272 | 8.46E-18 |
| 0.964589424 | 0.583333333 | 0.850372389 | 0.322573685 | 1.93E-35 |
| 90.89688133 | 1 | 0.893426925 | 0.891152182 | 6.92E-96 |

| | | | | |
|---|---|---|---|---|
| 2.159666832 | 0.722222222 | 0.753331357 | 0.453347883 | 3.10E-49 |
| 2.279562281 | 0.416666667 | 0.670996874 | 0.131036192 | 2.73E-15 |
| 0.367282786 | 0.5 | 0.465896556 | 0.160416221 | 2.20E-18 |
| 1.343361544 | 0.75 | 0.512683355 | 0.305200716 | 1.30E-33 |
| 0.223997454 | 0.333333333 | 0.892186224 | 0.229124347 | 1.31E-25 |
| 0.676147601 | 0.527777778 | 0.889275816 | 0.382865769 | 8.56E-42 |
| 2.1096411 | 0.805555556 | 0.315855146 | 0.238885696 | 1.23E-26 |
| 1.092533658 | 0.583333333 | 0.485434116 | 0.221572179 | 8.15E-25 |
| 0.154774773 | 0.305555556 | 0.804323549 | 0.189465896 | 1.94E-21 |
| 0.556132607 | 0.555555556 | 0.667858005 | 0.25144537 | 5.89E-28 |
| 1.131713503 | 0.694444444 | 0.74692507 | 0.392494445 | 8.25E-43 |
| 12.40946869 | 0.974443528 | 0.159926085 | 0.215341604 | 4.20E-278 |
| 2.660126333 | 0.68920033 | 0.213137052 | 0.133532752 | 1.49E-169 |
| 1.054852794 | 0.403957131 | 0.258352497 | 0.086024043 | 1.28E-108 |
| 1.374372359 | 0.483924155 | 0.299011463 | 0.134067112 | 3.02E-170 |
| 3.384023132 | 0.774113768 | 0.259930456 | 0.15056877 | 9.06E-192 |
| 2.390014177 | 0.709810387 | 0.157975087 | 0.028170516 | 2.81E-36 |
| 2.292326093 | 0.588623248 | 0.212784353 | 0.092446043 | 9.04E-117 |
| 12.041462 | 0.97856554 | 0.126527689 | 0.012109932 | 1.87E-16 |
| 5.256506266 | 0.916735367 | 0.130364833 | 0.010854539 | 6.68E-15 |
| 1.017710172 | 0.401483924 | 0.329231282 | 0.131393596 | 8.89E-167 |
| 1.434244345 | 0.459192086 | 0.258601091 | 0.090598455 | 2.01E-114 |
| 1.844371507 | 0.593569662 | 0.2185362 | 0.08949601 | 5.06E-113 |
| 17.8222509 | 0.991755977 | 0.121842595 | 0.01950604 | 1.39E-25 |
| 12.23782908 | 0.988458368 | 0.115637294 | 0.01507615 | 4.07E-20 |
| 2.502996877 | 0.71063479 | 0.191199316 | 0.101067354 | 9.37E-128 |
| 2.580665202 | 0.648804617 | 0.351418534 | 0.256002044 | 0 |
| 7.854820864 | 0.930750206 | 0.246518204 | 0.305847199 | 0 |
| 7.344960279 | 0.936521022 | 0.151077232 | 0.16811575 | 7.33E-215 |
| 2.441904494 | 0.613355317 | 0.274310747 | 0.201229452 | 4.92E-259 |
| 6.170239119 | 0.90354493 | 0.259026242 | 0.317625276 | 0 |
| 4.988754701 | 0.924154988 | 0.13961486 | 0.022476359 | 3.02E-29 |
| 2.357292504 | 0.65704864 | 0.187773472 | 0.054187308 | 1.40E-68 |
| 3.753874142 | 0.77493817 | 0.205328332 | 0.158796132 | 1.43E-202 |
| 30.48375752 | 1 | 0.160139894 | 0.169804961 | 4.27E-217 |
| 3.345643161 | 0.755976917 | 0.268921343 | 0.18582973 | 2.15E-238 |
| 5.316547841 | 0.882934872 | 0.302565113 | 0.323974059 | 0 |
| 3.134420277 | 0.755152514 | 0.16025643 | 0.087840316 | 6.38E-111 |
| 7.397665854 | 0.906842539 | 0.320817097 | 0.348928374 | 0 |
| 5.047898321 | 0.851607585 | 0.314013751 | 0.320903531 | 0 |
| 3.184221729 | 0.765869744 | 0.142816616 | 0.036130058 | 3.99E-46 |
| 2.312303028 | 0.63561418 | 0.182757487 | 0.087538602 | 1.54E-110 |
| 1.231691983 | 0.450948063 | 0.26096057 | 0.079126129 | 6.84E-100 |
| 40.72710021 | 1 | 0.195039509 | 0.293203332 | 0 |
| 1.347341651 | 0.474031327 | 0.253464394 | 0.102520044 | 1.31E-129 |

| | | | | |
|---|---|---|---|---|
| 14.11454215 | 0.990107172 | 0.121350235 | 0.068919053 | 4.95E-87 |
| 60.14988901 | 1 | 0.122317996 | 0.130057437 | 4.80E-165 |
| 21.87718754 | 0.999175598 | 0.164661936 | 0.251293456 | 0 |
| 24.94988824 | 1 | 0.165214395 | 0.250842689 | 0 |
| 7.109884007 | 0.949711459 | 0.145075348 | 0.14192158 | 1.81E-180 |
| 4.969755846 | 0.883759275 | 0.161724173 | 0.083851372 | 7.24E-106 |
| 21.62923576 | 0.997526793 | 0.163223478 | 0.258283327 | 0 |
| 4.552564773 | 0.865622424 | 0.16479085 | 0.095152594 | 3.26E-120 |
| 3.473242623 | 0.80708986 | 0.155865067 | 0.069234057 | 1.99E-87 |
| 42.45152371 | 1 | 0.157439418 | 0.25704166 | 0 |
| 9.649944773 | 0.974443528 | 0.114918736 | 0.009927992 | 9.35E-14 |
| 9.64781867 | 0.964550701 | 0.264463315 | 0.422617502 | 0 |
| 13.5314925 | 0.995053586 | 0.136298821 | 0.044759954 | 7.77E-57 |
| 4.628865927 | 0.866446826 | 0.170623932 | 0.141447344 | 7.53E-180 |
| 2.540721573 | 0.671063479 | 0.157771216 | 0.066557708 | 4.58E-84 |
| 7.399696424 | 0.936521022 | 0.161313545 | 0.120771077 | 4.86E-153 |
| 4.831892734 | 0.83214649 | 0.379610593 | 0.496345394 | 0 |
| 1.452782704 | 0.625635809 | 0.404556917 | 0.357174963 | 0 |
| 3.9803242 | 0.849440488 | 0.222102648 | 0.224856472 | 1.93E-261 |
| 3.836115622 | 0.82197355 | 0.252069448 | 0.238990248 | 1.08E-278 |
| 13.94670944 | 0.983723296 | 0.17478566 | 0.230742703 | 1.30E-268 |
| 0.459789126 | 0.321464903 | 0.450880443 | 0.182869318 | 6.91E-211 |
| 1.813835819 | 0.667344863 | 0.271852549 | 0.220596762 | 2.89E-256 |
| 7.460959352 | 0.854526958 | 0.280200543 | 0.26100668 | 8.20E-306 |
| 14.38652177 | 0.98982706 | 0.14431975 | 0.135534865 | 4.20E-155 |
| 1.154992653 | 0.563580875 | 0.234008917 | 0.139319576 | 1.62E-159 |
| 0.849094232 | 0.480162767 | 0.36855437 | 0.246351983 | 9.94E-288 |
| 0.835726719 | 0.424211597 | 0.385380376 | 0.200416537 | 6.88E-232 |
| 1.489811542 | 0.585961343 | 0.220005405 | 0.143154174 | 5.34E-164 |
| 1.923669638 | 0.696846389 | 0.239664511 | 0.182354752 | 2.84E-210 |
| 0.639636784 | 0.390640895 | 0.404054504 | 0.209973439 | 2.11E-243 |
| 1.978763989 | 0.708036623 | 0.205140263 | 0.180645748 | 3.08E-208 |
| 1.402700878 | 0.631739573 | 0.20403975 | 0.116669521 | 3.26E-133 |
| 5.804363631 | 0.858596134 | 0.405671414 | 0.432927918 | 0 |
| 2.426489486 | 0.765005086 | 0.282046063 | 0.266882727 | 4.21724390308045e-313 |
| 3.557884638 | 0.830111902 | 0.222359625 | 0.186148593 | 8.47E-215 |
| 9.380775825 | 0.961342828 | 0.181995645 | 0.217202266 | 3.79E-252 |
| 14.5509711 | 0.940996948 | 0.381853176 | 0.498068951 | 0 |
| 1.106134399 | 0.561546287 | 0.253220865 | 0.147695013 | 2.58E-169 |
| 3.429017551 | 0.77110885 | 0.277340534 | 0.256913068 | 9.52E-301 |
| 1.002191336 | 0.498474059 | 0.321901246 | 0.193535428 | 1.24E-223 |
| 6.095494025 | 0.869786368 | 0.331098924 | 0.387659222 | 0 |
| 2.051497632 | 0.687690743 | 0.392339491 | 0.349817531 | 0 |
| 1.988200918 | 0.634791455 | 0.330282539 | 0.234064348 | 1.14E-272 |
| 0.91195914 | 0.481180061 | 0.305088109 | 0.211085205 | 9.56E-245 |

| | | | | |
|---|---|---|---|---|
| 1.168756912 | 0.545269583 | 0.283170059 | 0.20171739 | 1.88E-233 |
| 1.912330726 | 0.652085453 | 0.312940016 | 0.24989828 | 4.30E-292 |
| 0.747501797 | 0.393692777 | 0.329928194 | 0.183414476 | 1.55E-211 |
| 2.169985648 | 0.722278739 | 0.263827883 | 0.278495175 | 0 |
| 9.193638821 | 0.939979654 | 0.345977882 | 0.46455729 | 0 |
| 3.40533823 | 0.778229908 | 0.311504348 | 0.326822282 | 0 |
| 1.715742735 | 0.620549339 | 0.397872819 | 0.332898684 | 0 |
| 2.08642902 | 0.761953204 | 0.190091266 | 0.101346713 | 1.43E-115 |
| 1.730161801 | 0.689725331 | 0.189066344 | 0.112046569 | 7.06E-128 |
| 8.504351333 | 0.96439471 | 0.146136428 | 0.147106668 | 1.26E-168 |
| 0.807594973 | 0.446592065 | 0.31499377 | 0.17415869 | 1.58E-200 |
| 4.436480833 | 0.912512716 | 0.146748657 | 0.08236679 | 6.91E-94 |
| 0.643823342 | 0.375381485 | 0.380233727 | 0.186618236 | 2.33E-215 |
| 3.904415473 | 0.886063072 | 0.149869177 | 0.108504221 | 8.50E-124 |
| 1.196373587 | 0.560528993 | 0.353556951 | 0.271078914 | 2.55227395722546e-318 |
| 2.493326253 | 0.673448627 | 0.420188429 | 0.353590486 | 0 |
| 10.89380849 | 0.951169888 | 0.212334521 | 0.29136701 | 0 |
| 2.396719249 | 0.778229908 | 0.226315523 | 0.243097714 | 9.89E-284 |
| 0.985035237 | 0.512716175 | 0.284920796 | 0.181965753 | 8.24E-210 |
| 1.305668868 | 0.482197355 | 0.369903316 | 0.231884225 | 5.27E-270 |
| 3.167191559 | 0.729399797 | 0.486381235 | 0.440861947 | 0 |
| 0.8026098 | 0.58056872 | 0.441100158 | 0.333914961 | 0 |
| 11.93369215 | 0.920616114 | 0.286324639 | 0.365251396 | 0 |
| 0.584154746 | 0.414691943 | 0.639870354 | 0.28161711 | 1.10E-304 |
| 0.832218853 | 0.623222749 | 0.482429833 | 0.334731737 | 0 |
| 1.998529631 | 0.678909953 | 0.492456171 | 0.368427642 | 0 |
| 0.426293643 | 0.43957346 | 0.552146781 | 0.264771439 | 1.69E-285 |
| 4.223802632 | 0.795023697 | 0.338133771 | 0.239948206 | 1.70E-257 |
| 1.777240436 | 0.792654028 | 0.346248222 | 0.360273602 | 0 |
| 2.095896409 | 0.816350711 | 0.697114127 | 0.615996069 | 0 |
| 0.431483849 | 0.394549763 | 0.717397498 | 0.291590739 | 4.09926790064067e-316 |
| 2.802192798 | 0.805687204 | 0.293431364 | 0.296264547 | 1.72922976044436e-321 |
| 2.259760614 | 0.806872038 | 0.674979592 | 0.580683985 | 0 |
| 0.259089758 | 0.321090047 | 0.705781868 | 0.213694825 | 3.20E-228 |
| 4.916167569 | 0.913507109 | 0.46484787 | 0.577867167 | 0 |
| 0.560629786 | 0.462085308 | 0.575214383 | 0.267695627 | 8.11E-289 |
| 0.538535516 | 0.522511848 | 0.680744239 | 0.360947799 | 0 |
| 0.781784728 | 0.524881517 | 0.562841844 | 0.292963624 | 1.08439108959303e-317 |
| 3.101664261 | 0.873222749 | 0.267960686 | 0.251171444 | 4.07E-270 |
| 0.80903243 | 0.597156398 | 0.665536815 | 0.404975752 | 0 |
| 0.845968455 | 0.523696682 | 0.682292789 | 0.35380848 | 0 |
| 3.833962303 | 0.856635071 | 0.675923481 | 0.62868777 | 0 |
| 1.423244118 | 0.757109005 | 0.323785491 | 0.273430518 | 2.43E-295 |
| 1.370976701 | 0.700236967 | 0.410534157 | 0.322838825 | 0 |
| 2.224603259 | 0.797393365 | 0.334455715 | 0.295374326 | 1.83100728348766e-320 |

| | | | | |
|---|---|---|---|---|
| 0.610405585 | 0.517772512 | 0.468970387 | 0.275482611 | 1.11E-297 |
| 1.22288371 | 0.567535545 | 0.765815442 | 0.436132137 | 0 |
| 0.309993355 | 0.311611374 | 0.724819479 | 0.208110223 | 4.87E-222 |
| 0.533718101 | 0.476303318 | 0.669096598 | 0.32379627 | 0 |
| 1.389482776 | 0.688388626 | 0.601939922 | 0.457033378 | 0 |
| 1.073185741 | 0.680094787 | 0.376064096 | 0.312151412 | 0 |
| 1.093723677 | 0.637440758 | 0.357140946 | 0.274952837 | 4.47E-297 |
| 0.519632253 | 0.476303318 | 0.492993334 | 0.258693658 | 1.30E-278 |
| 1.463240698 | 0.687203791 | 0.620759924 | 0.43891096 | 0 |
| 0.864597333 | 0.552132701 | 0.433685198 | 0.279550968 | 2.53E-302 |
| 0.674934819 | 0.517772512 | 0.73219025 | 0.361952901 | 0 |
| 2.247538994 | 0.787914692 | 0.444007596 | 0.358122538 | 0 |
| 0.383075803 | 0.362559242 | 0.665917949 | 0.240312558 | 6.64E-258 |
| 0.767660844 | 0.562796209 | 0.612941078 | 0.374988061 | 0 |
| 2.825785077 | 0.80450237 | 0.282859862 | 0.224571637 | 2.62E-240 |
| 0.637775892 | 0.533175355 | 0.438382483 | 0.268811396 | 4.37E-290 |
| 0.368934073 | 0.408767773 | 0.657591685 | 0.281816759 | 6.52E-305 |
| 0.533490453 | 0.491706161 | 0.705518362 | 0.359523267 | 0 |
| 0.787935369 | 0.604265403 | 0.627024362 | 0.395937977 | 0 |
| 1.371742201 | 0.725118483 | 0.556584228 | 0.425742153 | 0 |
| 0.769324529 | 0.409952607 | 0.711444495 | 0.280604069 | 1.59E-303 |
| 0.727007625 | 0.584123223 | 0.728350634 | 0.42149356 | 0 |
| 0.510521021 | 0.414691943 | 0.70025753 | 0.288018428 | 5.16126189727191e-312 |
| 1.523042057 | 0.549763033 | 0.649411934 | 0.332753498 | 0 |
| 2.468832298 | 0.848341232 | 0.427272978 | 0.481930478 | 0 |
| 0.926510045 | 0.590047393 | 0.387826293 | 0.266499768 | 1.85E-287 |
| 0.534213995 | 0.276315789 | 0.564696138 | 0.14729184 | 5.77E-119 |
| 3.923195155 | 0.885338346 | 0.11647887 | 0.00150832 | 0.020382531 |
| 0.742986491 | 0.411654135 | 0.235757586 | 0.085799642 | 2.05E-69 |
| 4.481286859 | 0.917293233 | 0.11021194 | 0.007085066 | 4.94E-07 |
| 0.635440779 | 0.313909774 | 0.516595814 | 0.135501308 | 2.16E-109 |
| 0.971489506 | 0.385338346 | 0.306415534 | 0.108355541 | 1.75E-87 |
| 3.984646012 | 0.612781955 | 0.758543263 | 0.408447982 | 0 |
| 1.68277089 | 0.411654135 | 0.619601588 | 0.280839291 | 4.93E-230 |
| 2.373486117 | 0.631578947 | 0.280924831 | 0.126677292 | 3.00E-102 |
| 0.568054988 | 0.268796992 | 0.63673906 | 0.173556822 | 2.01E-140 |
| 11.1587455 | 0.981203008 | 0.119320123 | 0.002533724 | 0.0026457 |
| 5.166283024 | 0.936090226 | 0.128641599 | 0.006859585 | 7.50E-07 |
| 14.28292829 | 0.996240602 | 0.102662552 | 0.000662517 | 0.1242996 |
| 13.18427137 | 0.996240602 | 0.122477727 | 0.015310771 | 1.39E-13 |
| 5.937093031 | 0.902255639 | 0.199514391 | 0.11983531 | 1.00E-96 |
| 1.328147171 | 0.5 | 0.513423371 | 0.264970105 | 1.42E-216 |
| 1.6830226 | 0.609022556 | 0.427428419 | 0.326474808 | 3.72E-269 |
| 0.360999262 | 0.191729323 | 0.590712083 | 0.092441424 | 1.01E-74 |
| 1.456359125 | 0.541353383 | 0.267725492 | 0.180639941 | 3.05E-146 |

| | | | | |
|---|---|---|---|---|
| 5.993662794 | 0.964285714 | 0.129784992 | 0.015365263 | 1.26E-13 |
| 2.147624014 | 0.437969925 | 0.370553112 | 0.124874619 | 8.59E-101 |
| 2.291346067 | 0.665413534 | 0.190891018 | 0.086089665 | 1.20E-69 |
| 5.009991055 | 0.941729323 | 0.140069917 | 0.015767728 | 6.04E-14 |
| 0.397849952 | 0.229323308 | 0.643592819 | 0.136208602 | 5.78E-110 |
| 0.771438538 | 0.415413534 | 0.276021087 | 0.119735039 | 1.21E-96 |
| 0.44665674 | 0.281954887 | 0.404896195 | 0.105775293 | 2.07E-85 |
| 0.391710683 | 0.22556391 | 0.607804547 | 0.129634376 | 1.22E-104 |
| 18.78483725 | 0.998120301 | 0.110423309 | 0.003599723 | 0.000339061 |
| 2.194750535 | 0.684210526 | 0.283028742 | 0.195568529 | 1.50E-158 |
| 2.800861035 | 0.783834586 | 0.147069274 | 0.055355943 | 3.27E-45 |
| 2.109687437 | 0.620300752 | 0.240383881 | 0.124172335 | 3.17E-100 |
| 1.071576705 | 0.437969925 | 0.745897357 | 0.311264565 | 4.77E-256 |
| 1.158390321 | 0.45112782 | 0.689845736 | 0.287179132 | 1.97E-235 |
| 0.870621101 | 0.345864662 | 0.299998874 | 0.08515513 | 6.70E-69 |
| 0.899007189 | 0.313909774 | 0.735863355 | 0.209382196 | 5.44E-170 |
| 19.19509447 | 0.998120301 | 0.109699097 | 0.001857324 | 0.010062588 |
| 0.896296165 | 0.454887218 | 0.232522246 | 0.094238104 | 3.68E-76 |
| 1.222448311 | 0.498120301 | 0.417313868 | 0.203734102 | 2.63E-165 |
| 9.712538904 | 0.986842105 | 0.115494248 | 0.009169617 | 1.05E-08 |
| 10.85902125 | 0.984962406 | 0.115063938 | 0.003933357 | 0.000179711 |
| 3.221056432 | 0.678571429 | 0.454229901 | 0.266735882 | 4.56E-218 |
| 0.716945945 | 0.334586466 | 0.337968668 | 0.092603413 | 7.49E-75 |
| 2.864991571 | 0.734962406 | 0.369475669 | 0.373724993 | 2.50551441914018e-310 |
| 0.473191506 | 0.259398496 | 0.497318641 | 0.145978889 | 6.74E-118 |
| 1.743704083 | 0.614661654 | 0.17788541 | 0.082476358 | 9.19E-67 |
| 1.102939461 | 0.492481203 | 0.308433448 | 0.143984014 | 2.82E-116 |
| 0.292688012 | 0.159774436 | 0.719592236 | 0.110543331 | 3.06E-89 |
| 8.362191724 | 0.988721805 | 0.118253289 | 0.01005431 | 2.07E-09 |
| 0.517957458 | 0.285714286 | 0.399924135 | 0.10241793 | 1.03E-82 |
| 37.59802005 | 0.994360902 | 0.101875365 | 0.065266575 | 4.54E-53 |
| 0.609483135 | 0.505703422 | 0.592323057 | 0.272797062 | 1.34E-221 |
| 0.512909001 | 0.522813688 | 0.564448902 | 0.310165399 | 2.96E-253 |
| 0.525610207 | 0.442965779 | 0.656819191 | 0.256120296 | 1.34E-207 |
| 1.949003313 | 0.760456274 | 0.50006461 | 0.369935474 | 9.48E-305 |
| 6.932676805 | 0.935361217 | 0.69372067 | 0.674773984 | 0 |
| 0.604348278 | 0.480988593 | 0.629845273 | 0.280063038 | 1.01E-227 |
| 0.602337469 | 0.501901141 | 0.642179508 | 0.300335964 | 6.87E-245 |
| 1.596163853 | 0.714828897 | 0.514870512 | 0.327933889 | 1.88E-268 |
| 1.235612786 | 0.646387833 | 0.483865378 | 0.354448182 | 2.65E-291 |
| 12.09195143 | 0.570342205 | 0.60750766 | 0.112329355 | 4.73E-90 |
| 3.02570371 | 0.874524715 | 0.61232944 | 0.52349491 | 0 |
| 0.747662771 | 0.608365019 | 0.628368433 | 0.345717112 | 9.37E-284 |
| 1.519251216 | 0.553231939 | 0.66759679 | 0.323226206 | 2.04E-264 |
| 0.54851756 | 0.41634981 | 0.723183396 | 0.268400439 | 6.69E-218 |

| | | | | |
|---|---|---|---|---|
| 1.981560851 | 0.828897338 | 0.491835941 | 0.430492114 | 0 |
| 0.765474494 | 0.589353612 | 0.660580041 | 0.389876046 | 3.6066792146411e-322 |
| 1.151005922 | 0.587452471 | 0.537193882 | 0.300600302 | 4.10E-245 |
| 0.533323005 | 0.47338403 | 0.679578361 | 0.300806785 | 2.74E-245 |
| 0.78518209 | 0.595057034 | 0.563587002 | 0.350485973 | 7.11E-288 |
| 1.307945604 | 0.714828897 | 0.488366117 | 0.376462164 | 1.94187822229593e-310 |
| 1.327891214 | 0.676806084 | 0.486889652 | 0.320271931 | 6.90E-262 |
| 0.67967641 | 0.566539924 | 0.706037099 | 0.357334967 | 8.37E-294 |
| 1.753643975 | 0.707224335 | 0.647388206 | 0.411857502 | 0 |
| 5.629523373 | 0.922053232 | 0.592701065 | 0.560696452 | 0 |
| 0.50863525 | 0.517110266 | 0.644656641 | 0.29989929 | 1.62E-244 |
| 1.011861772 | 0.538022814 | 0.699296815 | 0.336309698 | 1.20E-275 |
| 0.435784723 | 0.401140684 | 0.74301043 | 0.268156053 | 1.07E-217 |
| 0.665999804 | 0.517110266 | 0.703732576 | 0.335066797 | 1.41E-274 |
| 0.4442706 | 0.44486692 | 0.685750248 | 0.276339617 | 1.39E-224 |
| 7.293326886 | 0.880228137 | 0.714123376 | 0.578694786 | 0 |
| 0.523610769 | 0.511406844 | 0.710909364 | 0.357269194 | 9.54E-294 |
| 0.960470564 | 0.543726236 | 0.656673485 | 0.320510108 | 4.32E-262 |
| 0.366053096 | 0.412547529 | 0.767987233 | 0.286460022 | 3.98E-233 |
| 1.802073763 | 0.726235741 | 0.661666576 | 0.472916166 | 0 |
| 0.48939076 | 0.494296578 | 0.637272888 | 0.283259484 | 2.02E-230 |
| 0.500240686 | 0.524714829 | 0.712261167 | 0.366428169 | 1.07E-301 |
| 0.540785217 | 0.486692015 | 0.657723035 | 0.305476891 | 2.92E-249 |
| 0.740608591 | 0.564638783 | 0.654107165 | 0.34790009 | 1.22E-285 |
| 0.457272745 | 0.425855513 | 0.772526829 | 0.307060447 | 1.31E-250 |
| 3.511090055 | 0.488593156 | 0.616965896 | 0.189747207 | 1.17E-152 |
| 1.988260729 | 0.802281369 | 0.445548572 | 0.377859459 | 1.17117443932596e-311 |
| 0.754017264 | 0.564638783 | 0.694548387 | 0.381066849 | 1.84853964462502e-314 |
| 0.686926577 | 0.532319392 | 0.697140173 | 0.338038323 | 3.90E-277 |
| 0.675803928 | 0.585551331 | 0.549341775 | 0.318981373 | 8.77E-261 |
| 0.880922148 | 0.646387833 | 0.613528901 | 0.357406888 | 7.25E-294 |
| 1.208218197 | 0.711026616 | 0.513259636 | 0.36428165 | 7.84E-300 |
| 4.611644284 | 0.768060837 | 0.692083139 | 0.508966421 | 0 |
| 11.68239771 | 0.859315589 | 0.343434644 | 0.358457886 | 8.89E-295 |
| 0.671258351 | 0.5 | 0.603762181 | 0.274740248 | 3.10E-223 |
| 0.423629368 | 0.435361217 | 0.837401331 | 0.335155853 | 1.18E-274 |
| 8.954046839 | 0.952941176 | 0.275466294 | 0.334106601 | 2.45E-268 |
| 1.942921549 | 0.715686275 | 0.326366856 | 0.202280813 | 8.37E-160 |
| 12.01929178 | 0.960784314 | 0.280402537 | 0.292100395 | 3.11E-233 |
| 1.660775663 | 0.731372549 | 0.407900918 | 0.278640234 | 4.32E-222 |
| 6.309128043 | 0.917647059 | 0.373513578 | 0.35077243 | 2.10E-282 |
| 1.401148354 | 0.737254902 | 0.297343539 | 0.236204608 | 2.77E-187 |
| 3.070991198 | 0.688235294 | 0.349551199 | 0.176875712 | 2.02E-139 |
| 1.390492129 | 0.721568627 | 0.468446289 | 0.371317587 | 7.41E-300 |
| 14.88878642 | 0.982352941 | 0.554614382 | 0.750532445 | 0 |

| | | | | |
|---|---|---|---|---|
| 1.91750923 | 0.809803922 | 0.266221757 | 0.237843134 | 1.28E-188 |
| 11.91561334 | 0.984313725 | 0.554081838 | 0.699785732 | 0 |
| 2.376185715 | 0.682352941 | 0.408130239 | 0.27194755 | 1.44E-216 |
| 0.628380856 | 0.549019608 | 0.435357539 | 0.224595176 | 7.61E-178 |
| 0.382653001 | 0.343137255 | 0.660545245 | 0.176146078 | 7.74E-139 |
| 0.919486659 | 0.639215686 | 0.447296463 | 0.281346904 | 2.51E-224 |
| 0.845986678 | 0.607843137 | 0.555183501 | 0.299196825 | 3.95E-239 |
| 0.188907455 | 0.329411765 | 0.559240017 | 0.163276477 | 1.41E-128 |
| 1.090881268 | 0.668627451 | 0.675597959 | 0.392887657 | 2.58896327208563e-318 |
| 0.601579629 | 0.496078431 | 0.693473739 | 0.287124429 | 4.14E-229 |
| 2.260164668 | 0.801960784 | 0.255566327 | 0.220292261 | 2.35E-174 |
| 0.278030467 | 0.376470588 | 0.611147898 | 0.191735487 | 2.53E-151 |
| 0.707587083 | 0.570588235 | 0.365118158 | 0.22925357 | 1.25E-181 |
| 0.490388618 | 0.480392157 | 0.655610344 | 0.267413346 | 7.80E-213 |
| 0.863595666 | 0.631372549 | 0.297262983 | 0.20745097 | 5.68E-164 |
| 1.303161196 | 0.711764706 | 0.27383372 | 0.258290697 | 2.44E-205 |
| 0.887318962 | 0.37254902 | 0.512499241 | 0.167484078 | 6.30E-132 |
| 1.104466227 | 0.621568627 | 0.430049415 | 0.243459576 | 3.36E-193 |
| 0.447088292 | 0.429411765 | 0.501067483 | 0.179491989 | 1.64E-141 |
| 1.926095168 | 0.509803922 | 0.48135532 | 0.233368044 | 5.66E-185 |
| 0.414646262 | 0.454901961 | 0.64565644 | 0.232278641 | 4.35E-184 |
| 0.315527339 | 0.374509804 | 0.555678255 | 0.182525865 | 6.12E-144 |
| 1.657897658 | 0.768627451 | 0.669232464 | 0.506166879 | 0 |
| 0.591767765 | 0.405882353 | 0.656565185 | 0.217819281 | 2.36E-172 |
| 1.175100218 | 0.696078431 | 0.647655387 | 0.424611872 | 0 |
| 2.486535174 | 0.660784314 | 0.341818468 | 0.20552844 | 2.02E-162 |
| 5.04962345 | 0.947058824 | 0.597585768 | 0.614365259 | 0 |
| 2.149840478 | 0.817647059 | 0.255831142 | 0.211666021 | 2.24E-167 |
| 0.972478347 | 0.629411765 | 0.483706106 | 0.268284439 | 1.50E-213 |
| 3.744604436 | 0.831372549 | 0.587164479 | 0.427866475 | 0 |
| 0.535442497 | 0.545098039 | 0.430020645 | 0.223555316 | 5.31E-177 |
| 0.567254105 | 0.449019608 | 0.523190718 | 0.202298605 | 8.10E-160 |
| 0.365042069 | 0.423529412 | 0.532154615 | 0.215001854 | 4.50E-170 |
| 0.765625047 | 0.57254902 | 0.491063688 | 0.267453666 | 7.23E-213 |
| 11.04492177 | 0.858823529 | 0.269468977 | 0.089520321 | 1.50E-70 |
| 4.533063766 | 0.943137255 | 0.463846988 | 0.559943438 | 0 |
| 1.129681684 | 0.690196078 | 0.456051852 | 0.314107624 | 1.46E-251 |
| 17.40101536 | 0.992156863 | 0.302589911 | 0.473050344 | 0 |
| 13.00703233 | 0.858823529 | 0.631633581 | 0.410076495 | 0 |
| 0.587079948 | 0.523529412 | 0.560192789 | 0.283819823 | 2.26E-226 |
| 0.586080966 | 0.523529412 | 0.466726757 | 0.247561005 | 1.50E-196 |

| marker_test_q_value |
|---|
| 0 |
| 0 |
| 4.17E-70 |
| 1.18E-130 |
| 3.20E-113 |
| 1.17E-33 |
| 1.51E-31 |
| 2.51E-95 |
| 1.28E-14 |
| 3.22E-27 |
| 0 |
| 9.79E-98 |
| 0 |
| 0 |
| 0 |
| 1.04E-135 |
| 1.71E-220 |
| 3.84E-59 |
| 0 |
| 9.81E-17 |
| 6.86E-92 |
| 0 |
| 1.97E-108 |
| 0 |
| 4.88E-26 |
| 2.55E-89 |
| 2.05E-51 |
| 1.71E-59 |
| 0 |
| 3.35E-39 |
| 1.19E-103 |
| 3.79E-148 |
| 4.24E-122 |
| 1.18E-246 |
| 3.12E-19 |
| 3.42E-110 |
| 1.11E-164 |
| 0 |
| 3.49E-68 |
| 0 |
| 0 |
| 0 |
| 2.15E-52 |
| 9.07E-24 |

| |
|---|
| 4.49E-14 |
| 0 |
| 4.99E-20 |
| 4.84E-98 |
| 1 |
| 0 |
| 1.03E-79 |
| 8.87582659554987e-310 |
| 8.34E-189 |
| 2.96E-67 |
| 1.48E-91 |
| 8.05E-76 |
| 1.26E-62 |
| 2.70E-86 |
| 2.32E-66 |
| 1.22E-115 |
| 8.00E-109 |
| 7.05E-68 |
| 3.93E-221 |
| 1.58E-159 |
| 1.45E-74 |
| 2.16E-116 |
| 2.50E-52 |
| 6.88E-123 |
| 2.43E-93 |
| 2.86E-153 |
| 1.92E-121 |
| 5.94E-104 |
| 1.26E-95 |
| 1.23E-65 |
| 3.72E-130 |
| 1.70E-59 |
| 2.92E-70 |
| 2.46E-203 |
| 1.00E-71 |
| 1.46E-61 |
| 3.99E-99 |
| 1.42E-124 |
| 6.46E-75 |
| 4.48E-88 |
| 1.83E-111 |
| 3.95E-72 |
| 5.13E-64 |
| 1.01E-78 |
| 1.65E-114 |

| |
|---|
| 3.37E-132 |
| 1.30E-57 |
| 1.44E-223 |
| 5.16E-148 |
| 2.52E-137 |
| 7.65E-159 |
| 0 |
| 7.91E-117 |
| 6.67E-207 |
| 2.14E-59 |
| 2.07E-92 |
| 1.08E-91 |
| 1.10E-104 |
| 2.16E-53 |
| 9.80E-28 |
| 2.62E-42 |
| 1.61E-36 |
| 1.97E-42 |
| 3.69E-17 |
| 3.69E-40 |
| 1.89E-16 |
| 7.14E-37 |
| 8.56E-38 |
| 1.31E-31 |
| 3.99E-37 |
| 1.37E-50 |
| 6.23E-70 |
| 3.04E-55 |
| 4.74E-297 |
| 1.55E-258 |
| 1.24E-35 |
| 4.87E-148 |
| 1.42E-116 |
| 3.59E-42 |
| 1.15E-56 |
| 3.14E-45 |
| 8.61E-36 |
| 1.73E-26 |
| 1.60E-306 |
| 4.69E-112 |
| 3.45E-33 |
| 3.38E-39 |
| 1.92E-99 |
| 1.85E-55 |
| 3.57E-45 |

| |
|---|
| 2.69E-38 |
| 4.74E-35 |
| 7.96E-37 |
| 2.46E-63 |
| 3.09E-28 |
| 5.55E-71 |
| 3.74E-45 |
| 1.08E-59 |
| 2.71E-77 |
| 2.22656576196982e-310 |
| 8.15E-220 |
| 4.51E-54 |
| 9.58E-159 |
| 4.96E-35 |
| 1.70E-164 |
| 2.02E-66 |
| 0.091157931 |
| 0.032075998 |
| 1.09E-56 |
| 7.17E-17 |
| 5.43E-20 |
| 1 |
| 3.90E-40 |
| 1.26E-13 |
| 7.17E-117 |
| 0 |
| 2.91E-22 |
| 1.64E-116 |
| 7.47E-38 |
| 5.92E-09 |
| 3.88E-48 |
| 1.29E-68 |
| 2.04E-94 |
| 8.35E-180 |
| 3.25E-11 |
| 9.54E-19 |
| 1.06E-50 |
| 2.10E-168 |
| 1.30E-86 |
| 2.31E-66 |
| 1.35E-198 |
| 3.30E-65 |
| 5.91E-07 |
| 6.88E-42 |
| 3.00E-23 |

| |
|---|
| 1 |
| 1.30E-191 |
| 6.02E-43 |
| 1.41E-50 |
| 0 |
| 1.29E-55 |
| 4.26E-33 |
| 4.84E-52 |
| 0 |
| 0 |
| 1.83E-31 |
| 0 |
| 3.28E-57 |
| 1.23E-201 |
| 8.57E-08 |
| 0.000330254 |
| 1.48E-09 |
| 9.03E-216 |
| 1.61E-23 |
| 1.99E-18 |
| 6.50E-105 |
| 1.05E-125 |
| 6.49E-208 |
| 3.39E-140 |
| 4.33E-160 |
| 7.04E-177 |
| 5.74E-129 |
| 1.15E-110 |
| 3.44E-127 |
| 1.54E-124 |
| 2.48E-141 |
| 2.34E-118 |
| 1.04E-197 |
| 1.49E-247 |
| 3.25E-205 |
| 3.83E-137 |
| 1.22E-124 |
| 1.39E-247 |
| 1.87E-299 |
| 3.98E-302 |
| 2.68E-185 |
| 6.65E-124 |
| 4.08E-206 |
| 2.05E-176 |
| 3.22E-196 |

| |
|---|
| 2.87E-135 |
| 1.21E-160 |
| 1.19E-118 |
| 3.66E-174 |
| 1.31E-123 |
| 7.47E-160 |
| 1.97E-170 |
| 8.54E-132 |
| 5.41E-199 |
| 0 |
| 1.58E-135 |
| 5.08E-237 |
| 9.34E-132 |
| 1.96E-126 |
| 0 |
| 1.91E-157 |
| 2.28397392194325e-310 |
| 2.76E-231 |
| 1.38E-120 |
| 9.68E-200 |
| 3.94E-165 |
| 7.31E-224 |
| 5.94E-182 |
| 1.85E-203 |
| 9.34E-175 |
| 2.53E-200 |
| 4.16E-46 |
| 0 |
| 4.18E-42 |
| 4.79E-107 |
| 6.75E-47 |
| 9.36E-49 |
| 6.59E-93 |
| 8.01E-27 |
| 1.65E-31 |
| 2.59E-61 |
| 3.73E-60 |
| 1.81E-126 |
| 3.87E-40 |
| 3.01E-163 |
| 1.46E-208 |
| 1.77E-48 |
| 1.30E-48 |
| 1.14E-271 |
| 2.76E-196 |

| |
|---|
| 5.75E-26 |
| 6.52E-36 |
| 7.78E-30 |
| 5.28E-100 |
| 2.76E-56 |
| 1.65E-75 |
| 6.87E-155 |
| 1.10E-41 |
| 2.02E-40 |
| 5.27E-176 |
| 5.07E-252 |
| 3.96E-186 |
| 1.92E-220 |
| 3.18E-263 |
| 1.67E-58 |
| 6.20E-275 |
| 1.58E-169 |
| 3.30E-50 |
| 1.30E-106 |
| 5.41E-36 |
| 1 |
| 1 |
| 4.60E-39 |
| 7.76E-251 |
| 3.07E-46 |
| 4.14E-32 |
| 4.04E-145 |
| 1.93E-155 |
| 1.50E-71 |
| 1.92E-28 |
| 3.69E-42 |
| 4.31E-140 |
| 4.94E-186 |
| 3.76E-135 |
| 5.42E-125 |
| 6.46E-110 |
| 3.27E-134 |
| 1.87E-144 |
| 1.41E-156 |
| 7.64E-158 |
| 3.03E-194 |
| 2.41E-119 |
| 4.36E-161 |
| 9.30E-113 |
| 1.74E-153 |

| |
|---|
| 4.47E-247 |
| 5.60E-239 |
| 9.49E-214 |
| 4.56E-160 |
| 1.37E-129 |
| 3.18E-203 |
| 1.19E-194 |
| 1.18E-146 |
| 2.81E-116 |
| 8.70E-196 |
| 9.27E-146 |
| 5.59E-178 |
| 2.34E-115 |
| 7.26E-152 |
| 1.45E-131 |
| 2.26E-149 |
| 1.47E-122 |
| 2.11E-242 |
| 4.85E-237 |
| 2.14E-234 |
| 1.61E-133 |
| 2.69E-124 |
| 2.39E-143 |
| 3.58E-152 |
| 1.99E-143 |
| 3.13E-133 |
| 1.85E-120 |
| 3.82E-221 |
| 6.94E-157 |
| 1.22E-186 |
| 3.46E-126 |
| 8.31E-160 |
| 1.04E-154 |
| 1.84E-152 |
| 6.25E-176 |
| 4.84E-117 |
| 5.45E-73 |
| 1.43E-165 |
| 1.32E-171 |
| 1.67E-53 |
| 4.52E-68 |
| 8.84E-49 |
| 1.19E-106 |
| 7.04E-142 |
| 3.72E-57 |

| |
|---|
| 8.07E-140 |
| 1.06E-103 |
| 9.16E-65 |
| 2.60E-201 |
| 3.40E-134 |
| 3.23E-77 |
| 1.86E-82 |
| 1.97E-88 |
| 8.43E-167 |
| 6.47E-110 |
| 2.48E-72 |
| 1.37E-307 |
| 1.99E-83 |
| 2.16E-135 |
| 1.24E-87 |
| 8.55E-174 |
| 1.64E-111 |
| 8.92E-163 |
| 7.88E-79 |
| 1.10E-93 |
| 2.59E-126 |
| 4.56E-112 |
| 4.51E-73 |
| 3.00E-77 |
| 8.22E-79 |
| 5.96E-90 |
| 6.14E-115 |
| 4.10E-74 |
| 8.50E-82 |
| 2.80E-120 |
| 9.32E-75 |
| 5.02E-161 |
| 1.80E-65 |
| 5.61E-81 |
| 8.05E-96 |
| 5.49E-212 |
| 4.92E-96 |
| 3.92E-110 |
| 8.57E-72 |
| 1.18E-71 |
| 8.95E-81 |
| 2.55E-224 |
| 7.57E-199 |
| 1.98E-237 |
| 1.16E-245 |

| |
|---|
| 2.87E-202 |
| 5.10E-222 |
| 4.64E-262 |
| 7.72E-188 |
| 9.39E-208 |
| 1.63E-222 |
| 2.40E-177 |
| 4.20E-175 |
| 6.66E-168 |
| 6.05E-148 |
| 4.48E-135 |
| 5.29E-180 |
| 2.88E-229 |
| 1.54E-154 |
| 1.79E-178 |
| 2.49E-214 |
| 1.49E-187 |
| 0 |
| 1.31E-139 |
| 6.90E-214 |
| 7.95E-241 |
| 2.37E-159 |
| 9.82E-268 |
| 1.81E-144 |
| 1.57E-195 |
| 1.46E-186 |
| 9.05E-283 |
| 1.43E-262 |
| 6.06E-203 |
| 1.42E-192 |
| 7.87E-172 |
| 1.07E-228 |
| 1.07E-162 |
| 3.07E-243 |
| 2.42E-198 |
| 2.35E-224 |
| 4.97E-152 |
| 2.98E-174 |
| 6.85E-170 |
| 4.84E-186 |
| 2.30E-170 |
| 6.05E-239 |
| 3.96E-176 |
| 7.11E-212 |
| 5.97E-194 |

| |
|---|
| 5.69E-141 |
| 2.71E-16 |
| 2.82E-19 |
| 5.18E-30 |
| 7.05E-27 |
| 9.34E-11 |
| 2.27E-23 |
| 2.73E-22 |
| 8.36E-14 |
| 1.36E-09 |
| 9.84E-18 |
| 6.97E-17 |
| 4.93E-51 |
| 8.88E-41 |
| 3.89E-37 |
| 8.90E-32 |
| 2.06E-19 |
| 1.32E-13 |
| 1.18E-15 |
| 4.18E-18 |
| 7.76E-24 |
| 2.32E-51 |
| 3.92E-26 |
| 1.48E-16 |
| 1.26E-14 |
| 3.07E-32 |
| 5.64E-13 |
| 1.00E-31 |
| 4.04E-28 |
| 4.05E-12 |
| 1.84E-11 |
| 2.71E-25 |
| 4.80E-21 |
| 1.93E-22 |
| 8.07E-15 |
| 1.28E-30 |
| 2.59E-10 |
| 7.71E-32 |
| 2.40E-22 |
| 1.94E-43 |
| 1.65E-34 |
| 4.77E-17 |
| 8.39E-18 |
| 2.48E-13 |
| 7.97E-30 |

| |
|---|
| 2.78E-18 |
| 4.61E-26 |
| 9.25E-20 |
| 3.10E-21 |
| 2.36E-26 |
| 1.89E-22 |
| 2.89E-12 |
| 1.45E-60 |
| 7.91E-19 |
| 5.78E-64 |
| 1.33E-13 |
| 1.04E-15 |
| 3.41E-25 |
| 7.00E-32 |
| 1.75E-78 |
| 8.83E-13 |
| 6.04E-93 |
| 3.64E-27 |
| 1.39E-21 |
| 5.30E-17 |
| 3.93E-74 |
| 1.65E-44 |
| 3.90E-23 |
| 1.18E-17 |
| 3.46E-31 |
| 8.40E-12 |
| 7.40E-39 |
| 2.41E-26 |
| 4.36E-14 |
| 3.37E-10 |
| 1.85E-07 |
| 5.88E-20 |
| 2.07E-21 |
| 2.92E-21 |
| 1.52E-07 |
| 1.18E-46 |
| 7.38E-75 |
| 1.68E-10 |
| 8.04E-25 |
| 1.46E-36 |
| 6.78E-14 |
| 4.96E-15 |
| 7.97E-12 |
| 1.82E-29 |
| 6.51E-90 |

| |
|---|
| 2.92E-43 |
| 2.57E-09 |
| 2.07E-12 |
| 1.23E-27 |
| 1.23E-19 |
| 8.06E-36 |
| 1.16E-20 |
| 7.67E-19 |
| 1.82E-15 |
| 5.54E-22 |
| 7.77E-37 |
| 3.95E-272 |
| 1.40E-163 |
| 1.21E-102 |
| 2.84E-164 |
| 8.53E-186 |
| 2.64E-30 |
| 8.51E-111 |
| 1.76E-10 |
| 6.29E-09 |
| 8.37E-161 |
| 1.90E-108 |
| 4.76E-107 |
| 1.31E-19 |
| 3.83E-14 |
| 8.82E-122 |
| 0 |
| 0 |
| 6.90E-209 |
| 4.63E-253 |
| 0 |
| 2.84E-23 |
| 1.32E-62 |
| 1.35E-196 |
| 4.02E-211 |
| 2.03E-232 |
| 0 |
| 6.01E-105 |
| 0 |
| 0 |
| 3.75E-40 |
| 1.45E-104 |
| 6.44E-94 |
| 0 |
| 1.23E-123 |

| |
|---|
| 4.66E-81 |
| 4.52E-159 |
| 0 |
| 0 |
| 1.71E-174 |
| 6.82E-100 |
| 0 |
| 3.07E-114 |
| 1.87E-81 |
| 0 |
| 8.81E-08 |
| 0 |
| 7.32E-51 |
| 7.09E-174 |
| 4.32E-78 |
| 4.58E-147 |
| 0 |
| 0 |
| 1.82E-255 |
| 1.01E-272 |
| 1.23E-262 |
| 6.50E-205 |
| 2.72E-250 |
| 7.72E-300 |
| 3.95E-149 |
| 1.52E-153 |
| 9.36E-282 |
| 6.48E-226 |
| 5.03E-158 |
| 2.67E-204 |
| 1.98E-237 |
| 2.90E-202 |
| 3.07E-127 |
| 0 |
| 3.97E-307 |
| 7.98E-209 |
| 3.57E-246 |
| 0 |
| 2.43E-163 |
| 8.96E-295 |
| 1.17E-217 |
| 0 |
| 0 |
| 1.08E-266 |
| 9.00E-239 |

| |
|---|
| 1.77E-227 |
| 4.05E-286 |
| 1.46E-205 |
| 0 |
| 0 |
| 0 |
| 0 |
| 1.34E-109 |
| 6.65E-122 |
| 1.19E-162 |
| 1.49E-194 |
| 6.50E-88 |
| 2.19E-209 |
| 8.00E-118 |
| 2.40308078305585e-312 |
| 0 |
| 0 |
| 9.31E-278 |
| 7.76E-204 |
| 4.96E-264 |
| 0 |
| 0 |
| 0 |
| 1.04E-298 |
| 0 |
| 0 |
| 1.59E-279 |
| 1.60E-251 |
| 0 |
| 0 |
| 3.85964519550872e-310 |
| 1.62814763479759e-315 |
| 0 |
| 3.01E-222 |
| 0 |
| 7.63E-283 |
| 0 |
| 1.02100300845087e-311 |
| 3.83E-264 |
| 0 |
| 0 |
| 0 |
| 2.29E-289 |
| 0 |
| 1.72397575273139e-314 |

| |
|---|
| 1.05E-291 |
| 0 |
| 4.58E-216 |
| 0 |
| 0 |
| 0 |
| 4.21E-291 |
| 1.22E-272 |
| 0 |
| 2.39E-296 |
| 0 |
| 0 |
| 6.25E-252 |
| 0 |
| 2.47E-234 |
| 4.12E-284 |
| 6.14E-299 |
| 0 |
| 0 |
| 0 |
| 1.49E-297 |
| 0 |
| 4.86E-306 |
| 0 |
| 0 |
| 1.74E-281 |
| 5.43E-113 |
| 1 |
| 1.93E-63 |
| 0.464802946 |
| 2.04E-103 |
| 1.65E-81 |
| 0 |
| 4.64E-224 |
| 2.82E-96 |
| 1.89E-134 |
| 1 |
| 0.705948295 |
| 1 |
| 1.31E-07 |
| 9.45E-91 |
| 1.34E-210 |
| 3.50E-263 |
| 9.50E-69 |
| 2.87E-140 |

| |
|---|
| 1.19E-07 |
| 8.08E-95 |
| 1.13E-63 |
| 5.69E-08 |
| 5.44E-104 |
| 1.14E-90 |
| 1.95E-79 |
| 1.15E-98 |
| 1 |
| 1.42E-152 |
| 3.08E-39 |
| 2.98E-94 |
| 4.49E-250 |
| 1.85E-229 |
| 6.30E-63 |
| 5.12E-164 |
| 1 |
| 3.47E-70 |
| 2.48E-159 |
| 0.00992113 |
| 1 |
| 4.29E-212 |
| 7.05E-69 |
| 2.36E-304 |
| 6.35E-112 |
| 8.65E-61 |
| 2.66E-110 |
| 2.88E-83 |
| 0.001952317 |
| 9.68E-77 |
| 4.27E-47 |
| 1.26E-215 |
| 2.78E-247 |
| 1.26E-201 |
| 8.93E-299 |
| 0 |
| 9.50E-222 |
| 6.47E-239 |
| 1.77E-262 |
| 2.50E-285 |
| 4.46E-84 |
| 0 |
| 8.82E-278 |
| 1.92E-258 |
| 6.30E-212 |

| |
|---|
| 0 |
| 3.39585078114925e-316 |
| 3.86E-239 |
| 2.58E-239 |
| 6.69E-282 |
| 1.83E-304 |
| 6.50E-256 |
| 7.88E-288 |
| 0 |
| 0 |
| 1.52E-238 |
| 1.13E-269 |
| 1.01E-211 |
| 1.32E-268 |
| 1.31E-218 |
| 0 |
| 8.98E-288 |
| 4.06E-256 |
| 3.75E-227 |
| 0 |
| 1.90E-224 |
| 1.01E-295 |
| 2.75E-243 |
| 1.15E-279 |
| 1.23E-244 |
| 1.10E-146 |
| 1.10E-305 |
| 0.00E+00 |
| 3.67E-271 |
| 8.25E-255 |
| 6.82E-288 |
| 7.38E-294 |
| 0 |
| 8.37E-289 |
| 2.92E-217 |
| 1.11E-268 |
| 2.31E-262 |
| 7.88E-154 |
| 2.92E-227 |
| 4.07E-216 |
| 1.98E-276 |
| 2.61E-181 |
| 1.90E-133 |
| 6.98E-294 |
| 0 |

| |
|---|
| 1.21E-182 |
| 0 |
| 1.35E-210 |
| 7.17E-172 |
| 7.29E-133 |
| 2.36E-218 |
| 3.72E-233 |
| 1.33E-122 |
| 2.43762542401587e-312 |
| 3.90E-223 |
| 2.21E-168 |
| 2.39E-145 |
| 1.18E-175 |
| 7.34E-207 |
| 5.35E-158 |
| 2.30E-199 |
| 5.93E-126 |
| 3.17E-187 |
| 1.54E-135 |
| 5.33E-179 |
| 4.10E-178 |
| 5.77E-138 |
| 0 |
| 2.23E-166 |
| 0 |
| 1.90E-156 |
| 0 |
| 2.11E-161 |
| 1.41E-207 |
| 0 |
| 5.00E-171 |
| 7.63E-154 |
| 4.24E-164 |
| 6.80E-207 |
| 1.41E-64 |
| 0 |
| 1.37E-245 |
| 0 |
| 0 |
| 2.12E-220 |
| 1.41E-190 |

### Supplemental Table 3

*Percentage of NC-derived and non-NC-derived neurons in adult Wnt1-cre:tdTomato mice*

| | No of neurons | % Wnt1:tdT+ | % Wnt1:tdT- |
|---|---|---|---|
| **Mouse 1** | 185 | 61.62 | 38.38 |
| **Mouse 2** | 276 | 53.62 | 46.38 |
| **Mouse 3** | 239 | 45.60 | 54.40 |
| **Mouse 4** | 485 | 54.64 | 45.36 |
| **Mouse 5** | 686 | 60.64 | 39.36 |
| **Mouse 6** | 345 | 58.26 | 41.74 |

*Percentage of mesoderm-derived and non-mesoderm-derived neurons in adult Mesp1-cre:tdTomato mice*

| | No of neurons | % Mesp1:tdT+ | % Mesp1:tdT- |
|---|---|---|---|
| **Mouse 1** | 147 | 53.06 | 46.94 |
| **Mouse 2** | 137 | 53.28 | 46.72 |
| **Mouse 3** | 200 | 44.50 | 55.50 |

*Percentage of tdTomato-expressing neurons of all NOS1+ neurons in adult Wnt1-cre:tdTomato mice*

| | NOS1+ counted | % Wnt1:tdT+ | % Wnt1:tdT- |
|---|---|---|---|
| **Mouse 1** | 248 | 64.04 | 35.96 |
| **Mouse 2** | 200 | 78.00 | 22.00 |
| **Mouse 3** | 168 | 76.78 | 23.22 |

*Percentage of tdTomato-expressing neurons of all CGRP+ neurons in adult Wnt1-cre:tdTomato mice*

| | CGRP+ counted | % Wnt1:tdT+ | % Wnt1:tdT- |
|---|---|---|---|
| **Mouse 1** | 41 | 25.47 | 74.53 |
| **Mouse 2** | 56 | 23.37 | 76.63 |
| **Mouse 3** | 49 | 23.87 | 76.13 |

*Age-wise changes in percentages of MENs in myenteric plexus neurons of Wnt1-cre:tdTomato mice*

| | P11 mice | |
|---|---|---|
| | # neurons | % MENs |
| 1 | 374 | 2.14 |
| 2 | 534 | 10.00 |
| 3 | 419 | 0.23 |

| | P22 mice | | |
|---|---|---|---|
| | # neurons | % MENs | Mean neurons/ganglia |
| 1 | 245 | 29.79 | 16.34 ± 2.07 |
| 2 | 175 | 27.43 | 11.67 ± 2.07 |
| 3 | 322 | 31.68 | 21.47 ± 1.61 |

| | P60 mice | | |
|---|---|---|---|
| | # neurons | % MENs | Mean neurons/ganglia |
| 1 | 185 | 38.38 | 15.42 ± 3.59 |
| 2 | 276 | 46.38 | 21.23 ± 4.22 |
| 3 | 239 | 54.40 | 19.91 ± 3.26 |

| | P180 mice | | |
|---|---|---|---|
| | # neurons | % MENs | Mean neurons/ganglia |
| 1 | 171 | 60.97 | 15.54 ± 2.52 |
| 2 | 239 | 50.39 | 21.73 ± 3.69 |
| 3 | 176 | 60.52 | 14.67 ± 1.80 |

| | P510 mice | | |
|---|---|---|---|
| | # neurons | % MENs | Mean neurons/ganglia |
| 1 | 261 | 98.85 | 13.74 ± 2.22 |
| 2 | 360 | 93.22 | 17.14 ± 3.46 |
| 3 | 375 | 95.91 | 13.89 ± 1.30 |

*Percentage of MENs in myenteric plexus of P20 Wnt1-cre:tdTomato mice dosed with and without GDNF from the P10 age*

| | Control | | |
|---|---|---|---|
| | # neurons | % MENs | Mean neurons/ganglia |
| 1 | 477 | 22.85 | 22.71 ± 6.99 |
| 2 | 296 | 20.27 | 12.86 ± 2.37 |
| 3 | 577 | 34.49 | 26.23 ± 4.55 |

| | GDNF | | |
|---|---|---|---|
| | # neurons | % MENs | Mean neurons/ganglia |
| 1 | 418 | 3.83 | 19.90 ± 4.23 |
| 2 | 350 | 4.00 | 15.21 ± 3.38 |
| 3 | 533 | 3.75 | 38.00 ± 4.89 |

*Percentage of MENs in myenteric plexus of P20 Wnt1-cre:tdTomato mice dosed with and without HGF from the P10 age*

| | Control | | |
|---|---|---|---|
| | # neurons | % MENs | Mean neurons/ganglia |
| 1 | 325 | 23.69 | 15.48 ± 1.88 |
| 2 | 439 | 17.77 | 20.90 ± 3.20 |
| 3 | 454 | 36.12 | 21.62 ± 3.40 |
| 4 | 336 | 44.05 | 16.00 ± 2.46 |
| 5 | 416 | 15.38 | 27.73 ± 3.73 |

| | HGF | | |
|---|---|---|---|
| | # neurons | % MENs | Mean neurons/ganglia |
| 1 | 394 | 40.86 | 18.76 ± 2.51 |
| 2 | 409 | 40.83 | 19.48 ± 2.88 |
| 3 | 386 | 39.38 | 18.38 ± 2.71 |
| 4 | 326 | 62.27 | 15.52 ± 1.75 |
| 5 | 189 | 63.49 | 18.90 ± 2.01 |

*Percentage of MENs in myenteric plexus of 17 month old mice dosed with and without GDNF*

| | Control | | |
|---|---|---|---|
| | # neurons | % MENs | Mean neurons/ganglia |
| 1 | 141 | 88.65 | 15.67 ± 5.22 |
| 2 | 238 | 93.28 | 21.64 ± 6.52 |
| 3 | 225 | 89.78 | 20.45 ± 6.16 |
| 4 | 213 | 84.04 | 18.54 ± 5.59 |
| 5 | 92 | 88.04 | 11.5 ± 4.06 |

| | GDNF | | |
|---|---|---|---|
| | # neurons | % MENs | Mean neurons/ganglia |
| 1 | 111 | 84.68 | 12.33 ± 4.11 |
| 2 | 215 | 60.00 | 19.54 ± 5.89 |
| 3 | 160 | 88.12 | 14.54 ± 4.38 |
| 4 | 183 | 60.11 | 17.36 ± 5.23 |
| 5 | 130 | 77.69 | 16.25 ± 5.74 |

*Percentage of RET+ NENs in myenteric plexus of 17 month old mice dosed with and without GDNF*

| | Control | | |
|---|---|---|---|
| | # neurons | % RET | Mean neurons/ganglia |
| 1 | 170 | 8.23 | 17.00 ± 5.37 |
| 2 | 227 | 6.17 | 20.64 ± 6.22 |
| 3 | 232 | 5.17 | 21.09 ± 6.36 |
| 4 | 188 | 9.04 | 18.8 ± 5.94 |

| | GDNF | | |
|---|---|---|---|
| | # neurons | % RET | Mean neurons/ganglia |
| 1 | 137 | 19.70 | 13.70 ± 4.33 |
| 2 | 198 | 24.24 | 18.00 ± 5.43 |
| 3 | 200 | 11.00 | 18.18 ± 5.48 |
| 4 | 139 | 24.46 | 15.44 ± 5.14 |

*Percentage of Ret-CFP+ NENs in myenteric plexus of Ret+/- mice at 9 and 16 weeks*

| | 9 weeks | | |
|---|---|---|---|
| | # neurons | % CFP+ | Mean neurons/ganglia |
| 1 | 268 | 14.18 | 24.36 ± 6.27 |
| 2 | 275 | 35.65 | 19.64 ± 2.28 |
| 3 | 294 | 28.91 | 24.50 ± 3.40 |

| | 16 weeks | | |
|---|---|---|---|
| | # neurons | % CFP+ | Mean neurons/ganglia |
| 1 | 305 | 10.16 | 16.05 ± 3.68 |
| 2 | 306 | 13.40 | 13.91 ± 2.96 |
| 3 | 263 | 11.79 | 12.52 ± 2.73 |
| 4 | 122 | 15.57 | 11.09 ± 3.34 |
| 5 | 231 | 14.75 | 17.77 ± 4.93 |

*Percentage of MHCst+ MENs in myenteric plexus of Ret+/- and Ret+/+ mice at 9 and 16 weeks*

| Ret+/+ | 9 weeks | | |
|---|---|---|---|
| | # neurons | % MENs | Mean neurons/ganglia |
| 1 | 197 | 41.12 | 17.91 ± 2.96 |
| 2 | 286 | 49.65 | 26.00 ± 3.96 |
| 3 | 297 | 39.06 | 19.80 ± 3.62 |

| Ret+/+ | 16 weeks | | |
|---|---|---|---|
| | # neurons | % MENs | Mean neurons/ganglia |
| 1 | 217 | 56.68 | 19.73 ± 2.69 |
| 2 | 218 | 59.17 | 19.82 ± 3.12 |
| 3 | 316 | 60.44 | 19.75 ± 2.89 |
| 4 | 228 | 57.72 | 20.73 ± 4.45 |
| 5 | 297 | 38.38 | 18.56 ± 4.11 |

| Ret+/- | 9 weeks | | |
|---|---|---|---|
| | # neurons | % MENs | Mean neurons/ganglia |
| 1 | 220 | 44.54 | 20.00 ± 3.87 |
| 2 | 210 | 69.05 | 19.10 ± 2.73 |
| 3 | 214 | 62.62 | 13.37 ± 2.50 |

| Ret+/- | 16 weeks | | |
|---|---|---|---|
| | # neurons | % MENs | Mean neurons/ganglia |
| 1 | 182 | 91.21 | 16.54 ± 2.73 |
| 2 | 192 | 87.43 | 17.36 ± 1.25 |
| 3 | 215 | 83.25 | 13.43 ± 2.01 |
| 4 | 200 | 70.00 | 18.18 ± 3.74 |
| 5 | 147 | 82.84 | 21.00 ± 2.40 |

*Percent MET+ neurons*

| | # MET+ neurons | Wnt1-tdT+ | Wnt1-tdT- |
|---|---|---|---|
| 1 | 68 | 16.18 | 83.82 |
| 2 | 56 | 17.86 | 82.14 |
| 3 | 28 | 21.43 | 78.57 |

*Pattern Weights of the four MEN-specific NNMF patterns as represented in the post-natal mesenchymal single cell RNA sequencing data from Gut Cell Atlas*

| | | Pattern 16 | | | Pattern 27 | | | Pattern 32 | | | Pattern 41 | |
|---|---|---|---|---|---|---|---|---|---|---|---|---|
| **Age** | Mean | Standard Error | Sample size | Mean | Standard Error | Sample size | Mean | Standard Error | Sample size | Mean | Standard Error | Sample size |
| Juvenile | 0.0049 | 0.00023 | 5 | -0.0036 | 0.0012 | 5 | 0.024 | 0.0005 | 5 | -0.0029 | 0.0002 | 5 |
| Adult | 0.0049 | 0.0006 | 3 | 0.0085 | 0.0021 | 3 | 0.028 | 0.0030 | 3 | 0.0025 | 0.0028 | 3 |
| Aging | 0.0053 | 0.0007 | 3 | 0.0097 | 0.0019 | 3 | 0.033 | 0.0016 | 3 | 0.0058 | 0.0014 | 3 |

*Age-wise changes in whole gut transit time of Ret+/+ and Ret+/- mice*

| Age weeks | Whole gut transit time (in minutes) of *Ret +/+* mice | | | | | | | | Whole gut transit time (in minutes) of *Ret +/-* mice | | | | | | | | | |
|---|---|---|---|---|---|---|---|---|---|---|---|---|---|---|---|---|---|---|
| **9** | 131 | 119 | 59 | 135 | 113 | 88 | 103 | 86 | 73 | 105 | 93 | 91 | 87 | 83 | 134 | 143 | 153 | 132 |
| **12** | 83 | 89 | 89 | 100 | 106 | 86 | 110 | 78 | 89 | 124 | 106 | 98 | 77 | 71 | 128 | 73 | 82 | 124 |
| **14** | 107 | 103 | 120 | 141 | 111 | 117 | 166 | 110 | 100 | 189 | 190 | 93 | 107 | 106 | 190 | 207 | 125 | 96 |
| **16** | 111 | 130 | 106 | 116 | 130 | 122 | 116 | 140 | 200 | 237 | 194 | 122 | 185 | 157 | 102 | 101 | 156 | 119 |

*Fold change (transcript expression) of Gdnf in murine small intestinal LM-MP at different ages*

| | P30 | P90 |
|---|---|---|
| 1 | 0.8211 | 0.6884 |
| 2 | 0.6600 | 0.6081 |
| 3 | 1.6392 | 0.7047 |
| 4 | 1.6011 | 0.9263 |
| 5 | 0.7185 | 0.1380 |
| 6 | 0.9784 | 0.1683 |

*Expression levels (Fluorescence intensity) of Hgf, compared to Beta-actin, in murine small intestinal LM-MP at different ages*

| | P10 | P30 | P90 |
|---|---|---|---|
| 1 | 0.008 | 0.007 | 0.022 |
| 2 | 0.006 | 0.017 | 0.013 |
| 3 | 0.006 | 0.011 | 0.016 |

*Expression levels (Fluorescence intensity) of Gdnf, compared to Beta-actin, in murine small intestinal LM-MP at different ages*

| | P10 | P30 | P90 |
|---|---|---|---|
| 1 | 1.372 | 0.069 | 0.069 |
| 2 | 1.271 | 0.095 | 0.031 |
| 3 | 1.392 | 0.053 | 0.053 |

*Whole gut transit time (in min) of 17 month old mice before and after GDNF or Saline treatment*

| | Before Treatment | | After Treatment | |
|---|---|---|---|---|
| | Control | GDNF | Control | GDNF |
| 1 | 210 | 182 | 210 | 74 |
| 2 | 210 | 210 | 164 | 110 |
| 3 | 210 | 152 | 168 | 87 |
| 4 | 172 | 210 | 162 | 123 |
| 5 | 210 | 210 | 171 | 111 |

*Effect of GDNF on MHCst neuron proportions in 17 month old mice*

| | Control | | GDNF | |
|---|---|---|---|---|
| | Neurons counted | %MHCst | Neurons counted | %MHCst |
| 1 | 141 | 88.65 | 111 | 84.68 |
| 2 | 238 | 93.27 | 215 | 60.00 |
| 3 | 225 | 89.78 | 160 | 88.12 |
| 4 | 213 | 84.03 | 183 | 60.11 |
| 5 | 92 | 88.04 | 130 | 77.69 |

*Effect of GDNF on RET neuron proportions in 17 month old mice*

| | Control | | GDNF | |
|---|---|---|---|---|
| | Fields counted | RET+ | Fields counted | RET+ |
| 1 | 10 | 14 | 10 | 27 |
| 2 | 10 | 14 | 10 | 48 |
| 3 | 10 | 12 | 10 | 22 |
| 4 | 12 | 17 | 12 | 34 |

*Effect of GDNF on neuronal numbers in 17 month old mice*

| | Control | GDNF |
|---|---|---|
| 1 | 15.67 ± 2.63 | 12.33 ± 3.55 |
| 2 | 21.64 ± 4.18 | 19.54 ± 2.55 |
| 3 | 20.45 ± 3.19 | 14.54 ± 1.91 |
| 4 | 18.54 ± 2.84 | 17.36 ± 2.61 |
| 5 | 11.50 ± 3.81 | 16.25 ± 2.30 |

*Fold change (transcript expression) of Hgf in murine small intestinal LM-MP at different ages*

| | P10 | P30 | P90 |
|---|---|---|---|
| 1 | 2.146 | 5.442 | 6.114 |
| 2 | 1.867 | 4.747 | 5.337 |
| 3 | 0.543 | 2.919 | 4.745 |
| 4 | 0.659 | 2.683 | 4.123 |
| 5 | 0.789 | 2.703 | 5.593 |
| 6 | 0.884 | 2.928 | 5.824 |

*Percent MHCst neurons that express tdTomato in Wnt1-cre:tdTomato mice*

| | MHCst+ neurons counted | Percent tdTomato+ |
|---|---|---|
| 1 | 29 | 0 |
| 2 | 30 | 0 |
| 3 | 48 | 0 |

*UMI per cluster (6 month data)*

| | Average ± SEM | # Cells |
|---|---|---|
| **MENs** | 4023.91 ± 85.35 | 2223 |
| **NENs** | 3391.71 ± 386.52 | 77 |
| **Neuroglia** | 1137.93 ± 27.93 | 1660 |

*UMI per cluster (P21 data)*

| | Average ± SEM | # Cells |
|---|---|---|
| **MENs** | 8891.84 ± 393.74 | 504 |
| **NENs** | 3263.34 ± 142.29 | 502 |
| **Neuroglia** | 3493.49 ± 120.25 | 835 |

**Lengenivity Data**

| | Mean and SEM |
|---|---|
| **HGF** | 0.0027 ± 0.0009 |
| **GDNF** | -0.0008 ± 0.0004 |
| **RET** | -0.0162 ± 0.0013 |

*Percent ChAT+ neurons*

| | # neurons | Wnt1-tdT+ | Wnt1-tdT- |
|---|---|---|---|
| 1 | 440 | 68.77 | 31.23 |
| 2 | 594 | 74.84 | 25.16 |
| 3 | 648 | 77.31 | 22.69 |

*Feret diameter (in microns)*

| | NENs | MENs |
|---|---|---|
| 1 | 12.32 ± 0.34 | 17.38 ± 0.55 |
| 2 | 13.47 ± 0.42 | 18.38 ± 0.72 |
| 3 | 13.31 ± 0.44 | 16.65 ± 0.56 |

*Ret transcripts in small intestinal LM-MP*

| | Ret WT | Ret Het |
|---|---|---|
| 1 | 0.777546 | 0.435879 |
| 2 | 1.232852 | 0.123450 |
| 3 | 1.043912 | 0.815478 |
| 4 | | 0.583175 |

### Supplementary Table 4

scRNAseq data metrics from 6 month old mouse LM-MP cells

| n_cells | annotated_type | pct_batch1 | pct_batch2 | num_genes | mt_ratio | umi |
|---|---|---|---|---|---|---|
| 2248 | Macrophage-A | 0.551 | 0.449 | 238.8 | 0.09 | 440.98 |
| 2223 | MENs | 0.709 | 0.291 | 1255.92 | 0.05 | 4023.91 |
| 1736 | Macrophage-B | 0.548 | 0.452 | 1073.43 | 0.03 | 2654.48 |
| 1660 | Neuroglia | 0.74 | 0.26 | 633.96 | 0.06 | 1137.93 |
| 908 | Vascular endothelium | 0.792 | 0.208 | 854.27 | 0.07 | 1797.43 |
| 776 | Smooth muscle cells | 0.619 | 0.381 | 302.1 | 0.11 | 527.09 |
| 670 | RBC | 0.481 | 0.519 | 94.48 | 0.03 | 2380.12 |
| 240 | Pdgfra+ Fibroblasts | 0.542 | 0.458 | 776.06 | 0.06 | 1569.2 |
| 163 | B Lymphocytes | 0.067 | 0.933 | 602.87 | 0.05 | 1335.6 |
| 101 | Penk+ Fibroblasts | 0 | 1 | 1008.13 | 0.04 | 2272.25 |
| 80 | T cells | 0.125 | 0.875 | 763.98 | 0.05 | 1762.33 |
| 58 | NENs | 0.741 | 0.259 | 1031.98 | 0.07 | 3105.91 |
| 39 | NK cells | 0.256 | 0.744 | 752.21 | 0.04 | 1577.36 |
| 38 | Macrophage-C | 0.553 | 0.447 | 1631.5 | 0.03 | 4486.82 |
| 38 | NK cells | 0.184 | 0.816 | 711.11 | 0.04 | 1480 |
| 34 | Macrophage-C | 0.206 | 0.794 | 1334.94 | 0.02 | 4124.62 |
| 31 | Smooth muscle cells B | 0.742 | 0.258 | 588.87 | 0.05 | 975.58 |
| 22 | Macrophage-B | 0.727 | 0.273 | 1646.05 | 0.02 | 4178.14 |
| 22 | Unknown | 0.136 | 0.864 | 637.55 | 0.09 | 1617.18 |
| 19 | NENs | 0.105 | 0.895 | 1391.16 | 0.06 | 4264.16 |
| 17 | Macrophage-C | 0.294 | 0.706 | 1813.35 | 0.03 | 6721.53 |

scRNAseq data metrics from P21 mouse LM-MP cells

| n_cells | annotated_type | pct_batch1 | pct_batch2 | num_genes | mt_ratio | umi | umi_sem |
|---|---|---|---|---|---|---|---|
| 277 | Endothelial | 0.794 | 0.206 | 1712.19 | 0.05 | 5163.03 | 310.22 |
| 2165 | Fibroblasts | 0.824 | 0.176 | 1239.34 | 0.07 | 3187.54 | 68.51 |
| 312 | ICC | 0.76 | 0.24 | 1253.98 | 0.09 | 3001.8 | 169.94 |
| 231 | Macrophage | 0.879 | 0.121 | 2190.81 | 0.05 | 7270.13 | 478.34 |
| 510 | MENs | 0.751 | 0.249 | 1948.45 | 0.05 | 8891.84 | 393.74 |
| 526 | NENs | 0.743 | 0.257 | 1521.41 | 0.09 | 3263.34 | 142.29 |
| 844 | Neuroglia | 0.724 | 0.276 | 1501.25 | 0.06 | 3493.49 | 120.25 |
| 322 | RBC | 0.913 | 0.087 | 336.07 | 0.02 | 3792.37 | 211.34 |
| 5867 | SMC | 0.937 | 0.063 | 478.69 | 0.11 | 817.77 | 10.68 |
| 210 | Unknown | 0.91 | 0.09 | 1302.01 | 0.12 | 4767.06 | 328.96 |

**Table 1:** Top expressed genes by the Calcb-expressing clusters and the putative MENs cluster in data from May-Zhang et al, along with top-expressed genes in the MENs cluster in our data. Highlighted genes between MENs clusters from our data and data from May-Zhang et al and from Drokhlyansky et al show similar gene expression profiles between the clusters.

**Calcb+ cluster corresponding to Ileum cluster in May-Zhang et al. which expresses Calcb, Calb2, Nmu, Pcdh10, Cysltr2**

| Gene | FDR | summary.logFC |
|---|---|---|
| Zfp804a | 0 | 5.314372906 |
| Pcdh10 | 0 | 2.727131014 |
| Gpr149 | 0 | 2.928565459 |
| Necab1 | 0 | 3.126115371 |
| Grin3a | 0 | 2.477871618 |
| Fam19a1 | 0 | 4.658451729 |
| Ano2 | 0 | 3.443066822 |
| Ntrk3 | 0 | 4.380099903 |
| Cntn5 | 0 | 6.308342607 |
| Ccbe1 | 0 | 3.076522552 |
| Zbtb7c | 0 | 2.656109043 |
| Astn2 | 0 | 3.366649926 |
| Dapk2 | 0 | 1.703870376 |
| Nrxn3 | 0 | 5.160806272 |
| Dgkg | 0 | 4.15397884 |
| Galr1 | 0 | 1.673088209 |
| Nmu | 0 | 3.521779106 |
| Hspb8 | 1.42E-236 | 1.244433448 |
| Cpne4 | 0 | 4.163433255 |
| Tmeff2 | 0 | 3.49539549 |
| Myl1 | 0 | 3.653772641 |
| Calcb | 0 | 3.200679451 |
| Cysltr2 | 0 | 2.487957211 |
| Clstn2 | 0 | 2.655215539 |
| Robo1 | 1.98E-297 | 1.9960685 |
| Hunk | 0 | 1.918886092 |
| Snhg11 | 0 | 2.098574269 |
| Aff2 | 0 | 2.087576143 |
| Otud7a | 3.38E-263 | 2.545208984 |
| Pcdh9 | 0 | 4.216229123 |
| Htr3b | 0 | 1.553388667 |
| Scn11a | 0 | 1.908976823 |
| Iqgap2 | 0 | 1.75599309 |
| Rbfox1 | 0 | 4.70241843 |
| Tcf7l2 | 0 | 2.84136652 |
| Tbx2 | 0 | 1.871954414 |
| Robo2 | 0 | 4.088780154 |
| Kcnt2 | 0 | 3.926721104 |
| Srrm4 | 2.23E-261 | 2.303709088 |
| Plxna4 | 0 | 3.31892195 |
| Sgcz | 0 | 4.739595343 |
| 9530059O1 | 0 | 3.600146025 |
| Sox4 | 7.89E-259 | 1.211156367 |
| Hcn1 | 8.40016528 | 1.861283774 |
| Scube1 | 0 | 2.077291022 |
| Boc | 5.82E-199 | 0.854637195 |
| Cbln2 | 0 | 2.229769856 |
| Kcnn3 | 0 | 1.899022238 |
| Rph3a | 0 | 1.741082515 |
| Cpne8 | 0 | 2.360533782 |
| Calcrl | 1.12695240 | 1.788279732 |
| Lingo2 | 0 | 4.714175039 |
| Slc4a4 | 0 | 2.575716085 |
| Cux2 | 0 | 2.614397251 |
| Chat | 1.62E-252 | 1.075493057 |
| Htr3a | 0 | 1.82434275 |
| Hs3st2 | 1.18369186 | 1.564650584 |
| Cdh8 | 0.00E+00 | 2.544193879 |
| Gm45881 | 0 | 2.045509731 |
| Cdh6 | 0 | 2.996825969 |
| Gm21847 | 9.83E-301 | 2.668713923 |
| Frmd4b | 0 | 2.260822381 |
| Ndrg4 | 5.28E-172 | 1.033574626 |
| Efr3a | 0 | 2.922824682 |
| Syt2 | 0 | 1.996197662 |
| Pak7 | 2.65E-280 | 1.50563649 |
| Lhfpl2 | 0 | 1.681014338 |
| Adgrg6 | 0 | 3.290333725 |
| Spock3 | 0 | 3.61350554 |
| Smad6 | 3.00E-194 | 0.829179685 |
| Zeb2 | 1.58E-304 | 2.305509194 |
| Hs3st5 | 0 | 2.356357004 |
| Kcnh1 | 1.71E-232 | 1.145753317 |
| Scn9a | 0 | 1.985971133 |
| Ppp3ca | 0 | 2.635476275 |
| Agrn | 0 | 2.149338255 |
| Cntnap2 | 0 | 3.976062671 |
| Ngfr | 0 | 2.251817207 |
| Tnr | 0 | 2.255094218 |
| Mcf2l | 6.65E-174 | 0.813444301 |
| Kcnb2 | 0 | 4.170860912 |
| Gpr85 | 0 | 1.49922453 |
| Nell1 | 0 | 2.464621478 |
| Arhgef3 | 0 | 1.961289599 |
| Syt15 | 3.73E-275 | 1.216092892 |
| Lsamp | 7.25E-253 | 2.548136091 |
| 5330434G0 | 4.84E-185 | 2.358912068 |
| Ptger3 | 1.23E-256 | 1.370213397 |
| 1700111E1 | 4.26E-258 | 1.561674061 |
| Mgat4c | 0 | 3.086139006 |
| Fgf14 | 0 | 3.456636428 |
| Mafg | 2.98E-74 | 1.352690002 |
| Kif26b | 0 | 2.174461286 |
| Plppr5 | 1.60E-265 | 1.61715395 |
| Srrm3 | 5.65E-165 | 1.711562875 |
| Hmbox1 | 7.78E-79 | 0.6148506 |
| Rbfox3 | 1.95E-259 | 1.230611205 |
| Ncam2 | 0 | 2.906633451 |
| Slc8a1 | 0 | 2.937485628 |

**Calcb+ cluster corresponding to Ileum cluster in May-Zhang et al. which expresses Calcb, Calb2, Sst**

| Gene | FDR | summary.logFC |
|---|---|---|
| Nlgn1 | 1.56E-227 | 4.503778932 |
| Plxna4 | 5.40E-170 | 3.513261638 |
| Pcdh15 | 6.75E-269 | 4.552606891 |
| Nrg3 | 9.23E-246 | 4.051012028 |
| Eml6 | 1.02E-166 | 3.739252675 |
| Ryr2 | 4.15E-122 | 2.724692337 |
| Nrxn3 | 1.20E-156 | 4.845162261 |
| Rbfox1 | 0.00E+00 | 4.833341666 |
| Sst | 6.45E-135 | 5.862426929 |
| Chsy3 | 4.53E-122 | 3.602794203 |
| Gria2 | 3.32E-126 | 3.083755369 |
| Ptprd | 3.17E-139 | 2.906173926 |
| Pde4b | 3.69E-210 | 3.947198917 |
| Fam19a2 | 8.41E-100 | 3.227261386 |
| Mdga2 | 2.01E-132 | 3.667573403 |
| Snhg11 | 6.24E-112 | 4.344115315 |
| Ralyl | 1.59E-128 | 3.809356707 |
| Bnc2 | 1.58E-137 | 3.131344621 |
| Robo1 | 8.43E-84 | 2.405353602 |
| Nrxn1 | 1.51E-150 | 4.662080188 |
| Pcsk2 | 5.79E-169 | 3.258681935 |
| Syn2 | 1.17E-132 | 3.619085064 |
| Ntrk3 | 1.05E-108 | 2.657539356 |
| Scube1 | 1.03E-95 | 2.265338351 |
| Man2a1 | 9.20E-139 | 3.375536511 |
| Kcnn2 | 7.39E-85 | 2.418320757 |
| Fam19a1 | 1.79E-147 | 3.622699285 |
| Nell1 | 3.21E-100 | 2.793128177 |
| Galntl6 | 1.72E-109 | 3.815091788 |
| Sgcd | 5.24E-120 | 2.88253651 |
| Pde4d | 1.18E-97 | 2.285404689 |
| Slc8a1 | 1.08E-180 | 3.289543486 |
| Tcf4 | 1.52E-159 | 2.592805317 |
| Pak3 | 2.89E-91 | 2.580345128 |
| Kcnma1 | 1.27E-136 | 2.945051283 |
| Adgrb3 | 5.63E-143 | 4.396153392 |
| Unc5c | 2.09E-114 | 3.291001111 |
| Cpne4 | 4.42E-81 | 2.730862875 |
| Meg3 | 7.04E-124 | 4.308953415 |
| Snap25 | 4.22E-135 | 3.917101813 |
| Syt1 | 7.05E-140 | 4.346683131 |
| Grik4 | 8.53E-115 | 2.35016852 |
| Ptprn2 | 1.75E-115 | 3.282060606 |
| Arhgap26 | 1.46E-133 | 2.381300441 |
| Bcl2 | 1.53E-91 | 2.27434062 |
| Ppm1h | 6.46E-139 | 3.466729684 |
| Fgf14 | 3.30E-126 | 4.330212767 |
| Gm12216 | 9.54E-78 | 1.937925878 |
| Camk2a | 1.73E-88 | 2.352765766 |
| Fgf13 | 5.32E-123 | 3.932808506 |
| Ppp3ca | 1.96E-101 | 2.03501536 |
| Adamts9 | 1.24E-60 | 1.993937158 |
| Calcb | 4.08E-76 | 2.288867891 |
| Pcbp3 | 1.17E-136 | 3.893280923 |
| Dlgap2 | 1.21E-96 | 2.470978745 |
| Nkain2 | 1.01E-107 | 2.761261609 |
| Chat | 2.41E-66 | 1.619335139 |
| Gfra2 | 4.04E-106 | 2.379843878 |
| Pde10a | 7.37E-115 | 2.924056301 |
| Ptprr | 1.01E-130 | 3.346248491 |
| Plod2 | 7.61E-111 | 2.338515945 |
| Hs6st3 | 2.71E-78 | 3.027568338 |
| Rbms3 | 2.24E-117 | 1.826125797 |
| Ccbe1 | 2.81E-69 | 1.972703173 |
| Trpm3 | 3.63E-112 | 3.250658339 |
| Frmd4a | 4.22E-106 | 3.509014084 |
| Galnt17 | 5.29E-90 | 2.14675017 |
| Itpr1 | 1.09E-85 | 2.066072139 |
| Srrm4 | 1.17E-82 | 2.302220849 |
| Slc7a14 | 2.27E-93 | 2.415887276 |
| Sema6d | 4.44E-99 | 2.246929266 |
| Pcdh7 | 8.46E-105 | 3.303407952 |
| Nwd2 | 6.48E-47 | 1.597139988 |
| Slit3 | 2.09E-90 | 2.828190275 |
| Asic2 | 3.26E-63 | 3.678490279 |
| Stxbp5l | 1.22E-98 | 2.855884955 |
| Smarca2 | 2.65E-97 | 2.286413106 |
| Ptprt | 3.14E-93 | 2.816471863 |
| Dpyd | 1.72E-91 | 3.059193551 |
| Tln2 | 1.49E-97 | 2.183191767 |
| Satb2 | 1.74E-60 | 2.016803391 |
| Tshz2 | 5.54E-95 | 2.764995869 |
| Plcxd3 | 4.17E-89 | 2.590078205 |
| Tcf7l2 | 1.76E-95 | 2.141154837 |
| Caln1 | 9.30E-64 | 2.04008457 |
| Dgki | 6.15E-121 | 3.238923425 |
| Syt17 | 5.83E-76 | 1.809205079 |
| 9530059O1 | 5.05E-74 | 2.430525524 |
| Unc79 | 7.38E-102 | 2.530692527 |
| Pvt1 | 8.90E-90 | 2.163637608 |
| Mirg | 4.73E-63 | 1.711809423 |
| Ret | 1.04E-68 | 1.699484732 |
| Nrg1 | 5.72E-62 | 3.678288908 |
| Myt1l | 1.56E-120 | 3.008757293 |
| Mapk10 | 9.27E-117 | 2.871225578 |
| Gm15398 | 4.78E-73 | 2.683512094 |
| Arhgef28 | 5.64E-99 | 2.170007537 |
| Zbtb20 | 6.81E-94 | 1.566902687 |
| Dscam | 2.70E-101 | 2.808126988 |

**MENs-like cluster corresponding to Ileum cluster in May-Zhang et al. which expresses Upk3b and Igfbp6**

| Gene | FDR | summary.logFC |
|---|---|---|
| Abi1 | 4.98E-36 | 3.316005072 |
| Dcn | 2.08E-52 | 5.091738535 |
| Col3a1 | 7.95E-31 | 3.039859427 |
| C3 | 1.11E-38 | 4.827731701 |
| Slpi | 2.75E-31 | 4.598725251 |
| Lrrn4 | 7.45E-21 | 2.797813295 |
| S100a6 | 9.36E-29 | 3.230324987 |
| Upk3b | 1.90E-26 | 3.101085936 |
| Gpm6a | 1.13E-24 | 3.730157809 |
| Sparc | 1.37E-28 | 3.316214748 |
| Meg3 | 3.89E-23 | 2.952545493 |
| Igfbp6 | 4.99E-26 | 3.61531349 |
| Agap1 | 4.03E-21 | 2.507795342 |
| Pbx1 | 4.47E-25 | 2.530783608 |
| Efemp1 | 2.67E-22 | 2.915648257 |
| Gas6 | 2.54E-20 | 2.742900314 |
| Muc16 | 2.43E-21 | 3.168081514 |
| Gas1 | 2.13E-23 | 2.708510446 |
| Dpp4 | 4.03E-21 | 2.876742349 |
| Cobll1 | 1.10E-20 | 2.231586143 |
| Rarres2 | 1.64E-20 | 3.226437143 |
| Myo1d | 6.05E-21 | 2.158327438 |
| Tbce | 1.69E-12 | 3.139158995 |
| Il6st | 6.05E-21 | 2.339891442 |
| Nfib | 7.62E-23 | 2.57202827 |
| Crip1 | 7.17E-19 | 2.764477578 |
| Sema5a | 1.97E-19 | 2.852546212 |
| Fmo2 | 1.80E-18 | 3.253371192 |
| Ezr | 2.84E-20 | 2.166036039 |
| Ahnak | 6.05E-21 | 2.714253195 |
| Bnc2 | 6.54E-20 | 3.061680372 |
| Krt19 | 6.15E-20 | 2.380492921 |
| Crim1 | 3.00E-20 | 2.434254471 |
| Plxna4 | 2.07E-19 | 2.707560817 |
| Sox6 | 3.93E-19 | 2.803585619 |
| Aebp1 | 1.57E-19 | 2.548178022 |
| Cfh | 9.08E-18 | 2.422456138 |
| Col1a1 | 2.72E-13 | 2.033502612 |
| Wt1 | 2.03E-17 | 2.196826568 |
| Ccdc171 | 1.09E-14 | 2.022034827 |
| Cd2ap | 2.30E-18 | 2.290803108 |
| Arhgap29 | 3.72E-18 | 2.409211927 |
| Ano1 | 2.38E-18 | 3.039709984 |
| Col1a2 | 5.02E-13 | 2.132928671 |
| Cav1 | 5.10E-18 | 2.235452113 |
| Bicc1 | 2.27E-19 | 2.486459956 |
| Msln | 2.26E-14 | 1.829701191 |
| Dlc1 | 1.57E-18 | 2.561801087 |
| Upk1b | 1.96E-17 | 1.970013811 |
| Sulf1 | 3.39E-17 | 2.542517734 |
| Vim | 1.68E-16 | 2.45290991 |
| Sema3c | 1.17E-17 | 2.547300887 |
| Ifitm3 | 6.57E-15 | 2.071790767 |
| Rspo1 | 1.96E-17 | 1.964755911 |
| Aldh1a2 | 4.16E-15 | 2.160694165 |
| Serping1 | 4.91E-14 | 2.101997058 |
| Timp2 | 8.53E-10 | 1.596556868 |
| Sdc4 | 9.47E-13 | 1.640281568 |
| Cd34 | 2.04E-10 | 1.324668078 |
| Serpinh1 | 2.72E-11 | 1.617681506 |
| Heg1 | 1.56E-17 | 1.795303412 |
| Arhgap18 | 1.69E-15 | 2.494302228 |
| Igfbp7 | 3.27E-15 | 2.01497489 |
| Wdr17 | 9.82E-15 | 3.265554301 |
| Anxa2 | 1.55E-16 | 1.783316817 |
| Celf2 | 8.01E-17 | 2.016354207 |
| Slc4a4 | 2.64E-14 | 2.058956296 |
| Mast4 | 7.11E-16 | 2.591248418 |
| Bcam | 3.45E-09 | 1.453472527 |
| Cobl | 3.19E-14 | 2.113252622 |
| Airn | 1.00E-16 | 2.159009448 |
| Rbbp8 | 1.25E-07 | 1.460802144 |
| Piezo2 | 6.47E-15 | 2.155464659 |
| Wls | 8.02E-16 | 1.68899014 |
| Pcolce | 2.35E-09 | 1.299424275 |
| Prkg1 | 6.92E-15 | 2.705845188 |
| Syne2 | 1.79E-15 | 2.085470282 |
| Zfpm2 | 7.87E-15 | 2.689571135 |
| Adamtsl1 | 1.19E-15 | 2.937092165 |
| Gm26809 | 0.0001518 | 1.800107543 |
| Chst4 | 1.09E-13 | 1.534679673 |
| Ctdspl | 1.02E-13 | 1.587097481 |
| Ptgis | 1.96E-10 | 1.436656186 |
| Kcnab1 | 4.33E-13 | 2.347466136 |
| Nav2 | 2.26E-14 | 2.852022736 |
| C1ra | 6.88E-07 | 1.318506858 |
| Wnt5a | 4.94E-12 | 1.540387353 |
| Dab2 | 2.15E-13 | 1.86057873 |
| Ankrd12 | 0.0001106 | 1.841901593 |
| Cavin2 | 6.09E-12 | 1.976778556 |
| Rnase4 | 1.39E-11 | 1.688218465 |
| Meis1 | 1.26E-13 | 1.908870378 |
| Anxa1 | 1.86E-12 | 1.531275304 |
| Ildr2 | 9.47E-13 | 1.550620875 |
| Slc9a3r1 | 1.86E-10 | 1.325162783 |
| 9530026P05 | 2.17E-13 | 2.515612487 |
| Man1a | 3.13E-12 | 1.539400492 |
| Gpc3 | 1.20E-12 | 1.790707073 |
| Col14a1 | 3.60E-11 | 2.039467045 |

**MENs Cluster (Our Data)**

| Gene | FDR | summary.logFC |
|---|---|---|
| Bcam | 0 | 1.310889756 |
| Mgst1 | 0 | 1.466554292 |
| Krt18 | 1.02E-252 | 0.722876212 |
| Igfbp6 | 0 | 4.277079428 |
| Rspo1 | 0 | 1.356888212 |
| Gpm6a | 0 | 2.007236002 |
| Upk3b | 0 | 2.136663175 |
| Aebp1 | 0 | 2.502295384 |
| Dcn | 0 | 4.082052904 |
| Slpi | 0 | 4.928306229 |
| Rarres2 | 0 | 4.202866323 |
| Efemp1 | 0 | 1.672848073 |
| Aldh1a2 | 4.60E-277 | 0.796263425 |
| Fmo2 | 0 | 1.582315372 |
| Upk1b | 0 | 1.081912196 |
| Gas1 | 0 | 1.464482623 |
| Krt7 | 0 | 1.315380677 |
| Crip1 | 0 | 4.56883561 |
| Clu | 0 | 1.502811937 |
| C3 | 0 | 3.494711523 |
| Dpp4 | 1.12E-210 | 0.556370013 |
| Lrrn4 | 1.58E-271 | 0.736486954 |
| Nkain4 | 0 | 1.298360281 |
| S100a6 | 0 | 4.128650316 |
| Krt19 | 0 | 1.900927424 |
| Cfb | 8.74E-203 | 0.608707666 |
| Gabarapl1 | 5.26E-250 | 0.709728291 |
| Lgals3 | 0 | 1.26072921 |
| Krt8 | 1.39E-245 | 0.689105423 |
| Tm4sf1 | 0 | 1.367716307 |
| Slc39a8 | 7.17E-197 | 0.489802511 |
| Sema3c | 2.63E-255 | 0.71554962 |
| Pcolce | 0 | 1.823133603 |
| Gsta4 | 1.09E-226 | 0.62733526 |
| Ezr | 0 | 1.019712533 |
| Cldn15 | 4.76E-291 | 0.798661223 |
| Serpinb6b | 2.66E-212 | 0.562587863 |
| Nbl1 | 0 | 1.239754358 |
| Slc16a1 | 2.28E-244 | 0.65901005 |
| Smim1 | 2.76E-245 | 0.659125894 |
| Snhg18 | 3.11E-181 | 0.475288967 |
| Cavin3 | 6.15E-293 | 0.855731534 |
| Gpc3 | 0 | 1.393181642 |
| Serpinb1a | 3.54E-178 | 0.472277374 |
| Chst4 | 3.42E-232 | 0.599828027 |
| Esam | 1.50E-188 | 0.47512383 |
| Ptgis | 5.59E-231 | 0.598750529 |
| Slc9a3r1 | 7.03E-265 | 0.883988091 |
| Ecm1 | 1.88E-172 | 0.514360839 |
| Igfbp7 | 0 | 2.347561986 |
| Msln | 0 | 2.266588157 |
| Gpx3 | 0 | 2.209730215 |
| Col14a1 | 1.17E-163 | 0.423875564 |
| Anxa1 | 4.24E-246 | 0.691007825 |
| Bace2 | 7.94E-223 | 0.603024779 |
| Cfh | 0 | 1.190621234 |
| Tmem176a | 1.85E-170 | 0.915766916 |
| Tmsb10 | 0 | 2.299200297 |
| Timp2 | 0 | 1.918063461 |
| Pdpn | 2.36E-252 | 0.846386263 |
| Prss23 | 0 | 0.980677037 |
| Cd151 | 2.85E-279 | 0.795276372 |
| P3h4 | 8.94E-212 | 0.512747523 |
| Cd200 | 0 | 0.975177086 |
| Ltbp4 | 1.80E-268 | 0.859534481 |
| Gstt1 | 2.26E-142 | 0.347320974 |
| Ssbp3 | 1.42903506 | 0.888557965 |
| Tmem98 | 3.90E-240 | 0.62900078 |
| Spint2 | 1.42E-186 | 0.478143896 |
| Nupr1 | 3.42E-226 | 0.705142527 |
| Basp1 | 6.06E-239 | 0.625848924 |
| C1ra | 2.06E-263 | 0.752700756 |
| Serpinh1 | 4.94065645 | 1.065045464 |
| Adamts2 | 9.08E-129 | 0.322228993 |
| Dmkn | 3.43E-187 | 0.652807485 |
| C1s1 | 2.25E-276 | 0.820741368 |
| Sparc | 0 | 1.954167099 |
| Espn | 3.75E-166 | 0.392816171 |
| Anxa2 | 0 | 1.579365041 |
| Krt14 | 6.29E-117 | 0.368125408 |
| Sdc4 | 0 | 1.913582811 |
| Serping1 | 0 | 1.802367854 |
| Pmp22 | 5.43E-232 | 0.6401978 |
| Cplx2 | 1.77E-251 | 0.917909984 |
| Il18 | 6.70E-154 | 0.396176068 |
| Il6st | 1.73E-262 | 0.71225818 |
| Meg3 | 1.02E-173 | 0.558817278 |
| Cald1 | 2.88E-262 | 0.803200156 |
| Mt2 | 1.37E-218 | 0.736087263 |
| Nenf | 7.12E-165 | 0.869917467 |
| Col6a1 | 2.61E-252 | 0.693856752 |
| Rnase4 | 0.00E+00 | 1.162670662 |
| Vim | 0 | 1.805718628 |
| F11r | 6.92E-219 | 0.535570277 |
| Fam180a | 2.29E-131 | 0.294690768 |
| Npdc1 | 2.34E-164 | 0.415479342 |
| Cd2ap | 2.16E-225 | 0.595177252 |
| Pamr1 | 3.15E-177 | 0.443000918 |
| Stk26 | 4.71E-112 | 0.258746973 |

**Cluster annotated as Mesothelial (Drokhlyansky et al data)**

| Gene | FDR | summary.logFC |
|---|---|---|
| Lrrn4 | 5.4657E-206 | 16.886354 |
| Rspo1 | 3.6499E-210 | 9.4753895 |
| Wt1 | 1.5809E-227 | 10.114356 |
| Upk3b | 5.8625E-282 | 8.9449835 |
| Bnc1 | 4.04195E-85 | 15.045354 |
| Wt1os | 2.3697E-123 | 9.9661905 |
| Upk1b | 1.277E-215 | 8.0787953 |
| Aldh1a2 | 1.5275E-143 | 6.3791433 |
| Msln | 8.3474E-159 | 7.3877455 |
| Igfbp6 | 1.7674E-307 | 7.7565747 |
| Lvrn | 4.18762E-92 | 7.8098721 |
| C3 | 0 | 8.057149 |
| Muc16 | 7.8925E-269 | 7.3116341 |
| Gpm6a | 0 | 6.5940179 |
| Cybrd1 | 4.39858E-60 | 9.0017763 |
| Fam180a | 1.90191E-30 | 13.526168 |
| Myrf | 1.71228E-56 | 8.6635402 |
| Aebp1 | 4.5764E-247 | 6.6271763 |
| Crb2 | 2.91628E-52 | 7.3719161 |
| Zdbf2 | 3.25895E-75 | 7.0872463 |
| Sult5a1 | 2.85305E-24 | 13.31773 |
| Cfh | 4.6319E-250 | 5.7592104 |
| Wdr17 | 3.1501E-282 | 6.6320991 |
| Ildr2 | 3.4601E-136 | 6.7198792 |
| Slpi | 1.7684E-135 | 7.8326894 |
| Fmod | 2.27977E-40 | 8.7786829 |
| Cpxm1 | 7.29311E-19 | 12.558464 |
| Ifi205 | 2.62948E-18 | 12.864742 |
| A730046J19 | 8.01459E-17 | 12.793582 |
| Eya4 | 4.5788E-136 | 6.8974953 |
| Nkain4 | 1.08326E-83 | 5.7593624 |
| Col8a2 | 5.81471E-44 | 5.2884497 |
| Zfp185 | 1.12186E-43 | 7.3412329 |
| Dcn | 6.4597E-292 | 5.6187432 |
| Gm20400 | 7.4045E-135 | 7.0357895 |
| Gas1 | 2.6756E-182 | 5.7763744 |
| Cdh3 | 2.1841E-97 | 6.6406251 |
| Efna5 | 1.6397E-264 | 4.7533052 |
| Tmem151a | 4.7978E-88 | 6.2910937 |
| Bnc2 | 5.1982E-260 | 5.1670469 |
| Gm12381 | 7.0467E-102 | 7.6933315 |
| Serpinb6b | 1.16024E-44 | 5.3575251 |
| Plxna4 | 1.3438E-213 | 5.1141234 |
| Efemp1 | 6.3046E-195 | 5.3926198 |
| Gm765 | 4.94209E-43 | 6.7967853 |
| Fmo3 | 1.17753E-20 | 8.4299998 |
| Fmo2 | 2.9871E-243 | 5.3187177 |
| Prss12 | 3.97653E-61 | 5.4000171 |
| Vwa3a | 1.02294E-65 | 5.8157216 |
| Abcb1b | 1.7423E-115 | 6.470465 |
| Rarres2 | 6.6617E-142 | 5.7715796 |
| Lrp2 | 2.01232E-42 | 6.2276692 |
| Cldn10 | 6.74497E-17 | 5.6397899 |
| Spock2 | 1.9494E-138 | 5.1124954 |
| Chrd | 1.6918E-22 | 7.4139259 |
| Ltbp3 | 1.94368E-88 | 5.07873 |
| Pkhd1l1 | 4.889E-120 | 4.4235079 |
| Svopl | 4.52096E-42 | 6.0169611 |
| Ptgis | 1.487E-109 | 5.0355511 |
| Il16 | 1.59107E-97 | 4.7690361 |
| Wnt9a | 1.72493E-27 | 4.850231 |
| Gabra3 | 2.92934E-57 | 4.859601 |
| Medag | 2.3907E-99 | 4.7656656 |
| Atp6v0a4 | 1.46614E-75 | 5.0974539 |
| Rpp25 | 1.286E-19 | 5.4541715 |
| Bcam | 2.8668E-131 | 5.0323969 |
| Sulf1 | 4.0825E-179 | 4.5936014 |
| Evc2 | 1.99898E-30 | 5.6828103 |
| Arhgap28 | 6.6071E-111 | 4.5878415 |
| Adamtsl1 | 6.7922E-140 | 4.3437642 |
| Fam184a | 3.53509E-66 | 5.6445631 |
| Nxph1 | 9.70539E-83 | 6.4656245 |
| Wnt5a | 5.8364E-101 | 4.8191031 |
| Unc13c | 7.65838E-77 | 5.674392 |
| Smtnl2 | 1.59692E-27 | 4.2520936 |
| C1s1 | 7.3878E-125 | 4.5487014 |
| Gata5 | 1.77132E-31 | 5.1168633 |
| Tm4sf1 | 4.496E-113 | 4.5813419 |
| Anxa9 | 9.60666E-21 | 4.8493462 |
| Gpc3 | 4.9665E-99 | 4.7995673 |
| Trpv4 | 7.95066E-16 | 5.3205717 |
| Slit3 | 1.5771E-163 | 3.7434359 |
| Synpr | 1.67047E-53 | 5.2835493 |
| Krt14 | 7.46374E-25 | 5.109502 |
| Ifi204 | 4.97213E-20 | 5.7099144 |
| Isl1 | 1.73444E-33 | 5.327459 |
| Dnajc6 | 1.50196E-72 | 4.1858858 |
| Meis1 | 1.3366E-165 | 3.8477795 |
| Nnmt | 6.20949E-19 | 5.3840664 |
| Meis2 | 1.5618E-191 | 3.1626199 |
| Cd34 | 1.84466E-86 | 3.7841694 |
| Gm20186 | 1.72742E-38 | 4.7291369 |
| Npr1 | 4.88291E-40 | 4.4804827 |
| Gm11747 | 8.57184E-23 | 4.9058649 |
| Mmp16 | 1.46268E-89 | 5.0348242 |
| Loxl1 | 1.00489E-79 | 4.2443546 |
| Ar | 6.54875E-96 | 4.7109788 |
| Rab7b | 3.82339E-37 | 4.627583 |
| Tnfrsf11b | 1.46976E-58 | 5.0928081 |
| Ccdc85a | 1.53629E-69 | 4.4553075 |
