## Supplementary material for "Age-associated changes in lineage composition of the enteric nervous system regulate gut health and disease": Rebuttal to Review version 3

Responses to reviewer provided in blue

Reviewer #1:
Remarks to the Author:
The presence of mesenchymal marker genes in subsets of cells of the ENS in native gut tissue is convincing and the lack of retained fluorescent reporter expression in ENS from the many neural and Cre drivers used is indeed clear. Overall, the manuscript has revealed a new type of cell in the enteric nervous system that appears with age, has a mesodermal origin and show differential regulation to growth factors compared to neural-crest derived neurons. These data are convincing and should be revealed to the general public.

We thank the reviewer for his/her remarks and for accepting our central thesis and main claim that a significant population of enteric neurons in the post-natal gut is derived from a non-neural crest origin. We are glad that with the significant amount of data that we have provided by using various lineage fate mapping models, we were able to convince the reviewer that the mesoderm generates a large population of enteric neurons (MENs), whose proportions rival those derived from the canonical neural crest origin (NENs). The reviewer also agrees that MENs and NENs are responsive to different growth factors; that their proportions alter with age, and in animals and humans with reduced Ret signaling; and further supplementation with the NENs-specific growth factor GDNF reverts age-associated loss of NENs to rescue age-associated dysmotility.

The current state of the manuscript is however not acceptable as it has unsound interpretation of data at many places, most importantly there is no firm connection between the MENs identified in tissue to the scRNA cluster annotated as MENs.

We disagree with the reviewer as in prior and current versions of our manuscript, we demonstrate the expression of Cdh3, Calcb, and Met in the scRNAseq-MENs cluster and have validated the expression of these markers by immunofluorescence labeling in Wnt1-cre:tdTomato-negative non-neural crest-derived neurons (MENs). We have further expanded the validation of 18 additional scRNAseq-MENs markers to include robustly expressed mesenchymal markers Decorin, Aebp1, Slpi, Fmo2, Myl7, Smo, Clic3; neuronally significant markers Stmn2, Elavl2, Hand2, Gpr88, Vsnl1, Tubb2b, Pde10a, Stx3 (several of which are robustly expressed), and other equally important markers such as NT-3, IL-18, and Slc17a9 (VNUT). Along with the validation of the Myh11 marker for these ‘mesenchymal neurons’ identified by May-Zhang et al. (DOI: 10.1053/j.gastro.2020.09.032) in their scRNAseq data, we validate 22 markers that were identified via specific expression in an scRNAseq cluster annotated by us as MENs in the population of non-neural crest-derived mesodermal neurons identified through the various lineage fate mapping strategies. Importantly, the *in vivo* validation of these *in silico*-predicted markers, should be considered the gold standard of validation, and should be sufficient to demonstrate that these cells (A) are indeed MENs, (B) are intra-ganglionic, and (C) express a cohort of marker genes that are distinct from neural crest-derived cells. Recent studies have classified various enteric neuronal subsets often on the validation of 2 - 3 markers, and by contrast, we use 22 markers to provide a link between MENs and their molecular signature as discerned by single cell transcriptomics. We believe that this presents a strong case for our inference that the scRNAseq cluster annotated as MENs describes the molecular signature of MENs identified in tissue.

There are several reasons for this:
1) The most clear mismatch between "tissue-MENs" and the "scRNA-seq MENs" is the obvious neuronal expression in the first (high expression of Elavl2-4), but low abundance/level in the latter. Elavl3 and 4 are found in a minor proportion of "scRNA-seq MENs", while the claimed new marker Elavl2 also is very low and does not provide more strength to the validation. The absolute number of cells expressing Elavl3/4/2, Stmn2 (and a few other neuronal markers) in "scRNA-seq MENs" is similar or smaller than the cells within the NENs, yet the NENs dataset contains 77 cells, while the MENs has 2223 cells, thus 28 times more cells! It is not valid to claim that either the V1 or V3 10x platform isn't sensitive enough to capture highly expressed neuronal genes such as Elavl2-4, as previous efforts (FACS sorted, whole cells without RNAse inhibitors) using V1 captured 20-50 transcripts per enteric neuron (Zeisel et al., 2018; mousebrain.org; enteric neurons). Further, the number of Snap25+ cells in "scRNA-seq MENs" is similar to cells expressing

other neuronal markers, yet Snap25 is claimed negative, while Elavl2-4 are claimed expressed. Snap25 is expressed in the "MENs" of the May-Zhang atlas, but at the same time claimed equivalent to the "scRNA-seqMENs". Expression of Actl6b is also not shown in the "scRNA-seqMENs", yet the Baf53b-Cre line is used to claim the gene's limitation to NENs.

We wish to respond to this comment from the reviewer on the annotation of the MENs scRNAseq cluster by providing the following rationale.

1. Despite the validation of 22 scRNAseq marker genes (several of which are known mesenchymal ***and*** neuronal marker genes) for the MENs cluster in tissue, the reviewer has centered on the low but detectable expression of Elavl2/3/4 genes in the scRNAseq MENs cluster to suggest that our annotation of the scRNAseq MENs cluster is incorrect.

    There are two main reasons why the detectable expression of the Elavl gene transcripts may be lower in the MENs scRNAseq cluster than in the NENs cluster.

    a. Single cell or nucleus RNA sequencing methods are biased towards sequencing transcripts that are abundant relative to all other transcripts for that individual cell (detection and amplification bias). Thus, while the same transcript may be equally expressed at an absolute level in two different cells, it will be more readily sequenced and detected in the cell where the transcript is <u>*relatively*</u> more abundant. In this case, even if the expression of the Elavl gene transcripts is assumed to be equal between MENs and NENs, these transcripts will be more readily detected in NENs, if their relative abundance is higher in NENs than in MENs. Our data shows that MENs have a higher UMI per cell count, suggesting that even if the absolute expression of the Elavl genes is similar between NENs and MENs, the relative abundance of these gene transcripts in MENs would be lower than in NENs - making the detection of these transcripts more probable in NENs than in MENs. It is well known that the lack of detectable expression of transcripts in a cell cluster does not equate to lack of expression of those genes in that cell population. Hence, validation of the annotation of a scRNAseq cluster has to be performed by showing the corresponding protein expression of a number of genes (and not fixate on 1 or 2 genes) in the cell population, and we have performed such validation for ***22 different markers*** in our study that has allowed us to be confident in linking the identity of the scRNAseq cluster annotated as MENs to the mesoderm-derived neurons observed in tissue.

    b. In the Zeisel et al. (DOI: 10.1016/j.cell.2018.06.021) data that transcriptomically describes NENs at the P21 age, neurons show significant diversity in transcript abundance of the Elavl genes. After enriching for neural crest-derived cells, Zeisel et al.'s study showed that large populations of neurons (most neurons in the ENT3, 4, and 5 clusters) showed detectable

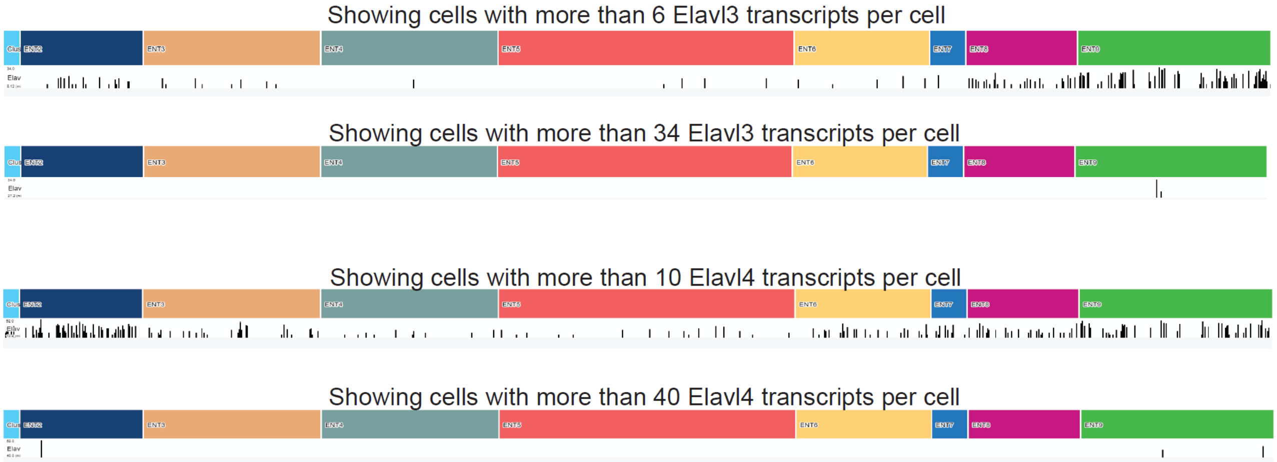


**Figure A:** Data from Zeisel et al. shows that the abundance of Elavl3 and Elavl4 transcripts, which encode for HuC and HuD proteins, is variable between different classes of Wnt1-cre:tdTomato-expressing neural crest-derived neurons.

| | Numbers of cells | Numbers of cells showing expression of gene | Percent of cells expressing gene | Average transcript count for all cells | Average transcript count in cells showing detectable expression |
|---|---|---|---|---|---|
| **Elavl2** | | | | | |
| **Mesenchymal neurons (MENs)** | 94 | 4 | 4.26 | 0.06 | 1.5 |
| **All other neurons (NENs)** | 11119 | 2816 | 25.33 | 0.47 | 1.87 |
| **Elavl3** | | | | | |
| **Mesenchymal neurons (MENs)** | 94 | 11 | 11.7 | 0.17 | 1.45 |
| **All other neurons (NENs)** | 11119 | 7758 | 69.77 | 2.61 | 3.73 |
| **Elavl3** | | | | | |
| **Mesenchymal neurons (MENs)** | 94 | 24 | 25.53 | 0.33 | 1.29 |
| **All other neurons (NENs)** | 11119 | 9296 | 83.6 | 10.8 | 12.92 |

**Figure B**: Analyses of May-Zhang et al. data shows that the Elavl gene transcripts are detected less frequently in MENs, when compared to all other neurons. In addition, data also shows that these transcripts are not detected as a high copy number transcript in all neuronal populations.

expression of less than 6 Elavl3 transcript per neuron, while other neurons showed detectable expression of more than 25 transcripts per neuron (Figure A). This observation extends to Elavl4 gene transcripts, where many neurons from the ENT3, 4, and 5 clusters showed detectable expression of less than 10 transcripts while some other neurons in their database showed expression of more than 30 - 40 transcripts per neuron (Figure A). These data first help establish that expression of Elavl genes is variable even within the canonical NENs population. Second, the four-fold difference in transcript abundance within various NENs subtypes does not correlate with our observed expression of Hu proteins by immunohistochemistry. This is not surprising since it is well known that transcript expression and protein expression poorly correlate for multiple tissue and cell types, including neurons (Jiang et al, DOI: 10.1016/j.cell.2020.08.036; Moritz et al, DOI: 10.1111/jnc.14664). Third, MENs have significantly different developmental origins and biology than the canonical neurons and hence it cannot be assumed that their absolute expression of Elavl genes would be similar to that of NENs. Indeed, our reanalysis of the May-Zhang et al. data (Figure B) shows that not all neurons show expression of the Elavl genes, with the cluster of 'mesenchymal neurons' or MENs in this dataset shows much sparser expression of all the Elavl genes. Only ~4%, ~11%, and ~25% of cells of the mesenchymal neuronal cluster in May-Zhang et al. data show detectable expression of the genes Elavl2, Elavl3, and Elavl4. This is in contrast to the larger population of canonical neurons, whose ~25%, ~70%, and ~83% of cells express these three genes. The correct inference here is that neurons in the mesenchymal neuronal cluster (MENs) have lower abundance of these gene transcripts, due to which they remain undetected in more cells of this cluster, when compared to the canonical neurons. Taken together, this argues that just because we observe a similar profile of Hu expression in myenteric neurons, it cannot be assumed that MENs and NENs have similar absolute abundance of the Elavl gene transcripts. Additionally, it is important to remember that, in the context of scRNAseq which is an inherently noisy assay, the expression (or lack thereof) of a given gene of interest should *always* be verified validated by a more sensitive and robust method. The interpretation of any single gene expression should be supported by additional validations, and this is why we have performed this assay for Elavl (IHC) and our additional 22 marker genes.

In addition to our validation of a significant number of MENs markers identified in scRNAseq data, these two issues, which highlight why it would be incorrect to expect or assume equal transcript abundance (*absolute and relative*) of the Elavl genes between MENs and NENs, provide the rationale for why it is erroneous to argue that the low detection of the Elavl genes in the scRNAseq MENs' data suggests a mismatch between MENs and their scRNAseq cluster.

2. The reviewer next asks about the expression of gene Snap25 in the MENs population, and particularly why its sparse expression in MENs scRNAseq cluster should not be held to the same argument applied for the Elavl genes. Expression of the Snap25 gene, among other genes such as Ret (which we show is specific to NENs), has been used by multiple groups in annotating the population of enteric neurons in their single cell/nuclear transcriptomic data. We had hypothesized then that the absence of detectable expression of these genes might be the reason for why the true neuronal annotation of cells we annotate as MENs was not correctly applied in prior reports (such as Drokhlyansky et al. DOI: 10.1016/j.cell.2020.08.003; and Elmentaite et al. DOI: 10.1038/s41586-021-03852-1). To test this hypothesis, we studied whether the Snap25 gene and its translated protein are expressed by MENs, and found that most cells within the MENs scRNAseq cluster did not show detectable expression of this gene, and that Snap25 protein expression was not observed in MHCst-expressing MENs. Then, we validated the lack of expression of Snap25 transcripts in a population of MENs by using a Snap25 knock-in reporter mouse that showed that a significant population of Hu-expressing and MHCst-expressing myenteric neurons did not express the Snap25 gene. Lack of Snap25 expression in a population of myenteric neurons has been previously documented in human tissue (Barrenschee et al. DOI: 10.1007/s00418-015-1310-x), which further validates our observations. This is in contrast to the case we discussed earlier with the Elavl genes, where their transcripts were sparsely detected in our scRNAseq data, but their robust protein expression was confirmed (suggesting that the Elavl genes are expressed in MENs but are not detected with scRNAseq). The lack of transcript and protein expression of Snap25 in a large population of MENs thus is different from cases where transcript expression may be undetected but protein expression can be observed.

However, at the same time, we continue to be careful by noting that Snap25 expression is lacking in a population of MENs (lines 402 - 403), and do not make a case that all MENs are devoid of Snap25 transcript and protein expression. It is certainly plausible that a small subpopulation of MENs do express the translated product for the Snap25, given that both our data and the data from May-Zhang et al. show detectable presence of Snap25 transcripts in MENs cluster.

3. The reviewer then asks why we do not mention the gene Actl6b (that codes for the protein Baf53b) in our analyses of the scRNAseq dataset. We did not detect Actl6b expression in the MENs cluster in our scRNAseq data. However, for the reasons explained above, the lack of detectable expression of the Actl6b gene does not equal the lack of expression of its translated product Baf53b. We found no commercially available antibodies that can definitively distinguish between the neuronal Baf53b and the more generally expressed Baf53a (translated product of the gene Actl6a), thus limiting us from commenting on the expression of this gene and its gene product in MENs. What we do note though is that the Baf53b-cre:tdTomato mouse (which is a BAC transgenic mouse line) does not label all myenteric neurons (which the reviewer agrees to below) and that all neurons sequenced by flow sorting the Baf53b-cre:tdTomato-expressing neurons at the P21 stage do express Snap25, which we show is expressed predominantly by NENs.

These arguments provide strong justification and extensive experimental validations for why our annotation of the MENs scRNAseq cluster is correct. Our study has provided strong linkage between the molecular identity of these neurons as discerned by scRNAseq and their presence in tissue by providing immunohistochemical validation of a large collection of markers, and by providing a nuanced interpretation of the transcriptomic data.

Correct interpretation of single cell/nucleus transcriptomic data relies on an understanding that not all transcripts of a cell can be sequenced and detected, and thus absence of the expression of transcripts in a cell does not imply absent gene expression. Thus, ascribing molecular identity to a cluster does not need to meet a threshold of whether all or majority of cells of that cluster show detectable expression of a particular gene. Instead, cluster identity is defined by whether a sufficiently large set of gene transcripts show detectable presence in that cluster, which can be used to establish that cluster’s unique identity, and whether the expression of those genes can be validated in tissue by other methods, including microscopic observation of protein or transcript expression. In our report, we have faithfully followed the correct methodology for annotating the MENs scRNAseq transcriptomic cluster.

2) The markers to identify MENs in the scRNA-seq tissue are Calcb, Met and Cdh3. Yet, the expression of the three is very sparse in the "scRNA-seq-MENs".

We again wish to reiterate our earlier point on the correct way to ascribe molecular identity for a cell cluster. Our report details the following chronology: Calcb and Met co-expression was found in a cluster that showed expression of Cdh3 and of various mesenchymal and neuronal markers. Upon validation, the expression of this set of markers (consisting of 22 gene products) was found to link this single cell transcriptomic cluster to the Hu-expressing neurons that did not express tdTomato in the neural crest fate mapping mouse model. No other cluster of cells identified in our scRNAseq experiment shows the co-expression of the 22 MENs-marker genes that were validated in tissue. In addition, while expression of Calcb and Met was restricted to a smaller number of cells in this cluster, ~30% of the cells in the MENs cluster showed presence of Cdh3 transcripts.

3) The proportion of "scRNAseq-MENs" is very much higher than expected. At this stage, authors find that 50% of the ENS neurons are MENS and 50% NENs. 28 times more cells are assigned a MEN identity than a NEN identity. Additionally, if the cells are larger, they are usually harder to maintain in good health, and hence would be lost in dissociation (spinal motor neurons are notoriously difficult for this reason).

The reviewer takes issue with the under-representation of NENs (and the over-representation of MENs) in our scRNAseq data. It is well known that enzyme-based dissociation methods are unequally tolerated by diverse cell types, which is known to cause over- or under-representation of several cell types in scRNAseq (Uniken Venema et al, DOI: 10.1038/s41598-022-13812-y; who showed that dissociation method drives detection and abundance of cells sequenced; Wu et al, DOI: 10.1016/j.stem.2018.10.010 showed the existence of similar dissociation bias in the kidney). Furthermore, the ***size*** of the cell has ***not*** been explicitly described to be a causal reason for this variability. The same is true for neurons, where certain neuronal subtypes may be significantly over- or under-represented in scRNAseq databases, compared to their true representation in tissue. Indeed, it is well known within the single cell community, that single cell assays using enzymatic dissociation should not be used to make determinations about cellular abundance in populations for this very reason. For example: Tiklova et al (DOI: 10.1038/s41467-019-08453-1) showed that specific subpopulations of Dat-expressing neurons in the developing mammalian brain were under-represented in scRNAseq. Thus, it is clear from previously published data that it is well accepted that the proportional size of individual cell clusters in scRNAseq data may not correlate with their true proportions in tissue.

There could be many reasons for why a particular cell type might be over- or under-represented, principle of which are the nature and amount of enzymes used, time allowed for dissociation, and cell tenacity to dissociation-mediated rupture. It is plausible that the expression of ECM proteins such as Decorin (which we have validated) among others by MENs make them more resistant to dissociation-mediated cell rupture, and that the dissociation materials and methods used enrich viable MENs (and not NENs) in the data. This was observed in the Gut Cell Atlas (Elmentaite et al.) data as well, where enzymatic dissociation of the adult human gut tissue was able to detect a significant population of MENs but not NENs.

Given this known limitation for scRNA-Seq analysis, we have chosen to provide the more rigorous and appropriate analysis of cell counting in replicate tissue sections, which has a storied history of being used to determine the cellular composition of a given tissue. It is this assay upon which the determination of MENs/NENs ratios can and should be determined, as performed in our study.

These arguments underscore the fact that over- and under-representation of specific cell types in single cell transcriptomic data is frequently observed in diverse studies and has been discussed at length in many articles as one of the pitfalls of this technique (For example Denisenko et al. DOI: 10.1186/s13059-020-02048-6). It would be unfair to expect our study to buck this known pitfall of this technique.

Hence, the claims that "tissue-MENs" equals "scRNA-seq MENs" should be milden. A possible explanation could be that the small proportion of cells expressing neural markers (3%?) in the "scRNA-seq MENs" indeed correspond to the "tissue-MENs". If authors want to include their scRNA-seq data in the paper, this interpretation is the most likely and could be presented. The manuscript would need to be rewritten accordingly. The new scRNA-seq from 10xv3 is not presented in detail. Does it show more homogeneous expression of markers of "tissue-MENS"? This dataset should no doubt have enough depth to capture neuronal markers including more robustly the ones used to confirm neuron identity in the tissue (Elavl2-4), but also synaptic machinery genes. The relation between "P21-scRNAseq-MENs" and "tissue-MENs" is hard to assess without more information.

As detailed in our earlier responses, in our report, we provide a significant amount of evidence to prove the linkage between the scRNAseq cluster we annotate as MENs, and the MENs we observe in tissue. Our responses to every comment of the reviewer here, and in earlier reviews, comprehensively show this linkage in addition to discussing the technical limitations of the scRNAseq methodology that currently do not allow for a deeper analysis of this newly annotated and newly-described neuronal population. We have been nuanced in our discussion not to over-interpret results and have validated every claim we have made with secondary independent assays.

Further, Supplementary Table 2 in the earlier and the current version of the manuscript, details the top-expressed genes for both the P21 and 6-month-old scRNAseq datasets. Similarly, Supplementary Table 4 details the scRNAseq metrics for both the scRNAseq datasets generated. P21 data analysis in Fig 11 shows the presence of ECT2-expressing MENs that are generated during this critical developmental phase associated with MENgenesis. Supplementary Figure 10 shows that using projectR pipeline and the expression of *Met* gene, the transcriptomic signature of P21 MENs is similar to that of 6 month old MENs. With these analyses of the P21 MENs performed in our report, we disagree with the reviewer that our study has not discussed this dataset in detail. We have previously stated that a deeper transcriptomic analysis of MENs (that can be validated by subpopulation-specific markers) at various time-points in their development requires the generation of dedicated tools and protocols for targeted enrichment of MENs and their deep sequencing, which currently do not exist.

A follow-up question of important consideration is also whether MENs, in case they are represented partial "scRNA-seq MENs" even are to be considered neurons, if they lack expression of synaptic markers. What is a neuron without the regular synaptic machinery? It is possible that the ENS consists of such cells, but should they then be considered neurons?

Hu-expressing cells that are located within the post-natal myenteric ganglia have been historically annotated as neurons. We show that MENs, despite their non-canonical developmental lineage, different biology and lack of Snap25 gene expression, show intra-ganglionic location, Hu protein expression, neurotransmitter expression, and express some synaptic markers such as Syntaxin 3. It is important to note that this definition meets the prior burden for calling these cells neurons, based on the current state of the field. Their specific biological functions, roles in gut function and disease, and physiological properties will likely be the subject of many future studies of these cells. And if, at a subsequent time, a different definition of enteric neuron is established, we will gladly re-evaluate the designation as it makes sense for these cells. In light of this however, the current definitions for enteric neurons by the field suggests that MENs are indeed enteric neurons that may employ alternate synaptic mechanisms as they partake in their neuronal functions. We have also cited prior literature that has reported on the lack of this marker expression in a population of CNS neurons, suggesting that non-canonical synaptic mechanisms occur in other neuronal populations as well.

Further Concerns:

A) It is important to mention that a proportion of NENs are Met+/Ret-. This is the neuron type which is called ENC7 (Morarach et al., 2021, P21 tissue) ENT8, (Zeisel et al., 2018) and EN7 (May-Zhang et al., 2021, P42-52 tissue). Even in the part where the manuscript focuses on P21, the previous findings are not clearly described (very little to no Ret is present in ENT8, while ENT9 and ENT6 clearly show expression of Ret in Supp 10d).

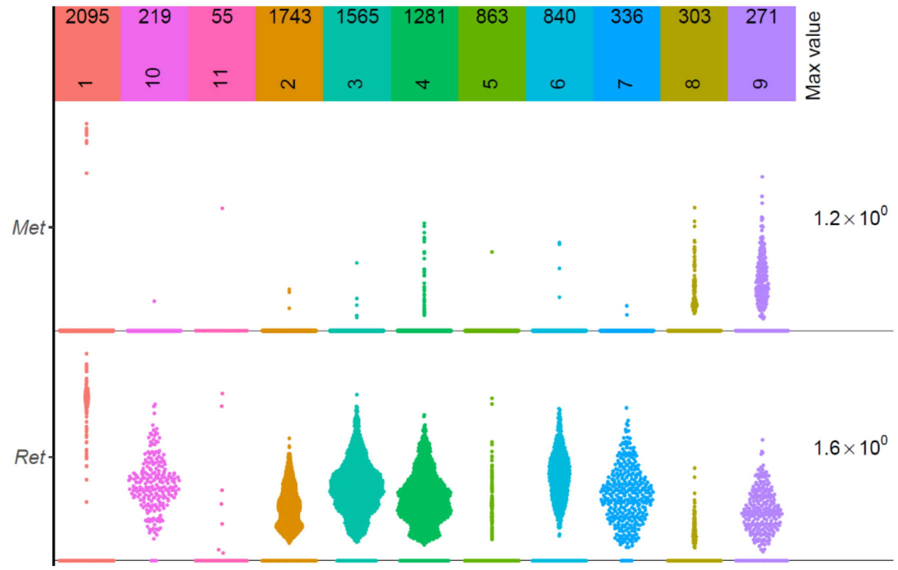


**Figure C:** Analyses of the Morarach et al. data shows that the ENC7 cluster shows detectable presence of Ret gene transcripts, but does not show detectable expression of Met gene. There is no cluster in the Morarach et al. data that shows detectable expression of Met transcripts without detectable expression of the Ret transcripts.

We disagree with the reviewer's statement that the ENT8 cluster from the Zeisel et al. and ENC7 cluster from the Morarach et al. datasets represent a population of Met+ and Ret- neurons. The data from Zeisel et al. (which we have presented in Suppl. Fig 10d), clearly shows that the ENT8 cluster shows detectable expression of both Met and Ret genes (albeit transcripts of both genes are detected in a smaller proportion of neurons). We further show here data (Figure C) from Morarach et al. (DOI: 10.1038/s41593-020-00736-x), which shows that the ENC7 cluster, whose cells express Ucn3 and Cck genes, again shows detectable expression of both Met and Ret gene transcripts. These observations extend to the May-Zhang et al. data as well, where ENC7 cluster is not a Met+ Ret- cell cluster. These data strongly show that Met-expressing neurons that do not express Ret (as our data and data from Avetisyan show make up a significant proportion of the population) are MENs and not NENs. The reviewer has incorrectly interpreted these datasets, possibly due to the low detectable expression and in general sparse expression profile of Met gene in the ENC7 (Morarach et al.) and ENT8 (Zeisel et al.) cell cluster.

B) Why were the Baf53b-Cre mice analysed in the colon, while the rest of the study was performed in the small intestine? It is possible that fewer neurons express Tomato in this region but the data does not match the general focus on small intestine in all other parts of the study. It is also not clear which stage was analysed, as the rebuttal letter states 6 weeks old mice, while the manuscript presents that the data is captured from P21 mice. Which one is correct? Moreover, the discussion claims that the initial analysis of Baf53bCre-Tom mice in the small intestine of P21 mice (Morarach et al) selectively included only neurons with cytoplasmic/nuclear HuC/D expression and not nuclear-only HuC/D neurons, while this was not the case for the analysis in question, but for adult ENS tissue in the same manuscript (pers. communication). Moreover, the limitation was for whole ganglia (not individual cells) that looked overall unhealthy due to large proportion of neurons only exhibiting nuclear localization and no cytoplasmatic HuC/D, features previously correlated to stressed neurons as discussed during the review-process. The manuscript discussion thus appears unbalanced and incorrect. The hostile remarks and comparisons should be corrected and possibly encompass possible explanations to the different results (colonic versus small intestine tissue, antibodies for HuC/D different, possible age difference). However, your actual data-analysis stands valid, and it possible that Baf53b-Cre mice do not capture the full ENS.

We are glad that the reviewer agrees with our data that the BAC transgenic Baf53b-cre:tdTomato transgenic mouse does not label all post-natal myenteric neurons. In addition, we direct the reviewer to Supplementary Figure 6f, where we have already shown the quantification of tdTomato-expressing neurons both in the small intestine and in the colon of this transgenic mouse line, which shows that this transgenic line does not label all myenteric neurons in large segments of the postnatal gut. We regret our error in the earlier rebuttal letter and confirm that these data were generated at the P21 age, which is the same age at which Morarach et al. used tissues from this transgenic mouse line for generating their scRNAseq data.

In addition, we respect the significant and groundbreaking work and data that has been generated by our colleagues in other labs (including the Marklund Lab) and thus, we have been careful not to negatively critique them. In prior reviews, reviewers asked us to compare and contrast why data from these laboratories were not able to detect MENs and we have complied with the reviewer's requests by listing the potential reasons for these discrepancies. However, we have further amended our Discussion (Lines 700 - 735) to ensure that we discuss these findings in a more collegial tone.

Minor points:
1) Add catalogue numbers of all antibodies used.

We thank the reviewer for bringing up this issue and have added the catalog numbers of all antibodies used.

2) Abstract first sentence: "The enteric nervous system (ENS), a collection of neurons contained in the wall of the gut" – too simplified – instead of "neurons" you could state "neural cells" (which encompasses both neurons and glia cells).

We thank the reviewer for this suggestion and have suitably revised the abstract.

Reviewer #2:
Remarks to the Author:
This re-revised manuscript from Kulkarni and colleagues is substantially improved from its initial submission and from the last revision. The addition of the concerted attempt to reconcile how the authors' data does or does not fit with existing scSEQ datasets in the field and to probe the causes of differences is helpful, strengthening the manuscript. The concept of a second developmental source of enteric neurons is an interesting one and is important for the field to consider, particularly with respect to aging. As with any interesting study, there are many questions left to answer, and further experiments with inducible lineage tracing, etc., will be needed to probe these observations further.

We are glad that the reviewer agrees that our data on the alternate developmental lineage of enteric neurons and their implications in age-associated diseases of gut physiology is important and we also thank the reviewer for highlighting the significant improvements made to the manuscript since the initial submission and through the three rounds of review. We are thankful that the reviewer agrees with us that this initial study opens the doors to further experiments and studies that will probe these observations further.

Only two concerns remain for me, 1 major and 1 minor.

1 – Major: This was raised in previous comments, but not addressed in revision. With respect to outcomes that are by nature binary (neurons can only be tdTomato positive or negative), it would be most appropriate to depict data showing mean and SEM only for % positive or % negative. Otherwise, it appears as if two different things were measured (For example, bar graph in Fig. 1d could be pie chart, split one column, or text alone rather. Two separate bars indicates two separate measurements but these data represent only one set of measurements). This is relevant to Fig. 1d, 1f, 3e-i. In Fig. 3, the two groups in each graph are even statistically compared even though each data point in the right column is always going to be 100 minus data point in the left column. A more appropriate test would be to compare how the % positive per animal

differs from expected theoretical average – ie. 100% if expecting that if all NOS neurons are derived from Wnt1+ population.

We thank the Reviewer for bringing up this issue and for providing suggestions on a better way for statistically analyzing these effects. While pie charts would not allow us to show individual data points and error bars, we have depicted data in Fig 1d, 2f, 3e-i as staggered bar graphs. Further, we have amended the statistical test used to Fisher's exact test to compare how proportions observed deviate from the expected theoretical average (where all neurons are expected to be derived from the Wnt1-cre-expressing neural crest).

2 – Line 161, Fig. 2h and Supp Fig 3a show the same cells, just at different magnifications. This should be clarified, redundancy eliminated, or a second example shown.

We thank the Reviewer for highlighting this issue. Figure 2h and Supplementary Figure 3a indeed show the same myenteric ganglion, as we wanted the reviewer and readers to appreciate the MHCst immunostaining in the myenteric ganglia with and without the red immunofluorescent signal from the neural crest-derived cells in the Wnt1-cre:tdTomato mouse line. However, we agree that this should have been better clarified and have amended our text in Lines 164 - 166 and in the figure legend of Supplementary Figure 3a (lines 1627 - 1633) to highlight that Supplementary Figure 3a and Figure 2h show the same myenteric ganglion.
